# Red bone marrow hosts metabolically active anucleate adipocytes that support hematopoiesis

**DOI:** 10.64898/2025.12.02.691590

**Authors:** Sauyeun Shin, Marine Hernandez, Stéphanie Dauvillier, Manuelle Ducoux-Petit, Emmanuelle Mouton-Barbosa, David Estève, Mohamed Moutahir, David Rengel, Océane Delos, Mathieu Roumiguié, Cédric Dray, Philippe Valet, Philippe De Médina, Marc Poirot, Sandrine Silvente-Poirot, Justine Bertrand-Michel, Odile Burlet-Schiltz, Nicolas Reina, Catherine Muller, Camille Attané

## Abstract

Bone marrow adipocytes (BMAds) are a major component of the bone marrow (BM) that regulate bone turnover and hematopoiesis. In rodents, two distinct adipocyte populations exist: constitutive BMAds (cBMAds), located in areas devoid of hematopoietic cells and resistant to metabolic cues, and regulatory BMAds (rBMAds), interspersed within hematopoietic niches and responsive to energy stress. Despite their potential importance, rBMAds have remained poorly characterized due to their scarcity in rodents. Here, we used a high-yield method to isolate human rBMAds, enabling structural, proteomic, lipidomic, and functional analyses. Remarkably, human rBMAds are anucleate yet retain organelles and maintain lipid and glucose metabolism, but unlike their rodent counterparts, they lack lipolytic activity. They actively secrete factors that support hematopoiesis *in vitro*, implicating them as functional contributors to the BM niche. These findings suggest that human rBMAds may arise from cBMAds through terminal differentiation and define a novel adipocyte subtype critical for BM homeostasis.

## Introduction

Bone marrow adipose tissue (BMAT), made up of bone marrow adipocytes (BMAds), is a major yet still incompletely characterized component of the bone marrow (BM)^1,2^. In both humans and rodents, BMAT displays structural and metabolic features that distinguish it from classical adipose tissues (AT) such as subcutaneous (SCAT) and visceral AT^1,2^. BMAT is unusual in forming two distinct AT types : “yellow” BMAT, containing densely packed adipocytes devoid of hematopoietic cells, that resembles other AT depots, and “red” BMAT, composed of adipocytes dispersed among hematopoietic cells, suggesting specialized context-dependent functions^3,4^. These regions occupy distinct anatomical sites within the skeleton, with transitional zones in between^1,3,4^. The preservation of BMAT during caloric restriction, and even its expansion under long-term energy deprivation, such as in anorexia nervosa, in contrast to all other AT depots, underscores its unique metabolic properties^1,2^. Evidence from histopathology and imaging indicates that BMAT regulates local bone turnover and hematopoiesis^1,2^. BMAT volume increases with age in rodents and humans, correlating with reduced hematopoietic activity and decreased bone mineral density, which can contribute to osteoporosis and fractures^1,5–8^. In rodents, BMAd density correlates negatively with hematopoietic stem cell function^9,10^. Paradoxically, BMAds are a major source of stem cell factor (SCF) after irradiation and are essential for hematopoietic recovery^11^, suggesting context-dependent functions that may differ between basal and acute hematopoiesis or among BMAd subpopulations. These observations underscore the need for deeper characterization of BMAds, particularly functional assessments of distinct subtypes. In rodents, BMAds located in hematopoietic-rich areas undergo dynamic remodeling and decrease in size in response to cold exposure, acute fasting, exercise, or hemolysis, leading to the concept of regulatory BMAds (rBMAds)^12–15^. By contrast, BMAds in areas devoid of hematopoiesis remain unaltered under such stressors and are termed constitutive BMAds (cBMAds)^14,15^. To date, lipolysis, a process involving catecholamine-mediated stimulation of triglyceride (TG) breakdown via sequential activation of three lipases ^16^, has been considered as a key metabolic feature distinguishing cBMAds from rBMAds^1^. cBMAds resist lipolysis and remodeling in response to β-adrenergic stimulation in mice, consistent with their preservation during caloric restriction^15^. We confirmed these results in humans using isolated human cBMAds obtained from aspirates of the proximal femoral metaphysis and diaphysis of patients undergoing hip replacement surgery^17,18^. Through mass-spectrometry (MS)-based proteomic analyses, we demonstrate that this phenotype is driven by profound downregulation of monoacylglycerol lipase (MGLL), the final enzyme of lipolysis, despite intact adrenergic signaling^17^. In rodents, rBMAds respond to β-adrenergic stimulation, although their glycerol release (the end product of lipolysis) remains modest compared to SCAT adipocytes (SCAds)^15^. Consistently, mice with BMAd-specific deletion of ATGL (Adipose triglyceride lipase), the first lipase of lipolysis, display enlarged rBMAds and mildly impaired forskolin-induced lipolysis^19^. However, whether rBMAds exhibit similar lipolytic behavior in humans remains unknown. Moreover, *in vivo* or *ex vivo* experiments in both rodents and humans have revealed differences in metabolic^20–23^ and endocrine^24,25^ functions between rBMAT and cBMAT, although these analyses performed on whole tissue limit the ability to distinguish between adipocyte subtypes.

Despite substantial progress in defining BMAd heterogeneity, an in-depth characterization of rBMAds has never been achieved, leaving fundamental questions unresolved regarding their cellular organization and function. Because rBMAds reside within active BM niches, they are the BMAd subtype most likely to influence hematopoietic cell function. However, their fragility and low abundance, especially in rodents, have hindered their isolation at yields sufficient for molecular and functional characterization. In the present study, we overcome this longstanding barrier by leveraging our recently developed protocol to obtain high-yield aspirates from the femoral metaphysis and diaphysis of patients undergoing hip surgery^18^, enabling, for the first time, the isolation of human rBMAds in quantities suitable for comprehensive structural, proteomic, lipidomic, and functional analyses. We show that human rBMAds lack a nucleus but retain other organelles, and although they display no lipolytic activity, they maintain key pathways of lipid and glucose metabolism. Moreover, rBMAds remain actively secretory and support normal hematopoiesis *in vitro*.

Together, our study provides the first functional and molecular characterization of isolated human rBMAds, identifying them as a distinct adipocyte population and redefining their role within the BM niche.

## Results

### Distinct structural and cellular features of rBMAT compared to cBMAT and SCAT

To characterize human rBMAds, BMAT was aspirated from the femoral proximal metaphysis and diaphysis of patients undergoing hip surgery, as previously described^18^. As we and others reported^1,7,18^, aspirates obtained contain interlaced regulatory (“red”) and constitutive (“yellow”) BMAT and the content of macroscopically-defined rBMAT and cBMAT areas varies between patients with overall higher amount of rBMAT relative to cBMAT (Figure S1A). Post-mortem slices of femur confirmed the presence of both BMATs within the metaphysis and diaphysis of patients, while the distal epiphysis and femoral head were enriched in cBMAT (Figure S1B). BM aspirates were dissected to isolate rBMAT and cBMAT and analyzed for lipid content and hematoxylin and eosin (HE) staining alongside paired SCAT. SCAT and cBMAT contained densely packed adipocytes with comparable lipid content, whereas rBMAT showed smaller and scattered adipocytes surrounded by numerous hematopoietic cells as well as reduced total lipid content, reflecting a higher proportion of non-adipocyte cells (Figure 1A, S1C). Post-mortem femur biopsies showed similar architectural differences between cBMAT and rBMAT regions (Figure S1D). HE staining also revealed the presence of transitional zones between rBMAT and cBMAT (Figure 1A) as previously reported in rodents^1^.

**Figure 1.**
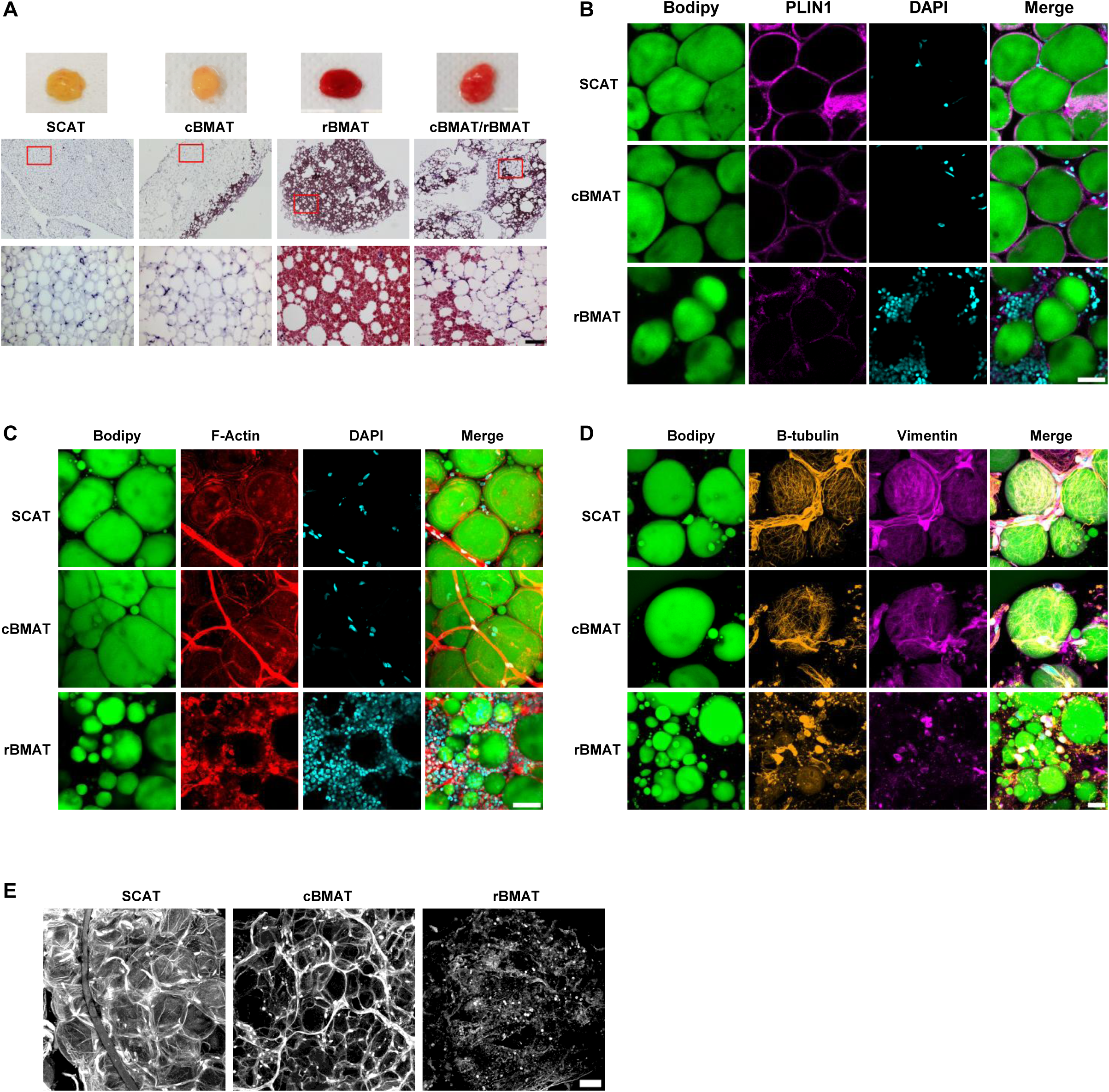
Distinct structural and cellular features of rBMAT compared to cBMAT and SCAT. **(A)** HE staining of SCAT, cBMAT, rBMAT and mixed cBMAT/rBMAT; Scale bar, 100µm. **(B-E)** Whole ATs were stained with Bodipy 493/503 for neutral lipids, DAPI for nuclei and Perilipin 1 (PLIN1); scale bar, 50µm (B) or Phalloidin for F-Actin; scale bar, 50µm (C) or Beta tubulin and Vimentin; scale bar, 20µm (D) or with picrosirius red to show collagen fibers, scale bar, 50µm (E). Representative maximum intensity projection of z-stack images is shown.

BMAT from both locations contained adipocytes with a unilocular lipid droplet (LD) filled with neutral lipids (as assessed by Bodipy staining) and expressing perilipin-1 (PLIN1), a protein localized to the surface of LDs^26^, as found in control SCAds (Figure 1B). Actin staining revealed blood vessels in SCAT and cBMAT, and numerous hematopoietic cells around adipocytes in rBMAT, while showing weaker staining intensity in rBMAT adipocytes compared with SCAT and cBMAT (Figure 1C). Staining for additional cytoskeletal component, including microtubules (β-tubulin) and intermediate filaments (vimentin)^27^ showed again weaker signals in rBMAT adipocytes compared with those in SCAT or cBMAT (Figure 1D). The lower expression of these proteins that contribute to contractile force generation and stiffness in response to mechanical stress^28^, might reflect a looser extracellular matrix organization in rBMAT, where adipocytes are less cohesive. Accordingly, the collagen network was substantially less dense and had thinner fibers in rBMAT compared to cBMAT and SCAT (Figure 1E). Taken together, these results validate the tissue samples obtained, which reflect the tissular and cellular organization of cBMAT and rBMAT and demonstrate that rBMAds display lower expression of cytoskeletal proteins compared to SCAds or cBMAds.

### rBMAds are devoid of nucleus but retain metabolic organelles

Adipocytes were then isolated as previously described^18^, yielding pure cells that retained the tissue morphology with a unique LD (Figure 2A). SCAds and cBMAds were similar in size (mean 90□µm) as we previously reported^17^, while rBMAds were smaller (mean 60□µm) with a heterogeneous distribution (Figure 2B). Nevertheless, all three adipocyte types contained comparable total lipid amounts per suspension volume (Figure 2C). Surprisingly, we did not detect signal with the commonly used fluorescent DNA stain Hoescht 33258 (Figure 2A) or with DAPI, another nuclear stain (Figure S2A) in rBMAds while a positive staining was observed in SCAds and cBMAds using both dyes. To confirm the absence of nuclei in rBMAds, we measured DNA and RNA content and found them barely detectable in rBMAds compared to SCAds and cBMAds (Figures 2D–E). Using transmission electron microscopy (TEM) on the three ATs, we observed that both SCAds and cBMAds display a large LD surrounded by a thin cytoplasm and nucleus located at the cell periphery whereas rBMAds exhibited a thinner cytoplasm surrounding a large LD, with no nucleus observed in any sample (Figure 2F).

**Figure 2.**
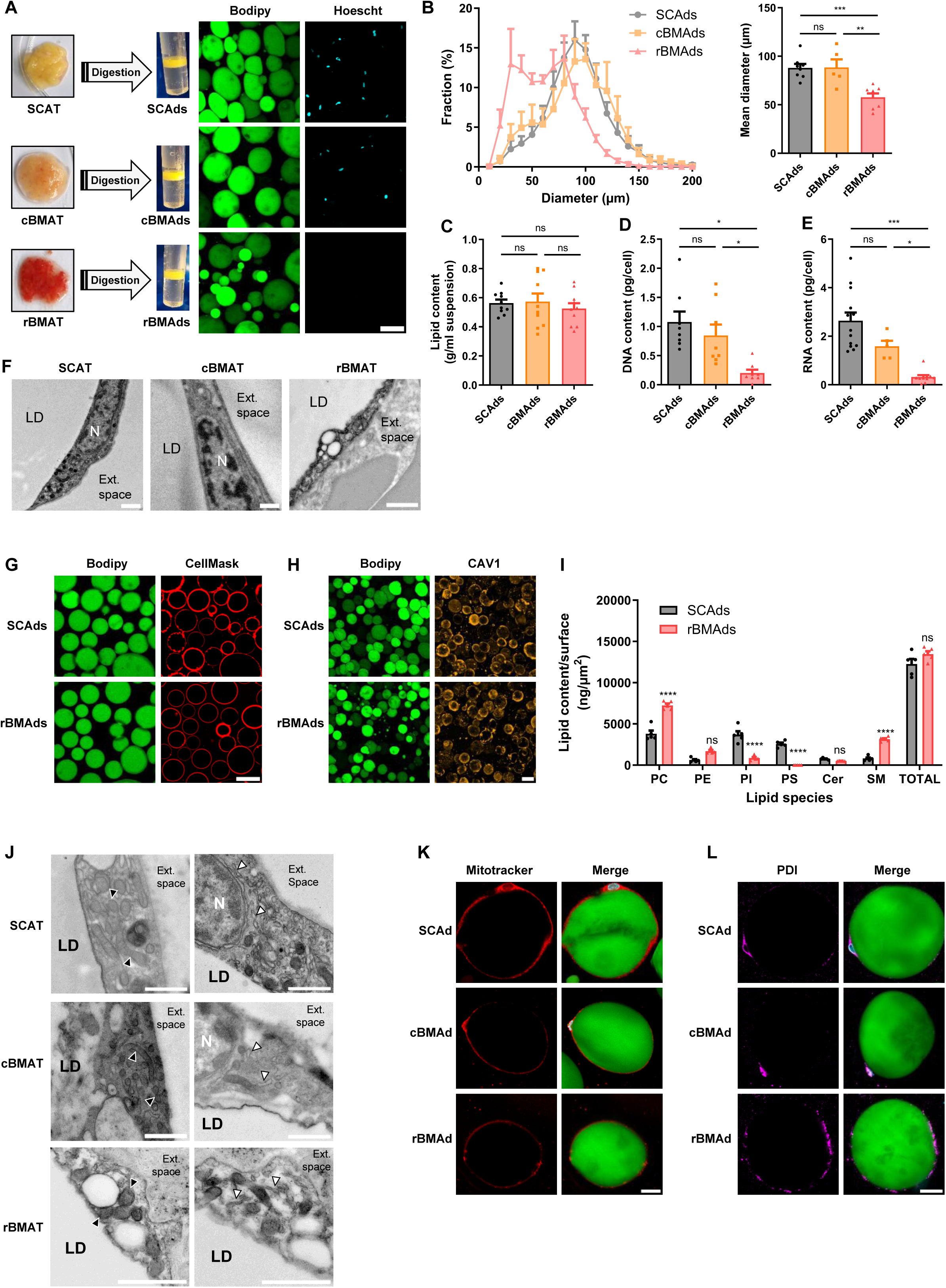
rBMAds are devoid of nucleus but retain metabolic organelles. **(A)** Primary adipocytes were isolated and stained with Bodipy 493/503 and Hoescht 33258. Representative maximum intensity projection of z-stack images is shown. Scale bar, 100µm. **(B)** Size distribution (left panel) and mean diameter (right panel) of isolated adipocytes (n=5-8). **(C)** Total lipid content in paired isolated adipocytes (n=10). **(D)** DNA content in paired isolated adipocytes (n=8). **(E)** RNA content in isolated adipocytes (n=5-13). **(F)** TEM images of ATs. N: Nucleus, LD: Lipid Droplet, Ext. space: extracellular space. **(G-H)** CellMask (G) or Caveolin (CAV1) (H) stainings in isolated adipocytes. Representative maximum intensity projection of z-stack images is shown. Scale bar, 100µm. **(I)** Quantification of membrane lipids: Phosphatidylcholine (PC), Phosphatidylethanolamine (PE), Phophatidylinositol (PI), Phosphatidylserine (PS), Ceramides (Cer) and Sphingomyelin (SM) (n=5 paired samples). **(J)** TEM images of SCAT, cBMAT and rBMAT. N: Nucleus, LD: Lipid Droplet, Ext. Space: extracellular space. Black arrowheads show mitochondria and white arrowheads show endoplasmic reticulum. **(K-L)** Mitotracker (K) or Protein disulfide-isomerase (PDI) (L) stainings in isolated adipocytes. Representative maximum intensity projection of z-stack images is shown. Scale bar, 20µm. The data are the mean ± SEM. ns: non-significant; *p<0.05; **p<0.01; ***p<0.001; **** p<0.0001 according to One-way ANOVA followed by Bonferroni post-test (B, C, D and E) or two-way ANOVA followed by Bonferroni post-test (I).

We confirmed the presence of a plasma membrane in rBMAds using CellMask™ (Figure 2G), which incorporates into the phospholipid bilayer^29^, and caveolin-1 (CAV1) (Figure 2H), a protein essential for caveolae formation in adipocyte membranes^30^. Total plasma membrane lipids (phospholipids and sphingolipids) were similar between SCAds and rBMAds in five paired samples, although their composition differed: phosphatidylcholine (PC) and sphingomyelin (SM) were increased, while phosphatidylinositol (PI) and phosphatidylserine (PS) were decreased in rBMAds (Figure 2I). PC and SM are major components of the outer leaflet of the plasma membrane associated with membrane rigidity and stability and their increase is highly susceptible to reduce fluidity^31^. On the contrary, both PI and PS are located in the inner leaflet and their decrease could contribute to reduced signaling capacity^31^. Both types of adipocytes contained organelles with the typical morphology of mitochondria, as shown by TEM of ATs (Figure 2J, left panel) and these findings were confirmed using MitoTracker (Figure 1K). rBMAds, like other adipocytes, also exhibited structures resembling endoplasmic reticulum (ER) in TEM (Figure 2J, right panel). Expression of protein disulfide isomerase (PDI), a key ER component^32^ was detected in adipocytes from all tissues, although with distinct patterns. In SCAds and cBMAds, the signal was concentrated around the nucleus, while in rBMAds it was diffusely present throughout the cytoplasm (Figure 2L). These findings suggest that SCAds and cBMAds predominantly express rough ER (RER), which is continuous with the nuclear envelope and facilitates mRNA transfer from the nucleus to ribosomes on the RER for protein synthesis^33^. In contrast, rBMAds exhibit structures resembling smooth ER (SER), which is distributed throughout the cytoplasm^33^. Thus, despite lacking a nucleus, rBMAds retain key metabolic organelles such as mitochondria and ER and their distinct cellular organization underscores their unique identity among adipocyte subtypes and suggests specialized functional roles. In humans, cells within hematopoietic tissue, such as erythrocytes, are also anucleate; however, these cells lack organelles entirely. Platelets, which are derived from megakaryocytes, are another example of anucleate elements, but they too differ structurally from rBMAds. Although their cellular architectures are markedly distinct, these analogies with terminally differentiated states hematopoietic derivatives raise the hypothesis that rBMAds might also represent terminally differentiated cells.

### Lipidomic and proteomic characterization define rBMAds identity in humans

Previous studies have shown that BMAd lipid composition differs by subtype and location. In rodents, rBMAd are enriched in saturated fatty acids (SFAs) and depleted in monounsaturated fatty acids (MUFAs) compared to cBMAds resulting in a lower unsaturation index^14^. Similar trends have been reported in humans using ¹H-MRS^14^ and GC-MS on whole BMAT, although the extent of these differences varies across studies depending of anatomical site studied^34^ and context (post-chemotherapy samples)^35^. To investigate potential differences in lipid content and saturation, we performed untargeted lipidomic on five paired samples. A total of 465 lipid species across 14 classes were detected. Consistent with prior findings in cBMAds^17^, rBMAds predominantly contained TG (>95%) and had minor levels of diglycerides (<2%) and other lipid species, including phospholipids and sphingolipids (Figure 3A). Principal component analysis of these data did not separate samples by adipocyte type or patient (Figure S3A). Consistently, the different lipid classes were not significantly different between the 3 types of adipocytes, although some trends were observed for membrane and signaling lipids that were decreased (ceramides and PI) or increased (SM) in rBMAds compared to the two other types of adipocytes (Figure 3B), in accordance with our characterization of membrane lipid composition (Figure 2I). FA profiling in five additional paired samples showed increased SFAs (palmitate, stearate) and decreased MUFAs (palmitoleate, oleate) in both BMAd types relative to SCAds (Figure 3C and S3B), leading to a lower unsaturation index in both BMAds (Figure 3D). Using paired purified adipocytes, these findings show that while BMAds exhibit a distinct FA profile and reduced unsaturation index relative to SCAds, no major differences distinguish rBMAds from cBMAds regarding their lipid composition.

**Figure 3.**
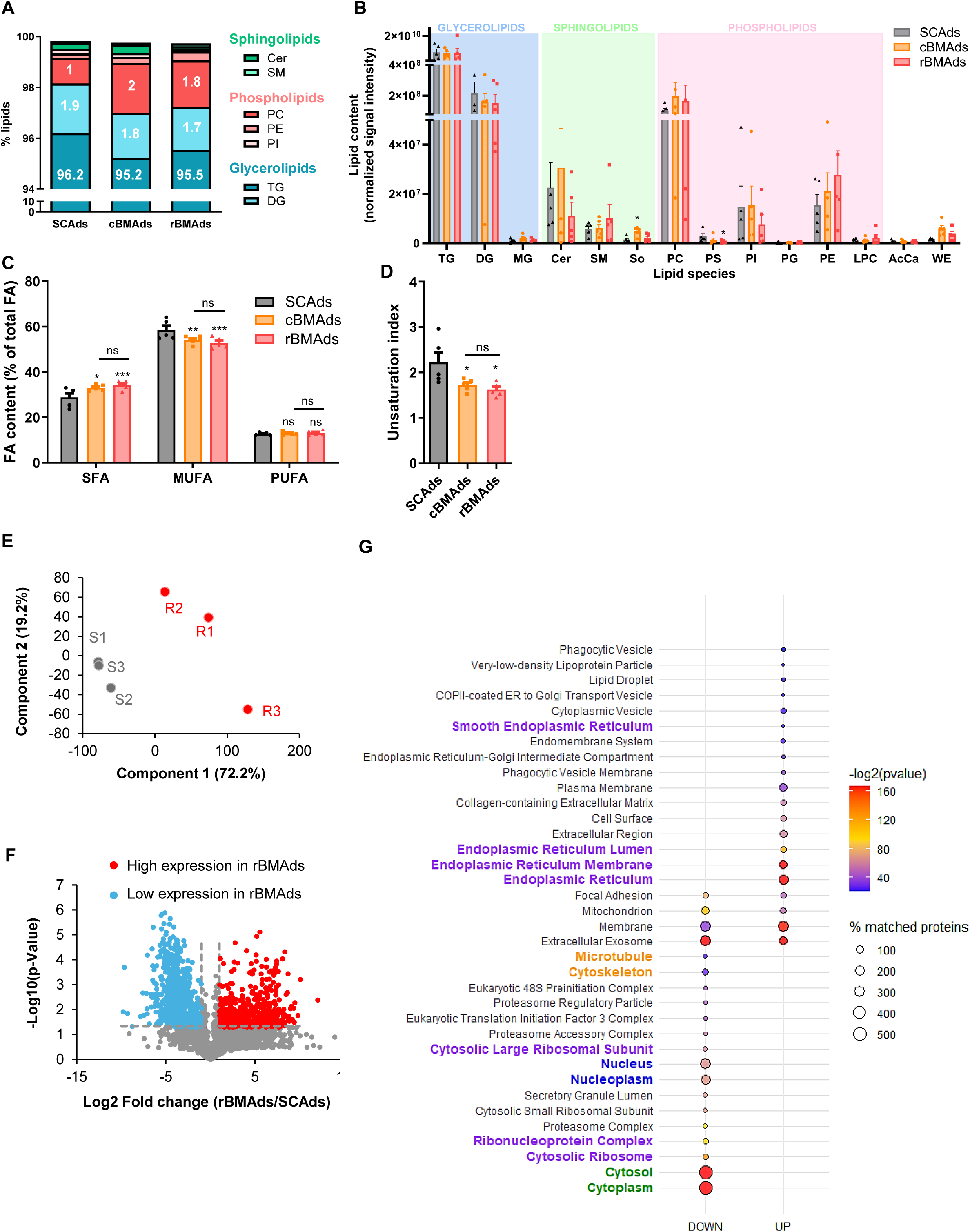
Lipidomic and proteomic characterization define rBMAds identity in humans. **(A-B)** Proportion (A) and relative abundance (B) of lipid classes detected by LC-MS/MS in paired isolated adipocytes (n=5). **(C-D)** Proportion of saturated FA (SFA), monounsaturated FA (MUFA) and polyunsaturated FA (PUFA) in paired isolated adipocytes (C) and unsaturation index (D) (n=5). The data are the mean ± SEM. Ns: non-significant; *p<0.05; **p<0.01; ***p<0.001 according to One-way ANOVA followed by Bonferroni post-test (D) or two-way ANOVA followed by Bonferroni post-test (B and C). **(E)** Principal component analysis performed on the proteomic dataset from paired rBMAds **(R)** and SCAds (S) samples (n=3). **(F)** Volcano plot representing the log2-transformed fold-change of rBMAds compared to SCAds in the proteomic dataset and the -log10 of the p-Value. **(G)** Pathway enrichment analysis of cellular component gene ontology terms performed with Gene Analytics. The top 20 pathways upregulated or downregulated in rBMAds compared to SCAds are presented.

To gain deeper insight into the specific features of rBMAds, we performed high-throughput MS-based proteomic analyses on paired rBMAds and SCAds. This approach was chosen given the lack of paired samples containing sufficient quantities of cBMAds and rBMAds for proteomic analyses. After quality control, 2673 proteins were detected across SCAds and rBMAds (Table 1). Principal component analysis of the proteomic dataset clearly partitioned the samples according to the adipocyte type with the first component explaining 72% of the total variance (Figure 3E). Differential expression analysis identified over 1 000 proteins significantly modified between SCAds and rBMAds, with 453 upregulated and 554 downregulated in rBMAds. (Figure 3F). Pathway enrichment analysis of cellular component (Figure 3G) and biological processes (Figure S3C) gene ontology terms confirmed the structural specificities of rBMAds that were initially observed through imaging, notably a marked decrease in proteins associated with the nucleus, cytoskeleton, and cytoplasm in rBMAds (Figure 3G). Furthermore, proteins associated with the SER were overrepresented, whereas those associated with the RER and translational processes were underrepresented in rBMAds relative to SCAds (Table 2, Figure 3G and S3C), again aligning with our imaging data (Figure 2L). Notably, a similar downregulation of translation-related pathways has been described during erythrocyte maturation, where enucleated erythrocytes exhibit a sharp reduction in translation compared to reticulocytes^36,37^. The overrepresentation of SER proteins in rBMAds raises the hypothesis that these cells may play a specific role in glycerolipid and phospholipid synthesis, suggesting that rBMAds exhibit unique membrane remodeling dynamics linked to the distinct phospholipid profile we observed (Figure 2I). Overall, proteomic analyses uncover a striking divergence between rBMAds and SCAds and confirmed the remodeling of intracellular architecture, including depletion of nuclear and cytoskeletal components and enrichment of SER-associated proteins, positioning rBMAds as a functionally distinct adipocyte population.

**Table 1:**
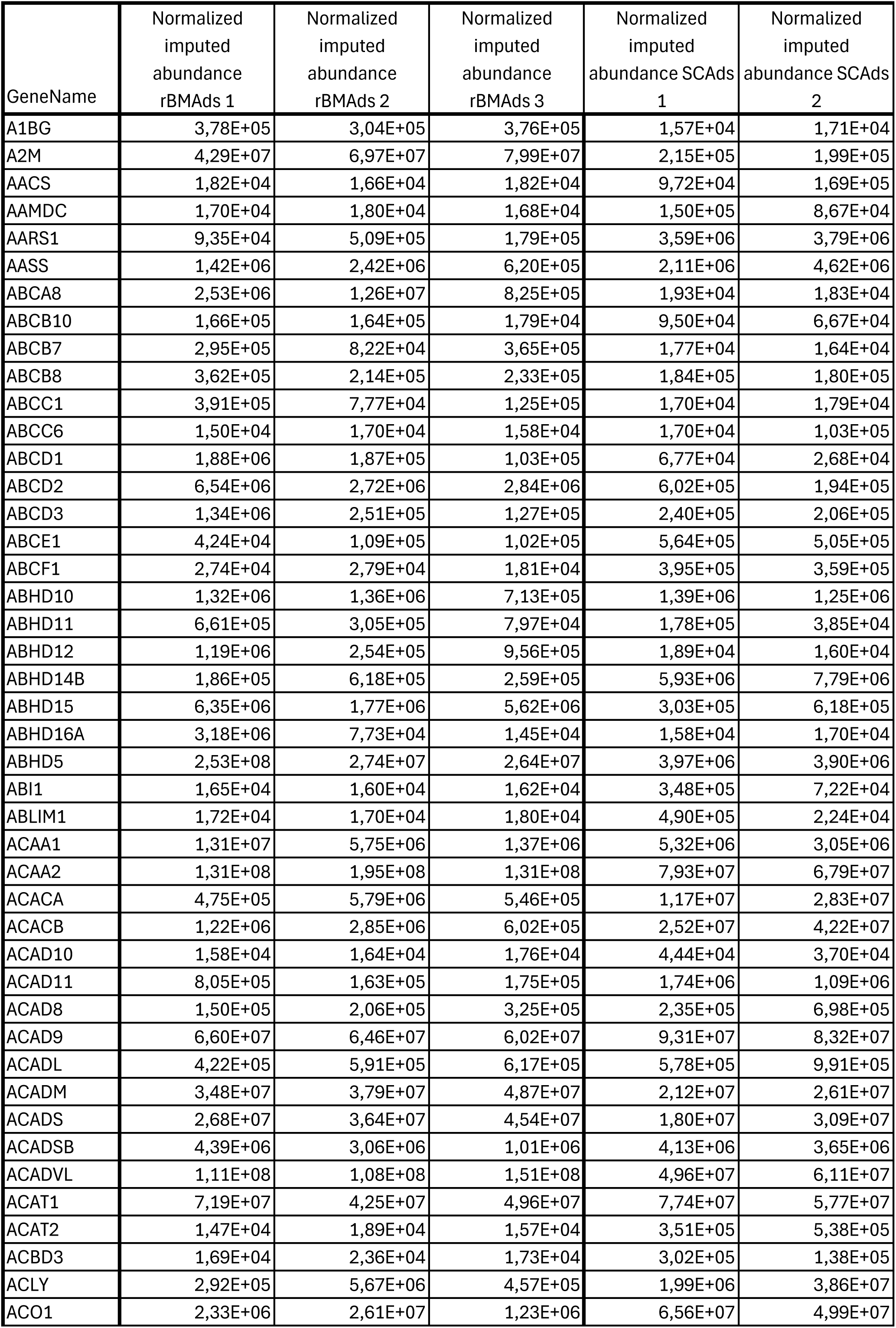
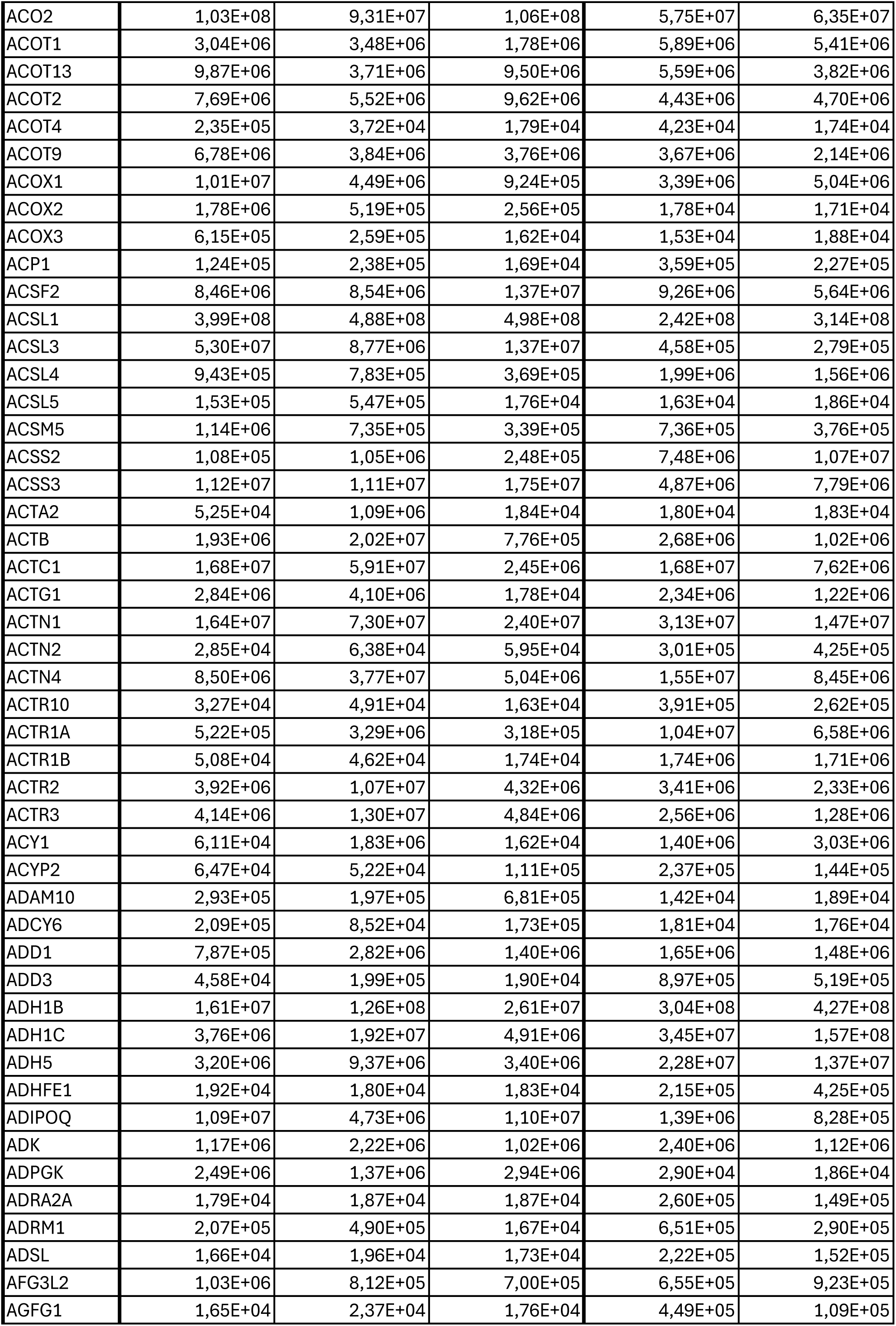
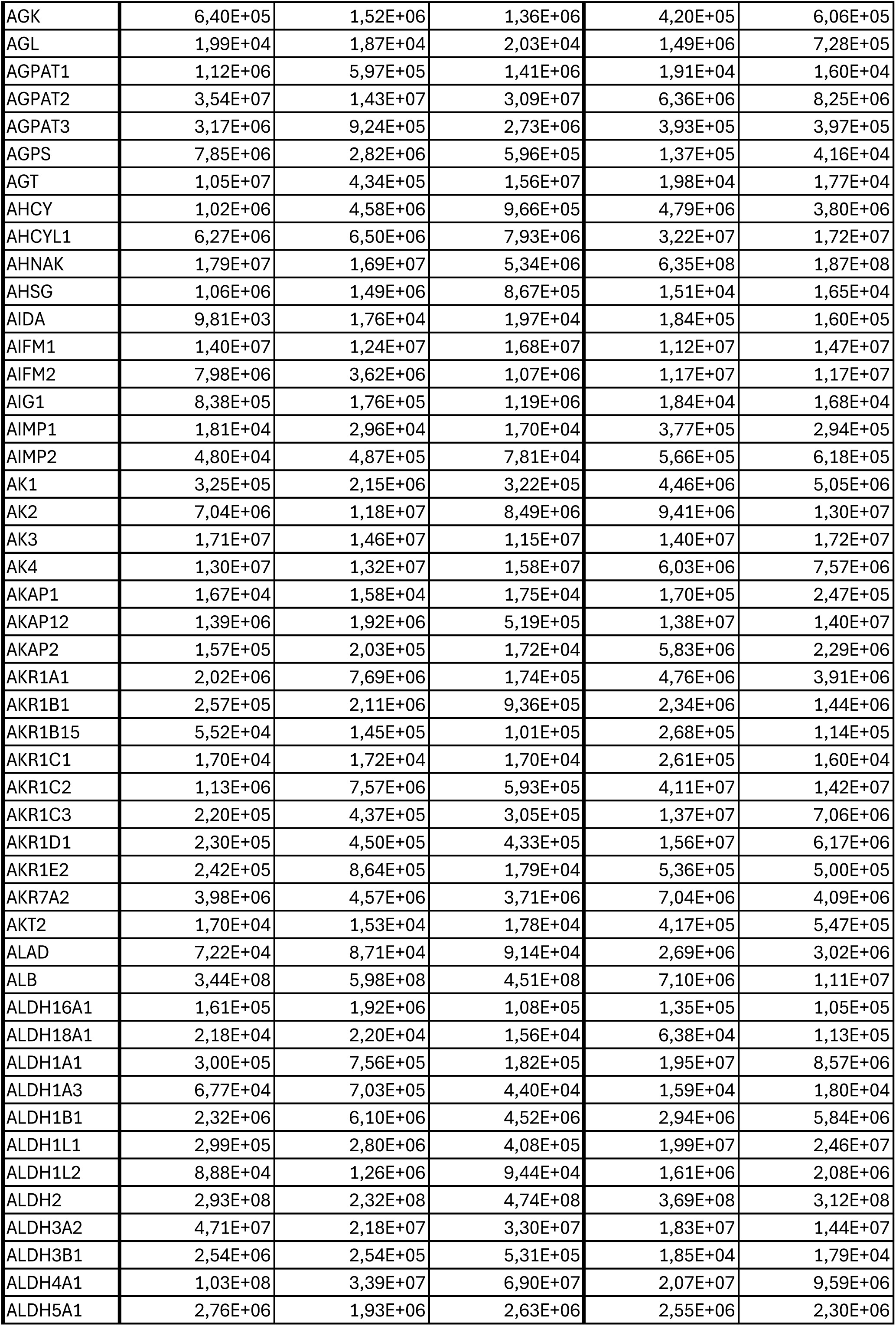
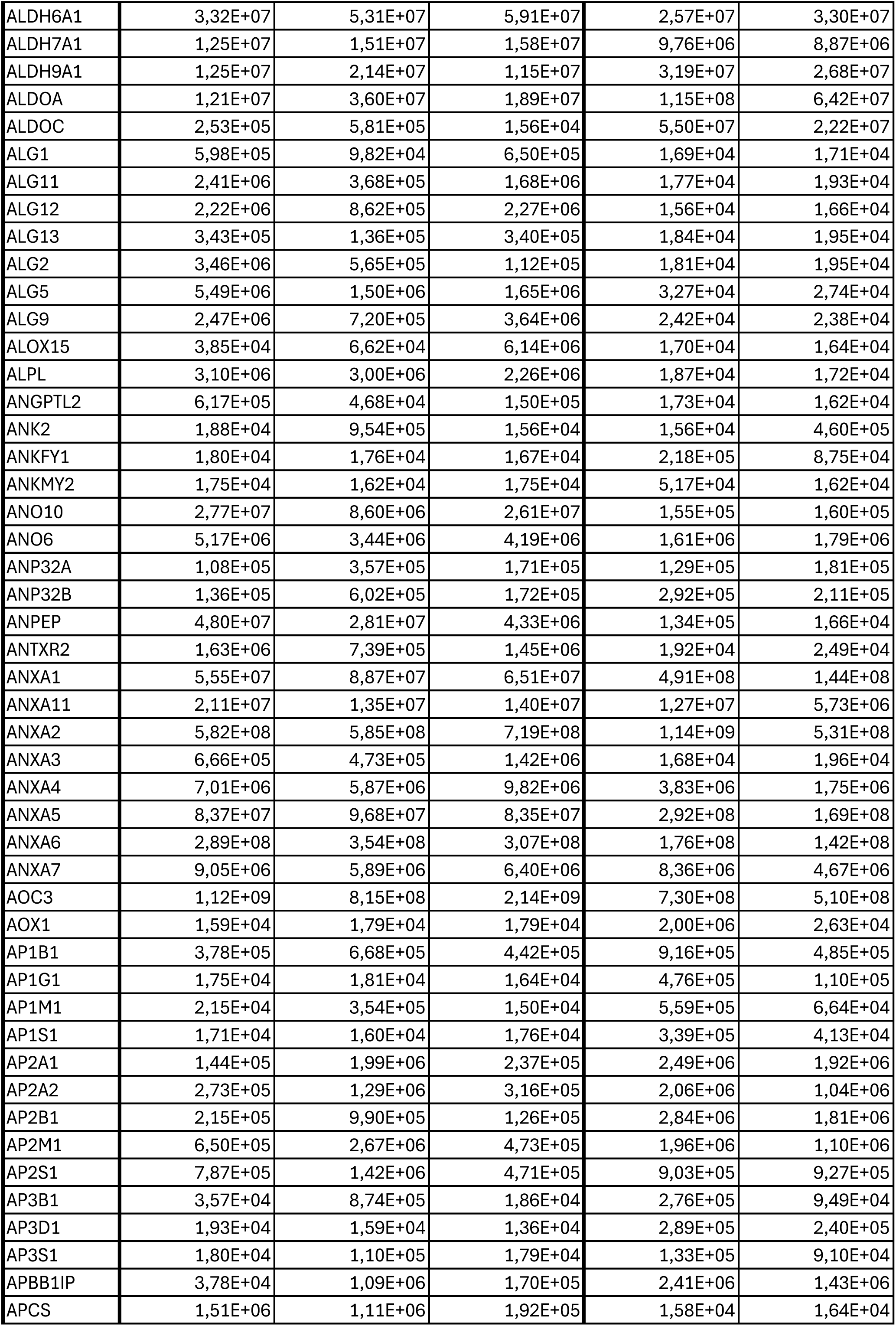
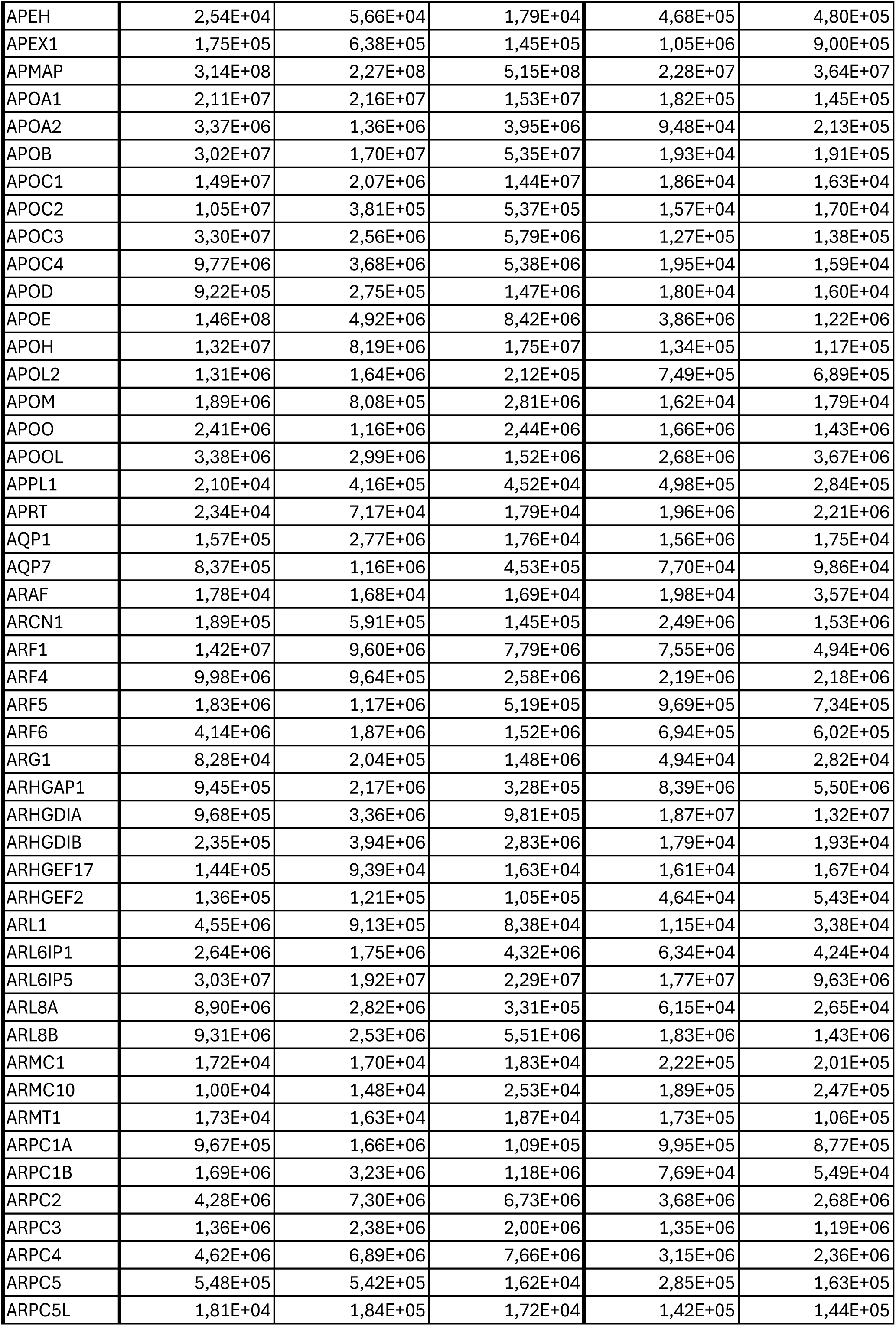
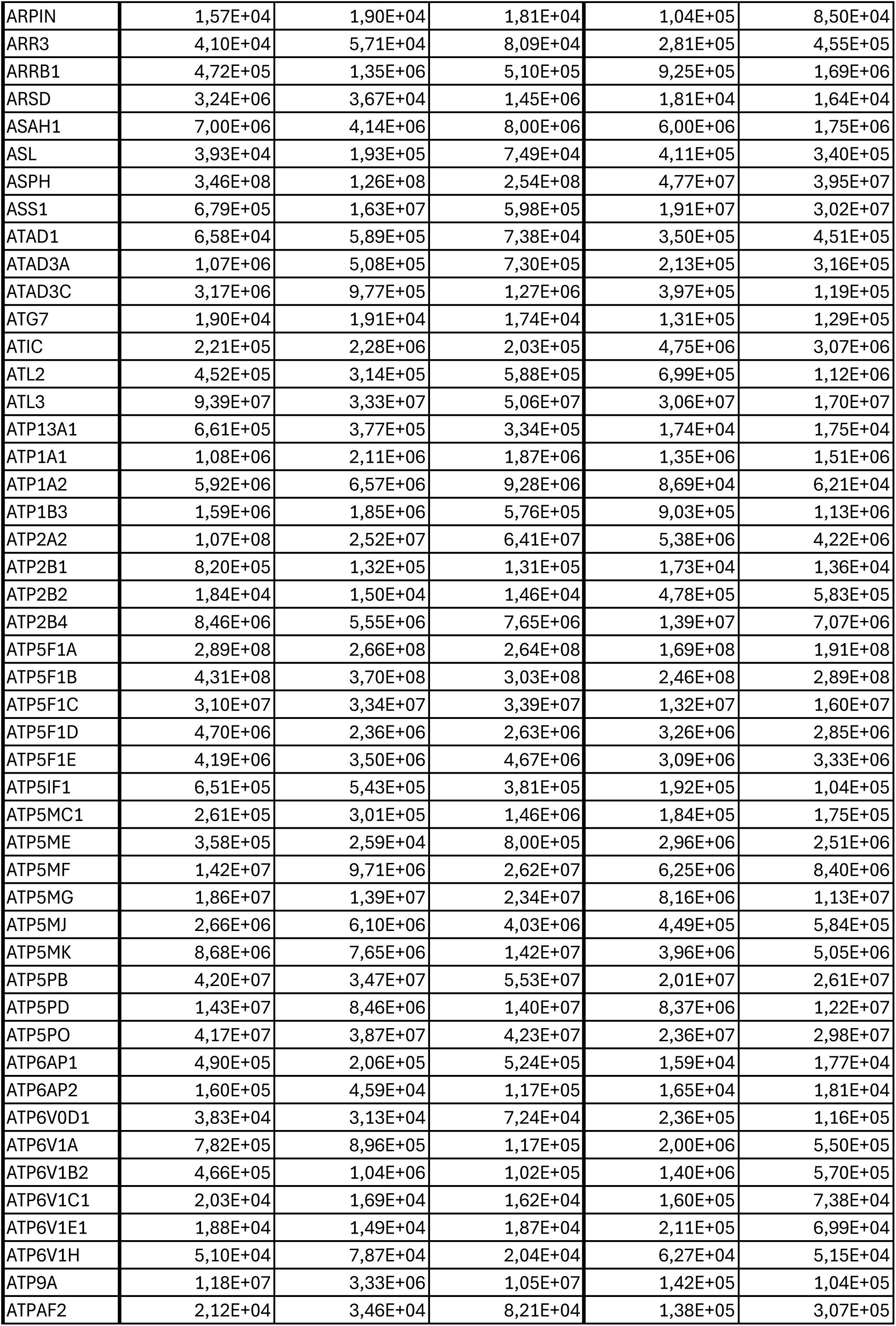
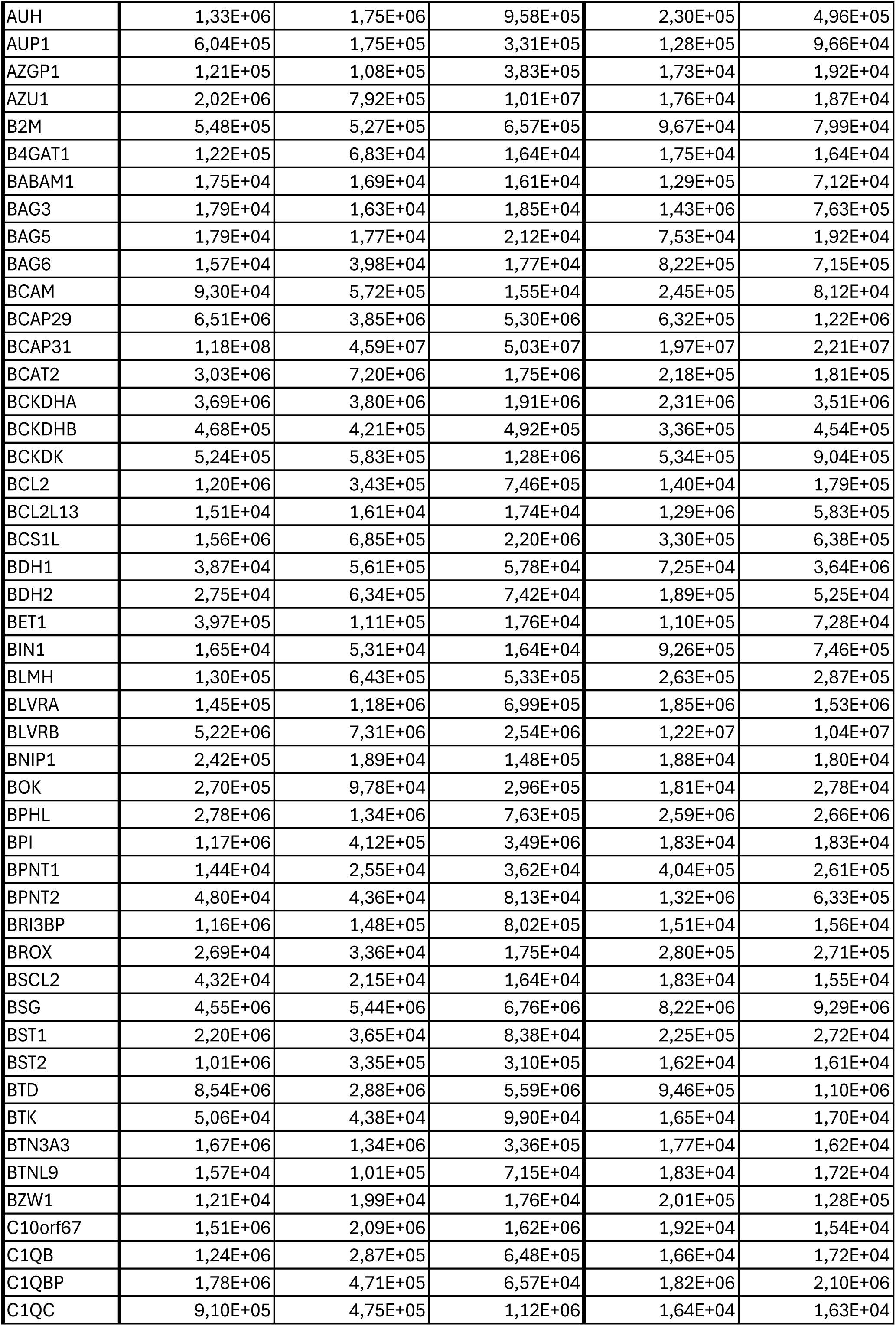
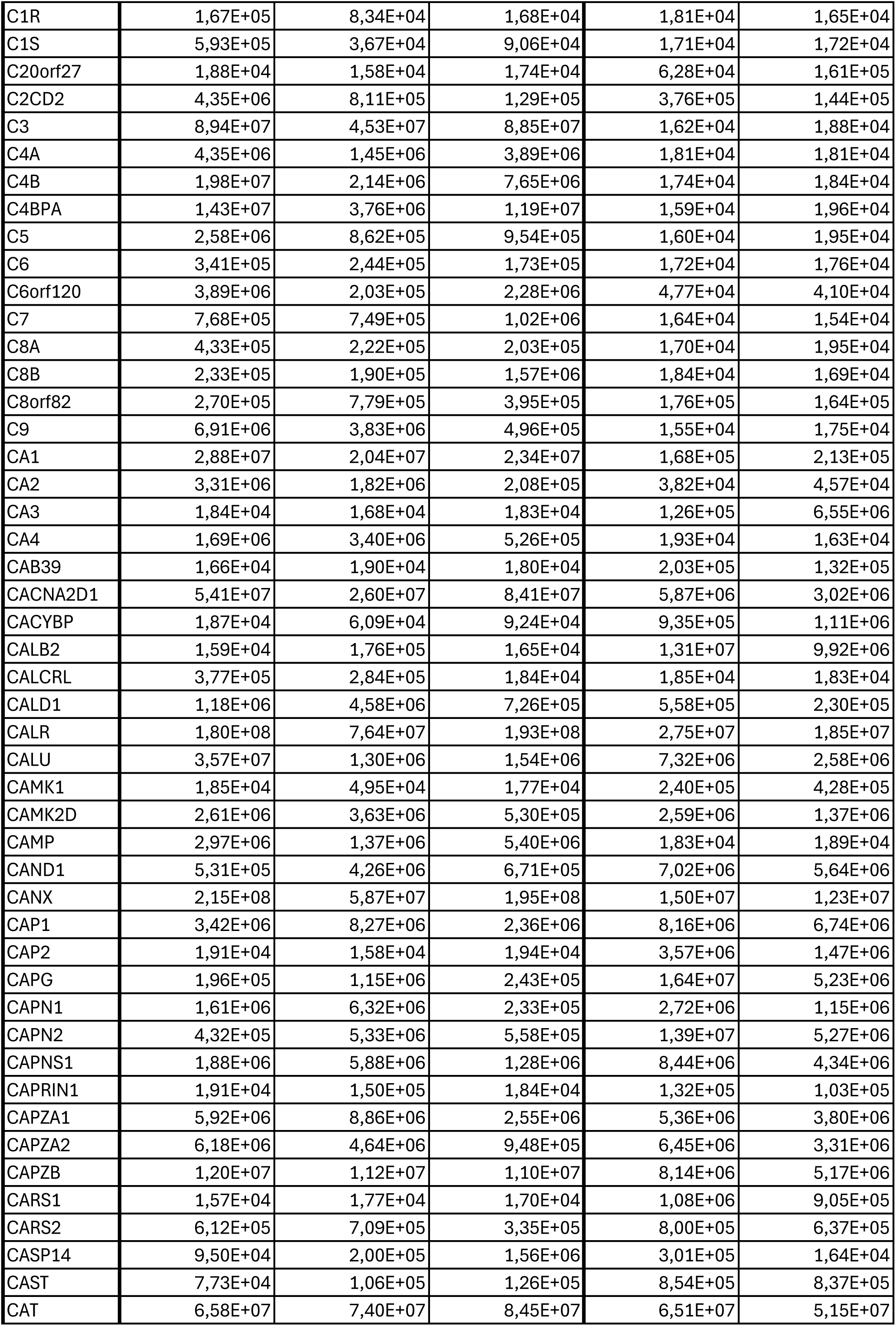
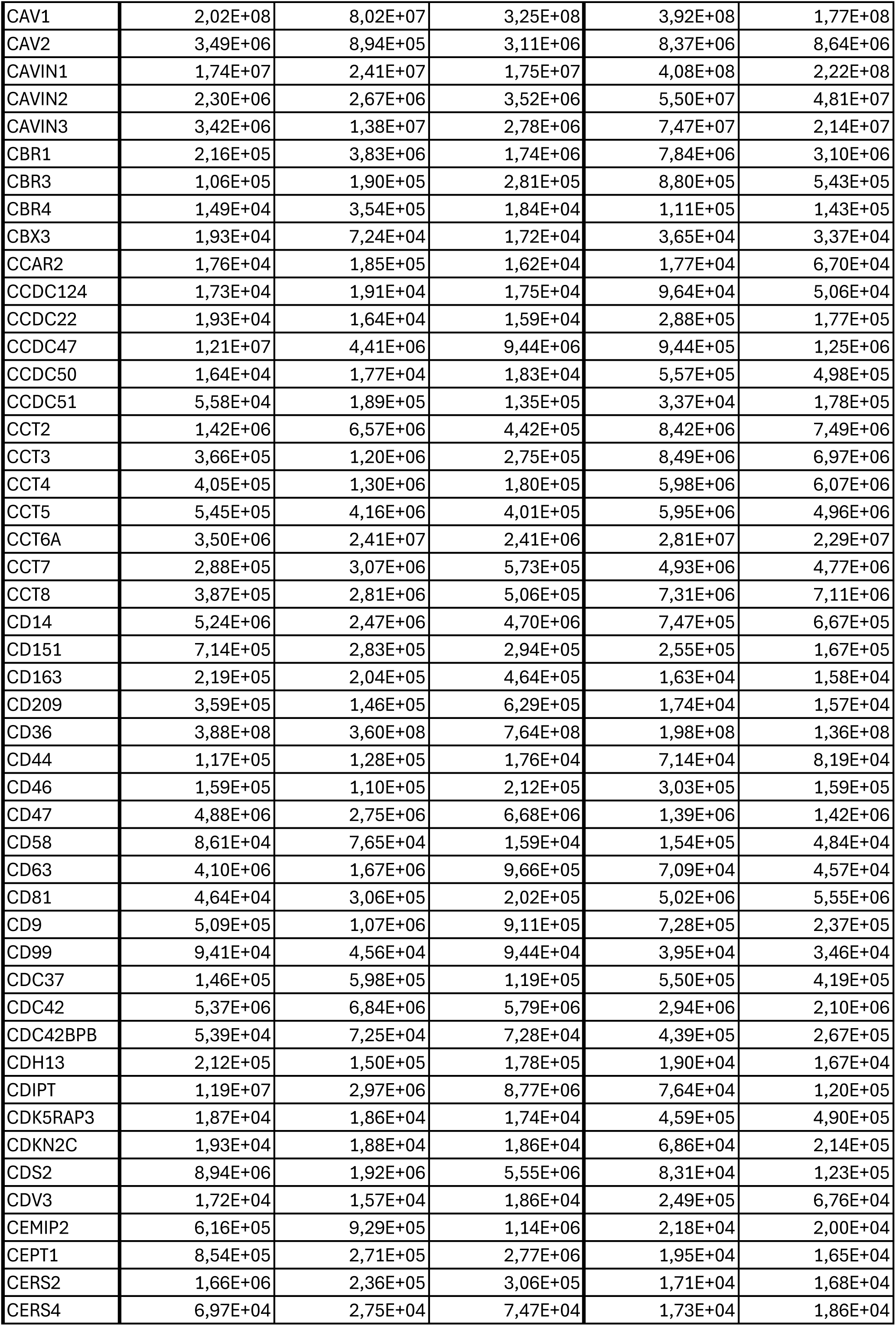
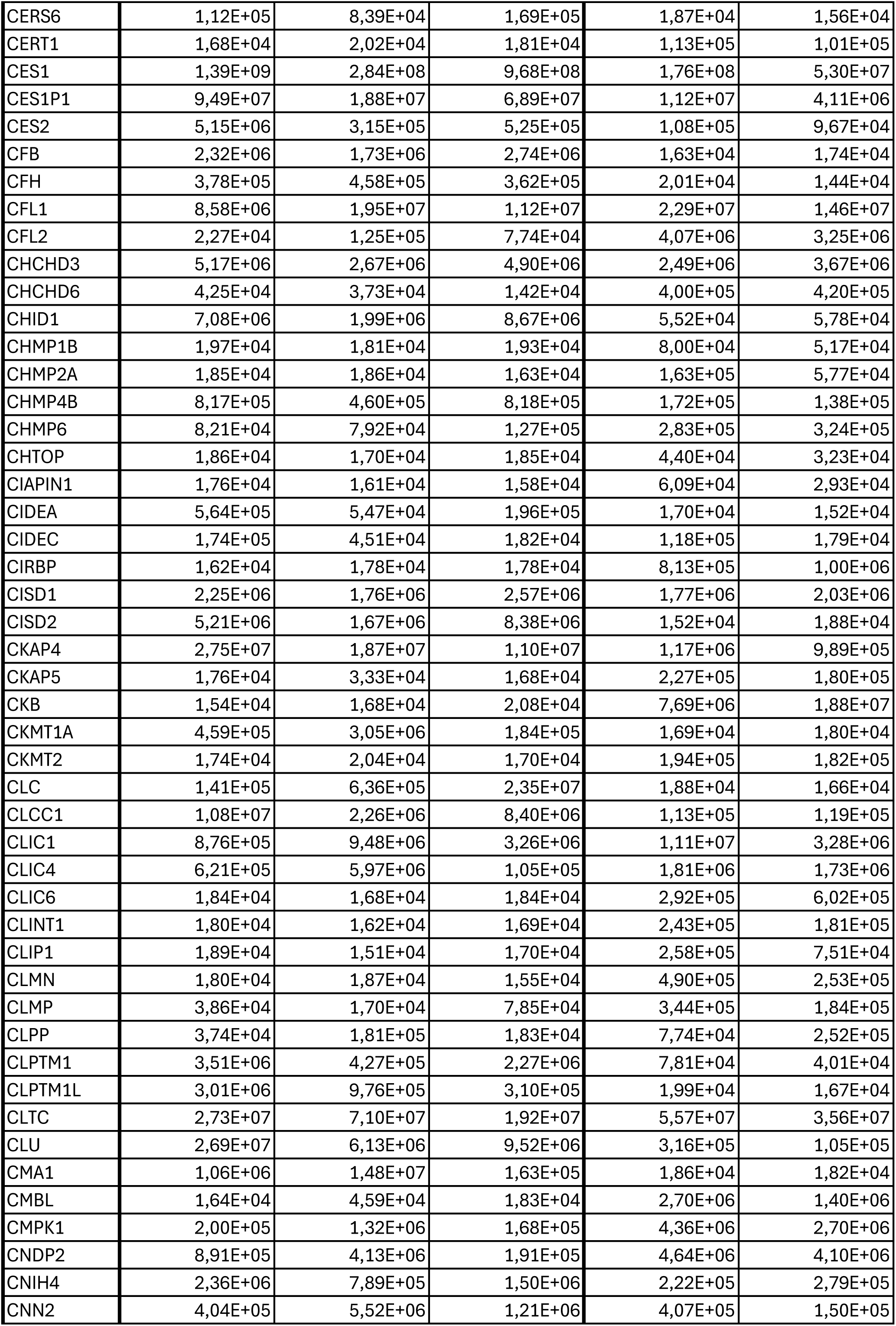
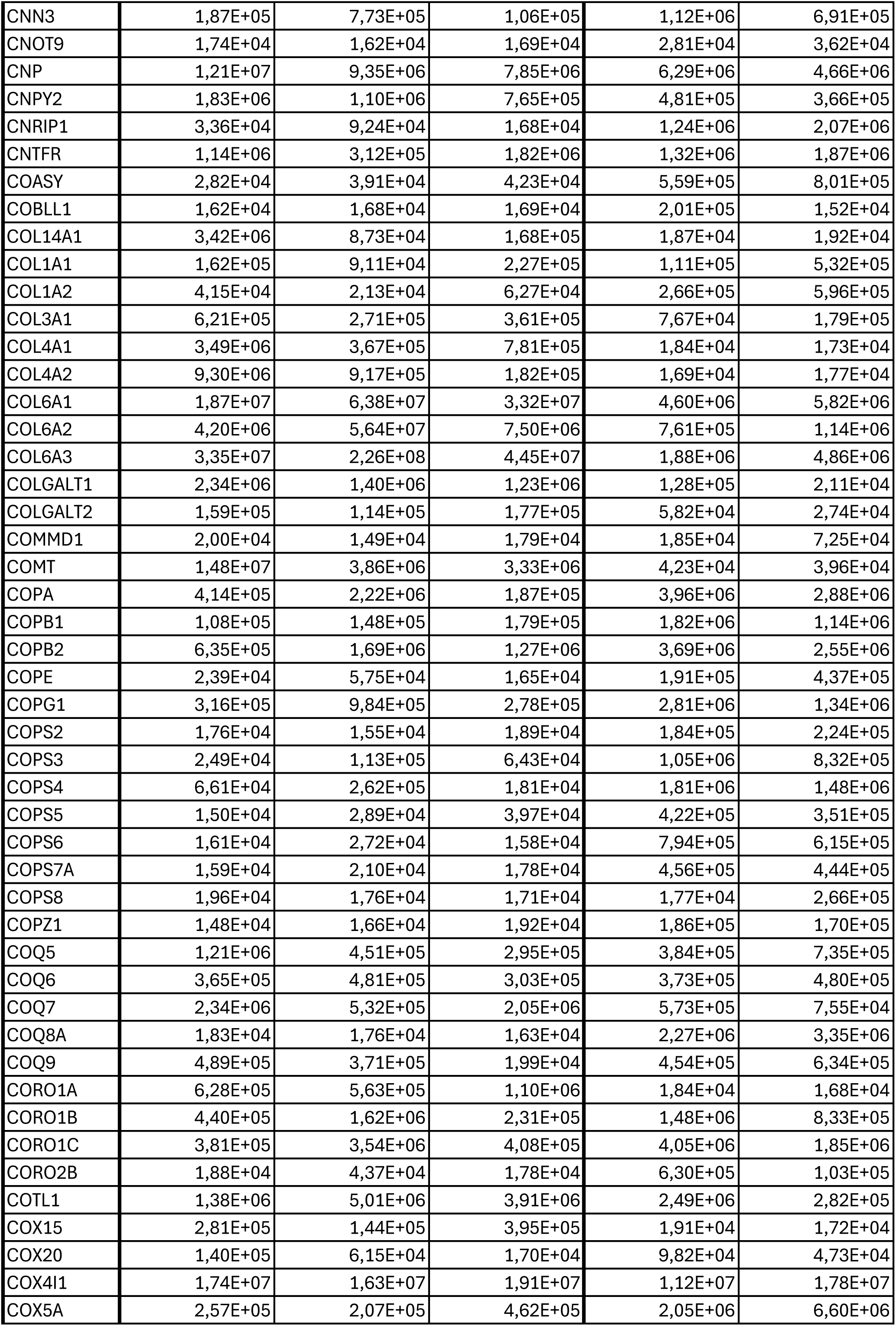
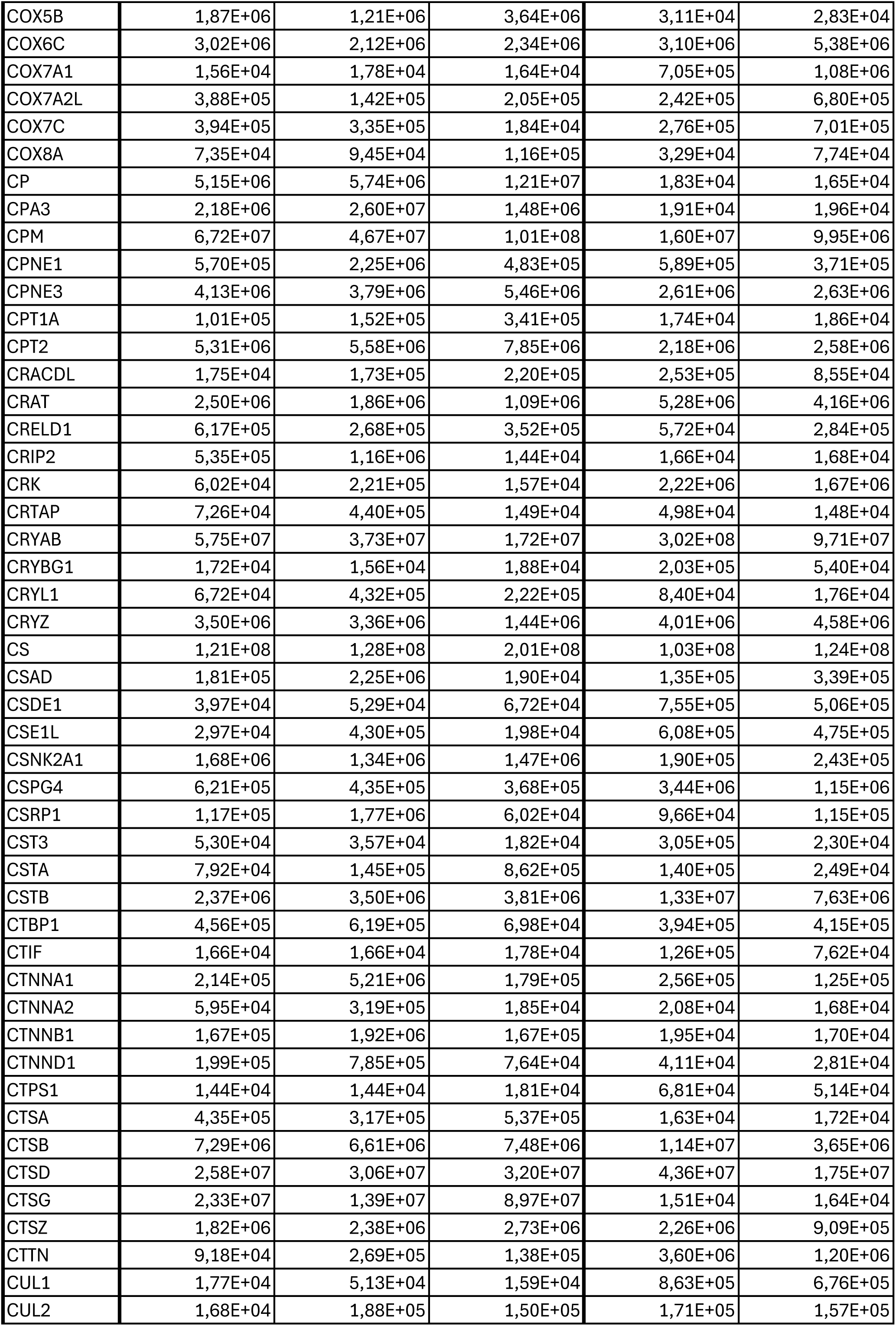
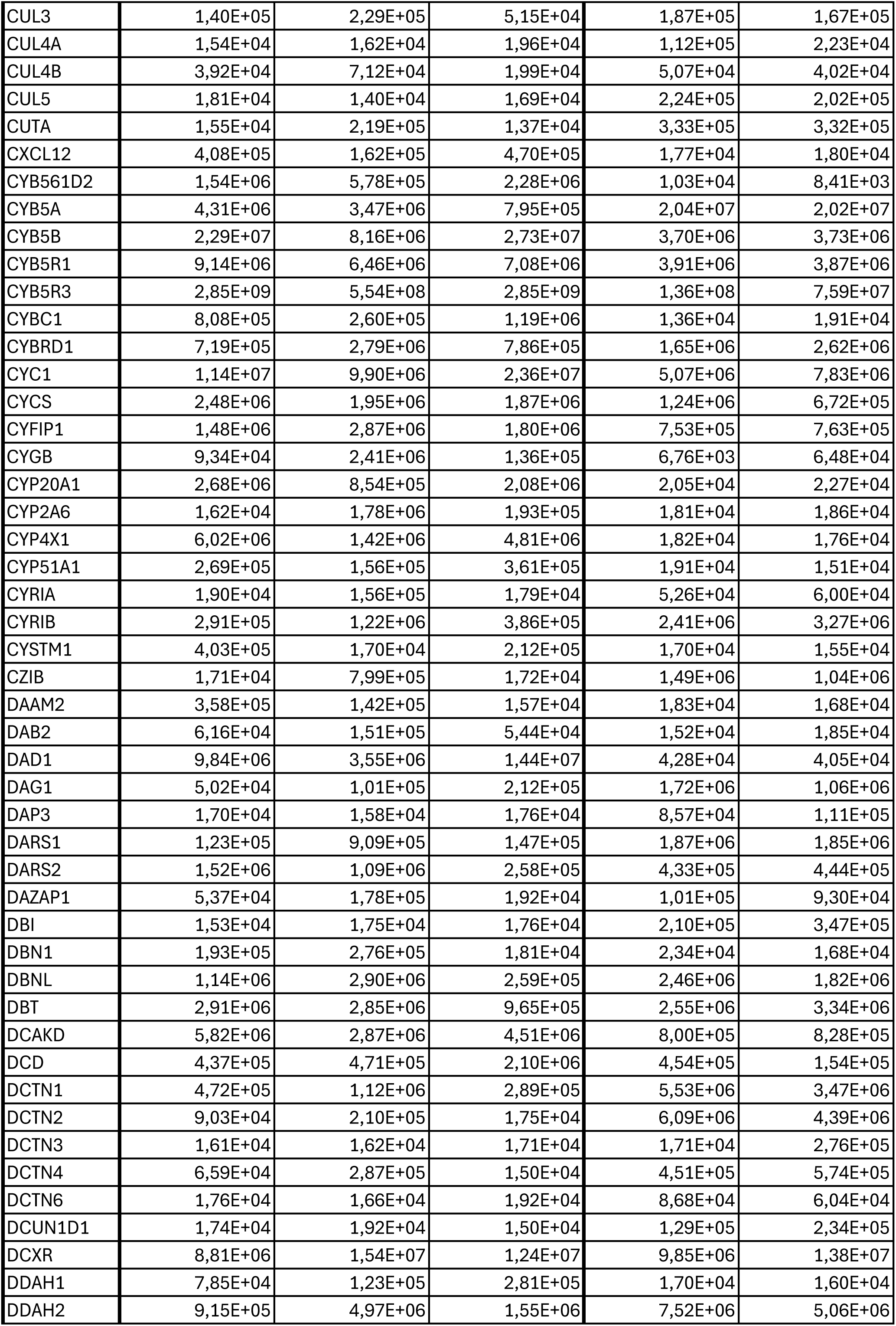
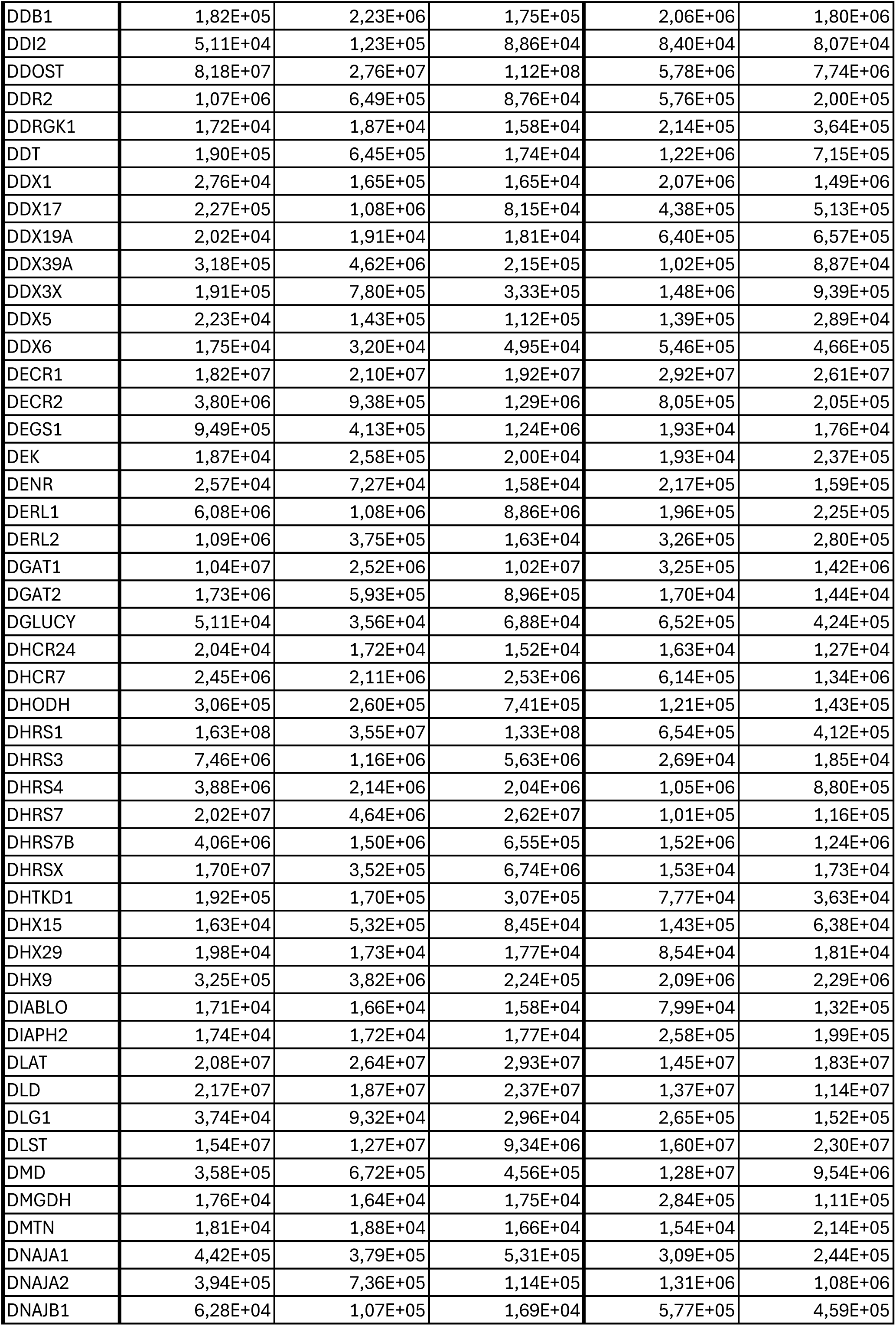
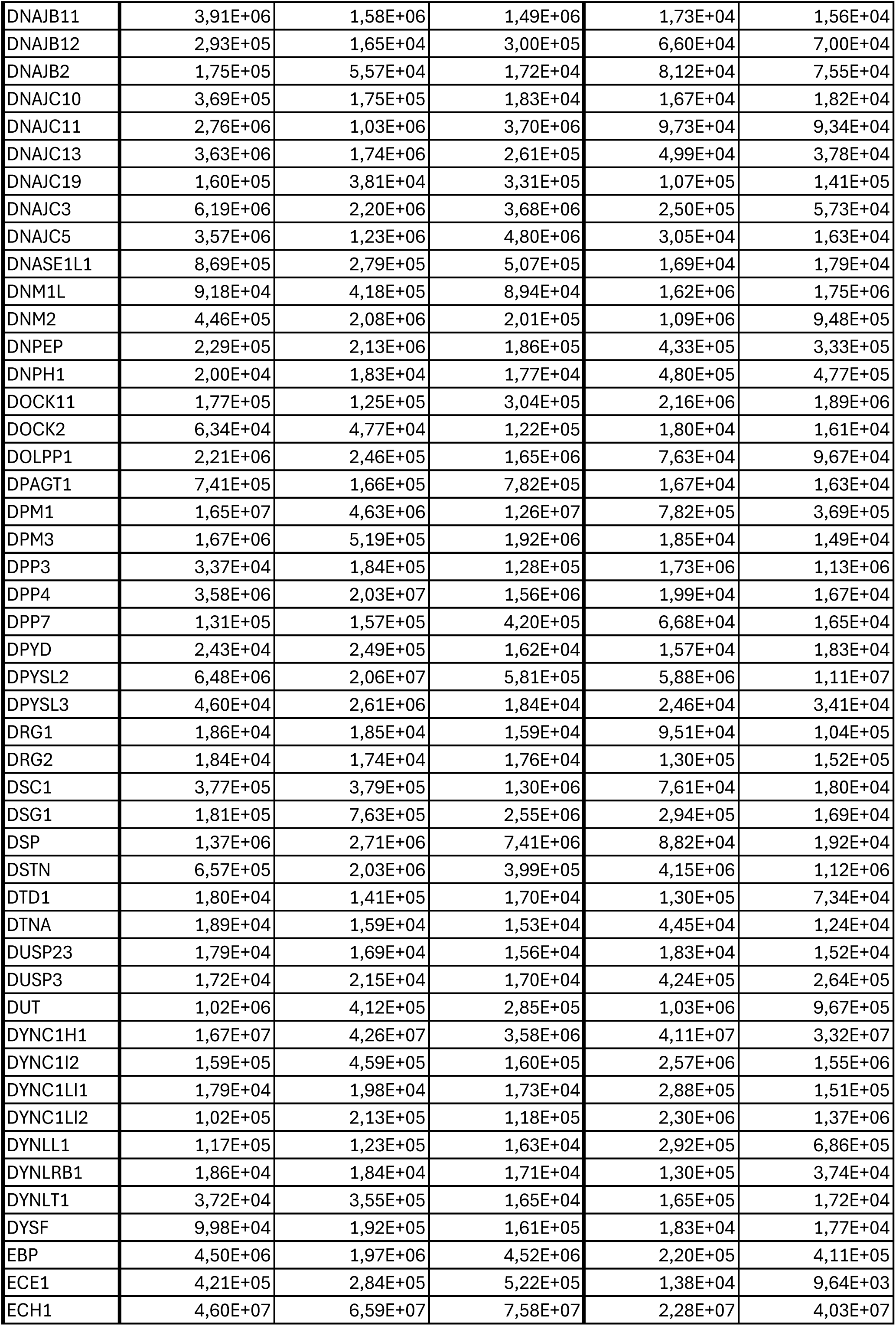
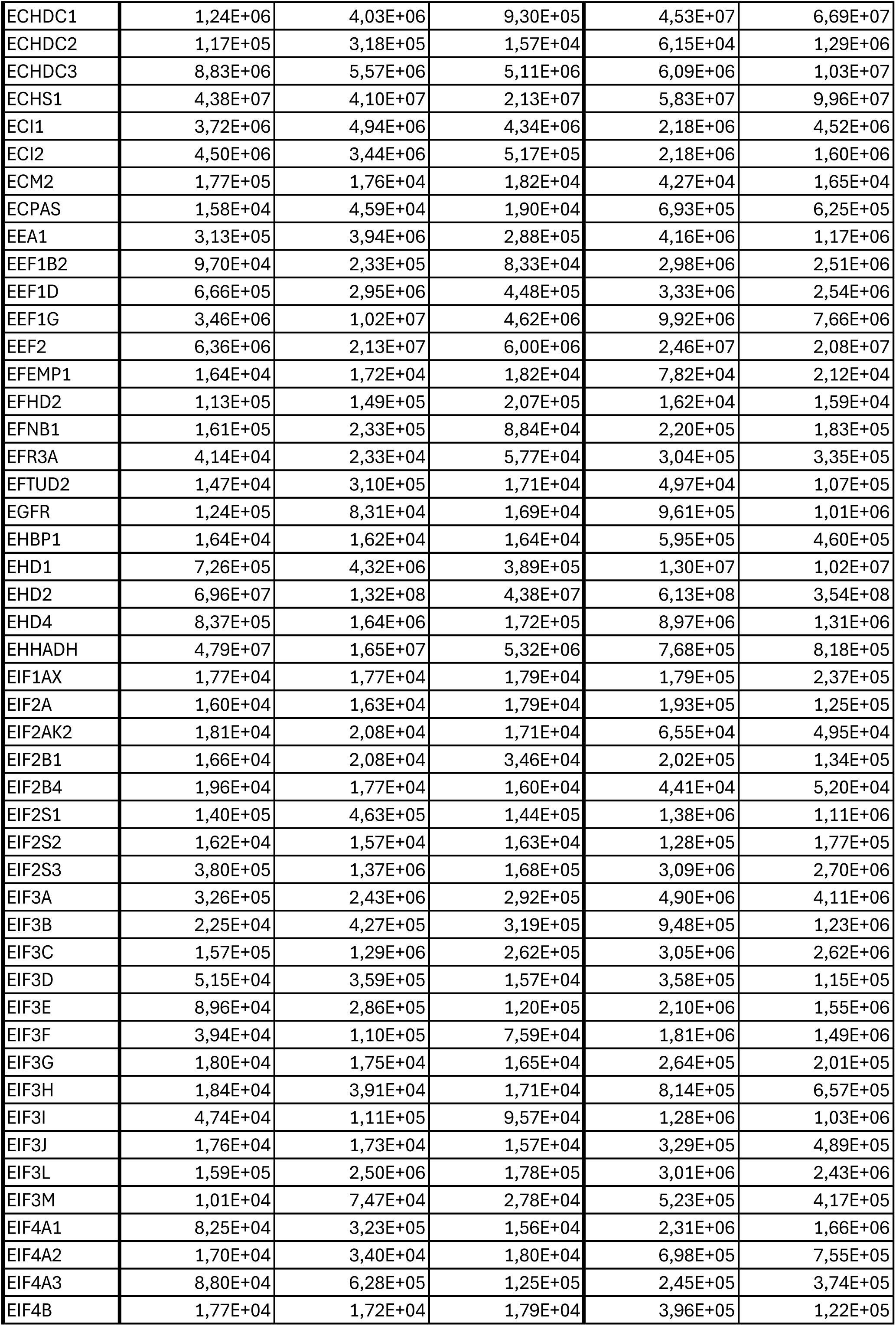
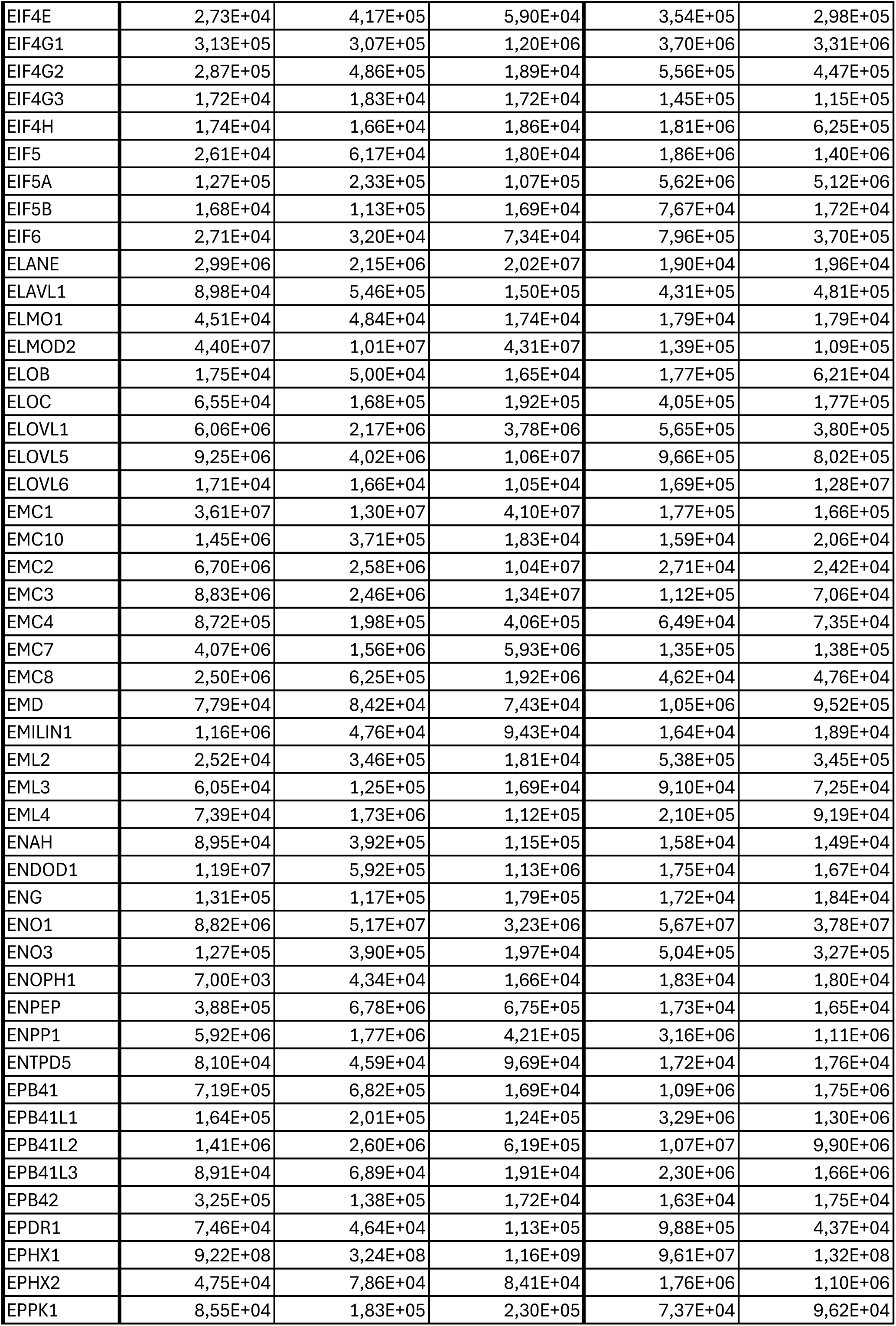
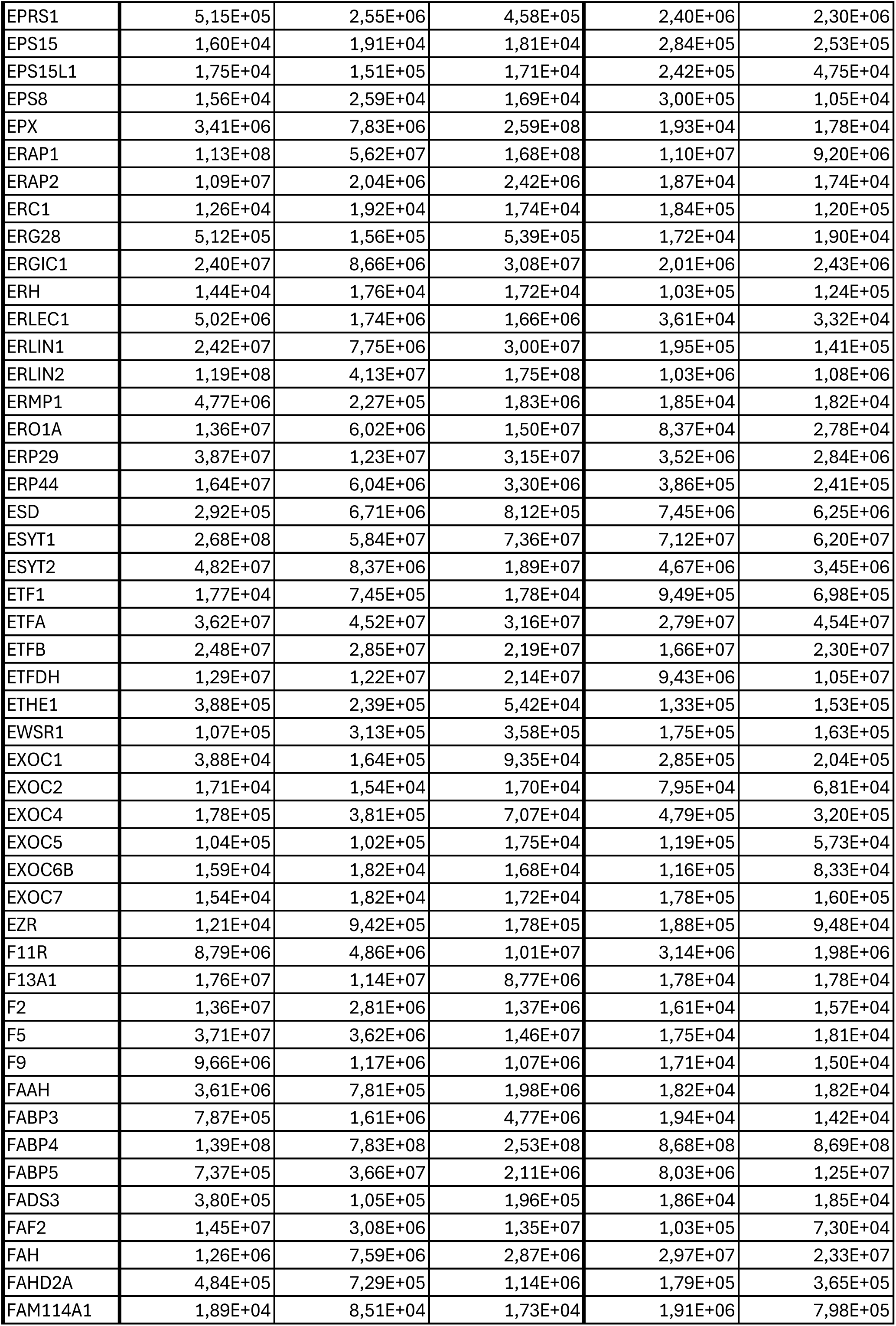
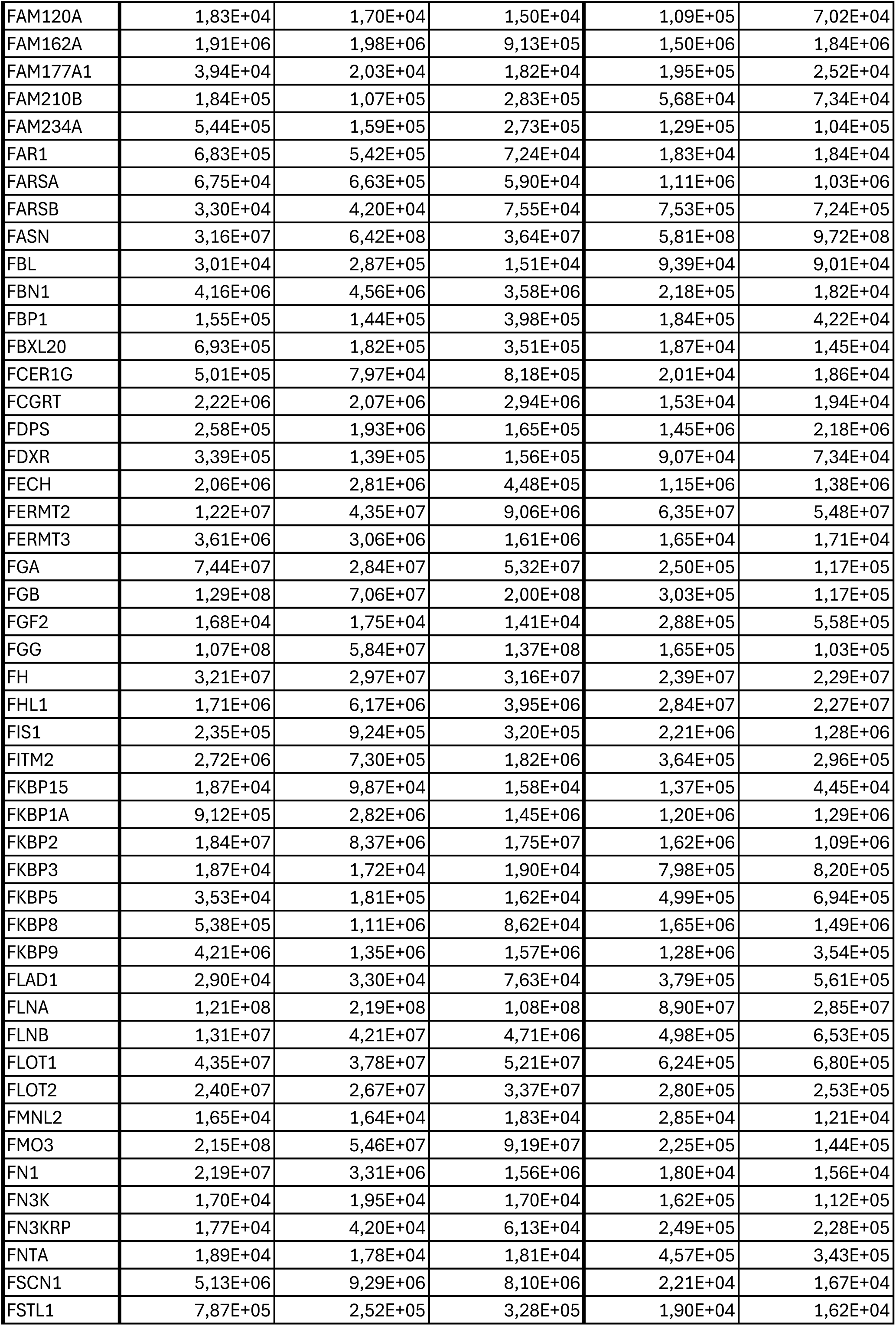
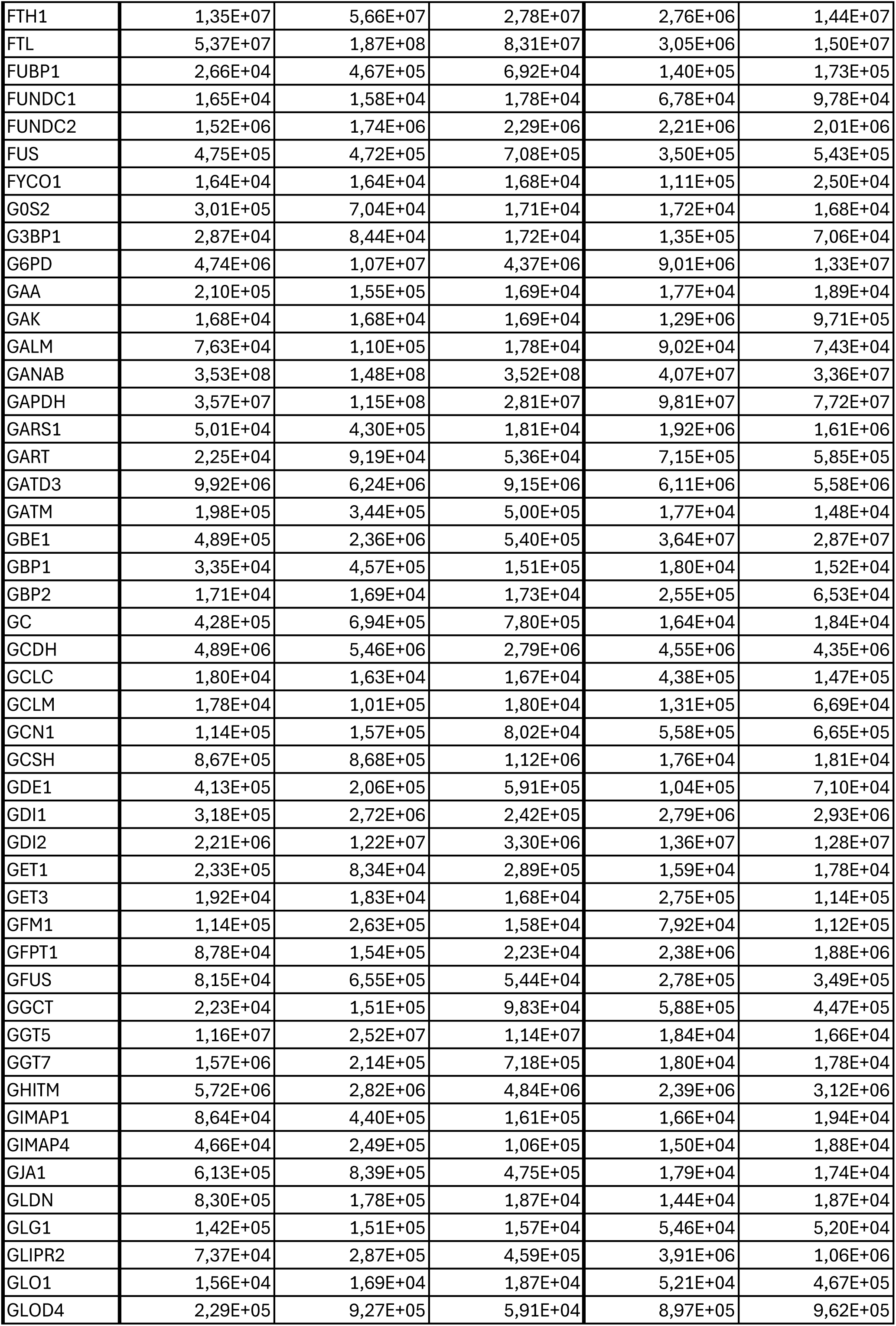
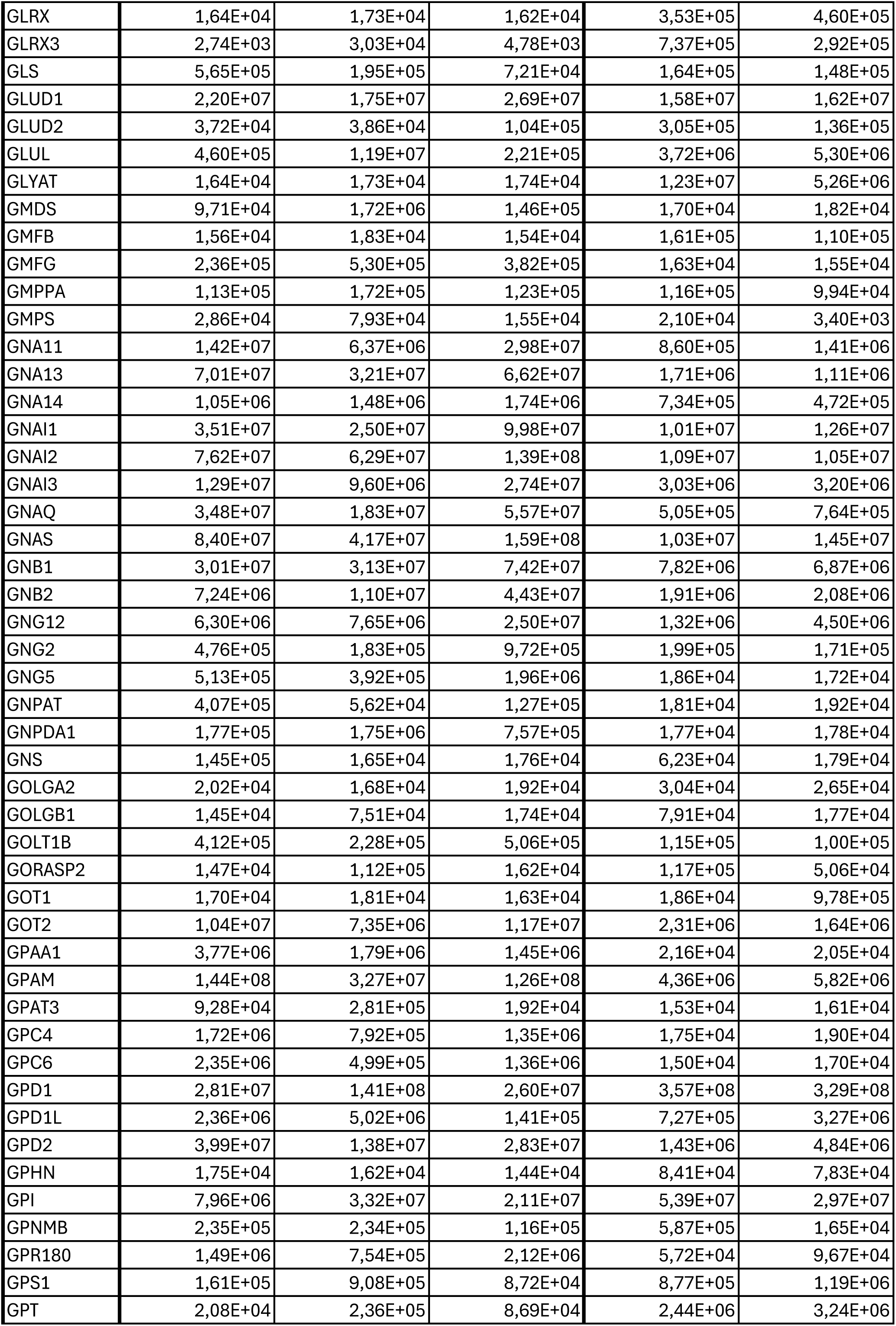
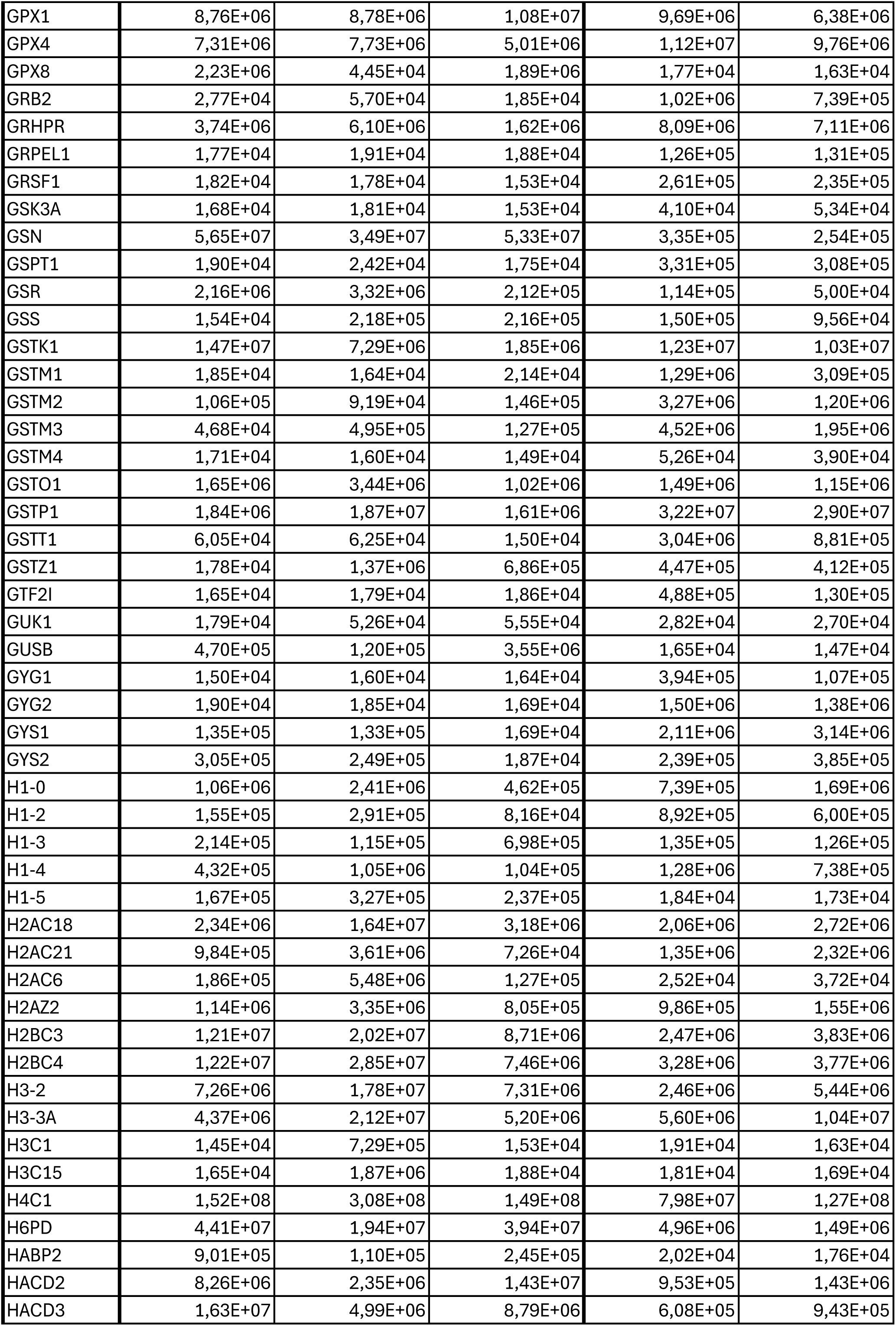
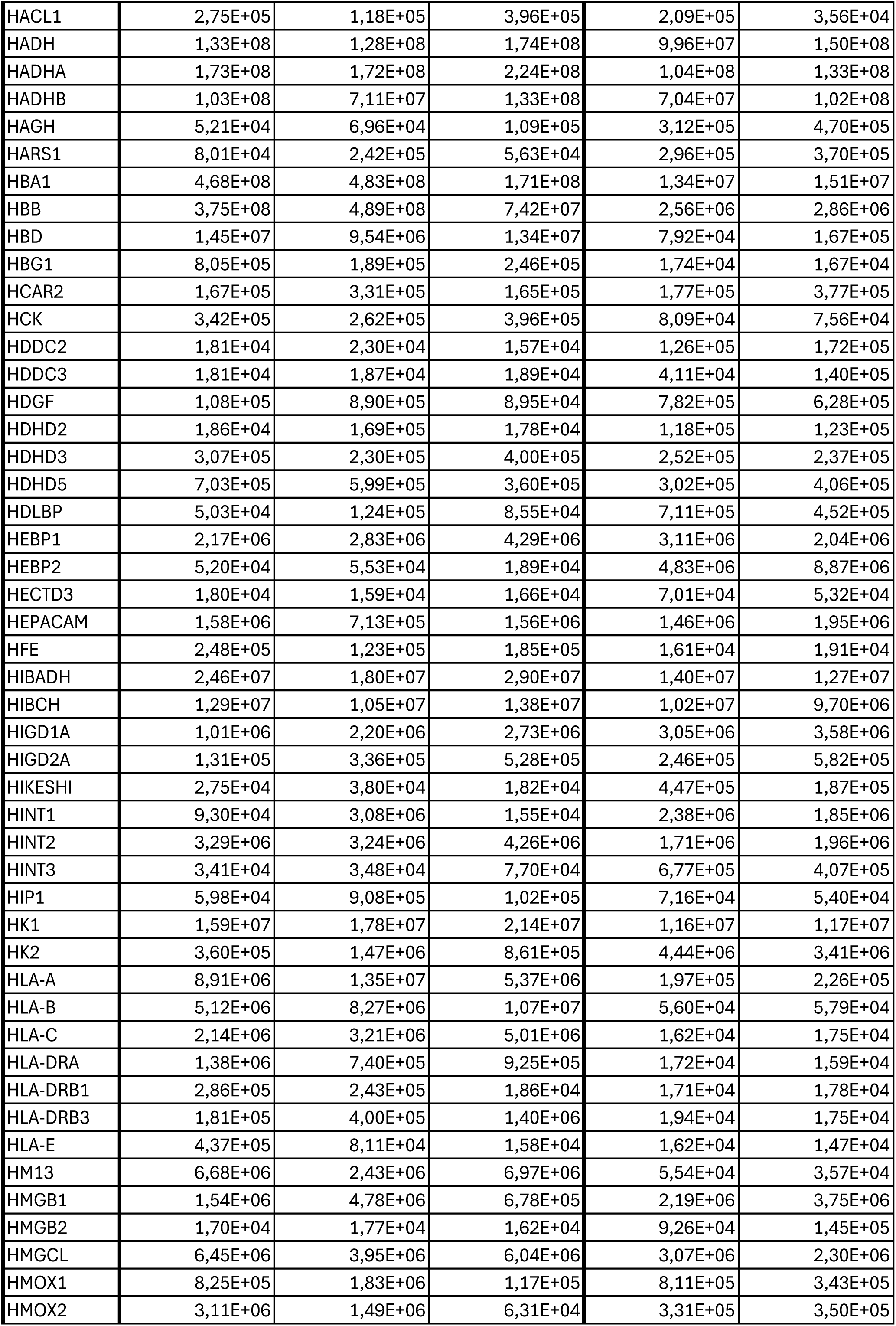
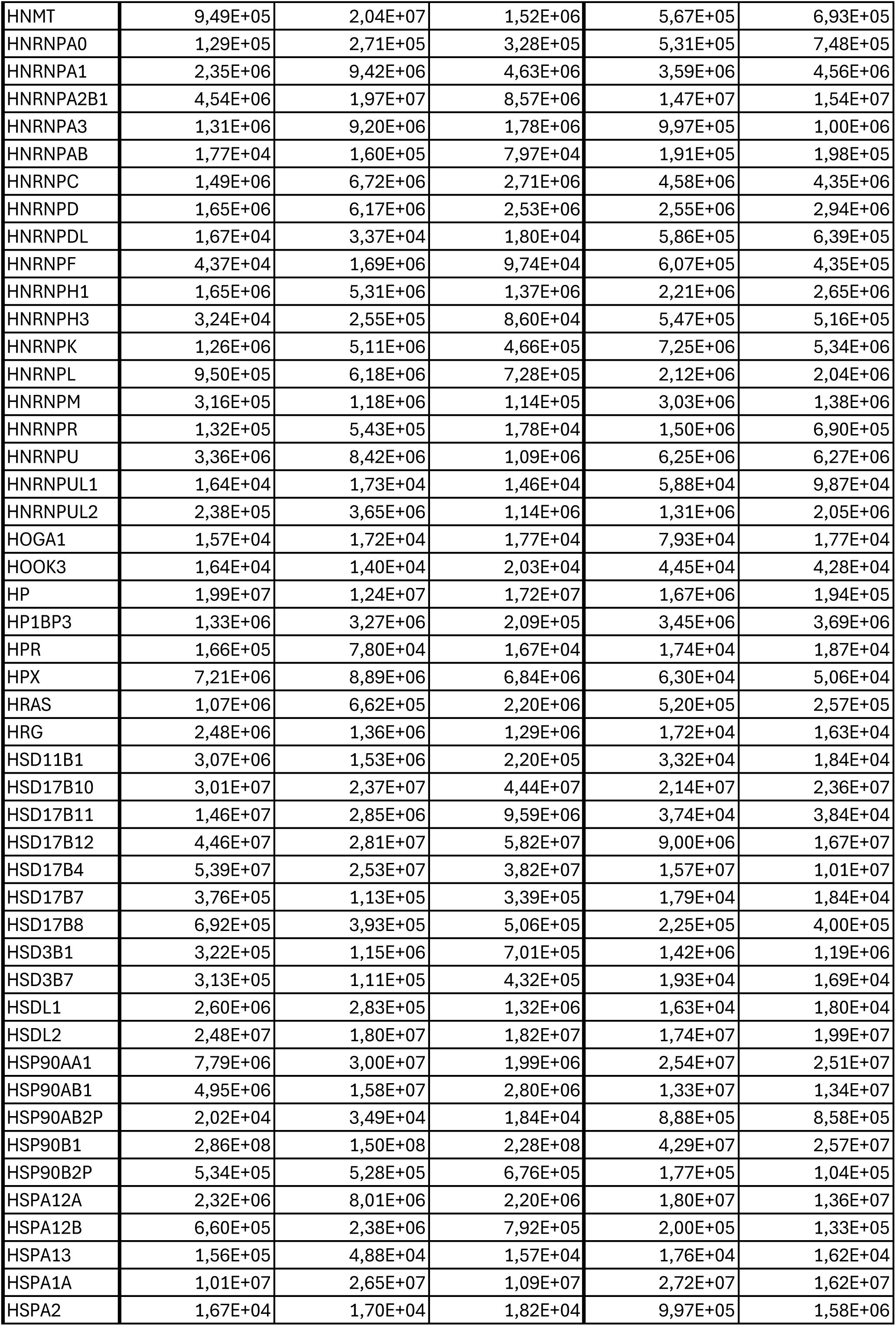
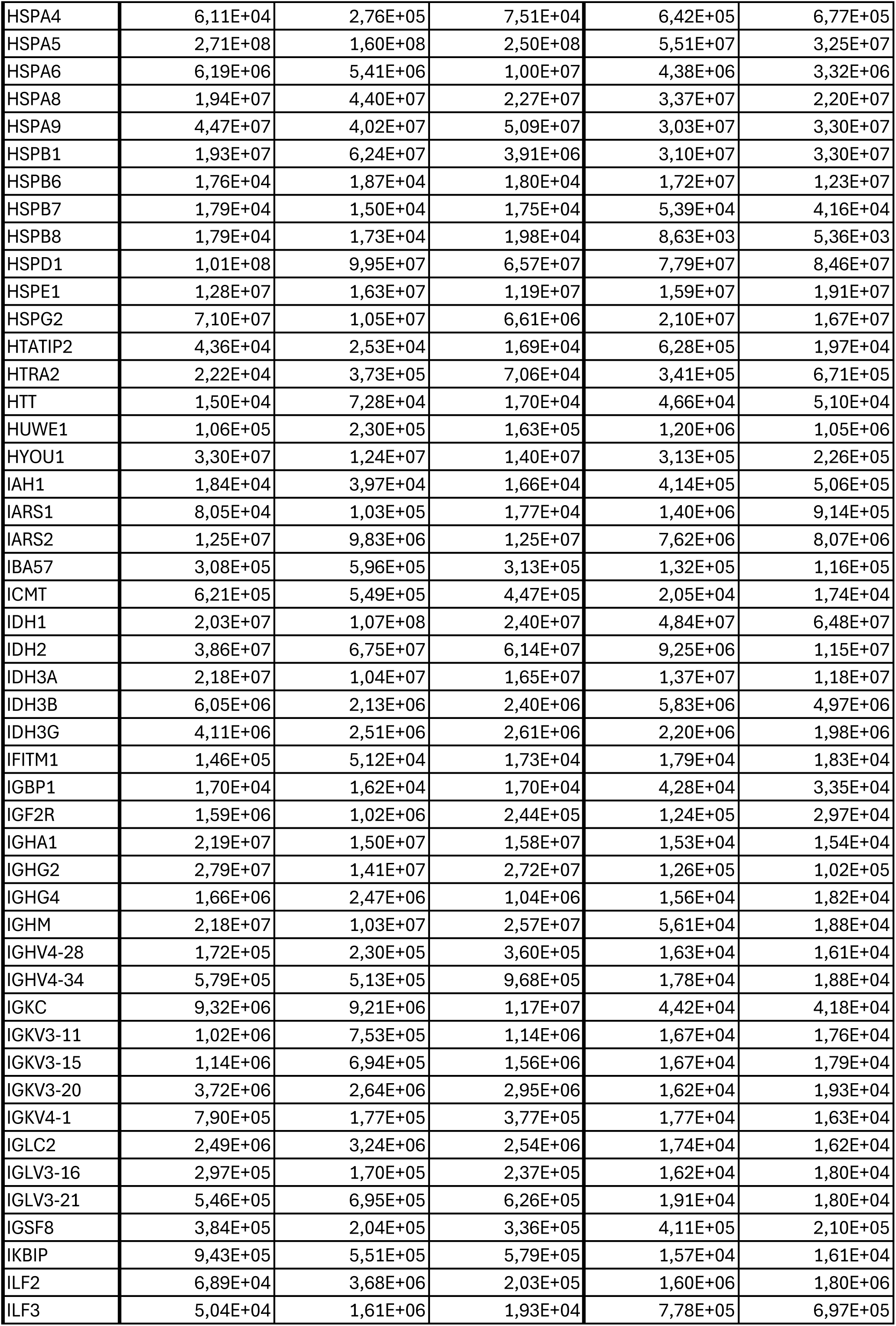
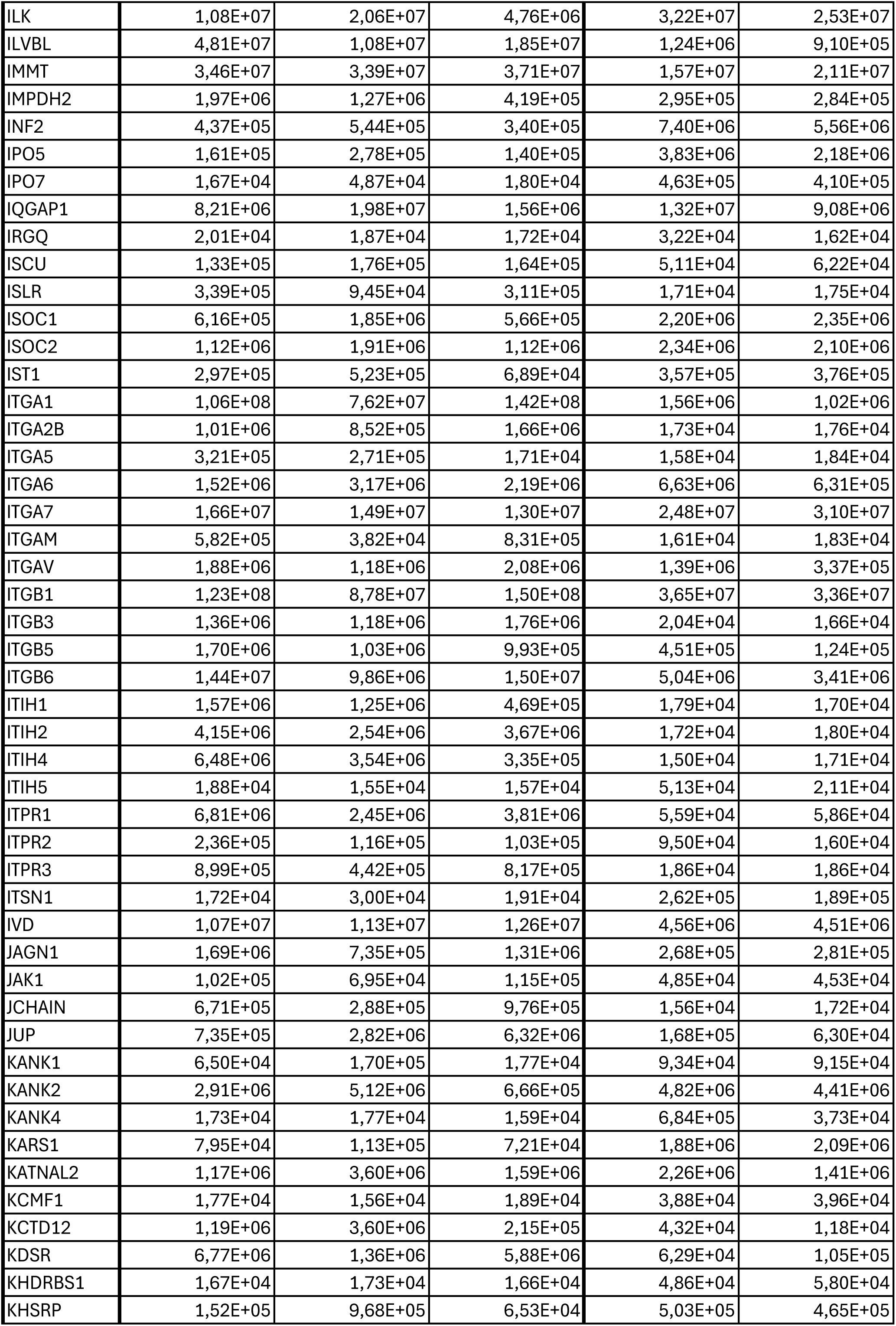
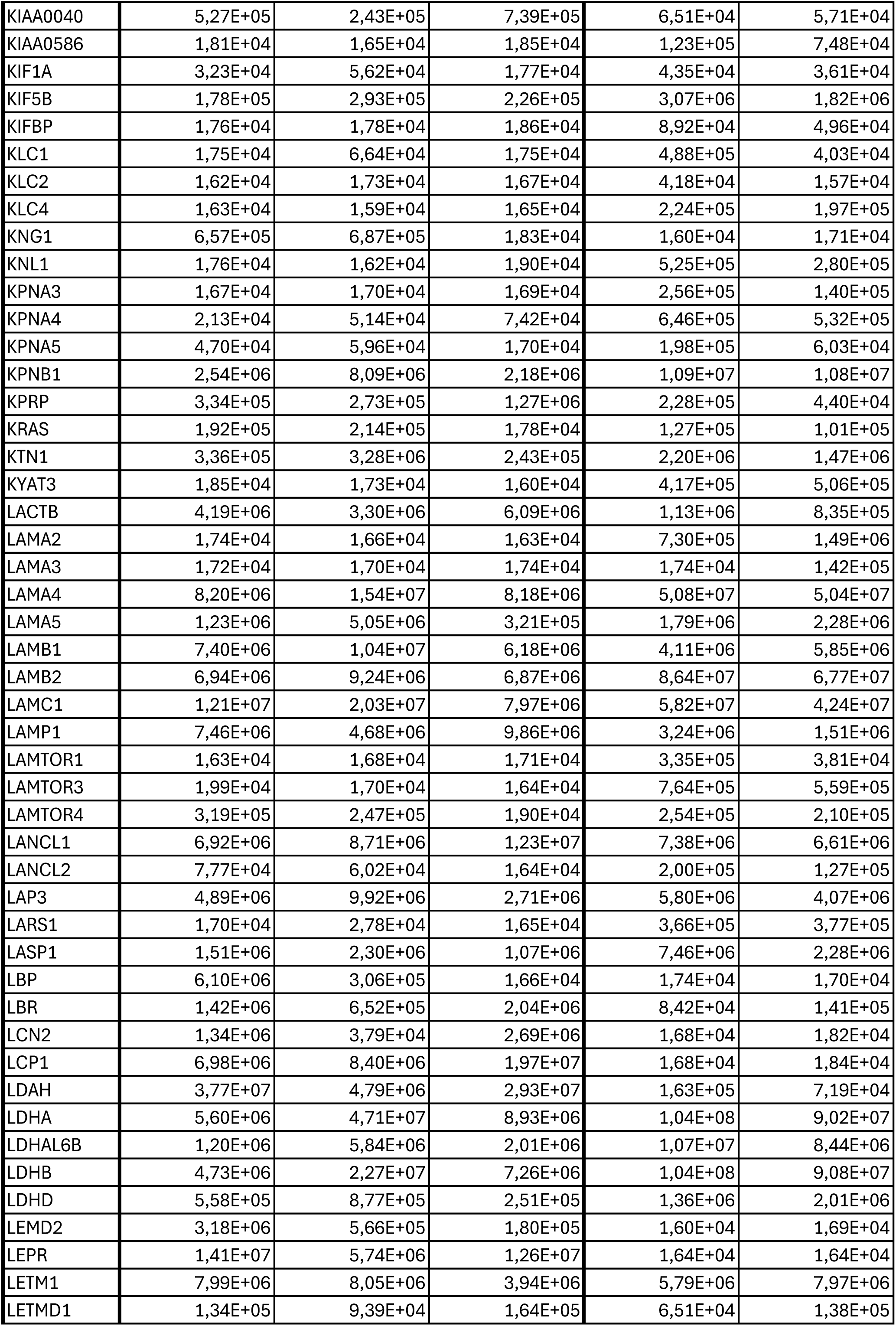
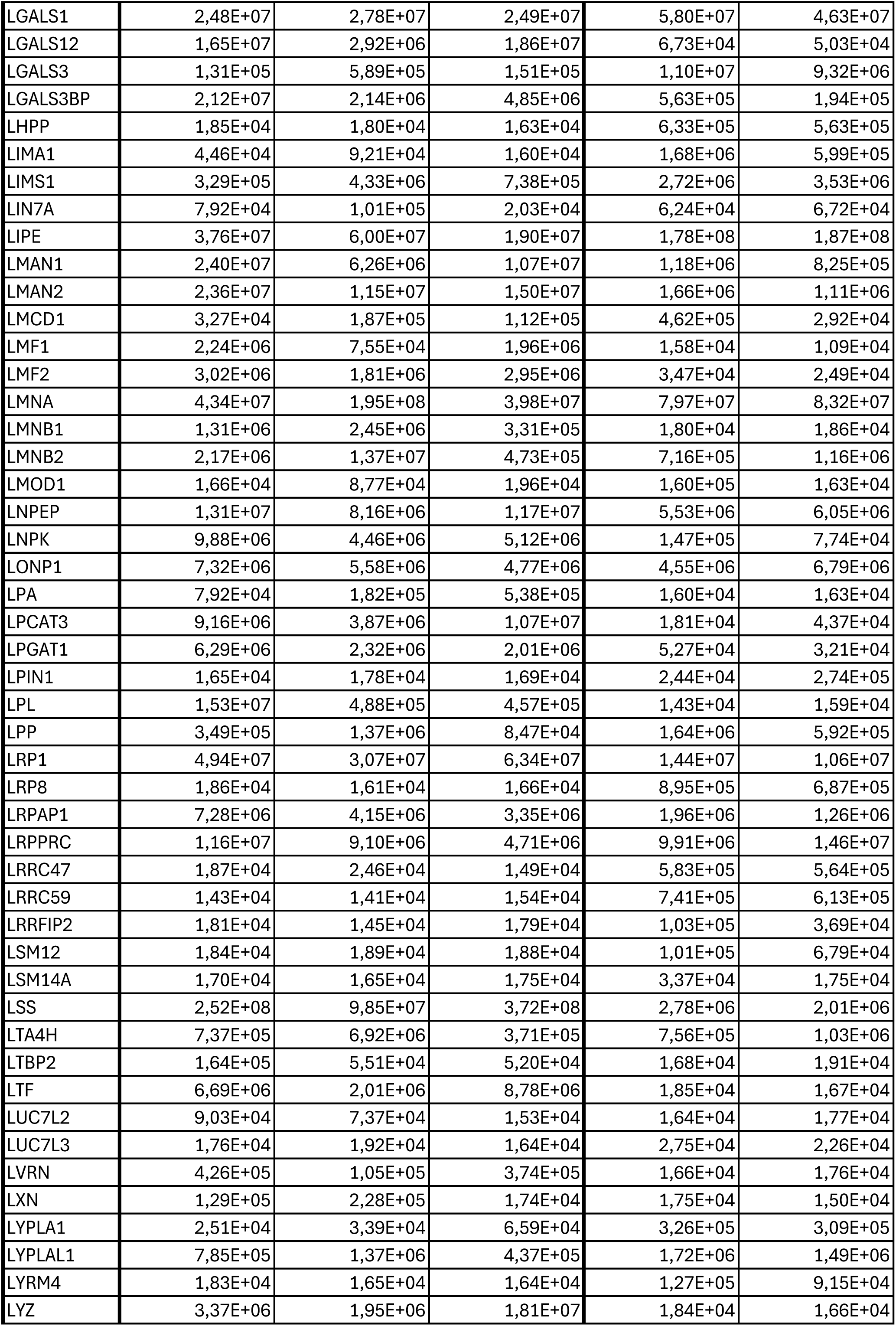
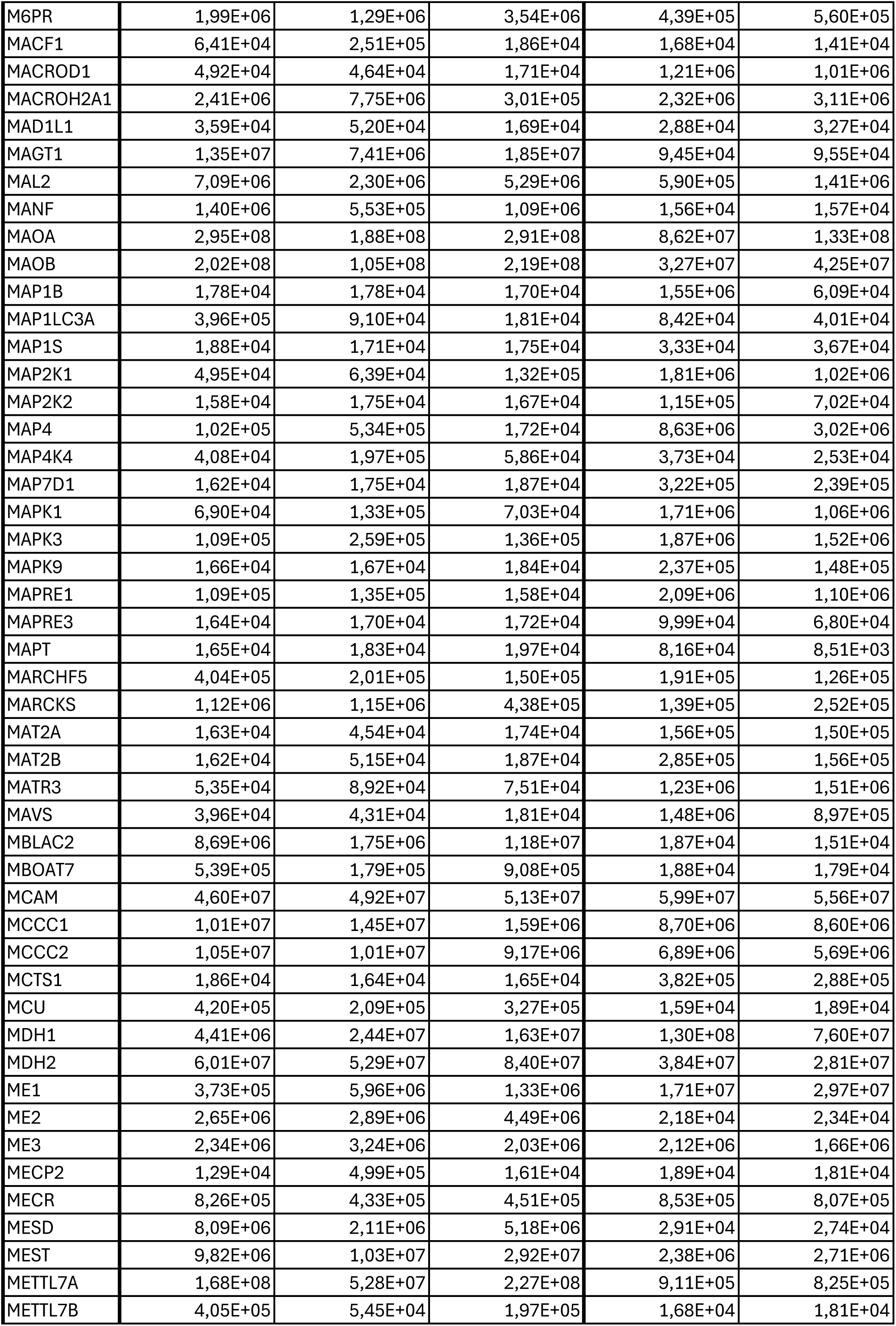
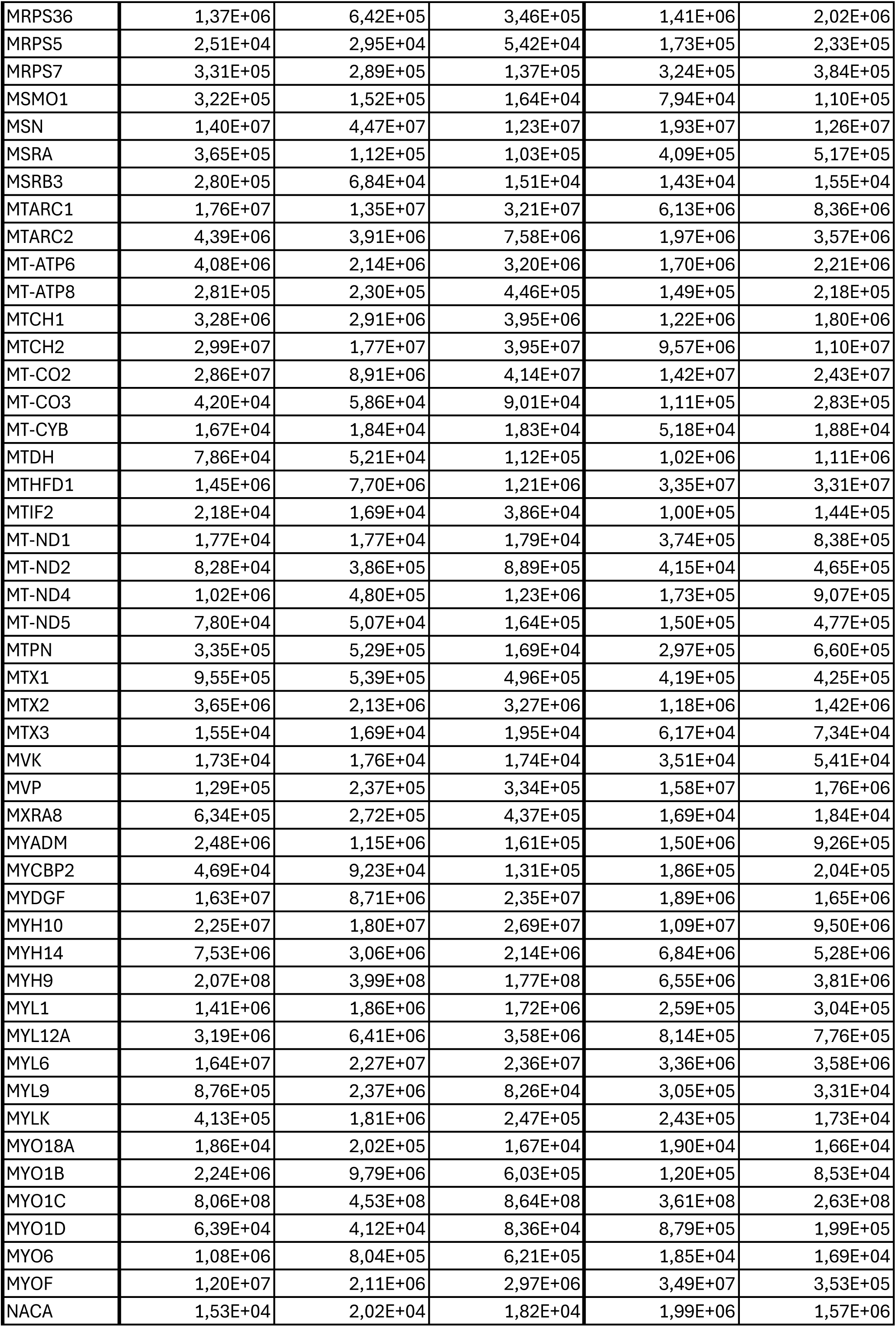
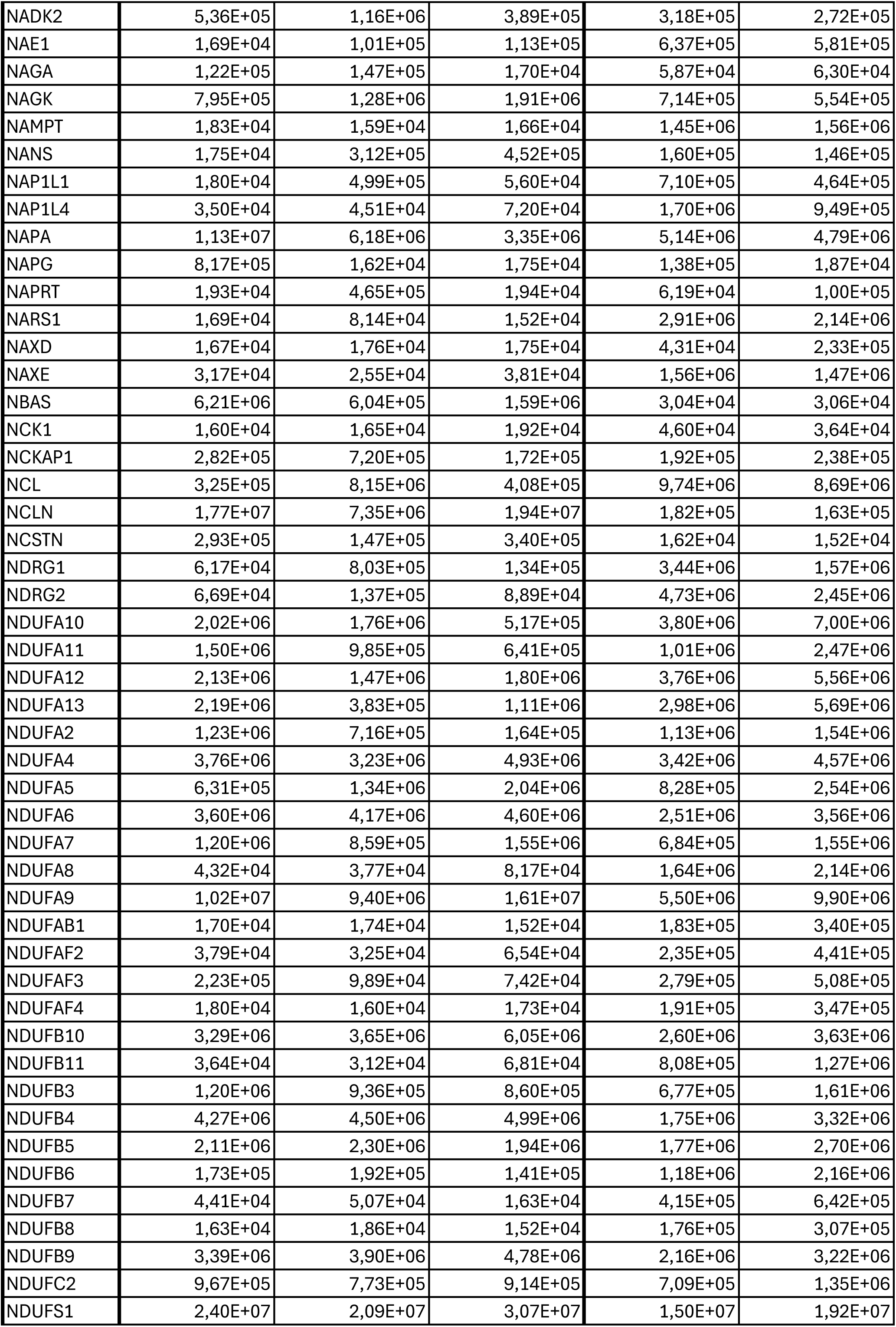
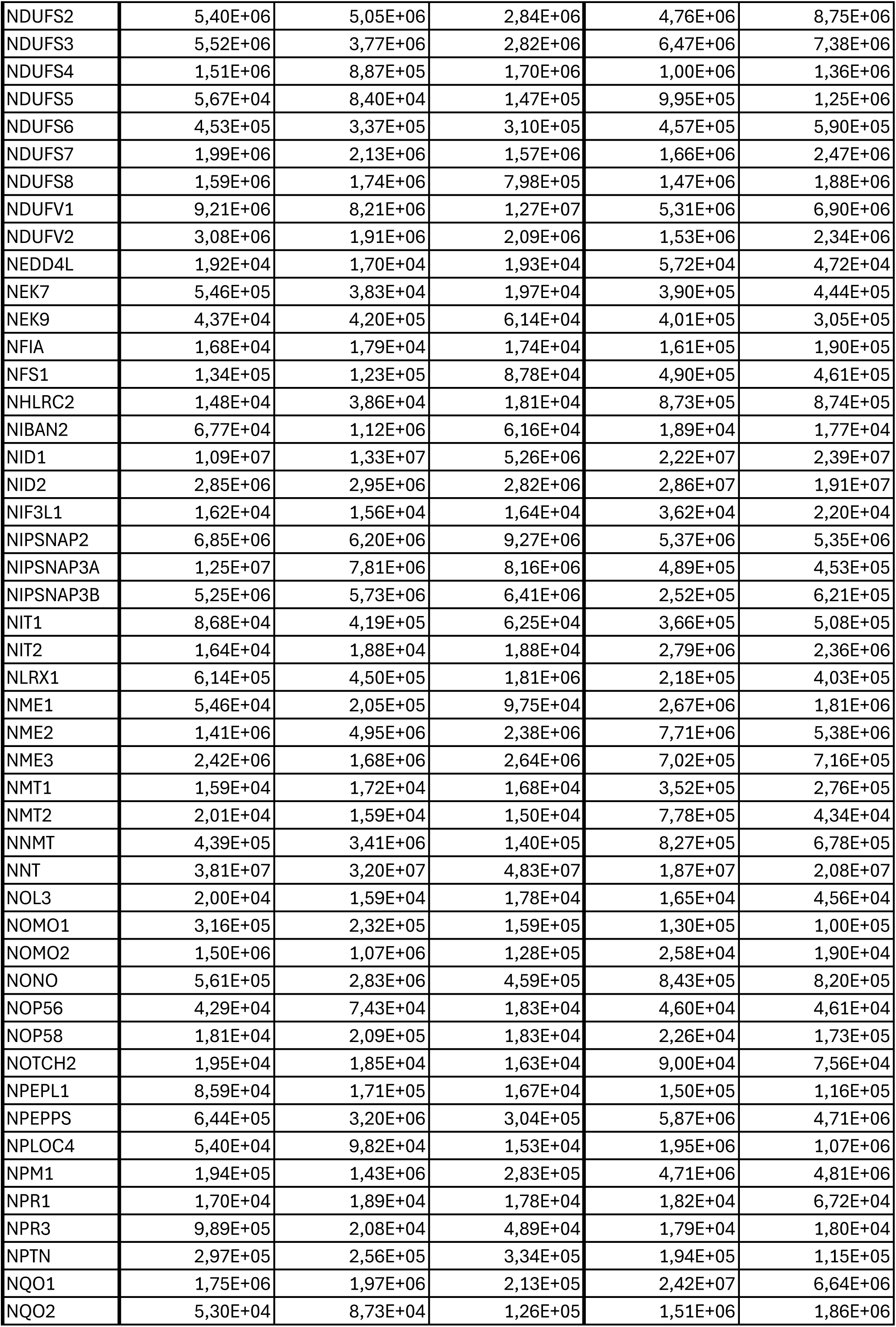
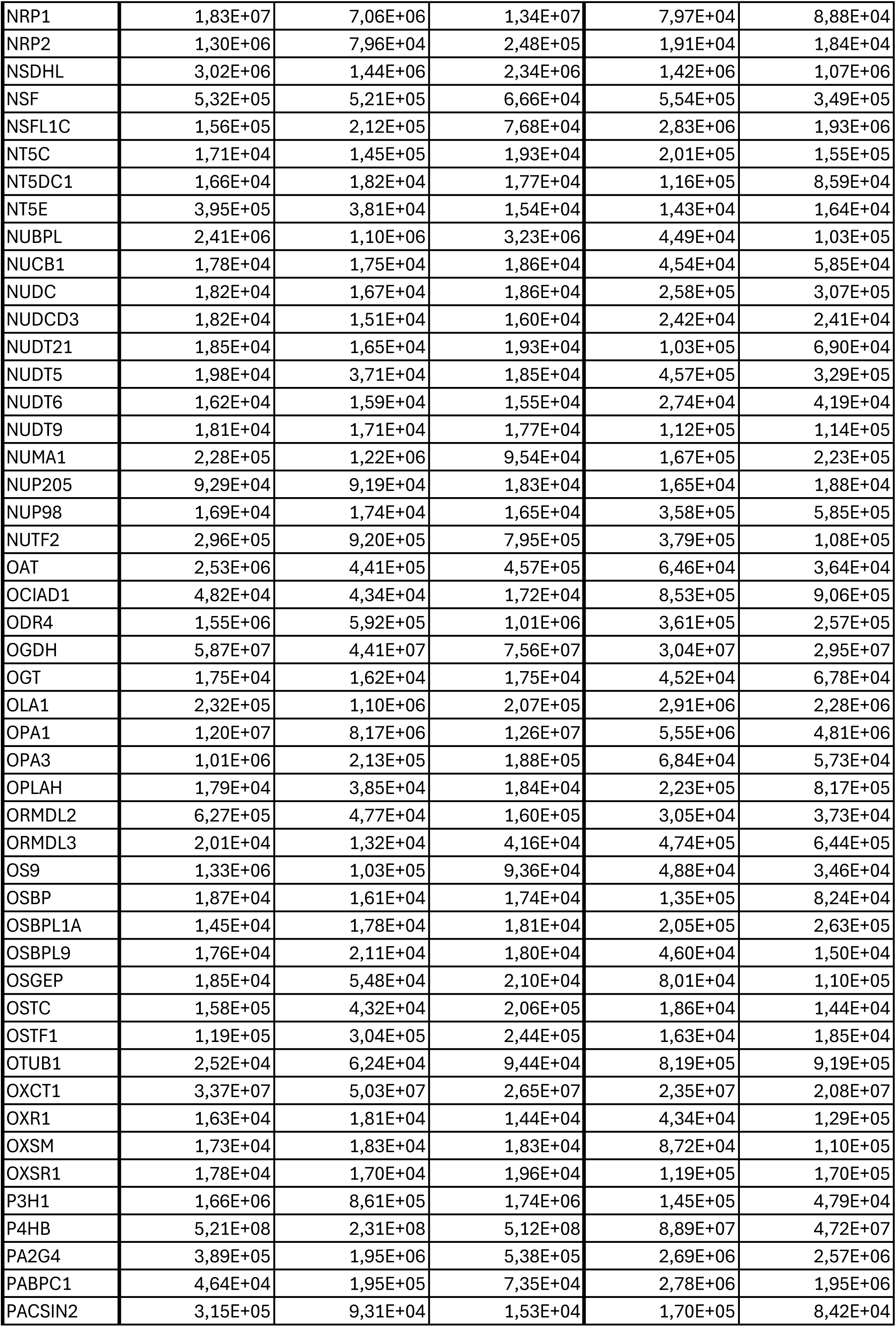
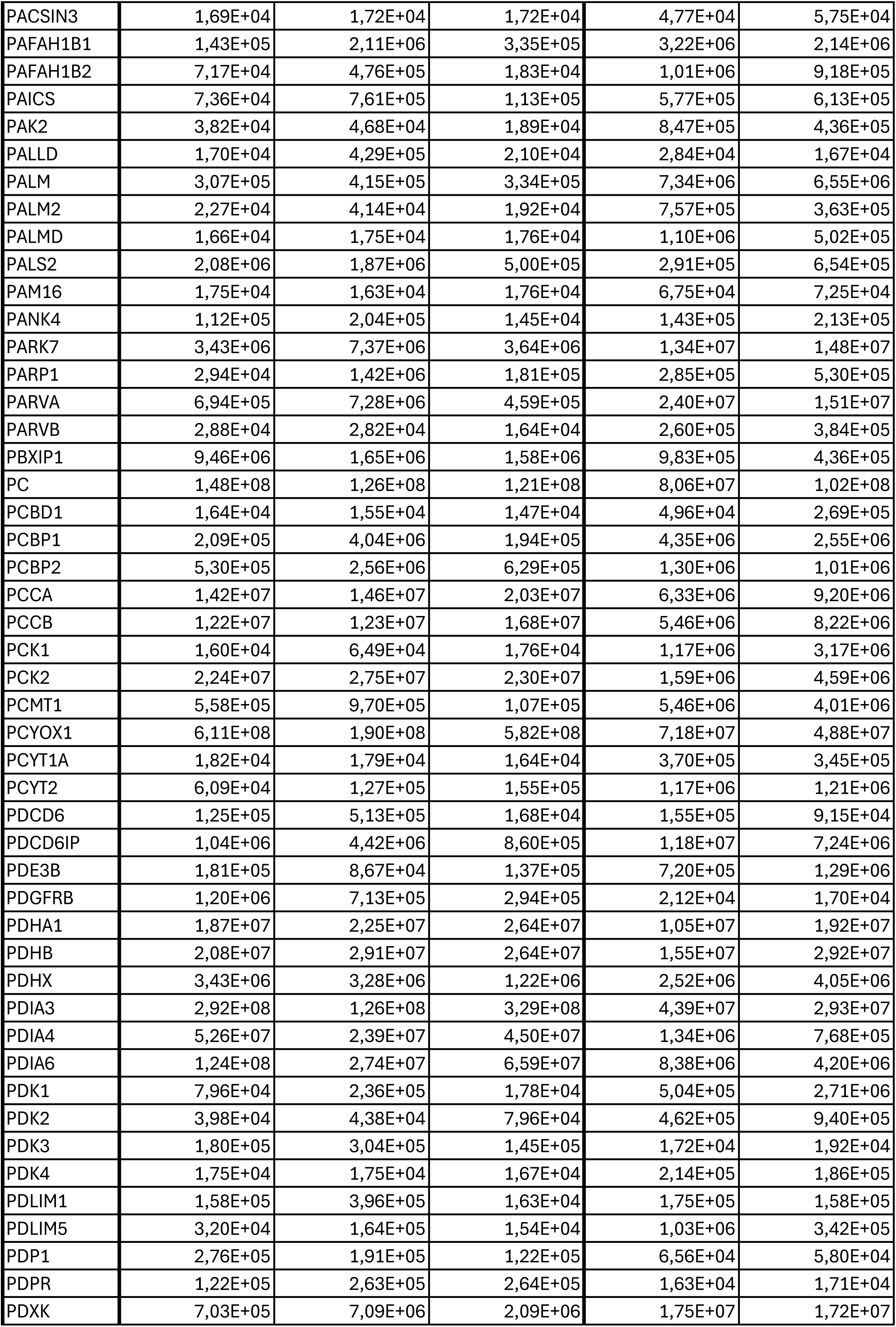
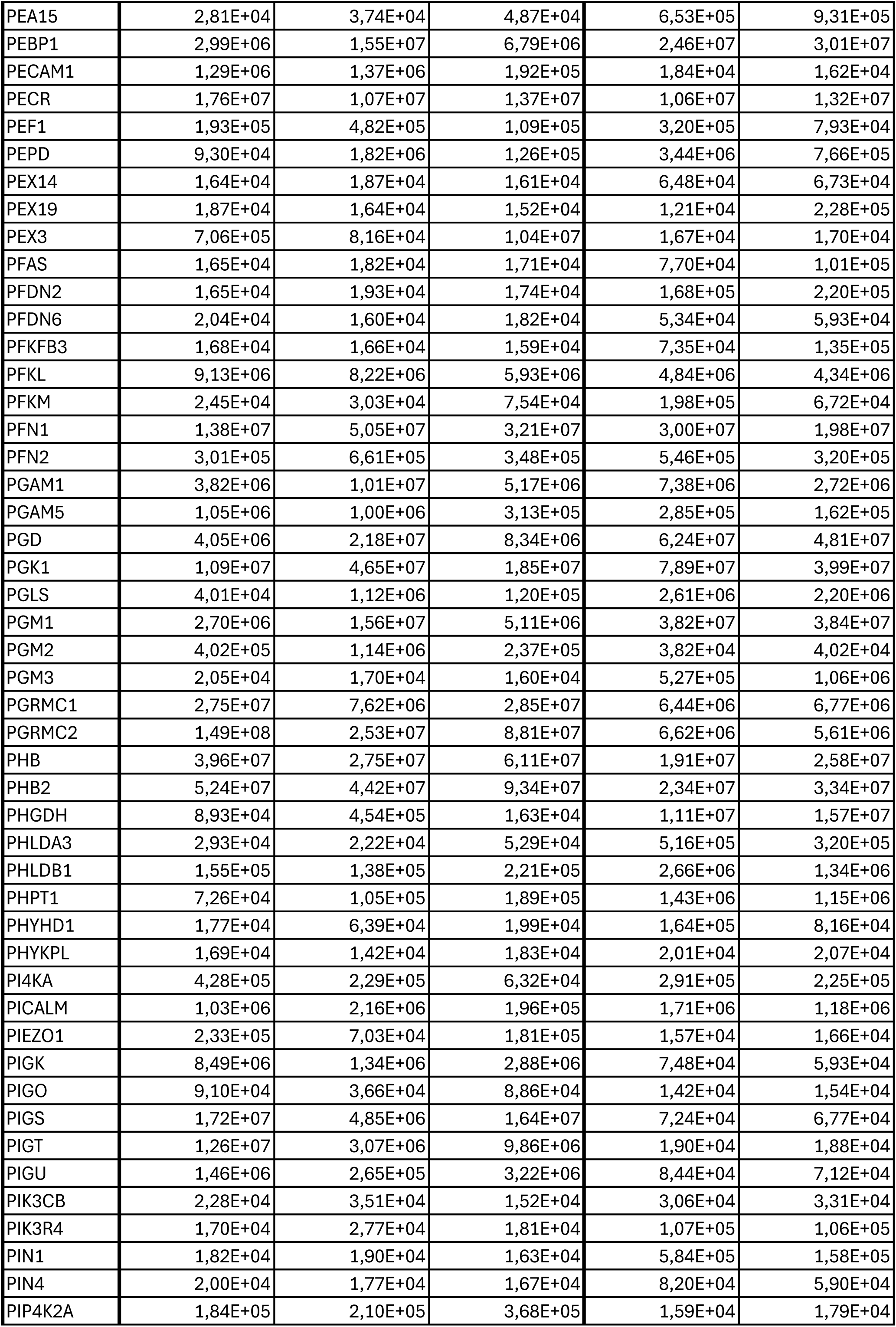
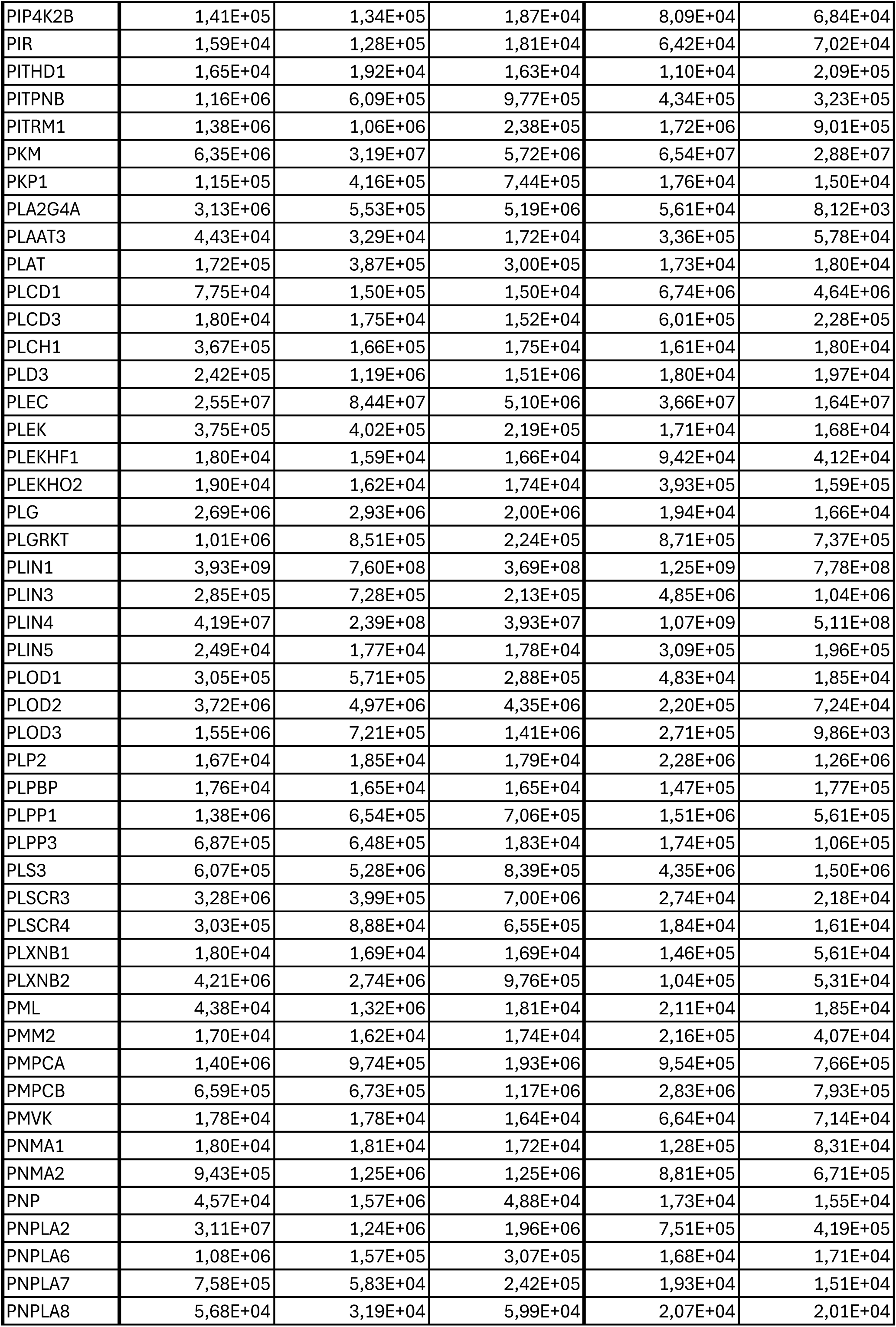
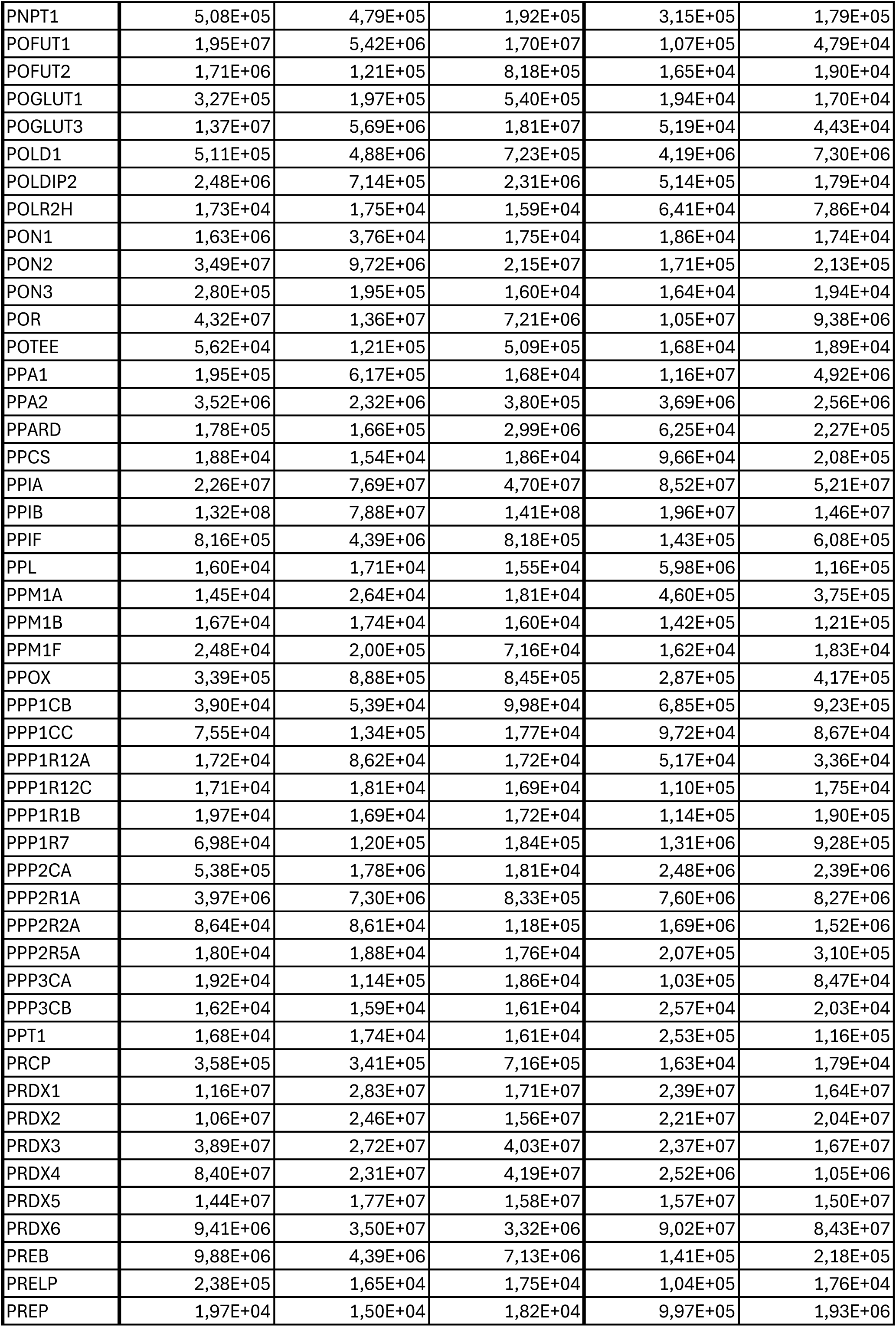
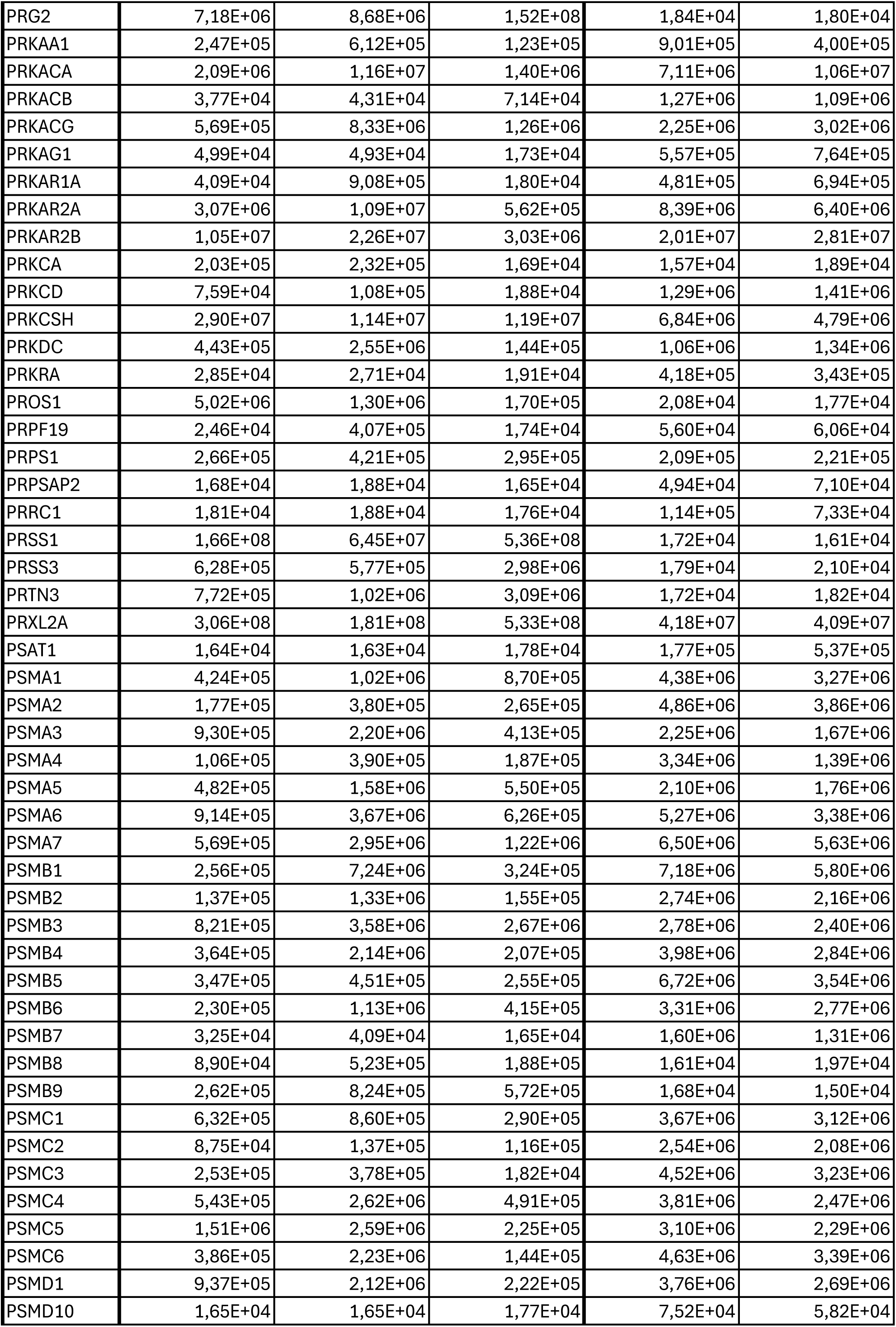
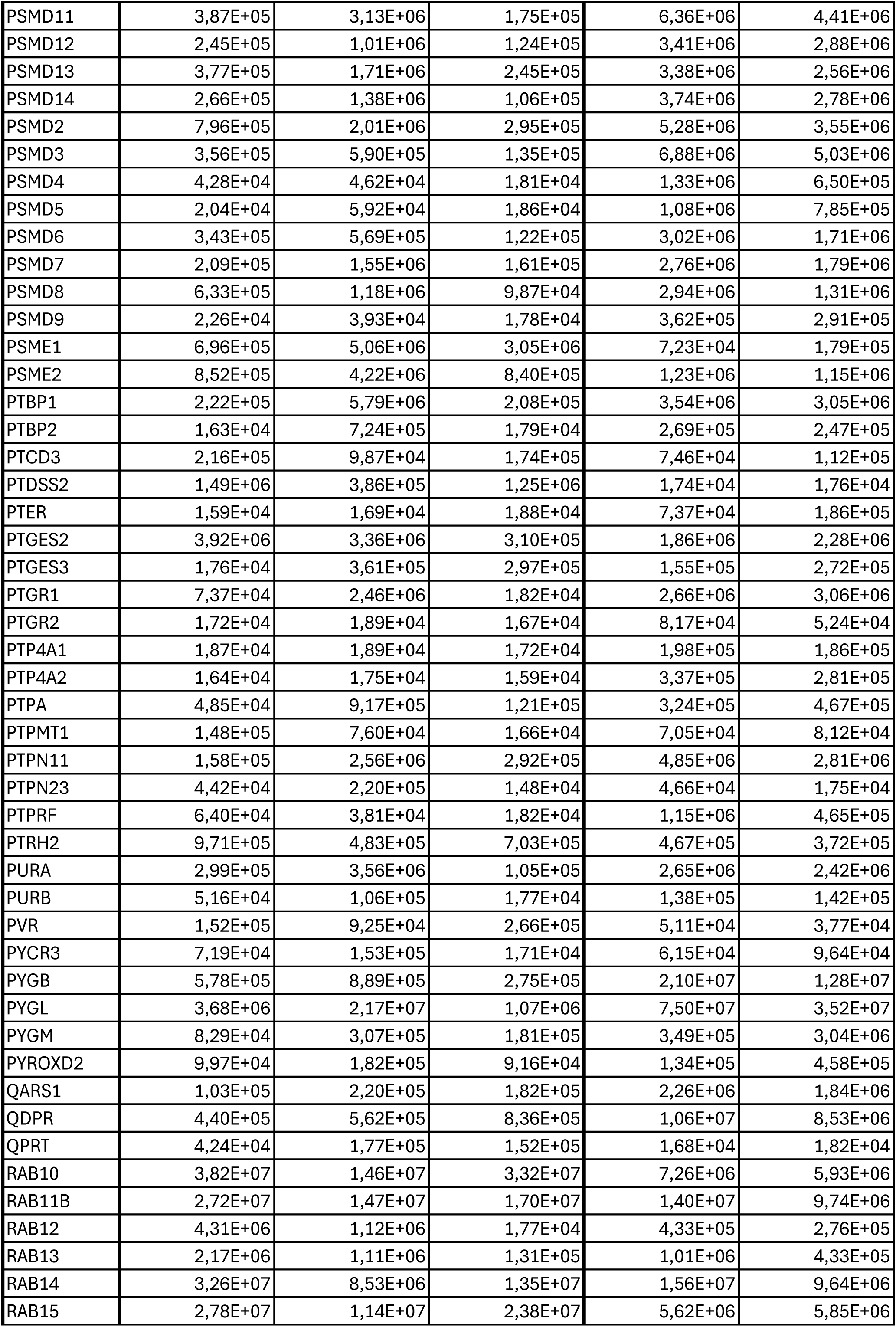
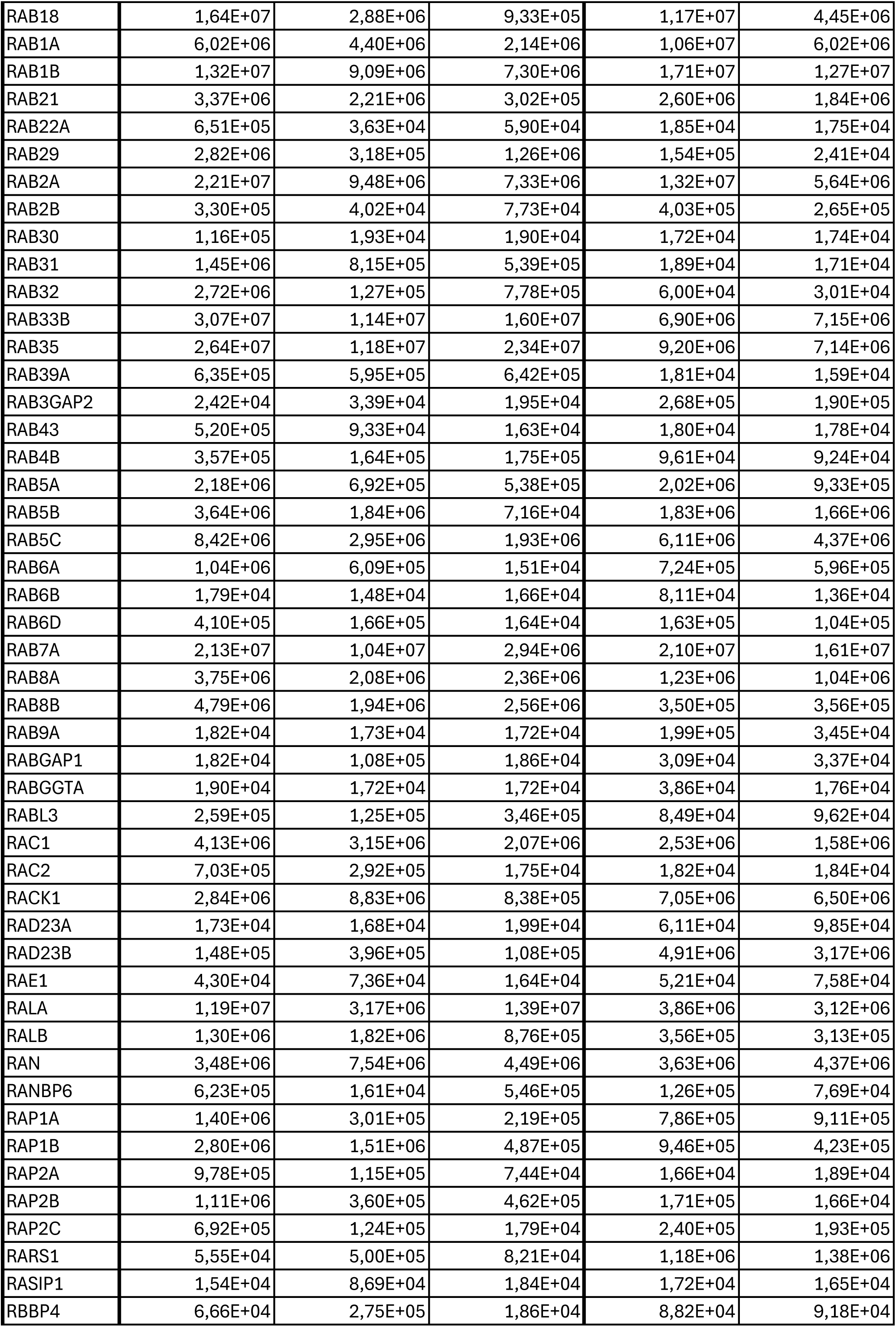
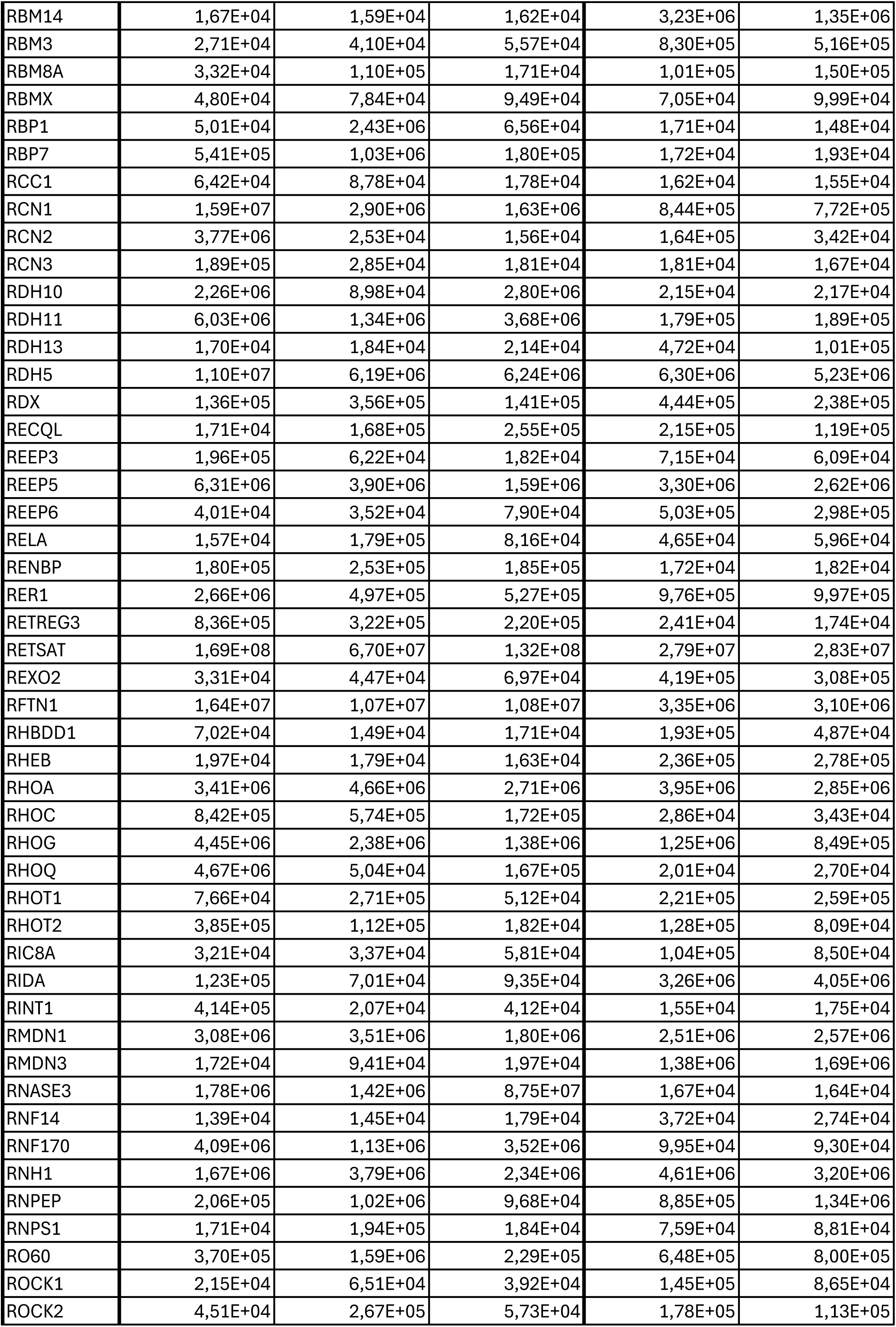
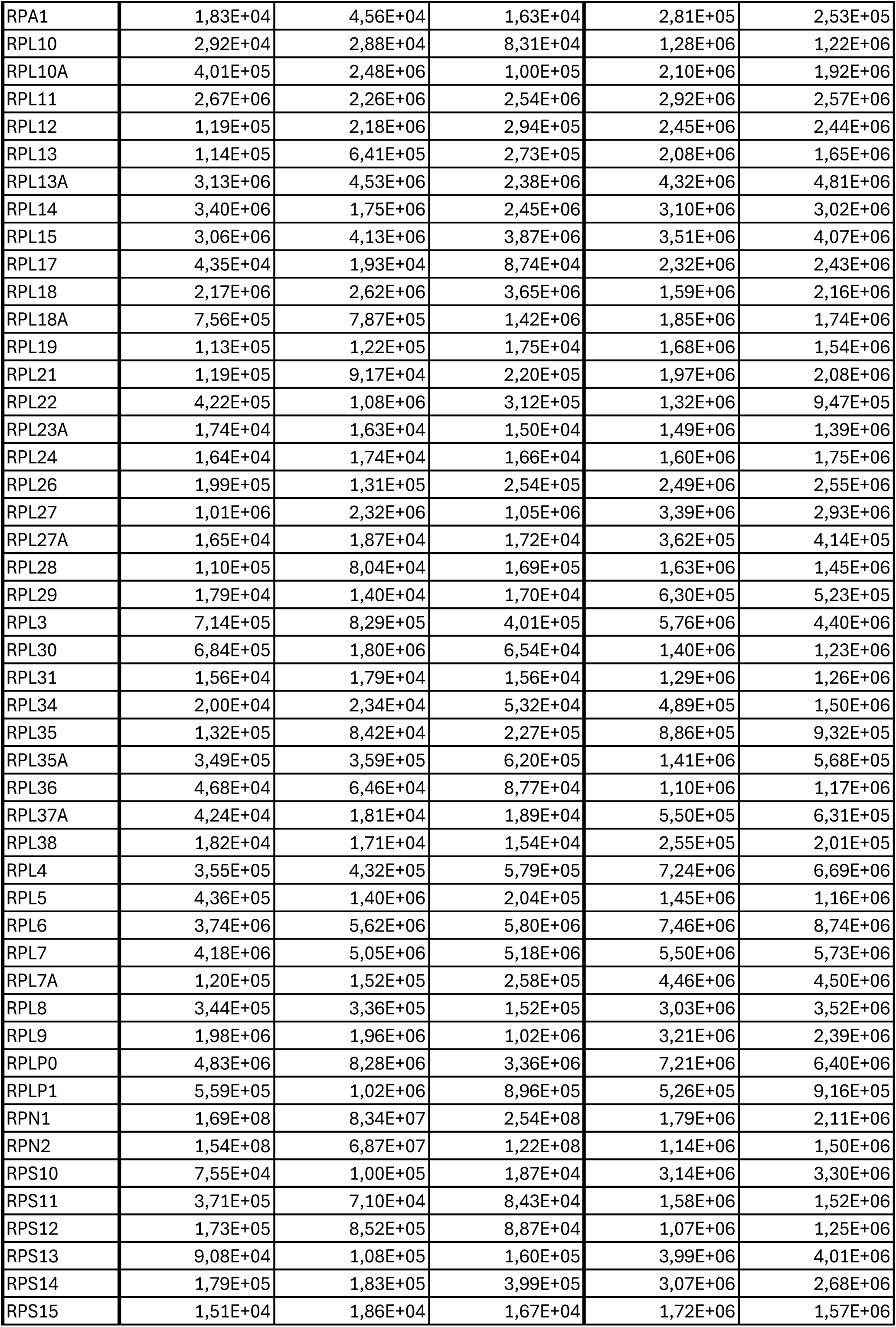
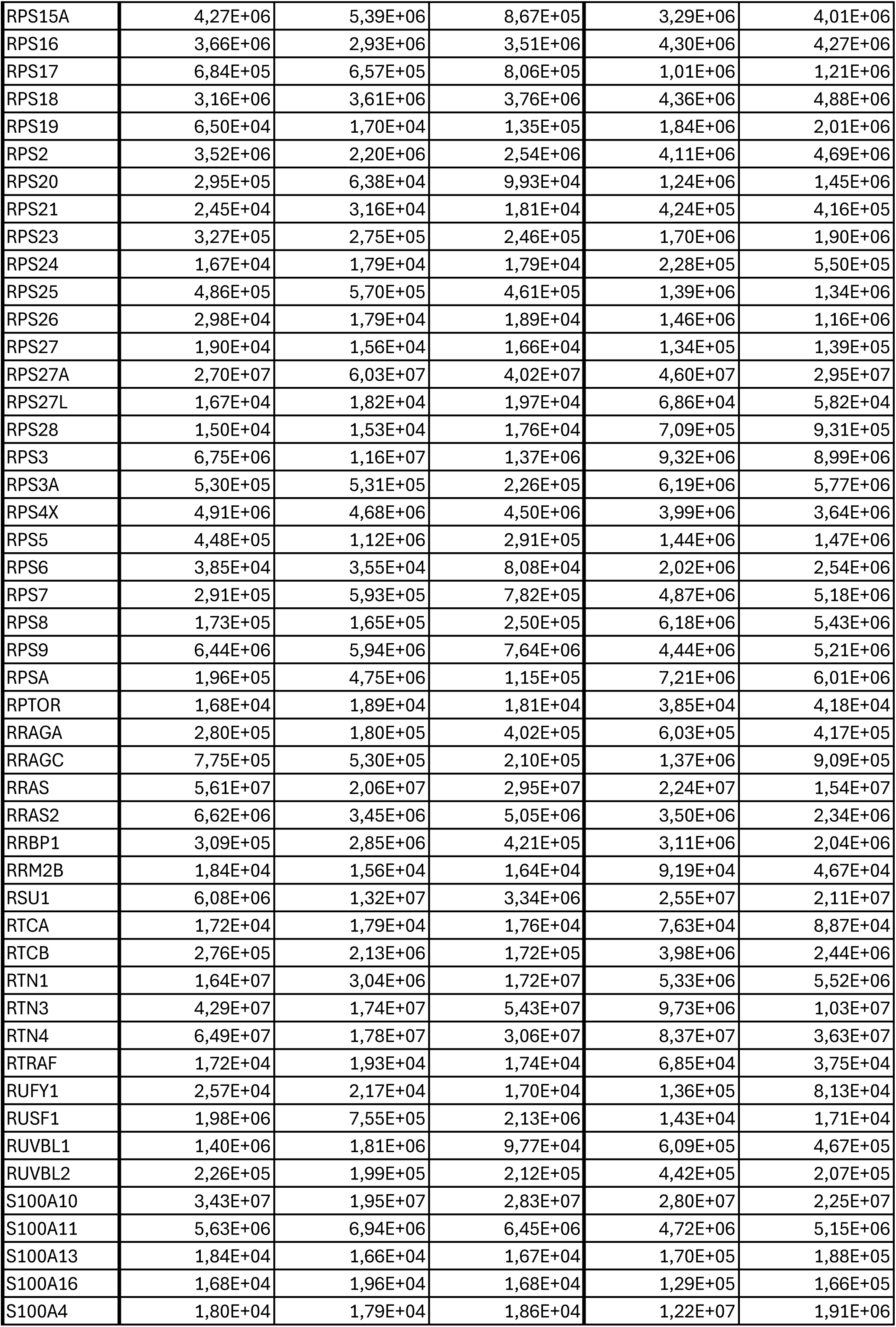
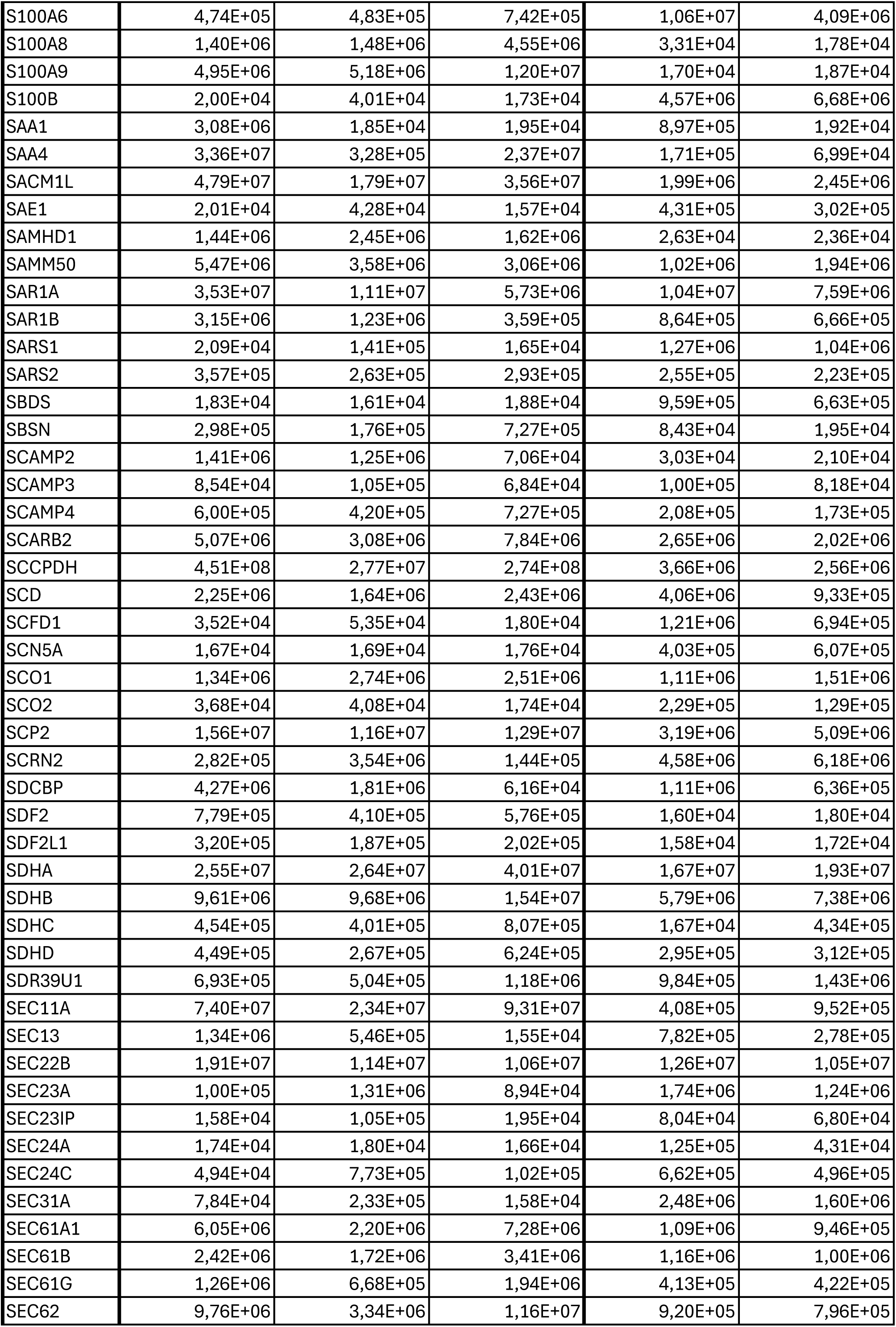
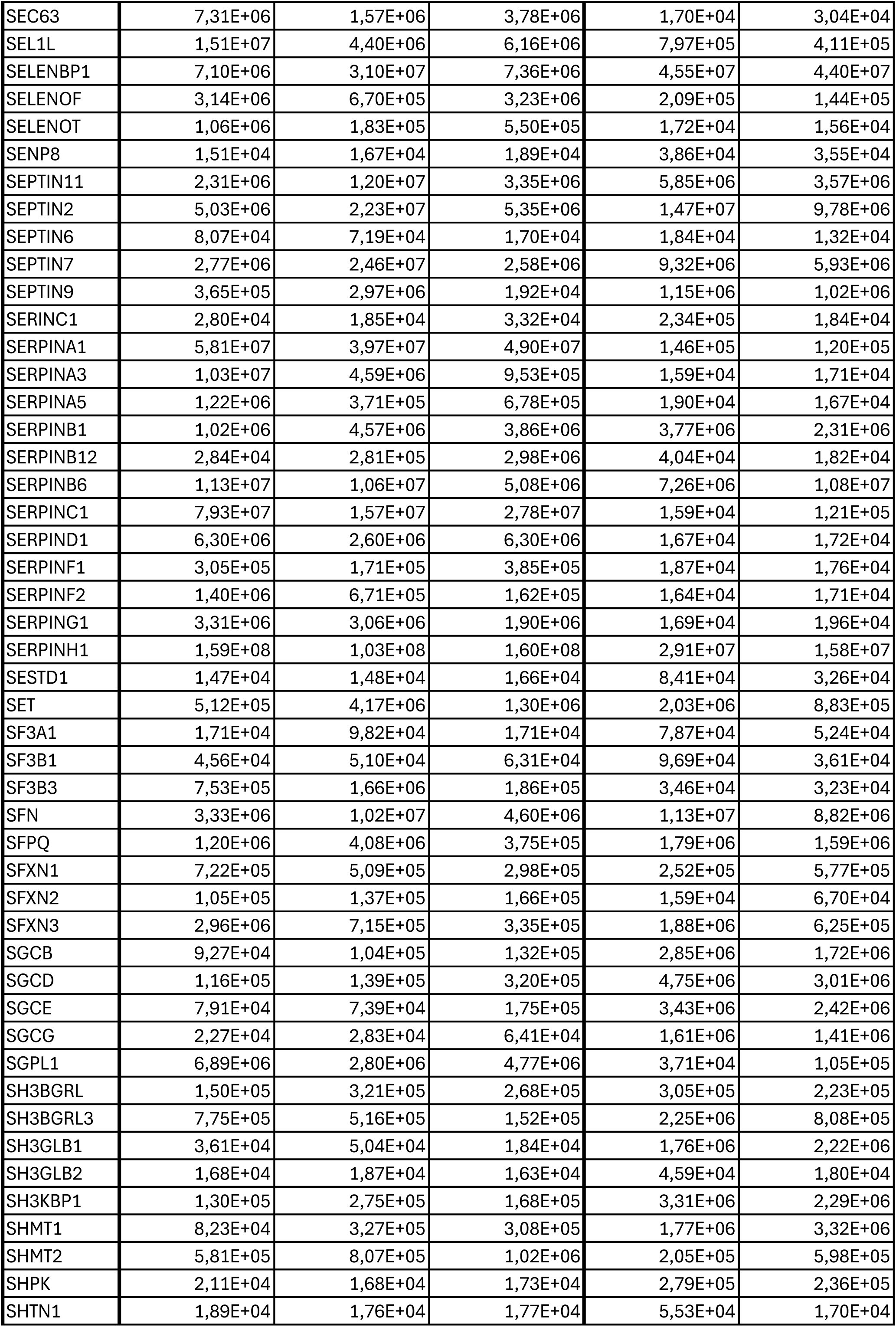
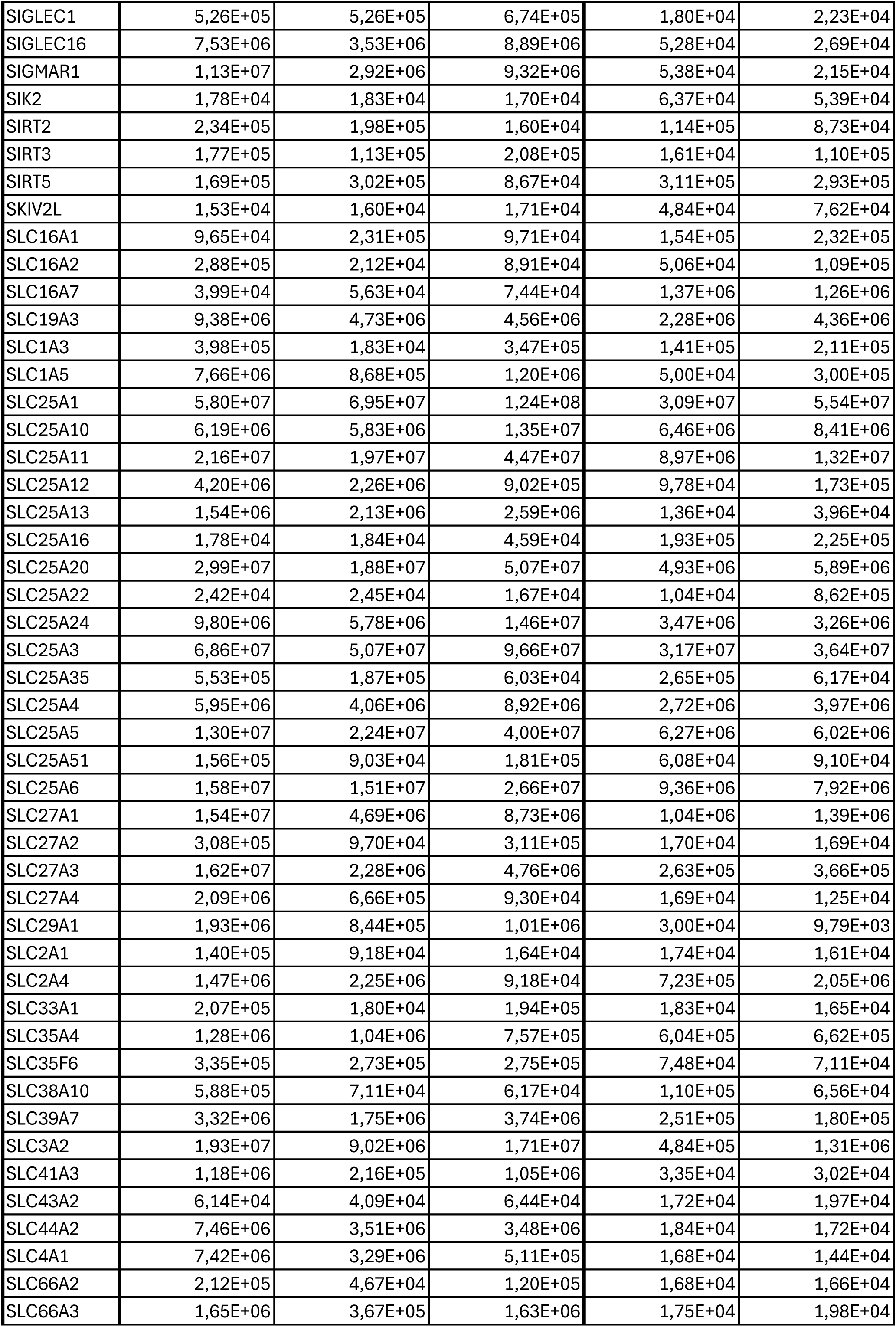
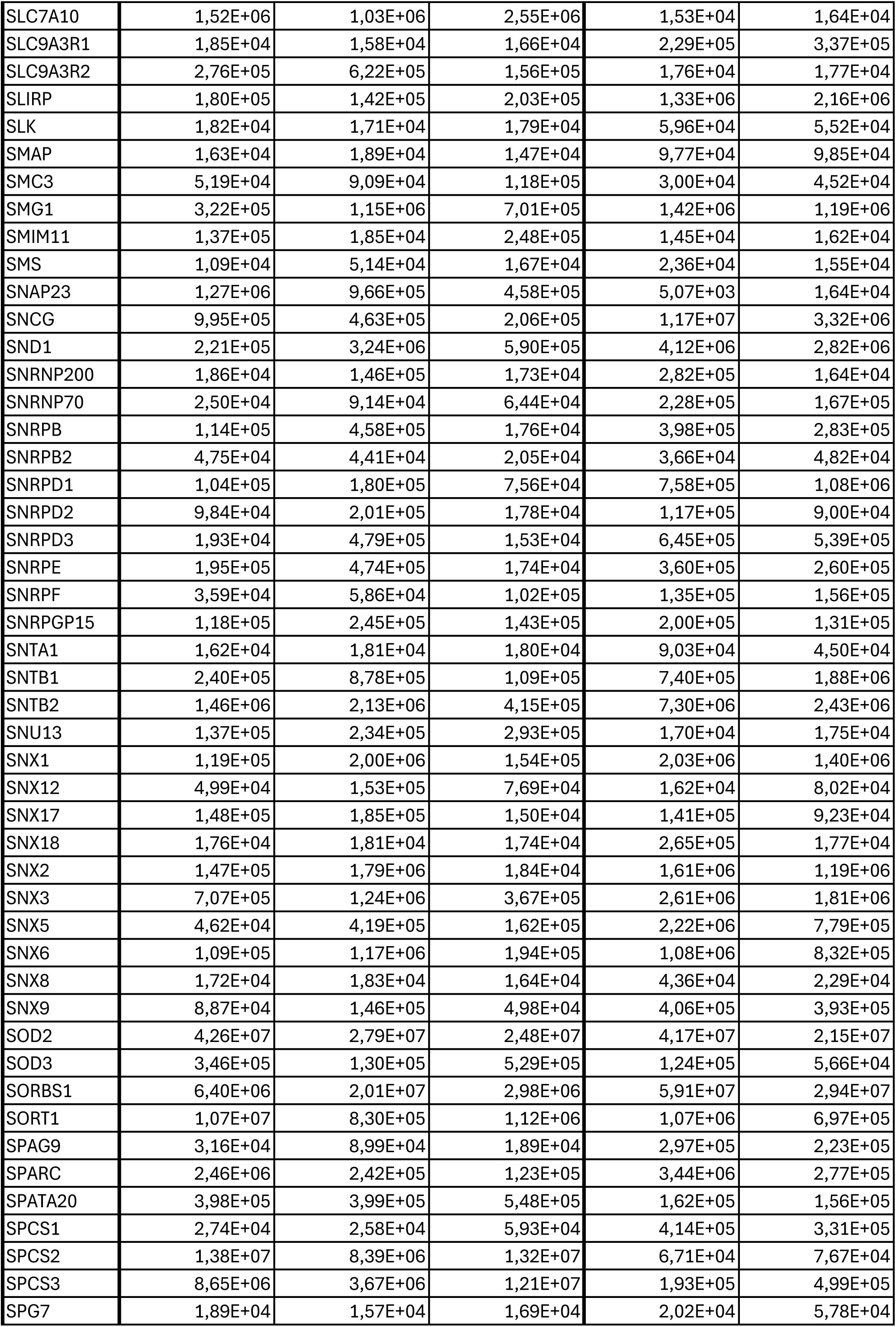
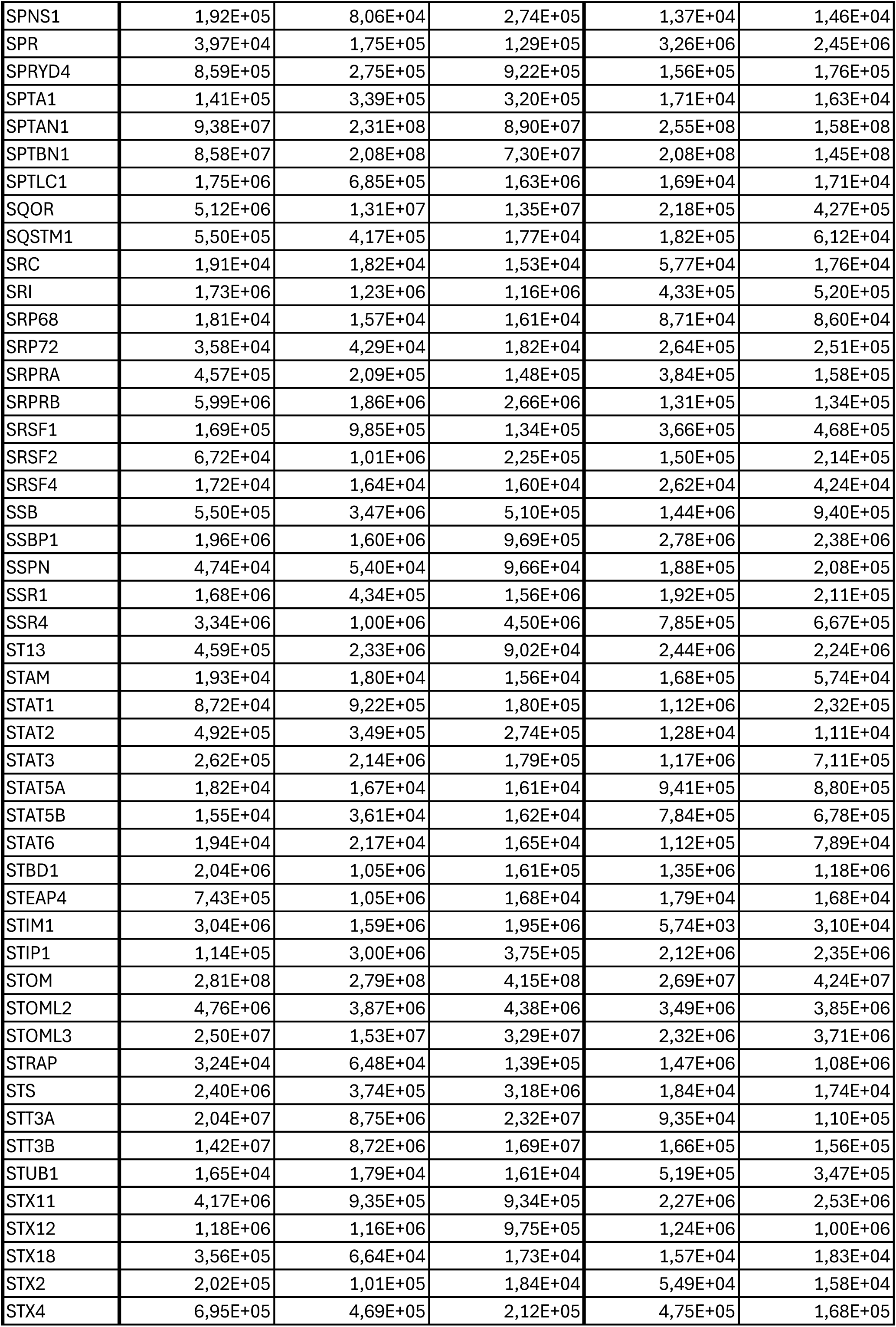
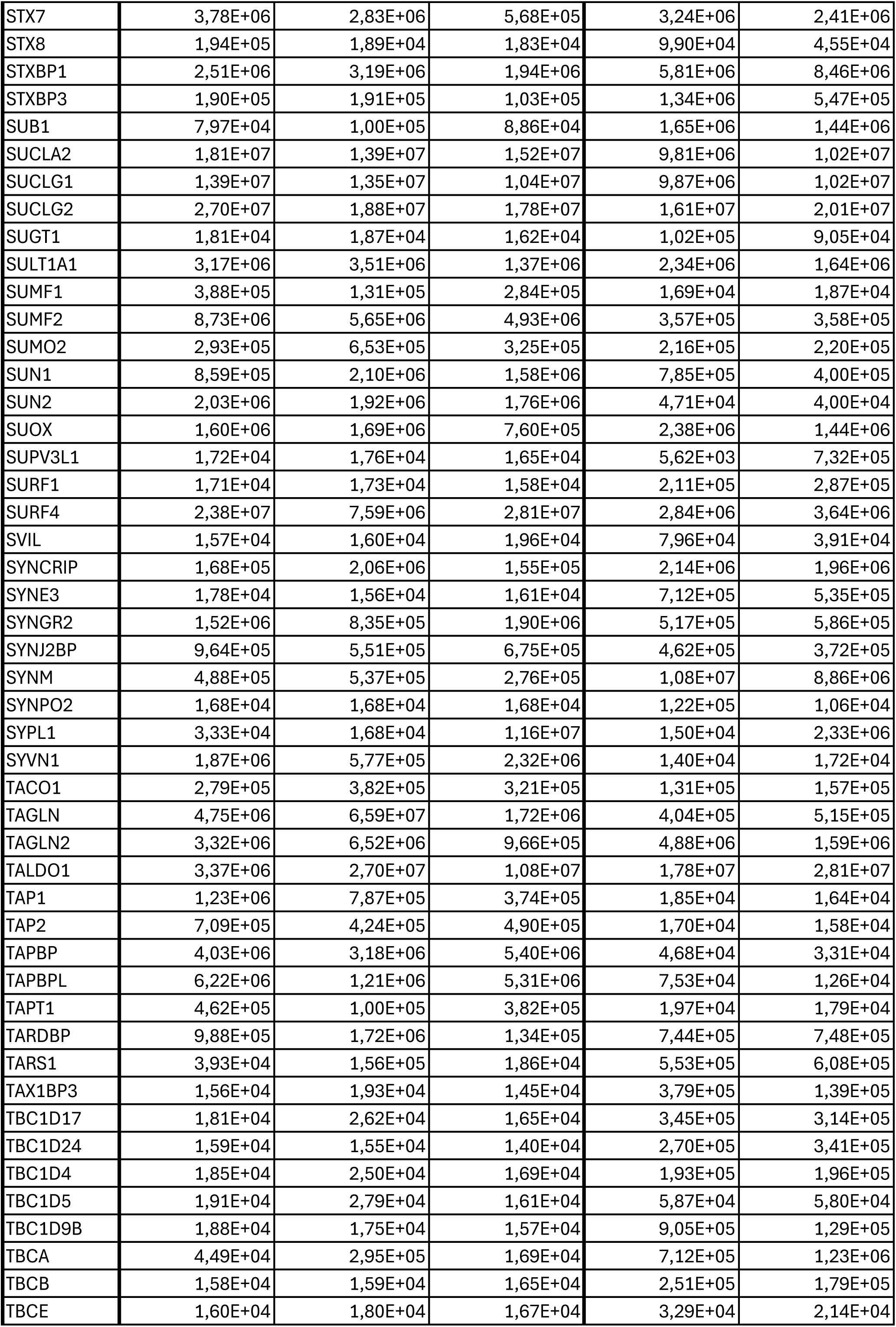
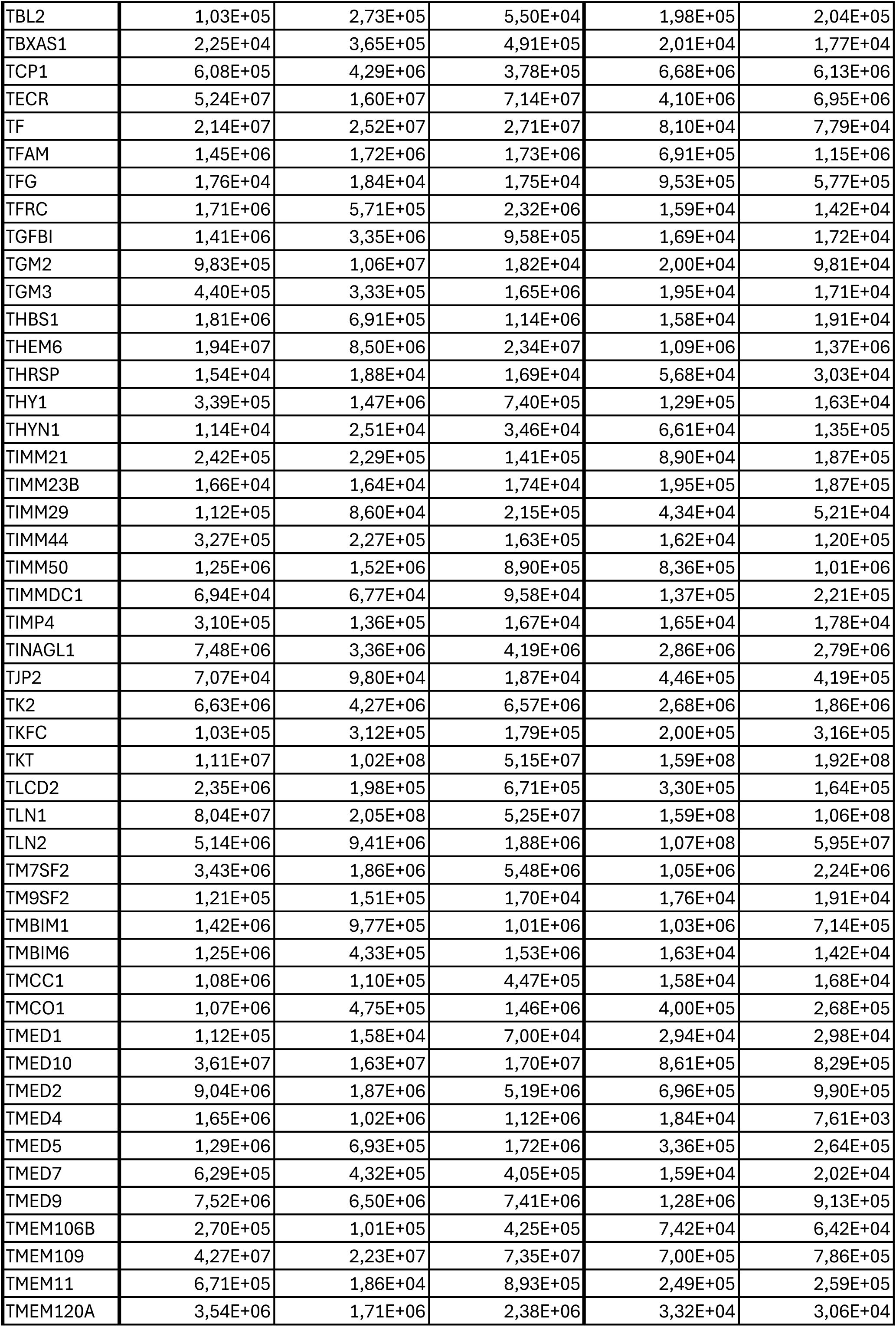
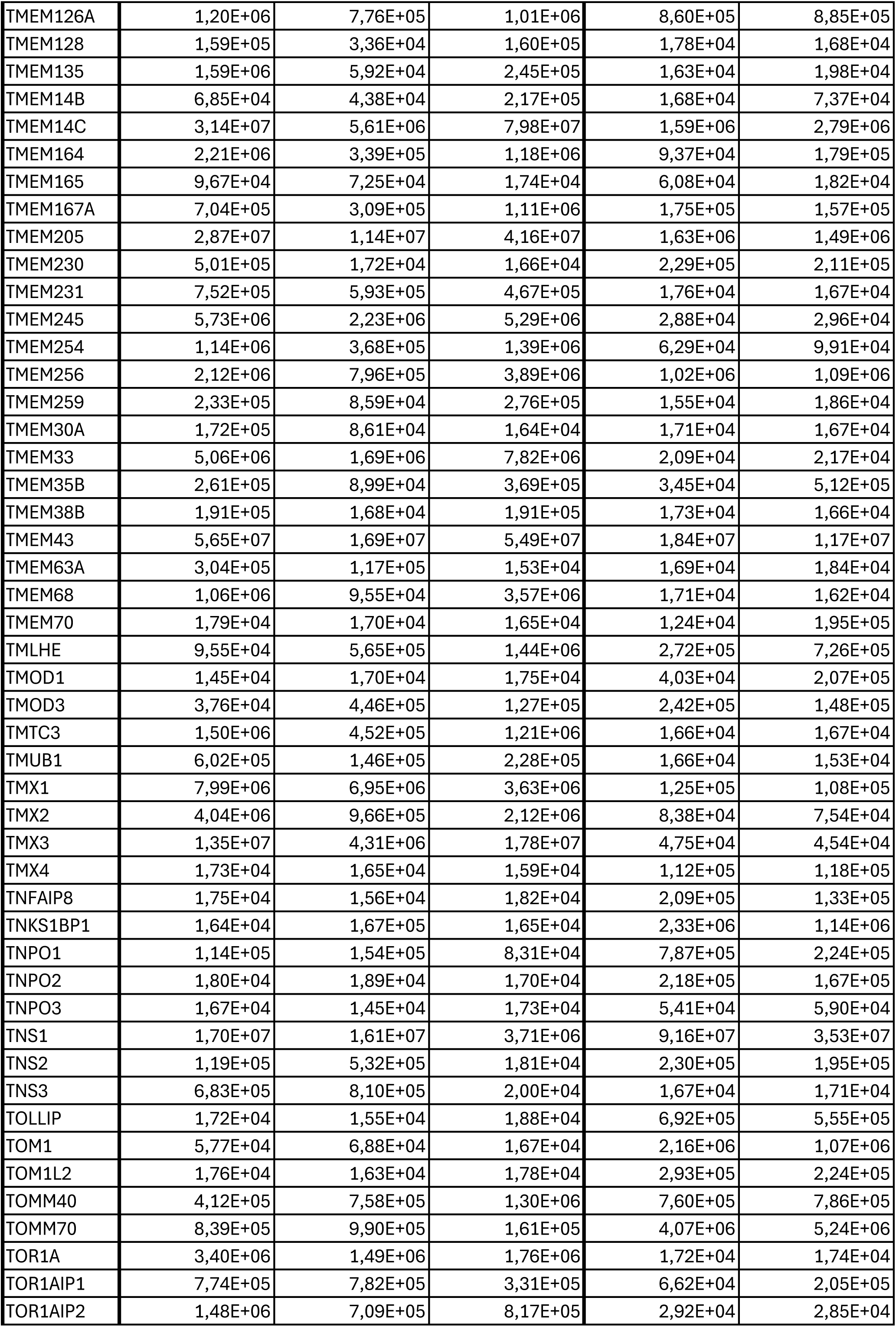
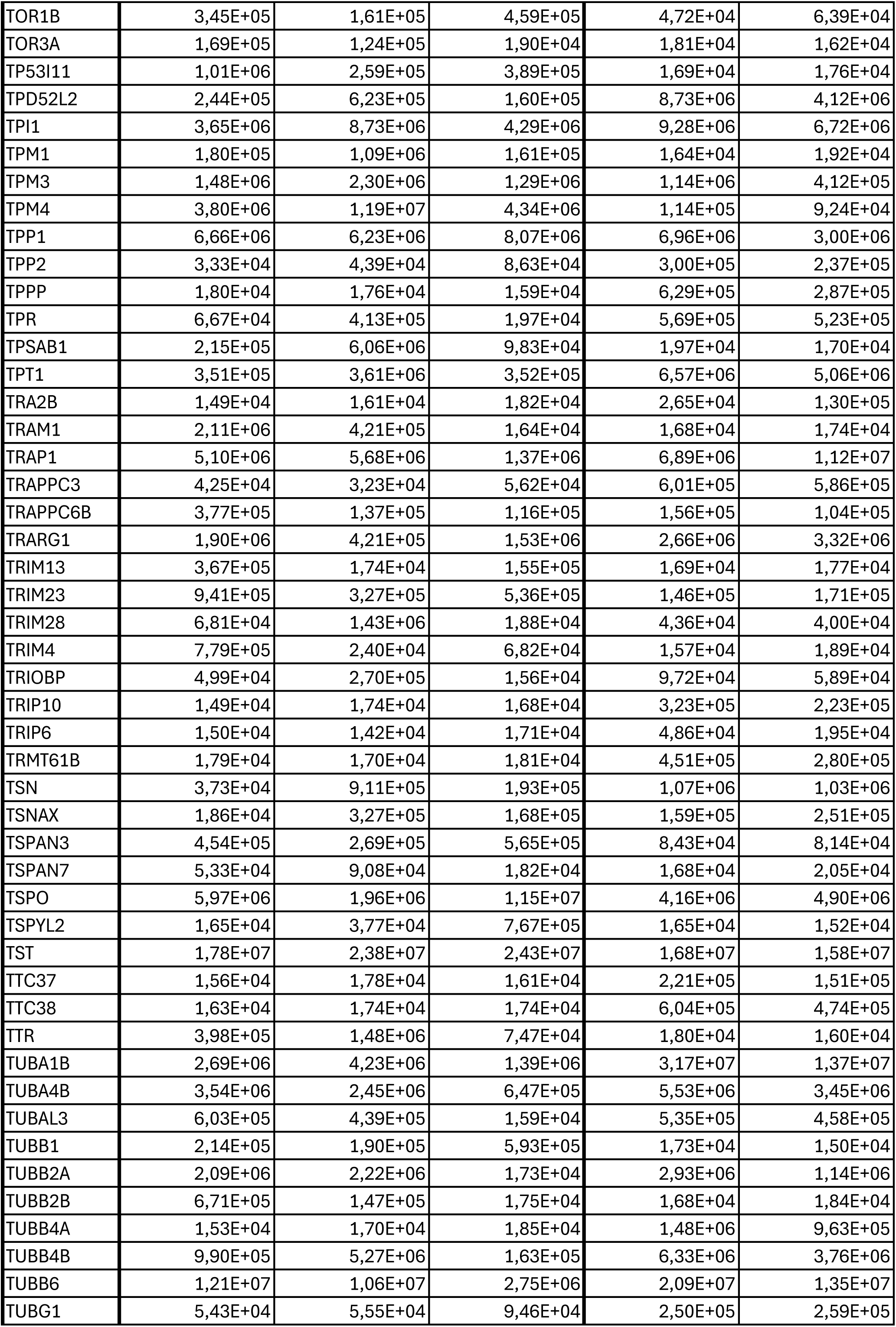
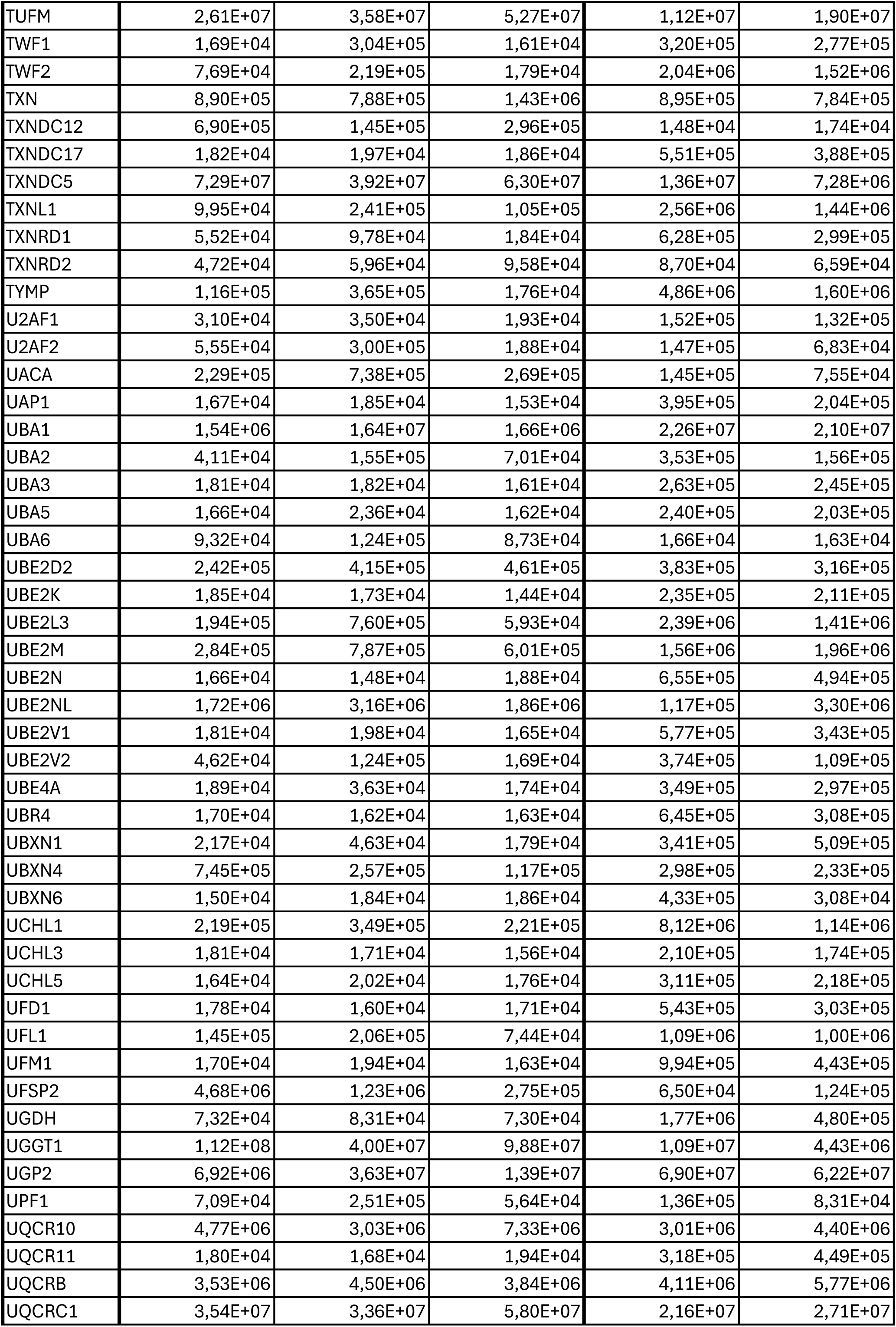
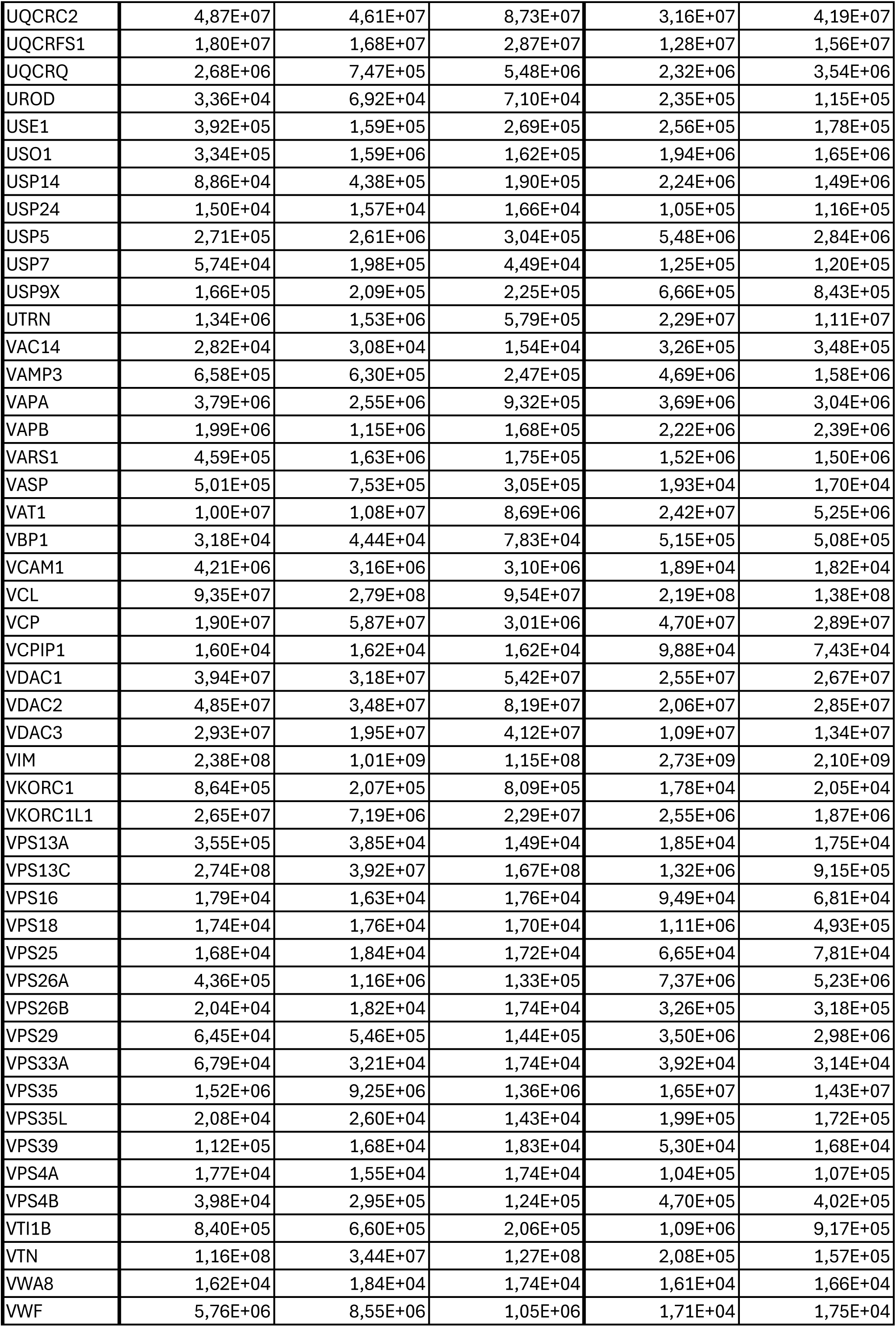
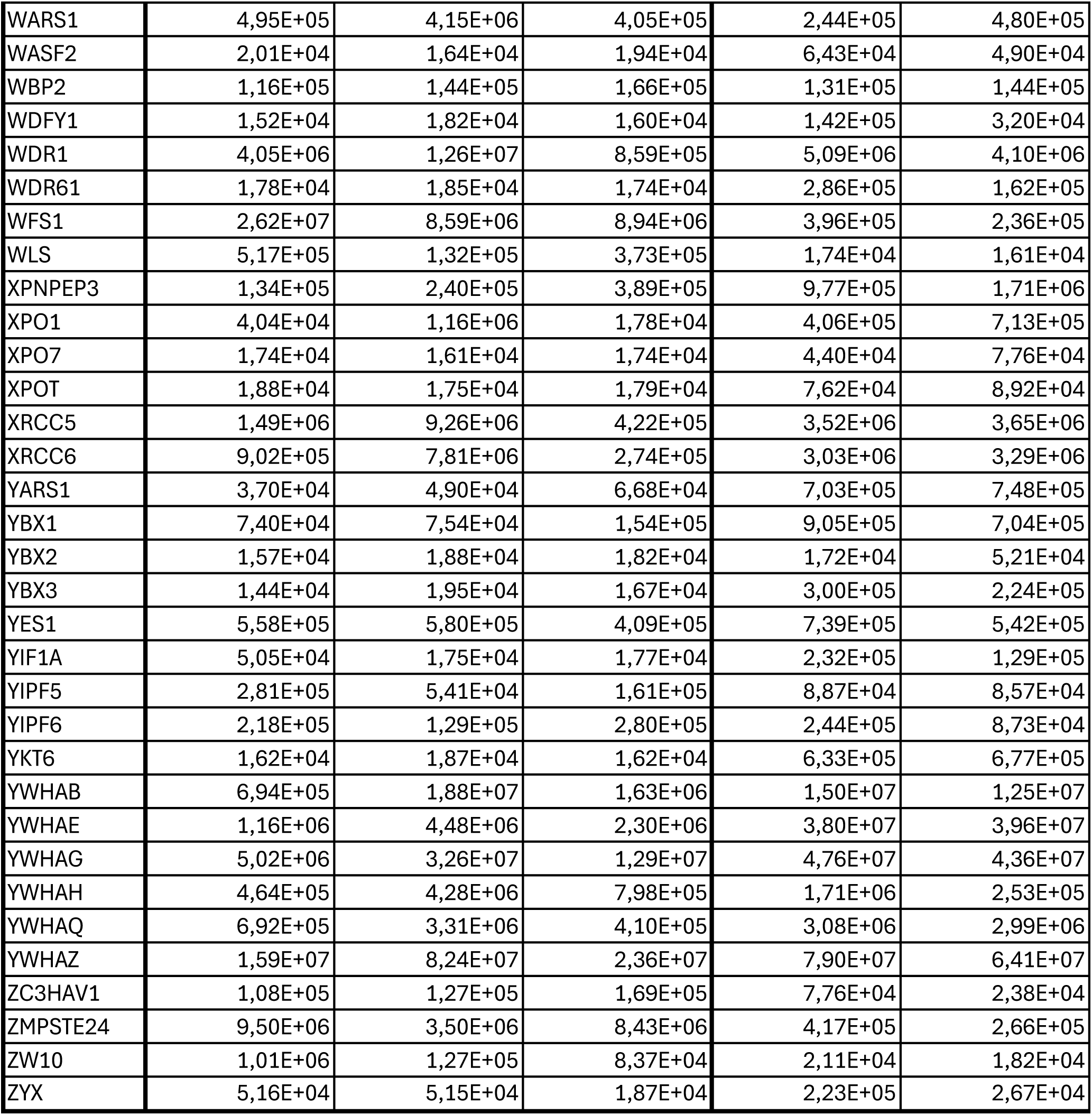
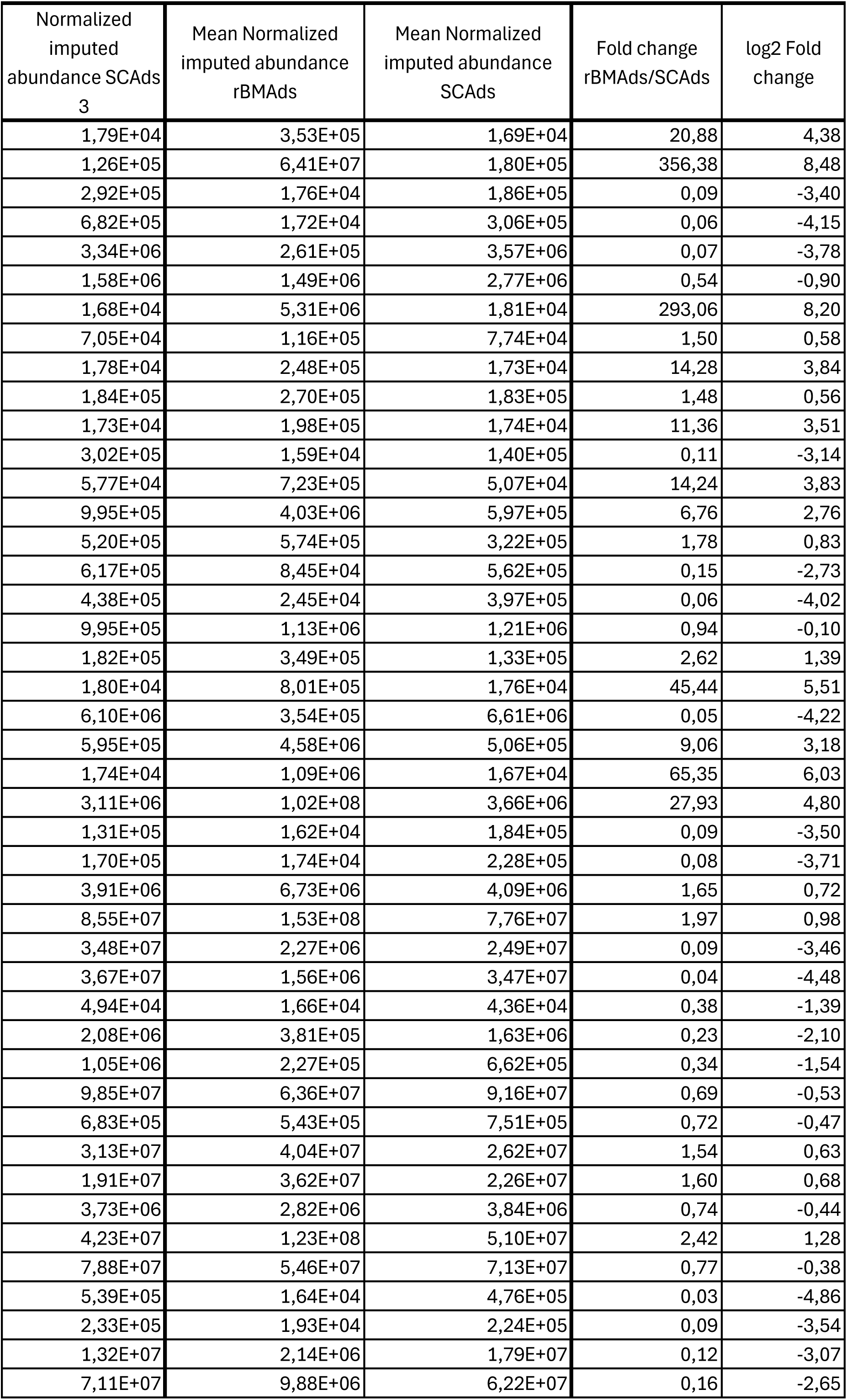
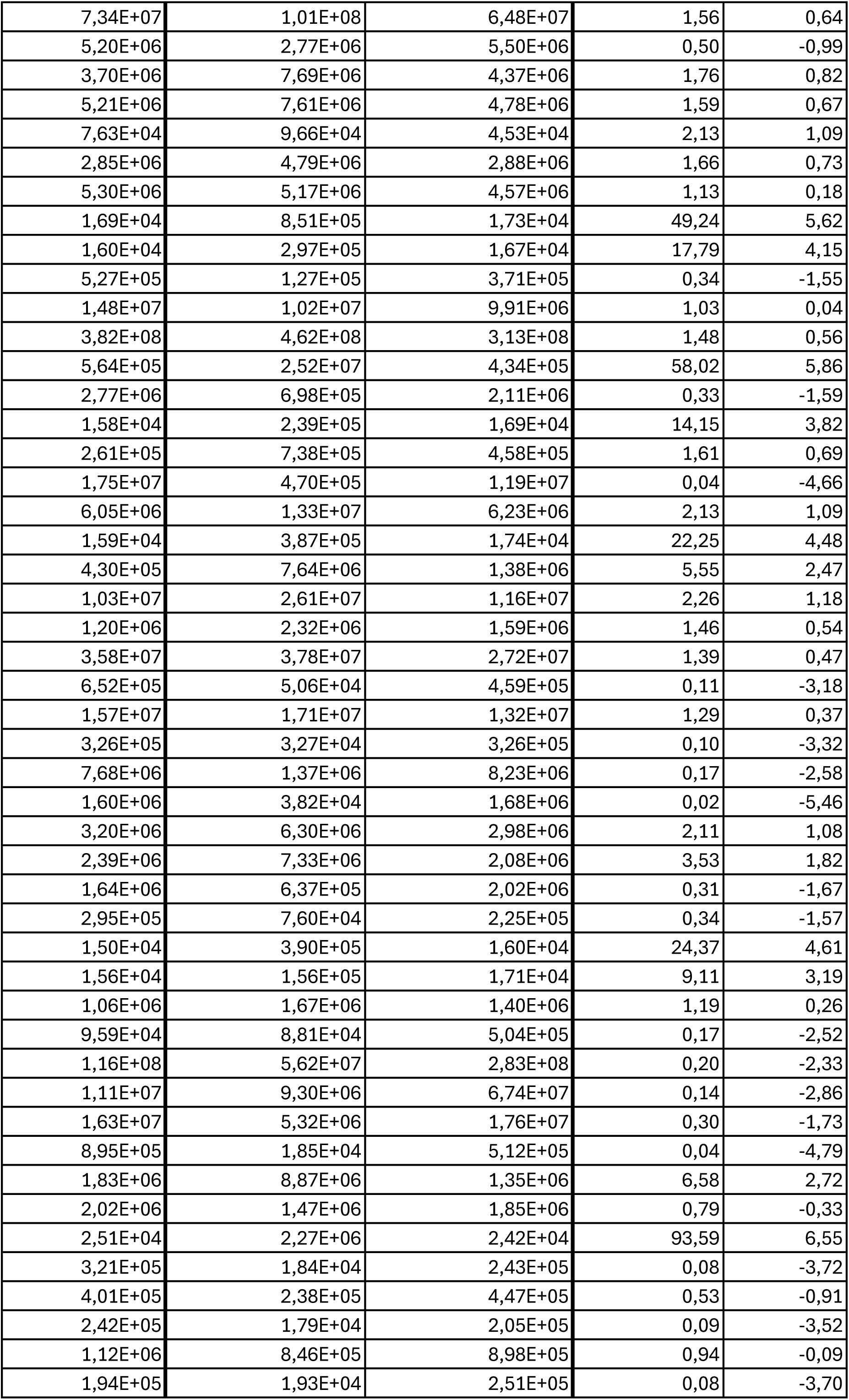
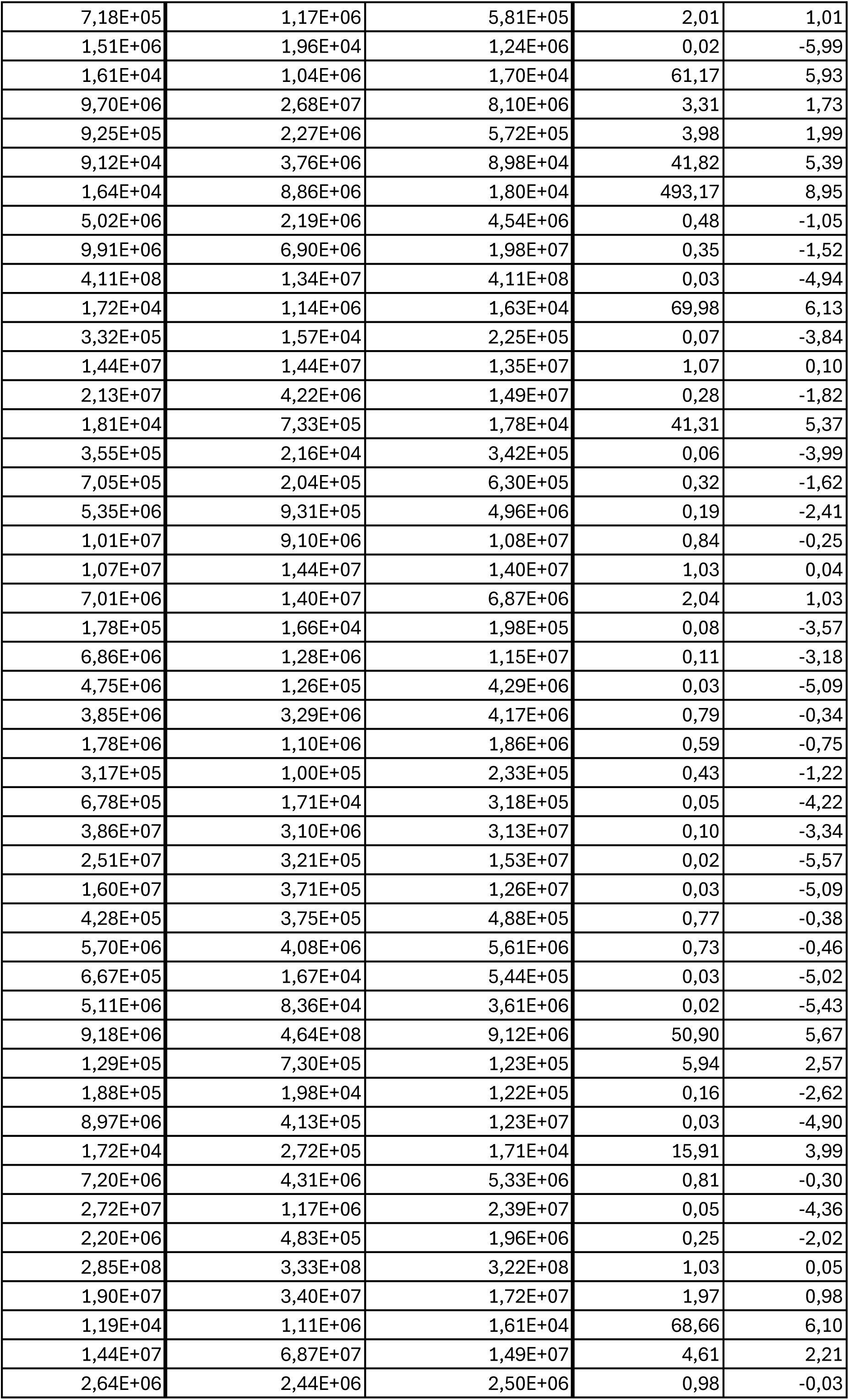
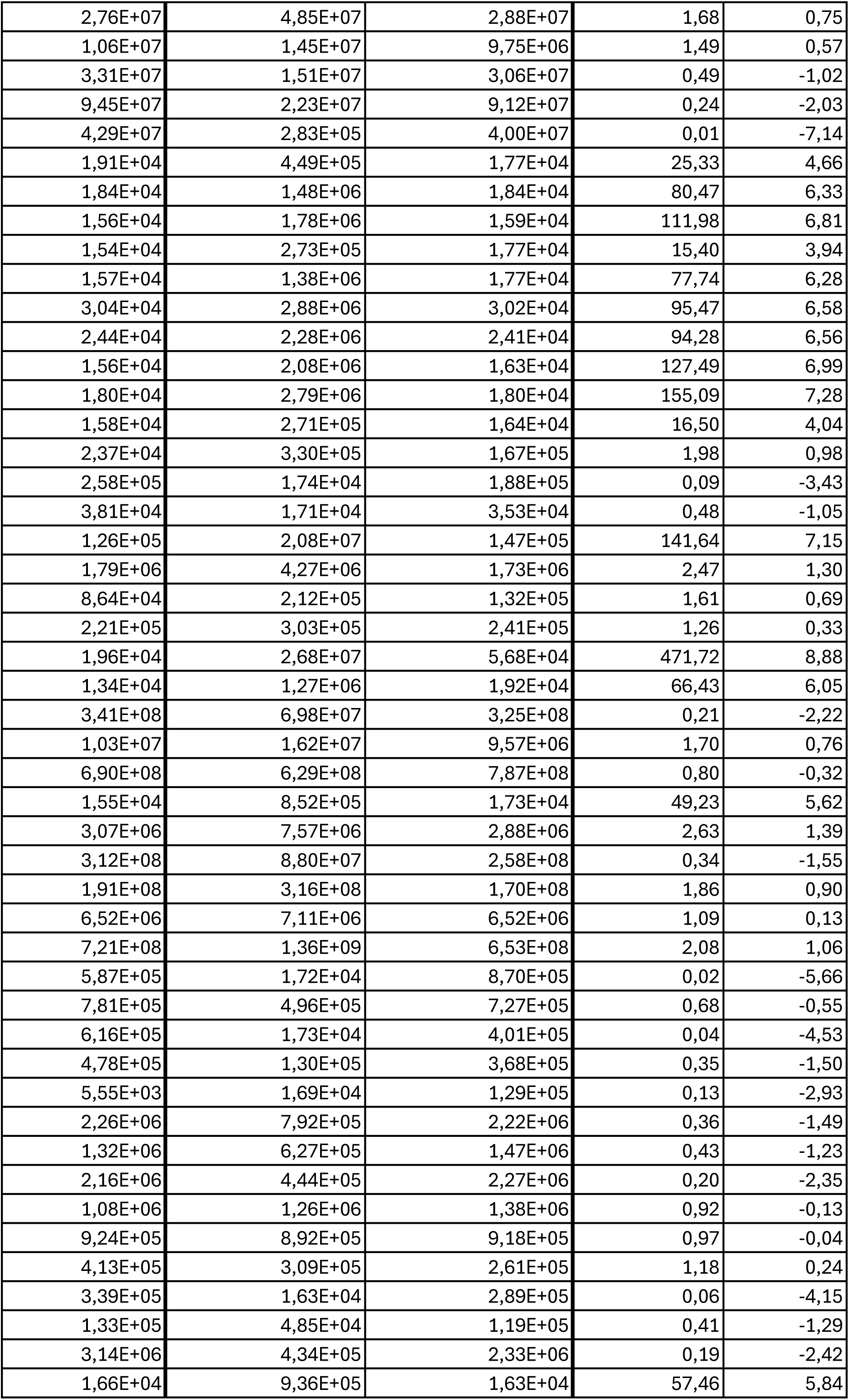
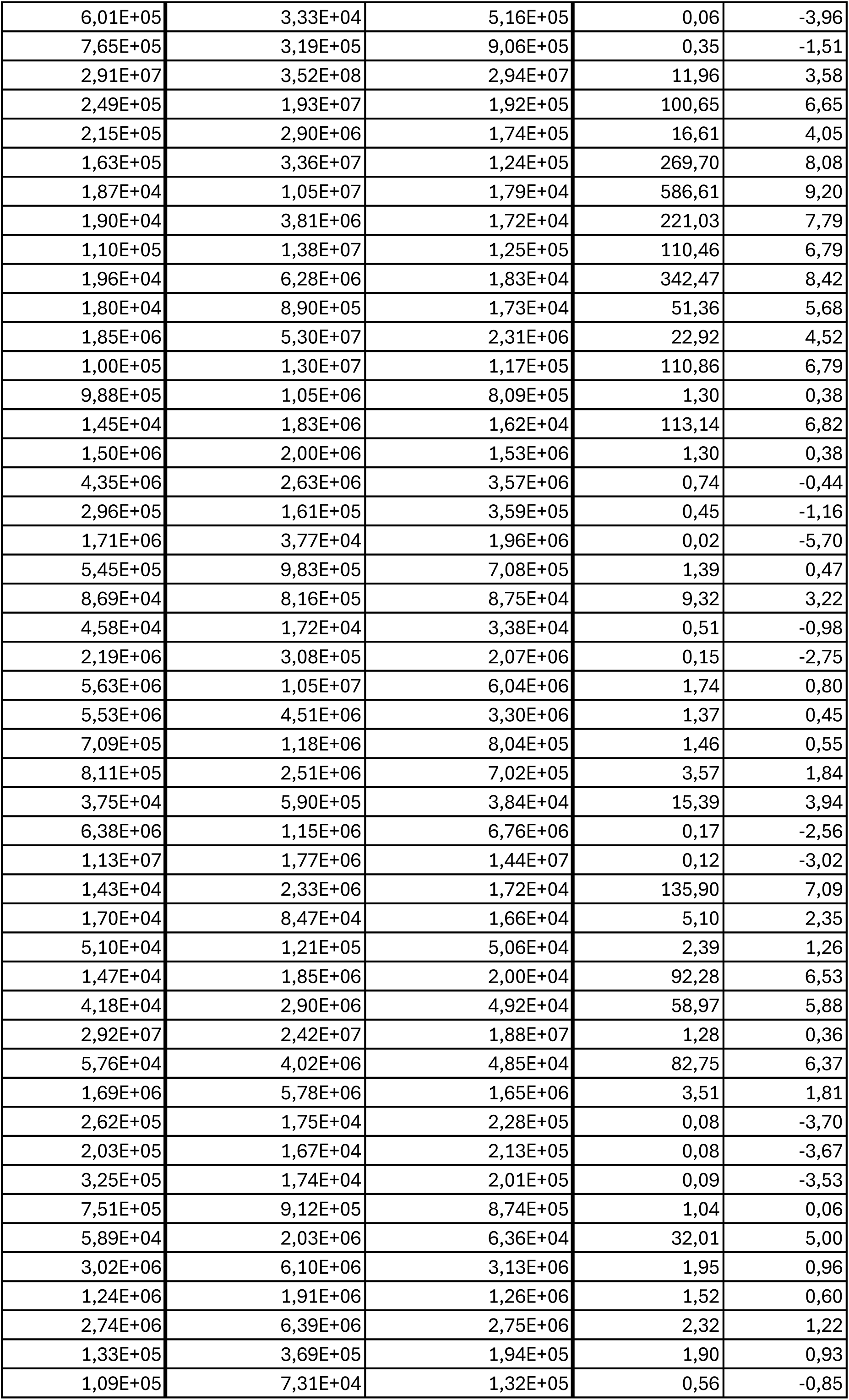
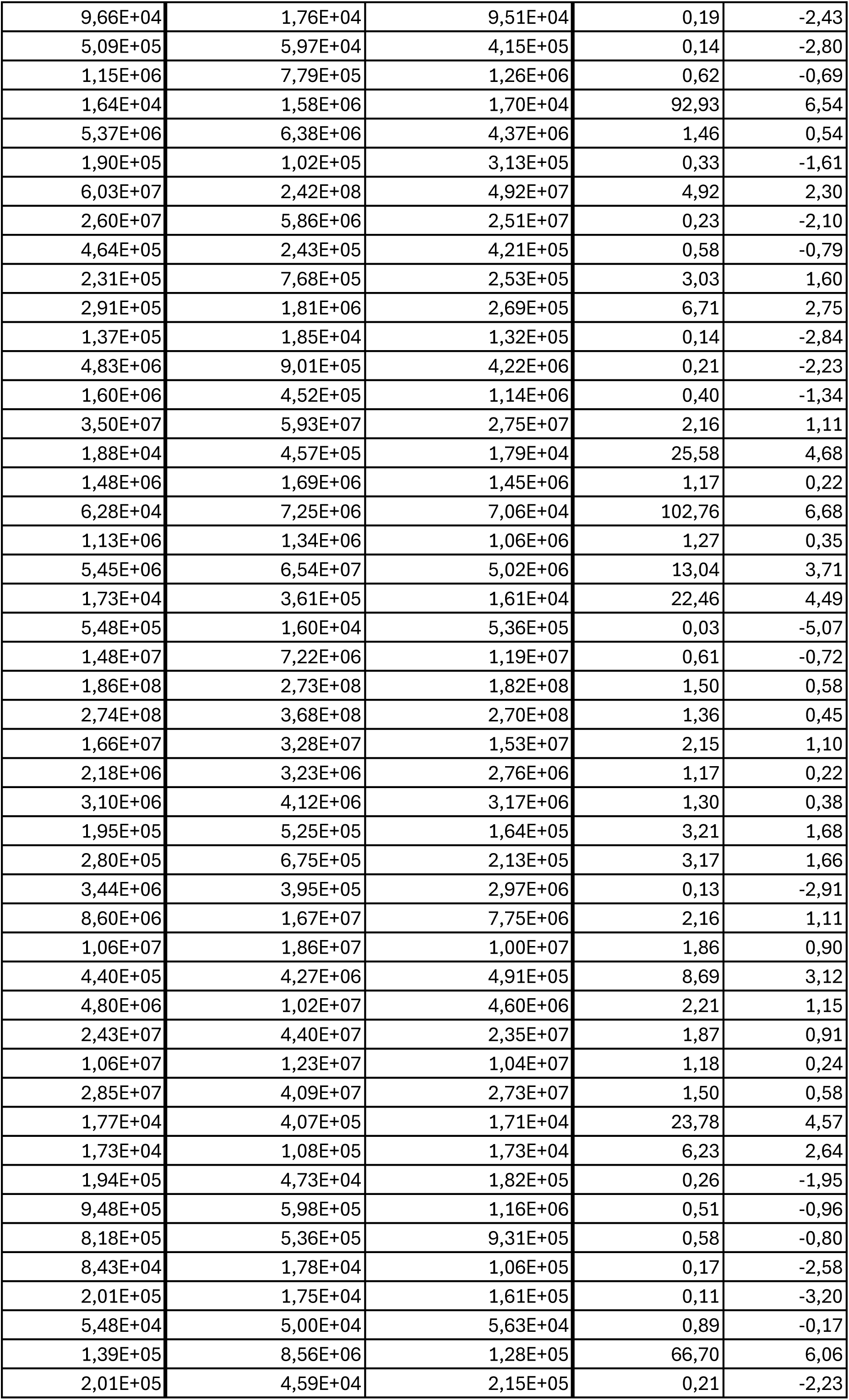
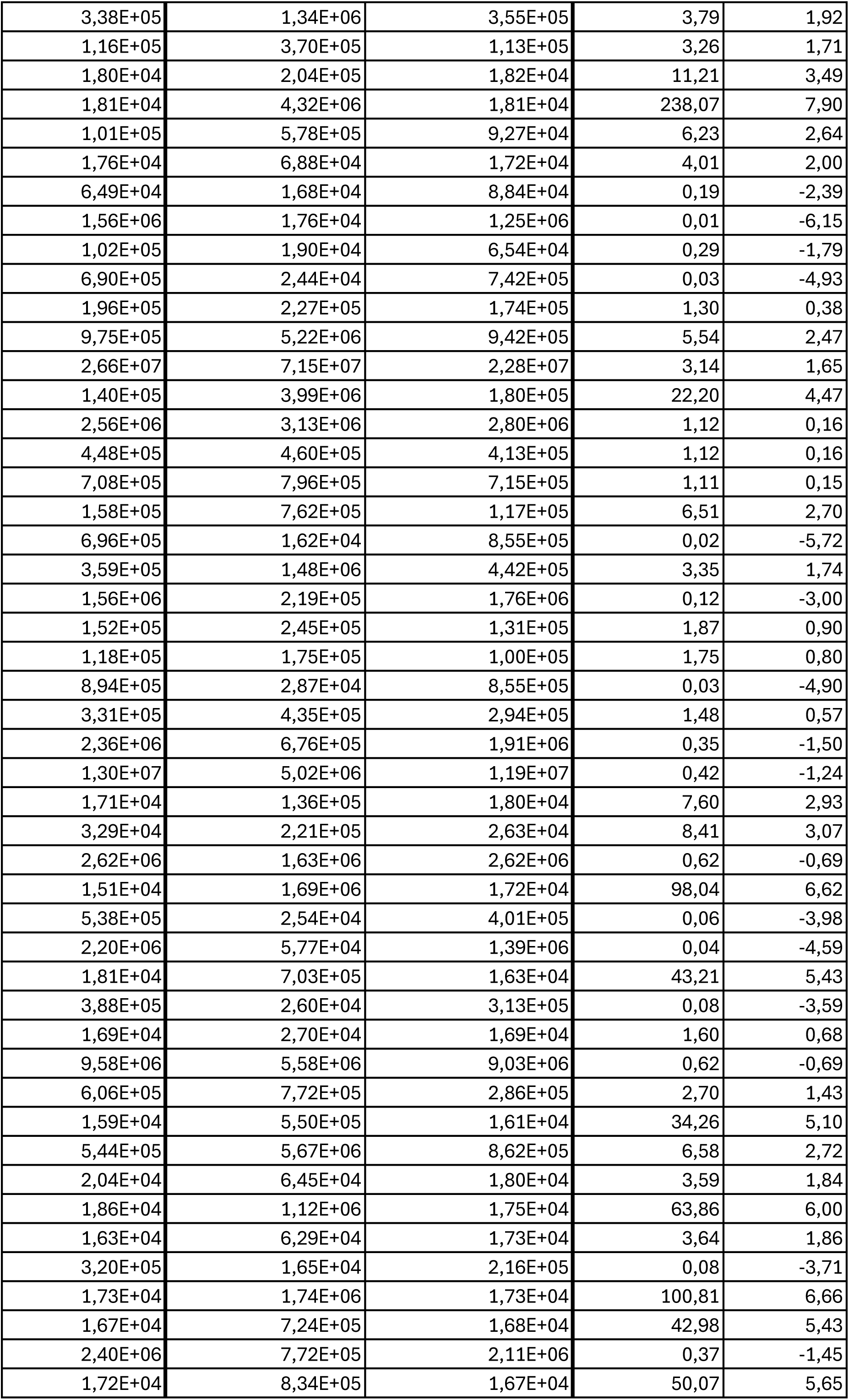
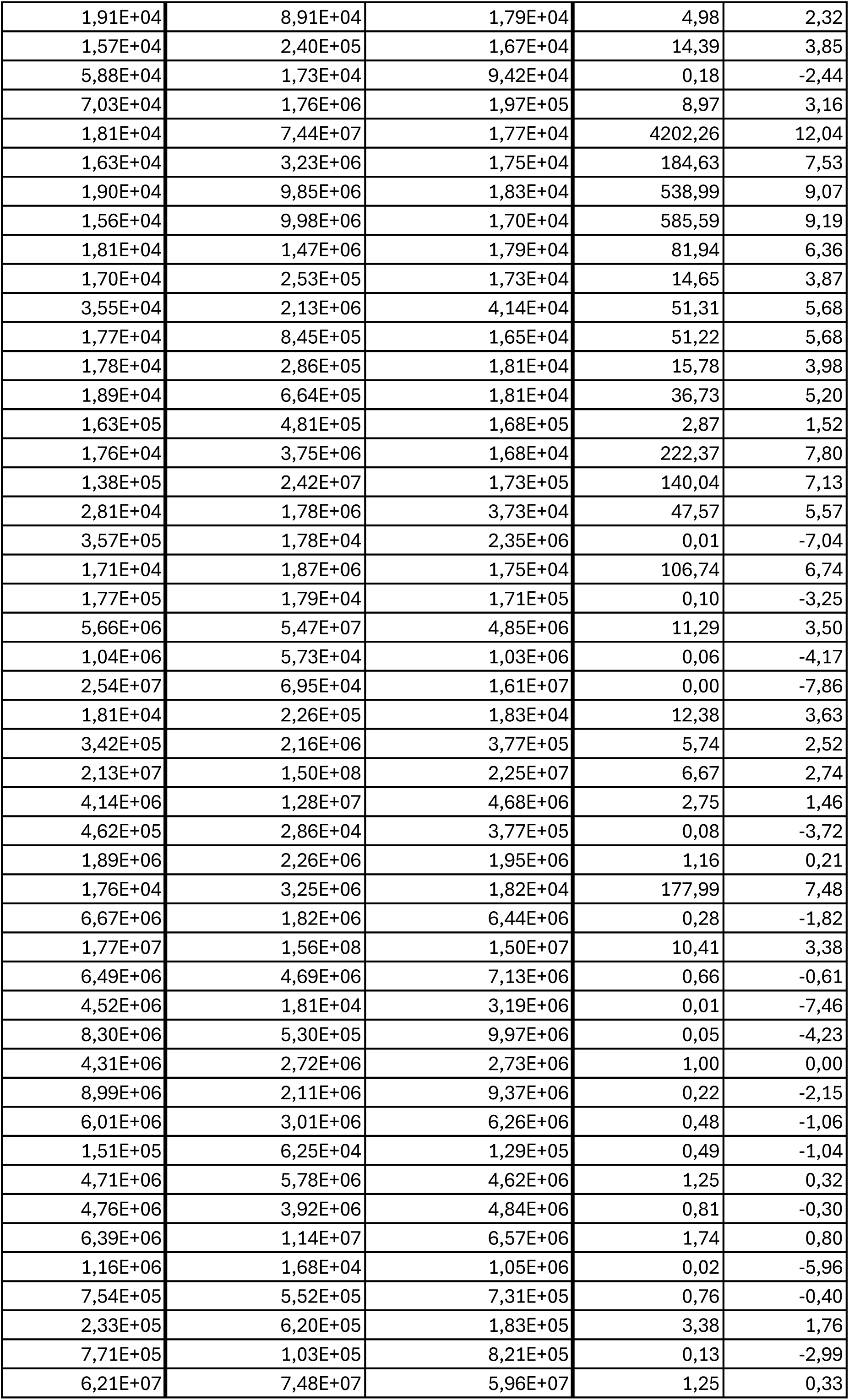
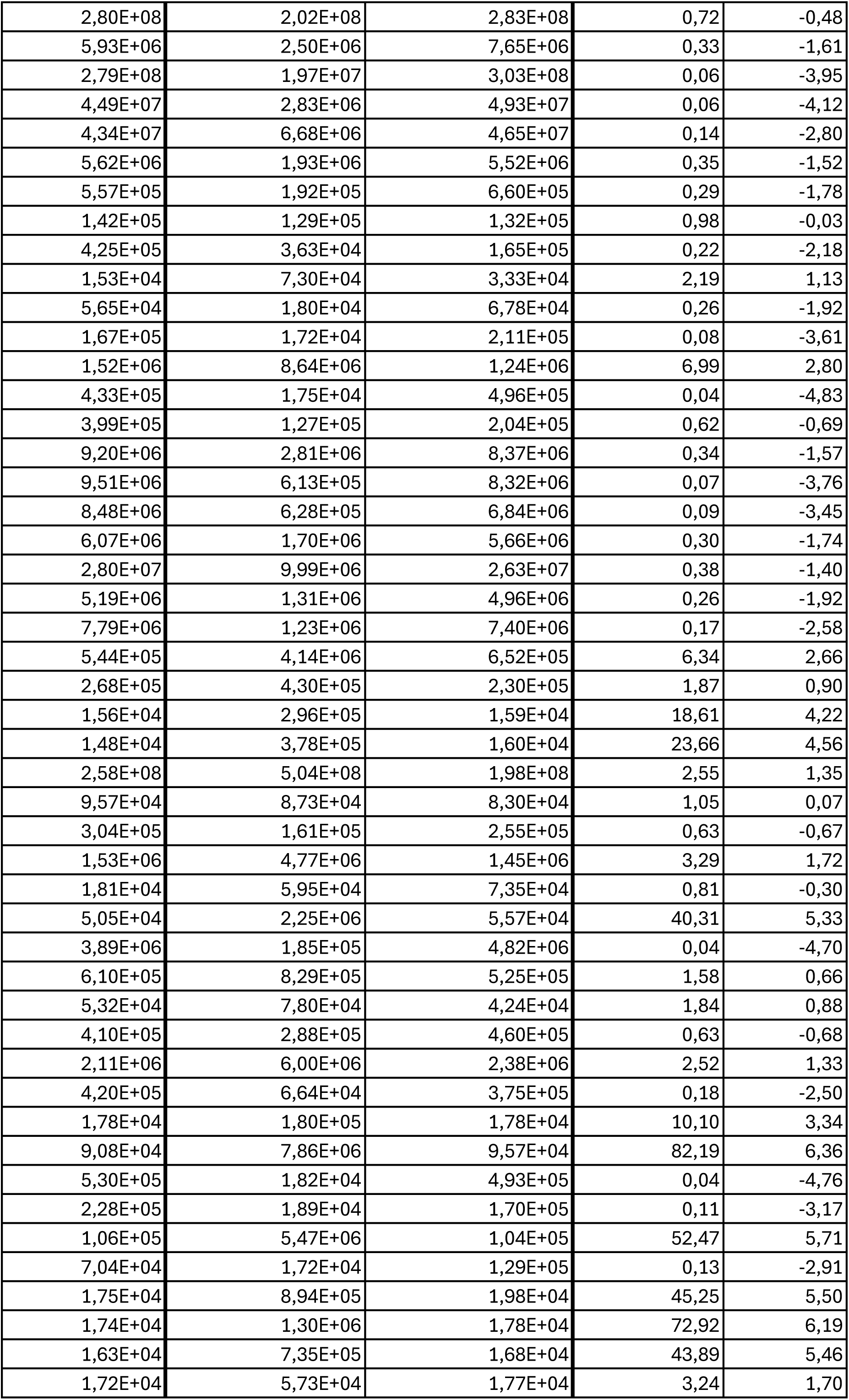
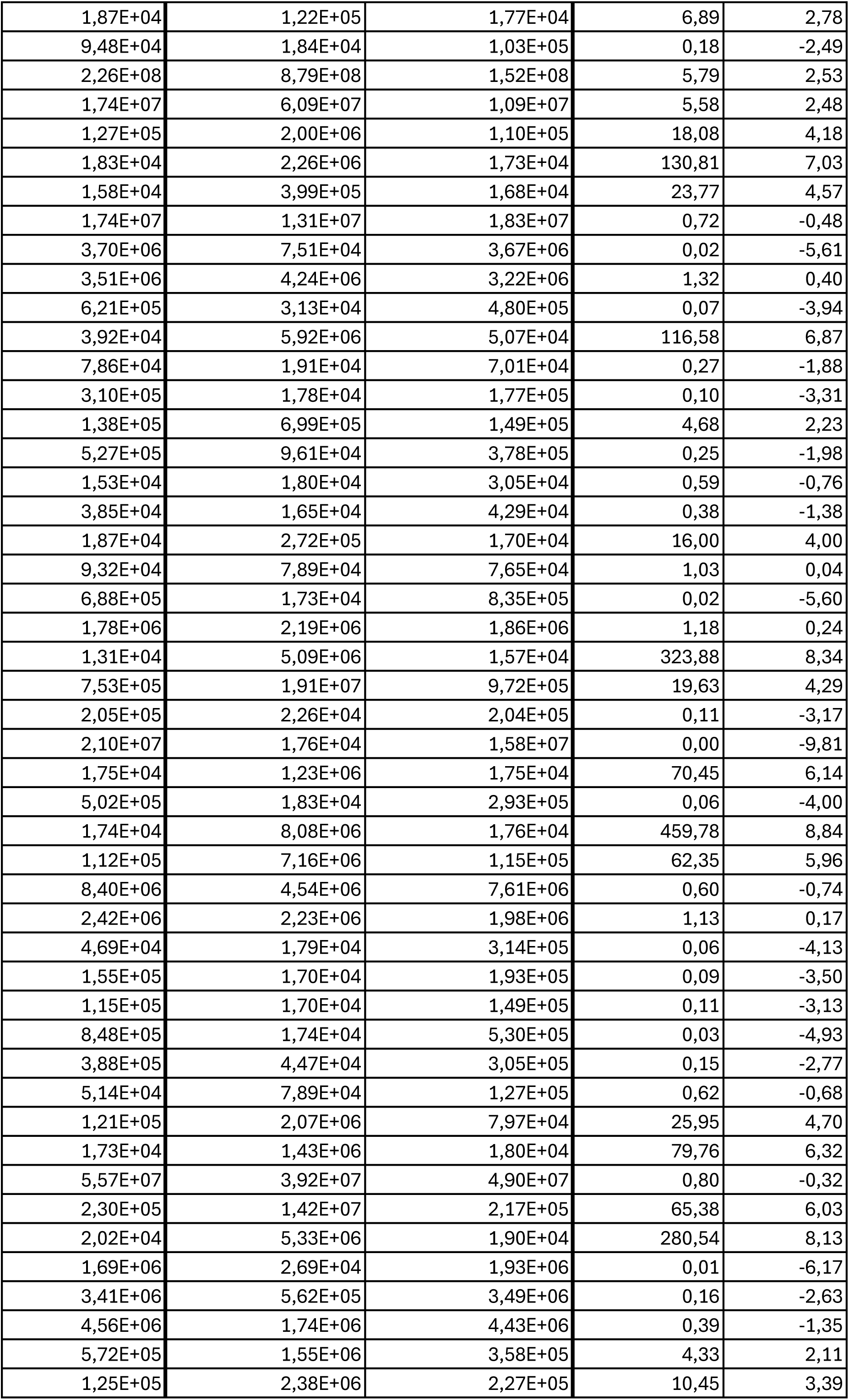
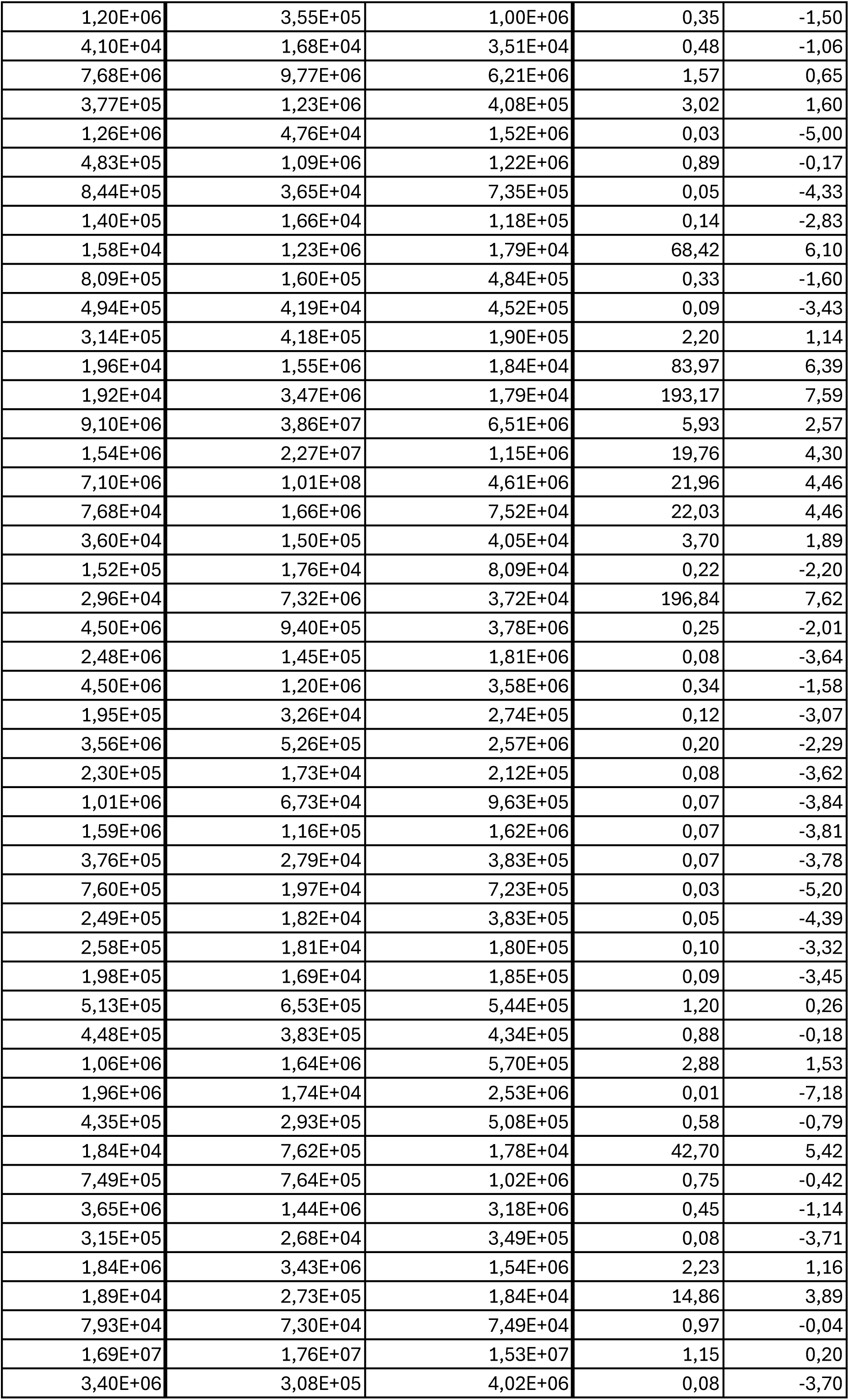
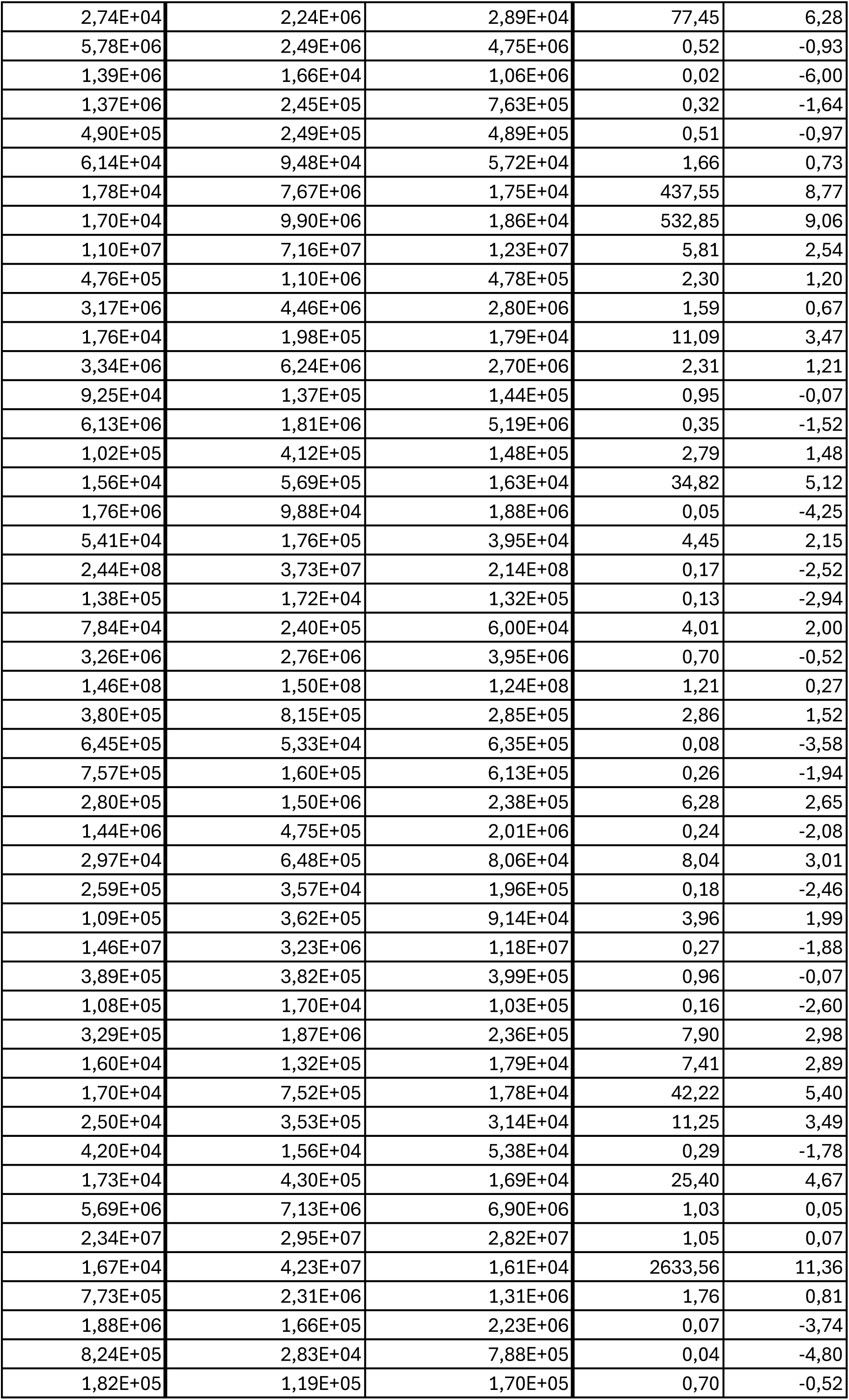
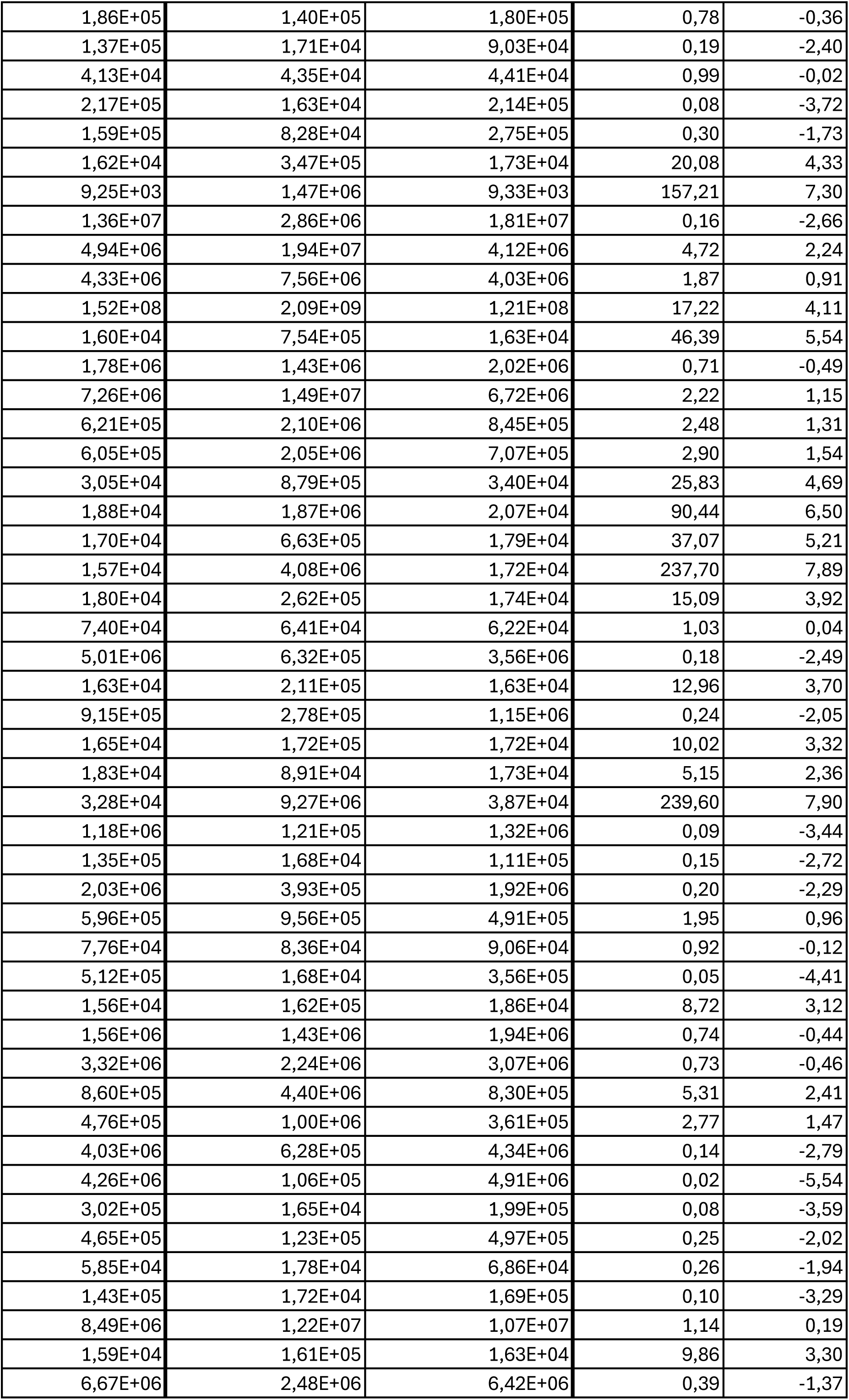
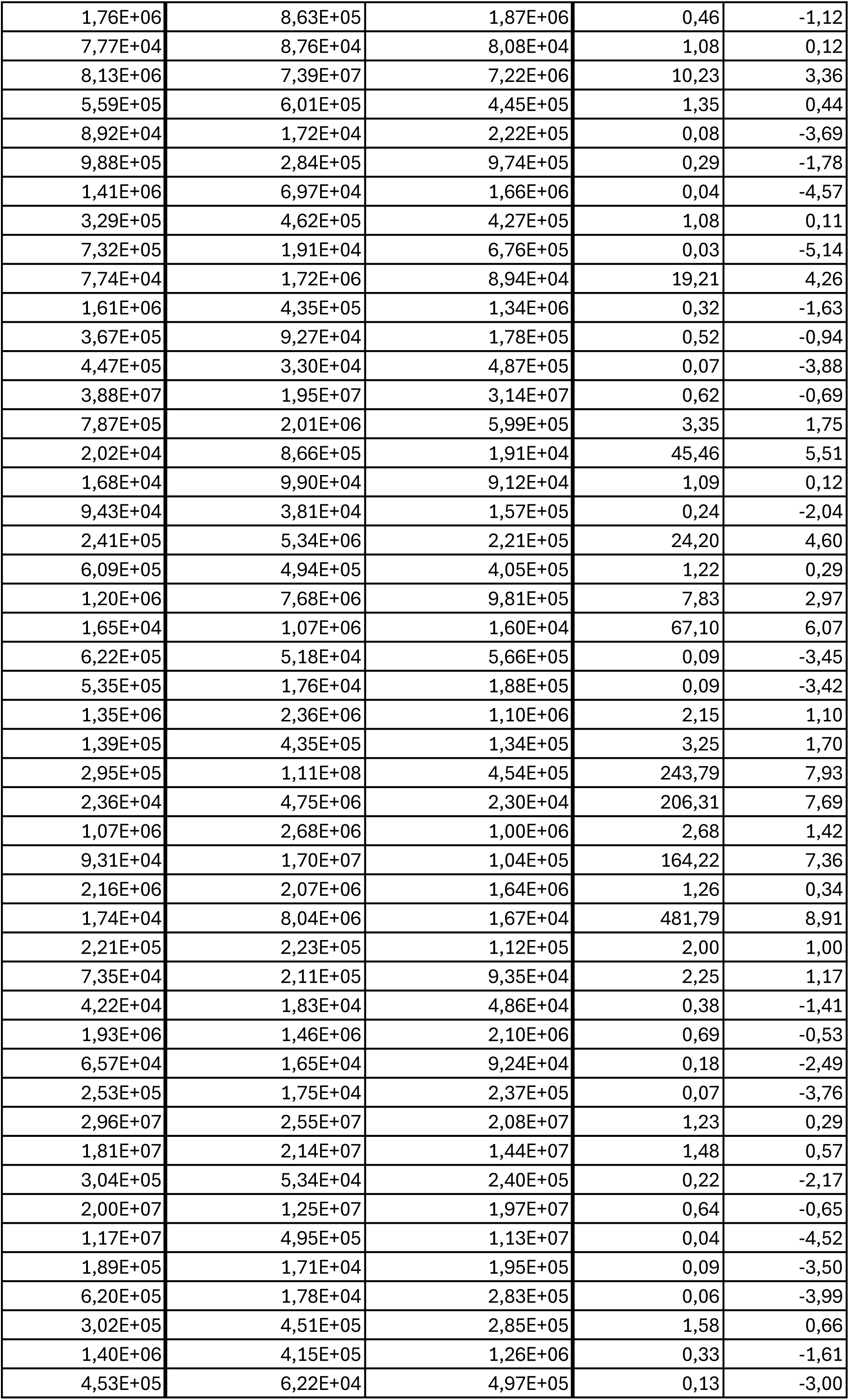
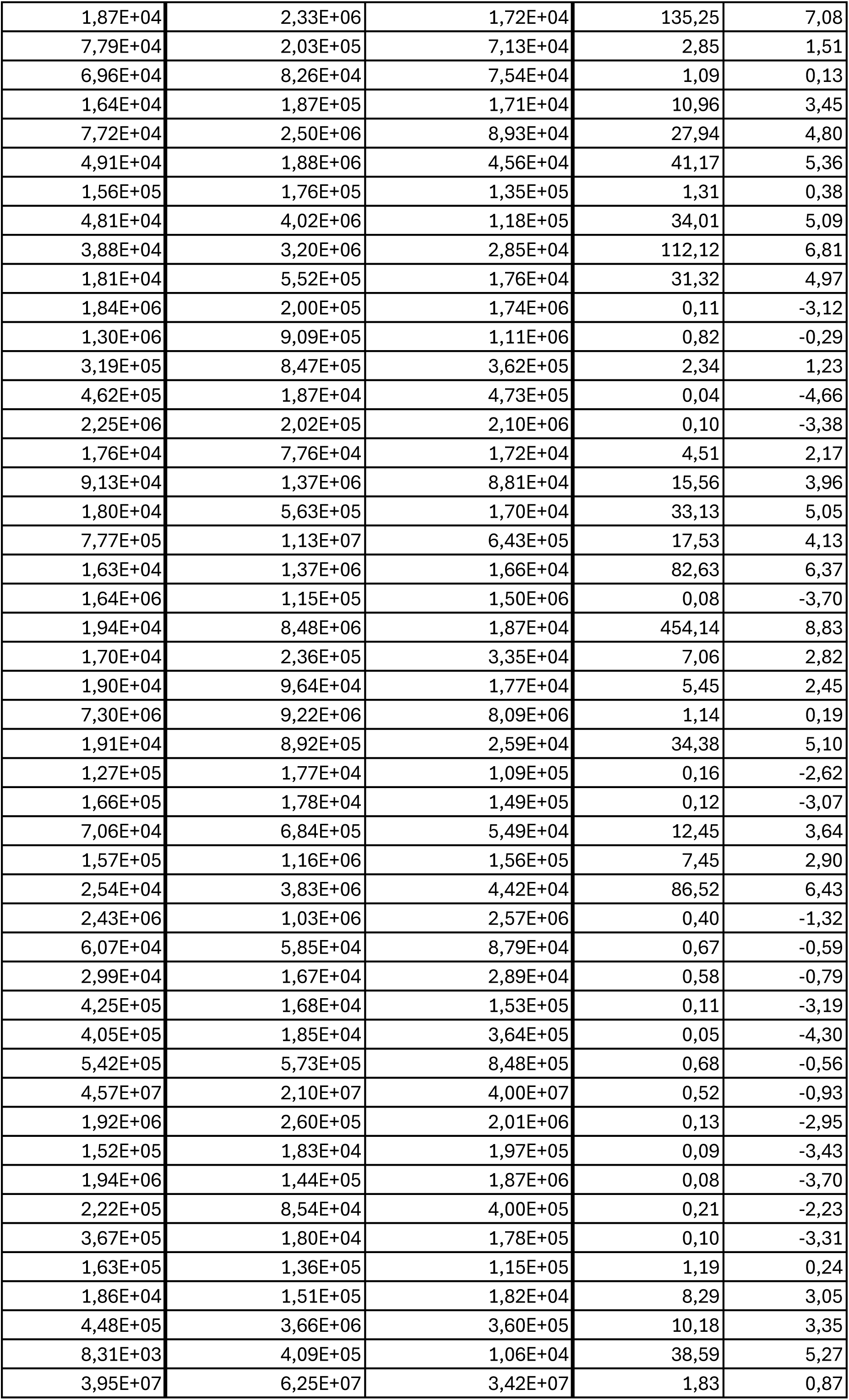
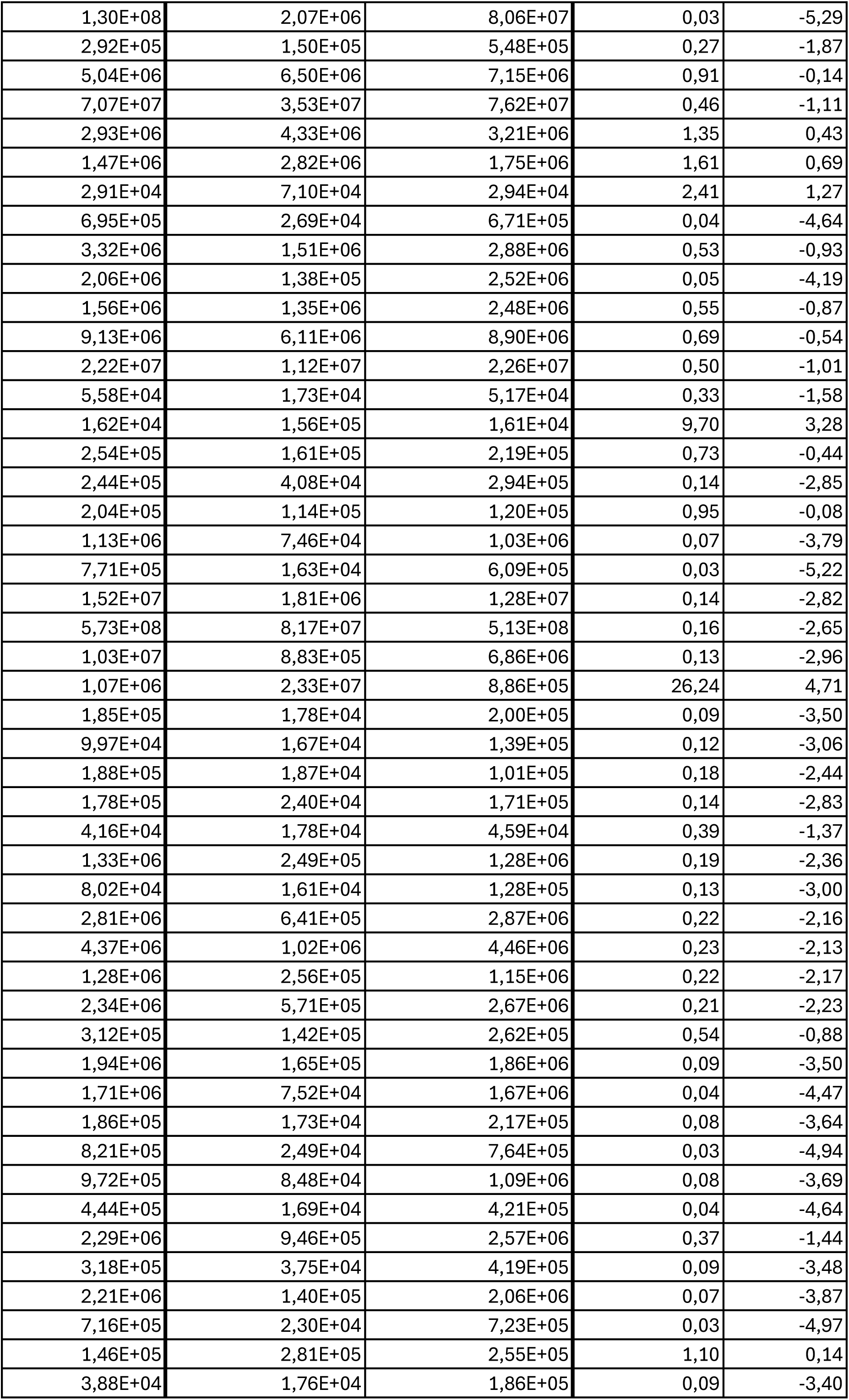
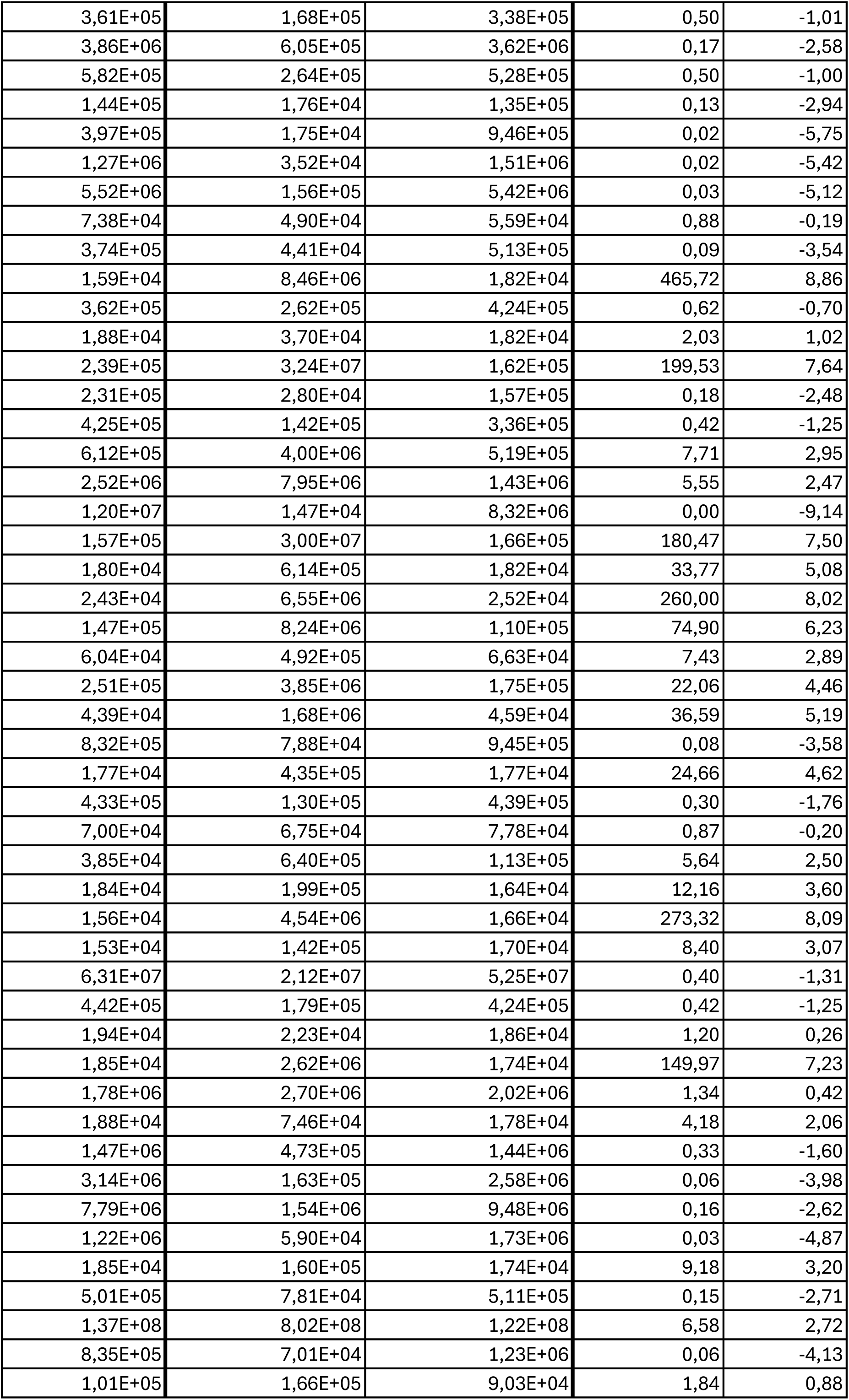
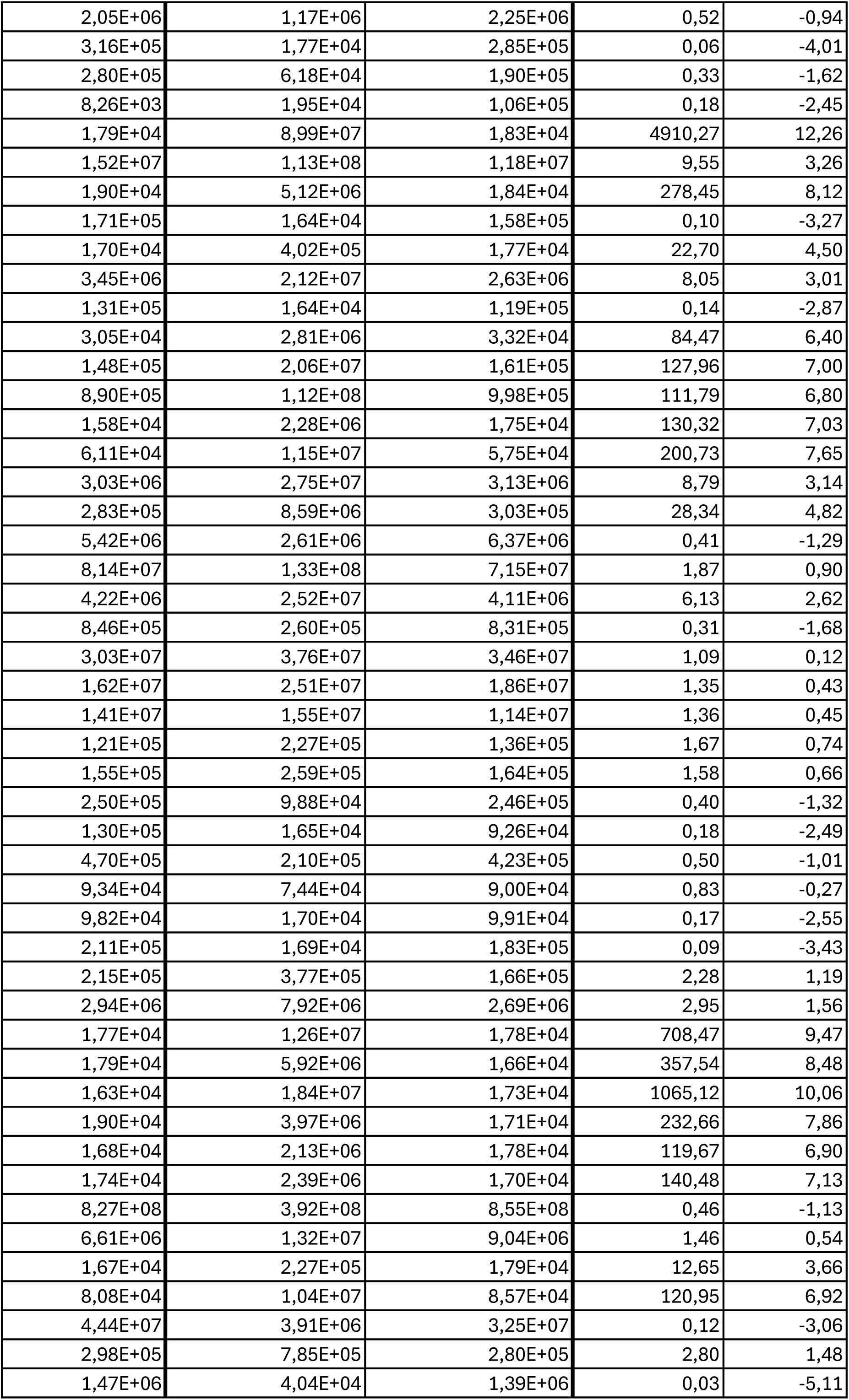
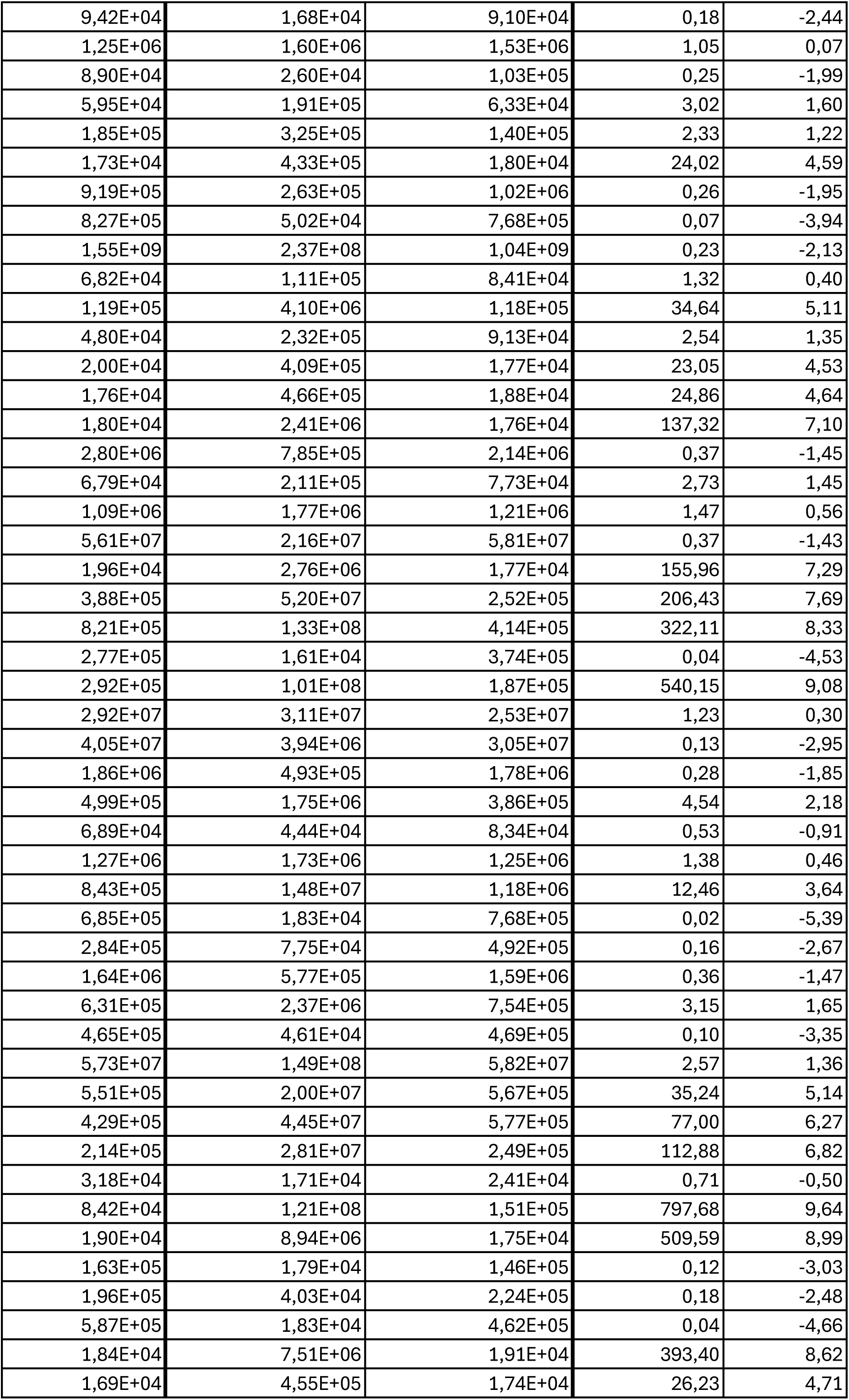
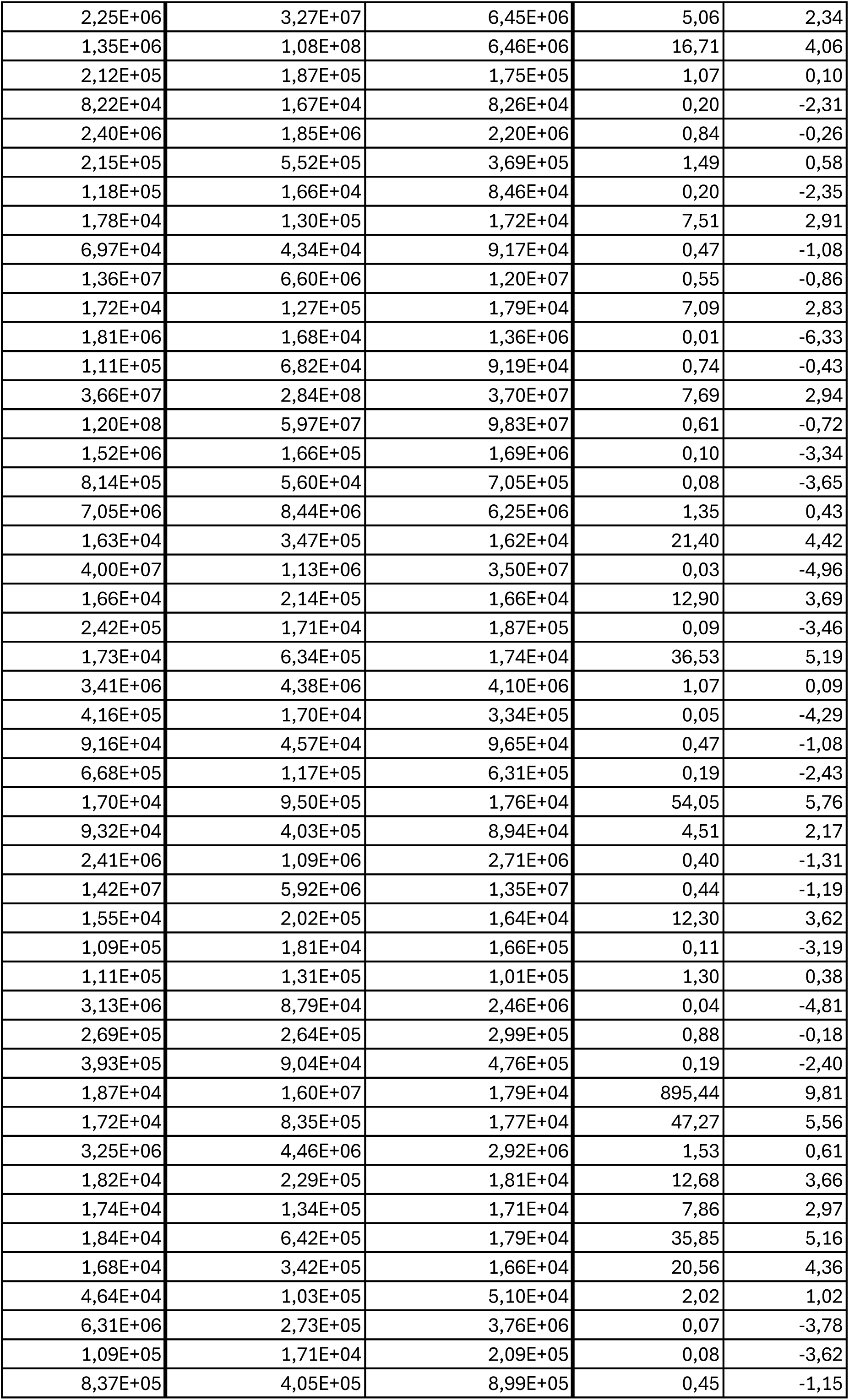
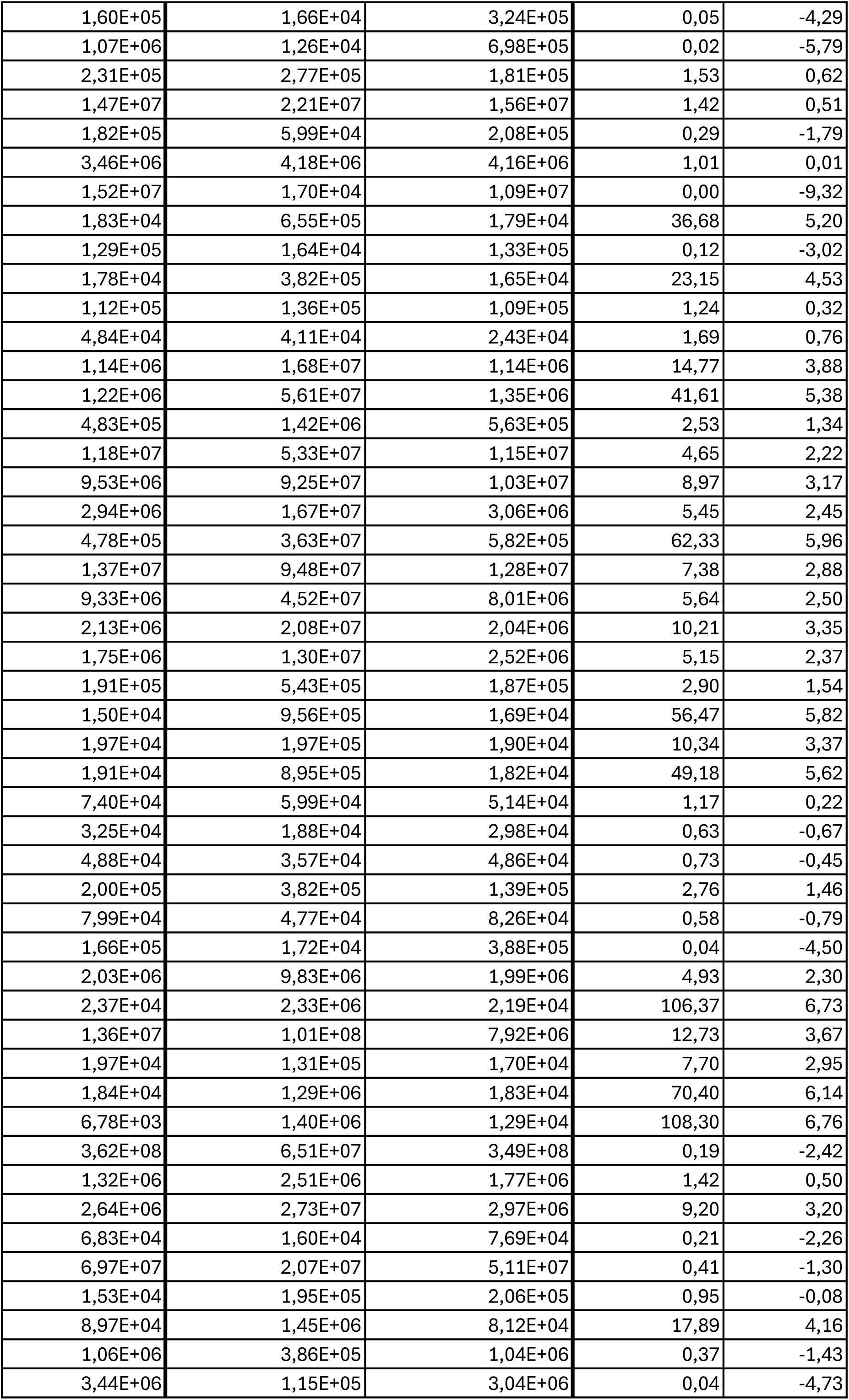
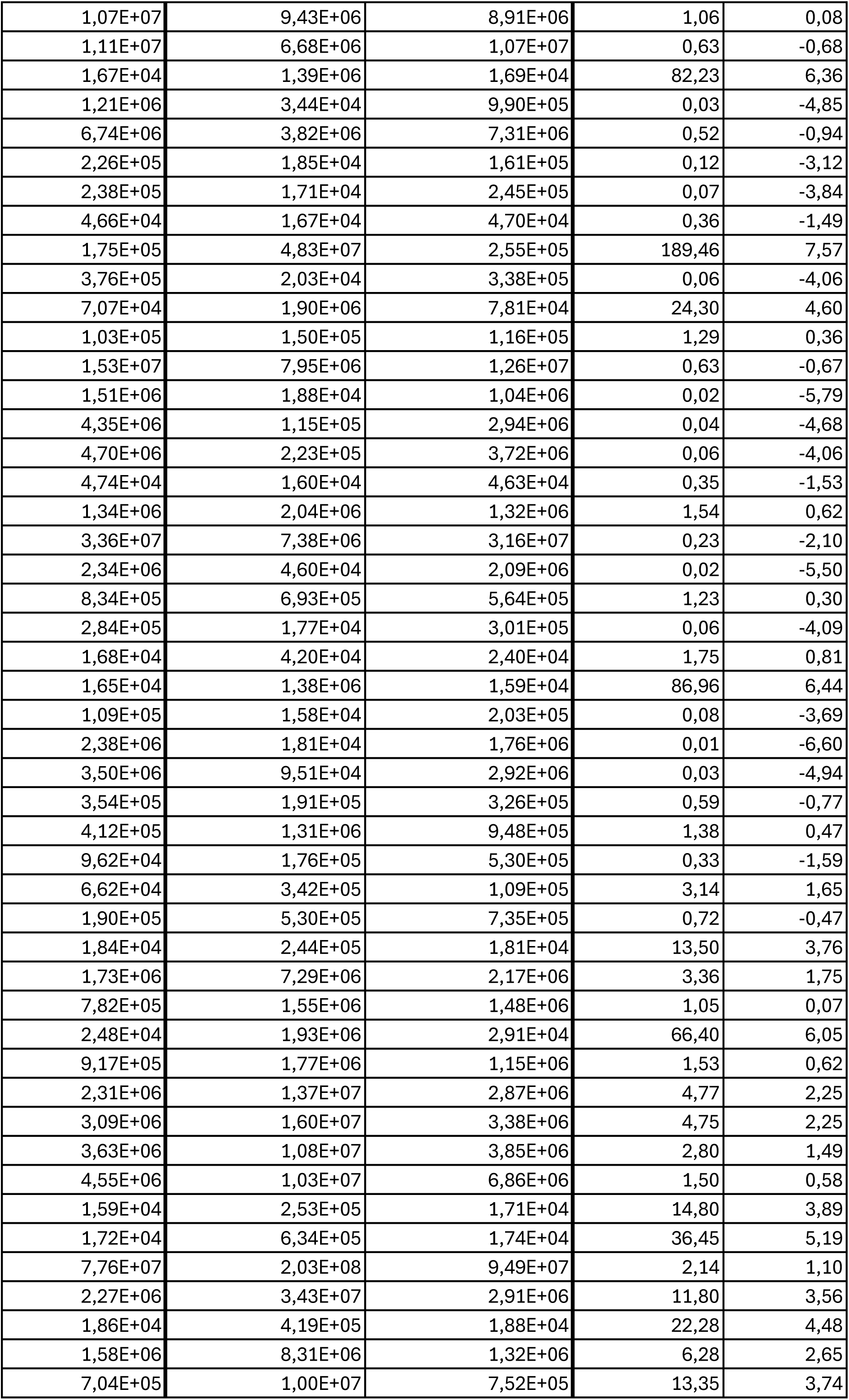
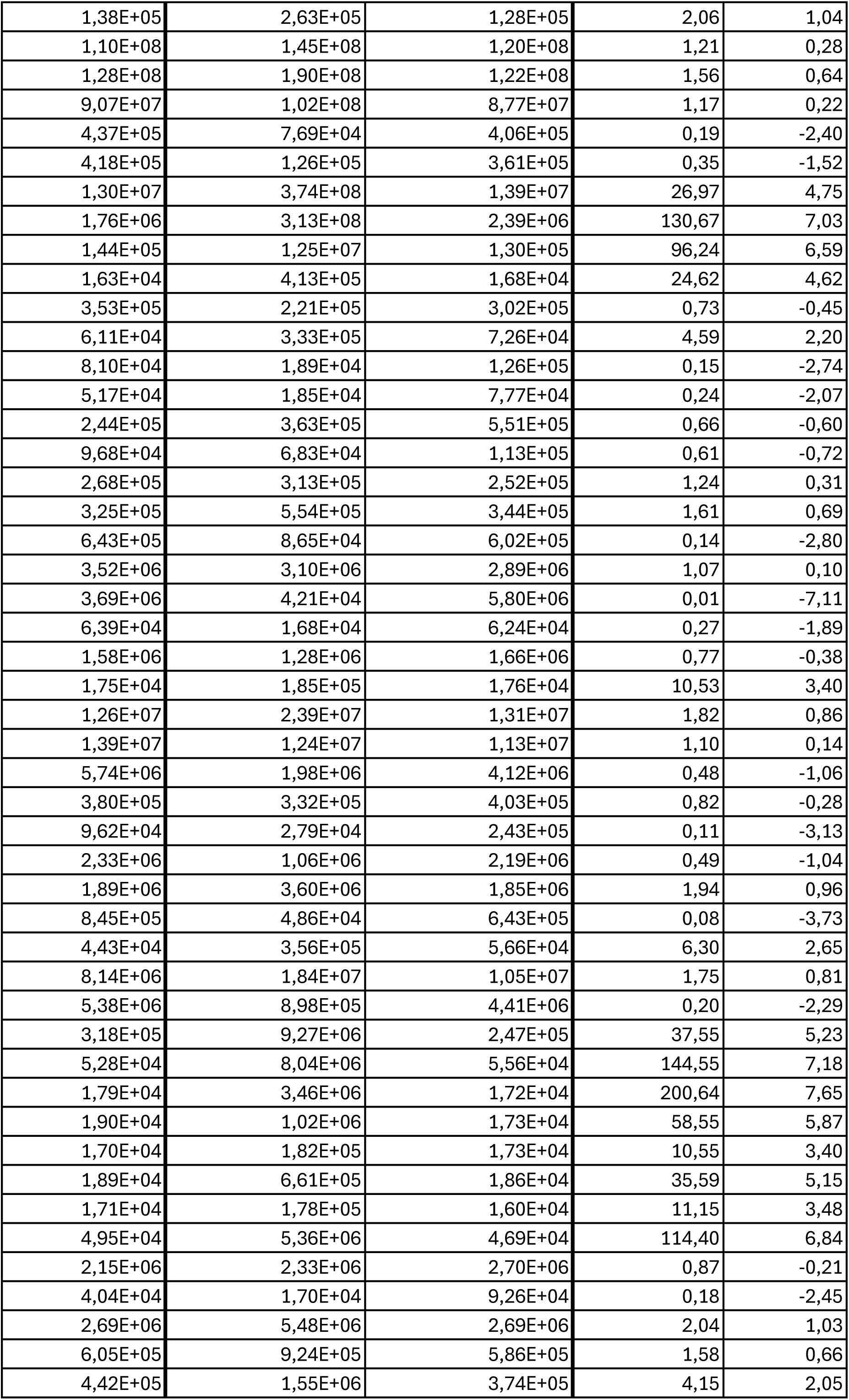
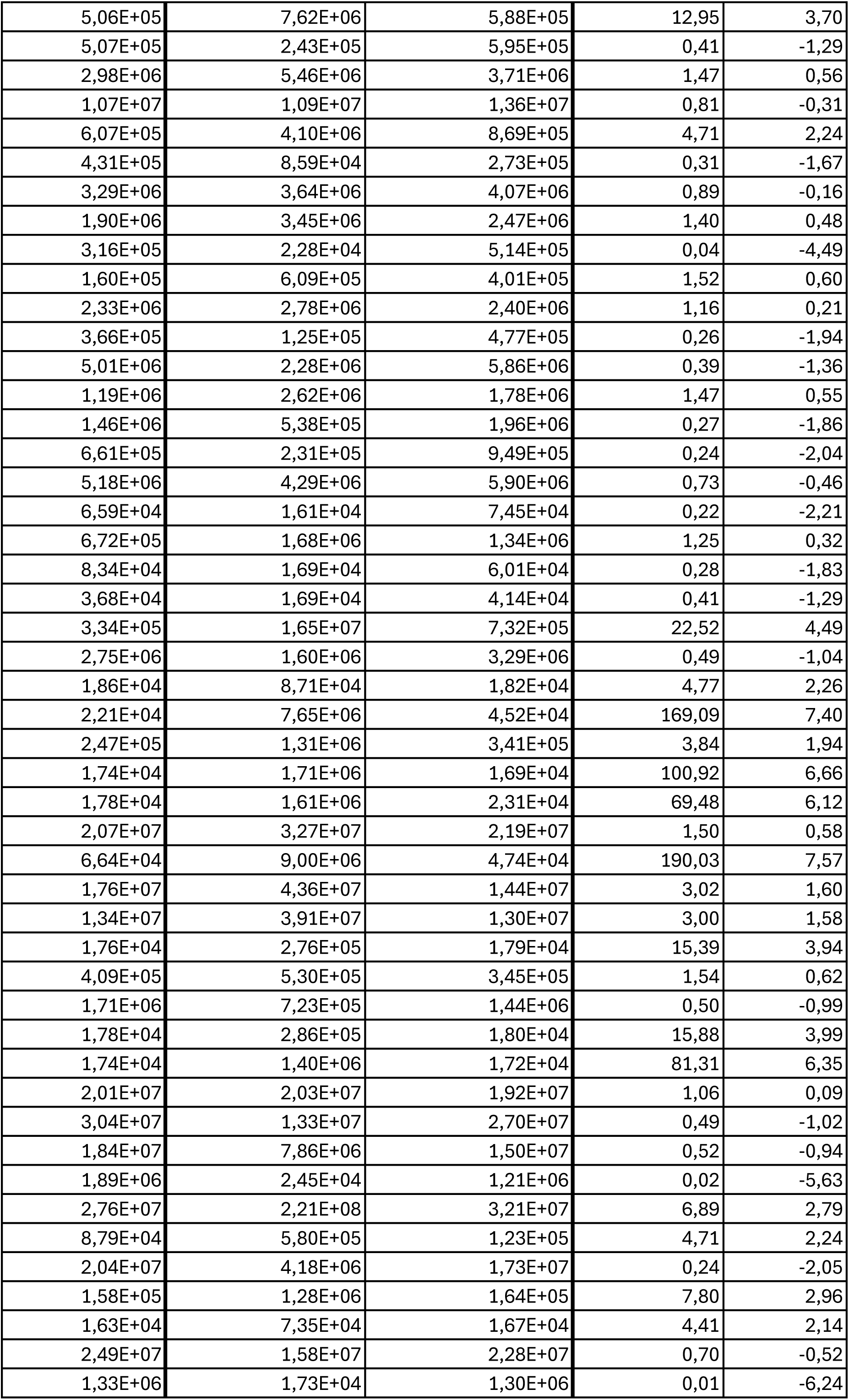
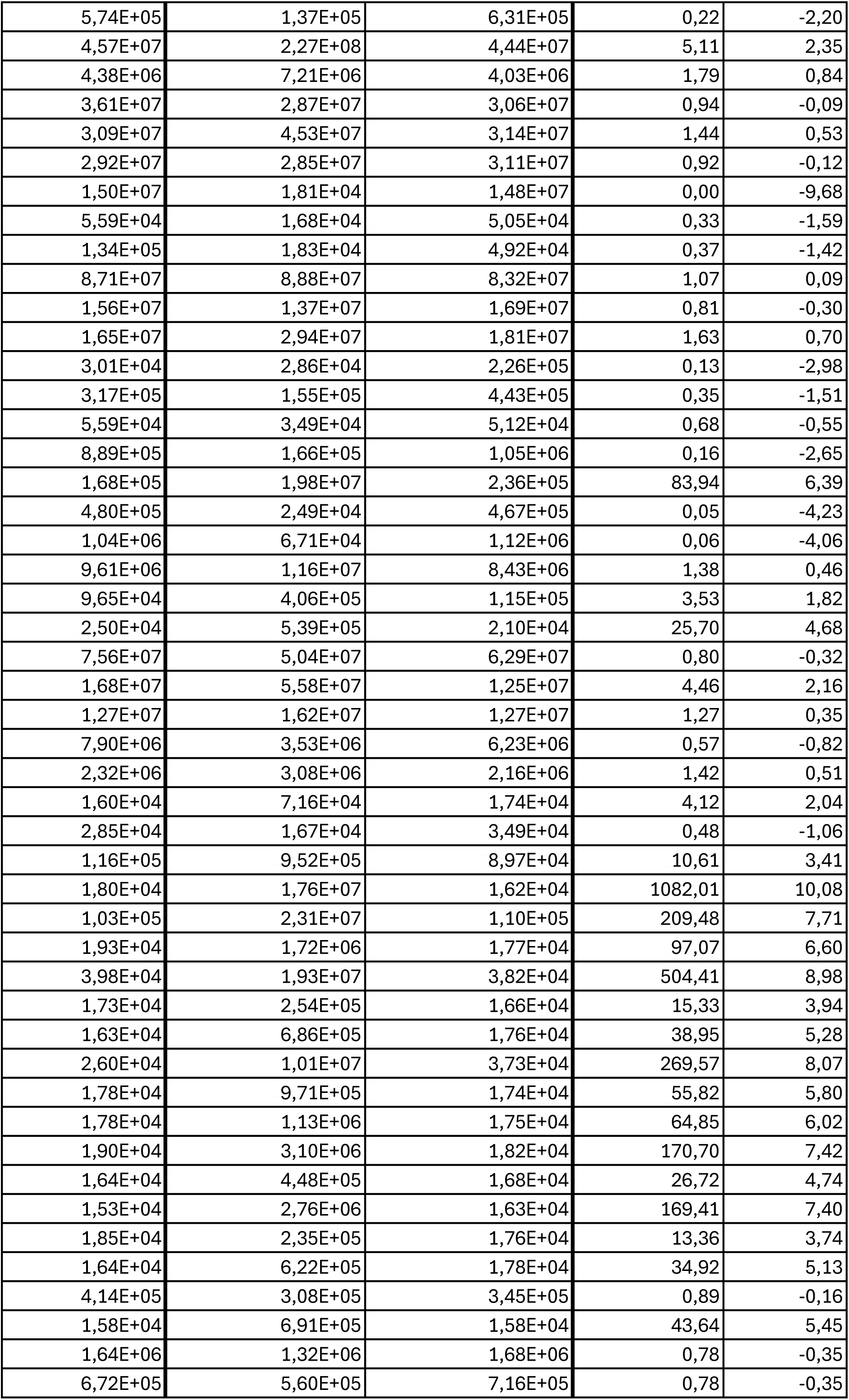
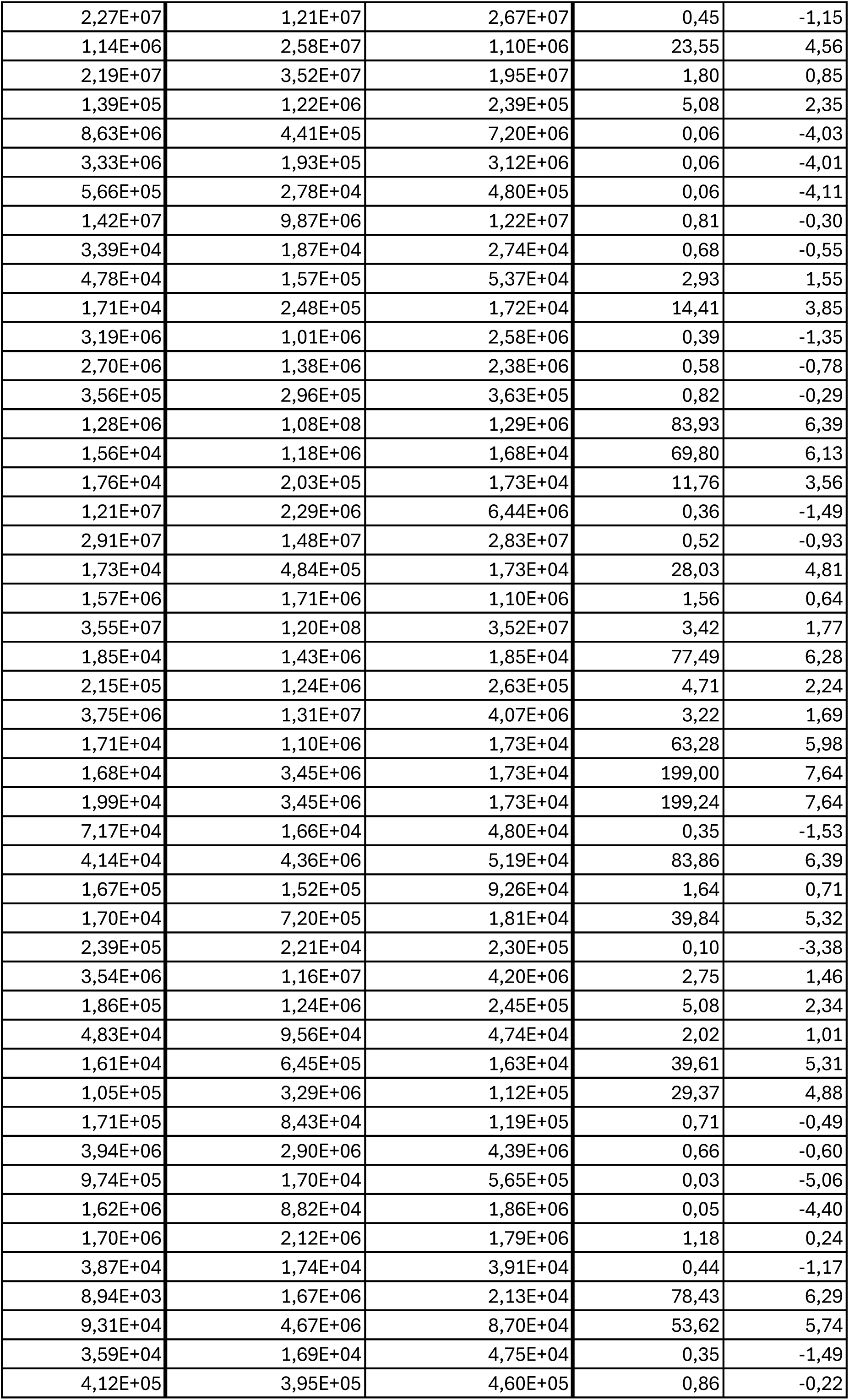
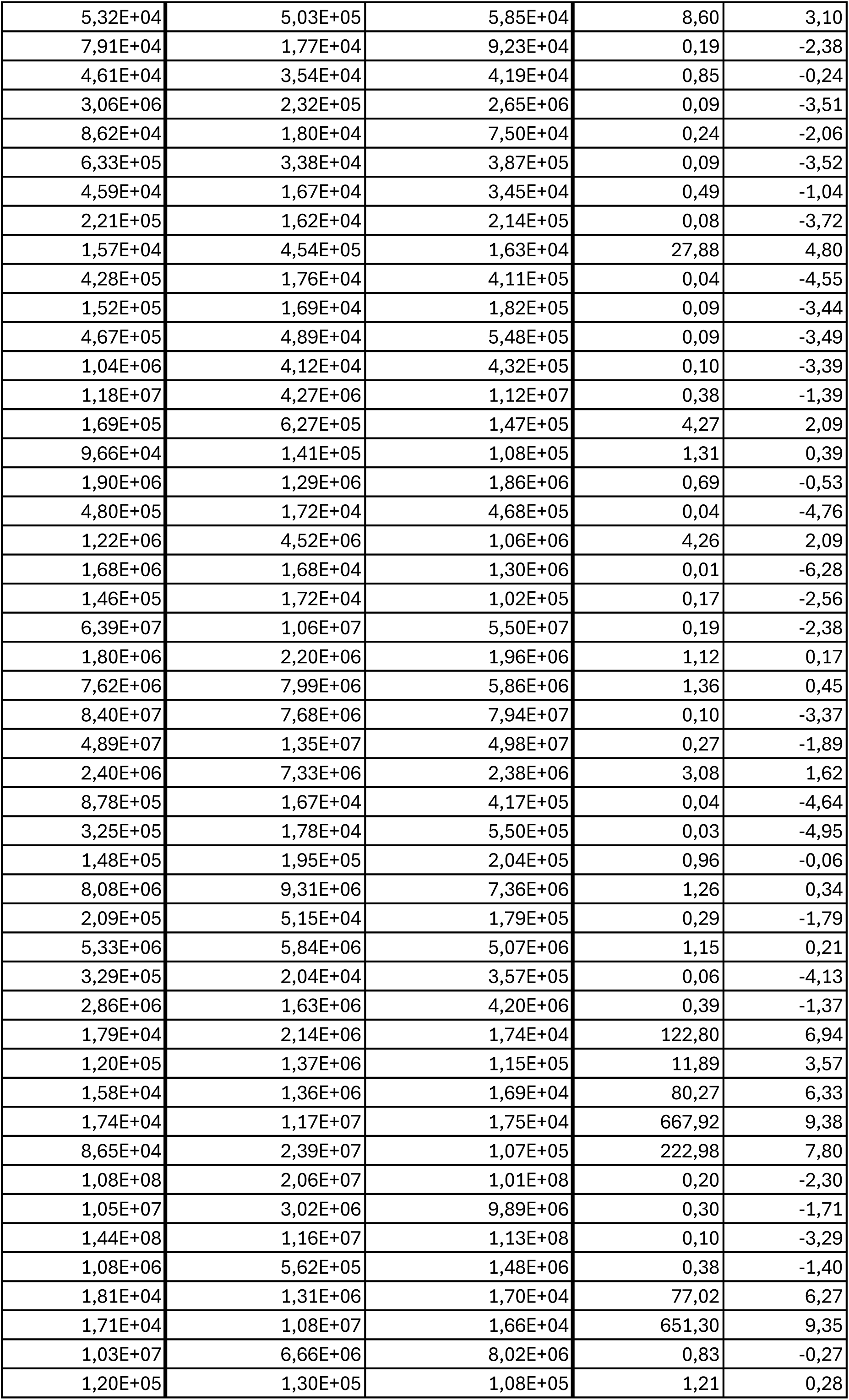
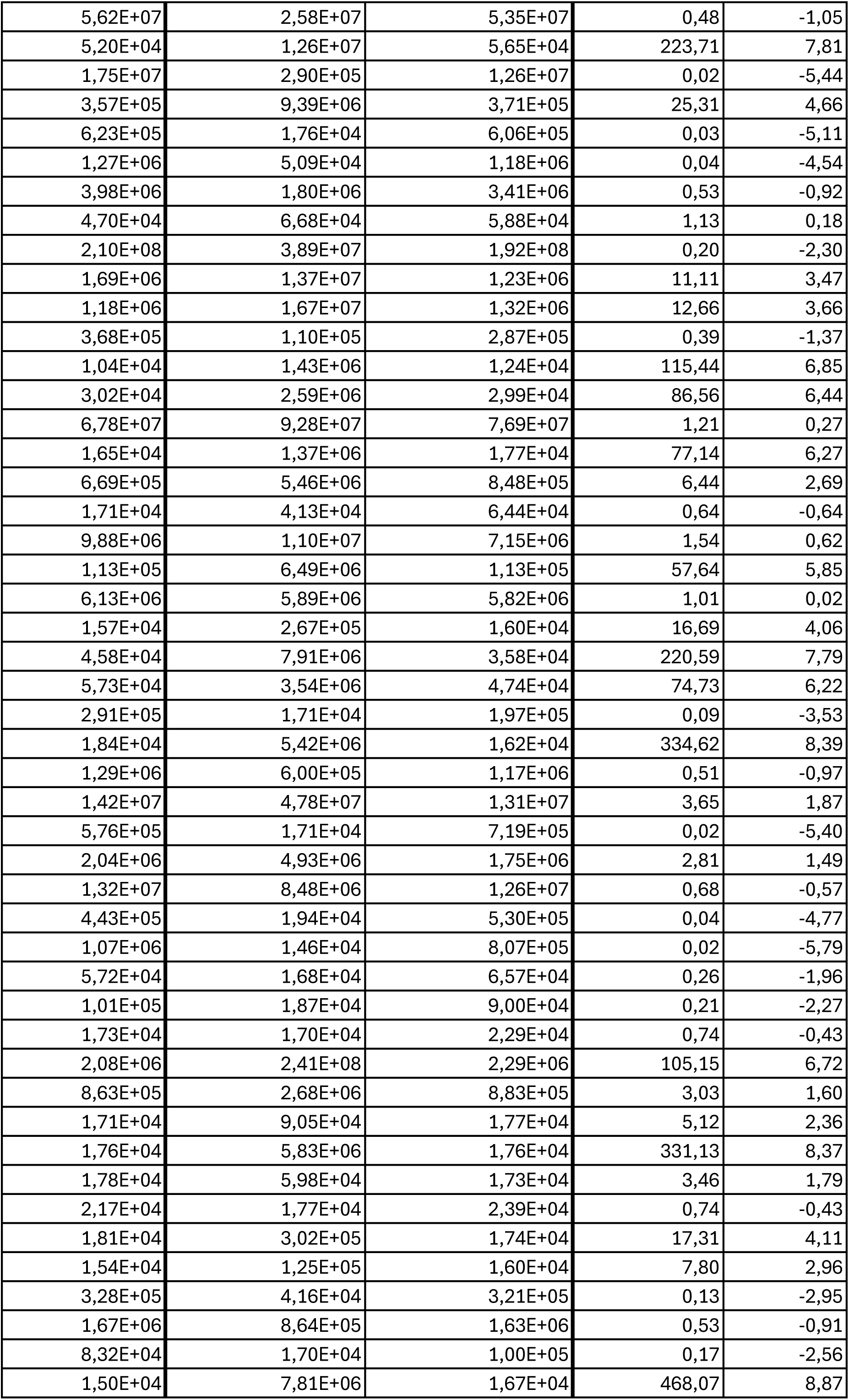
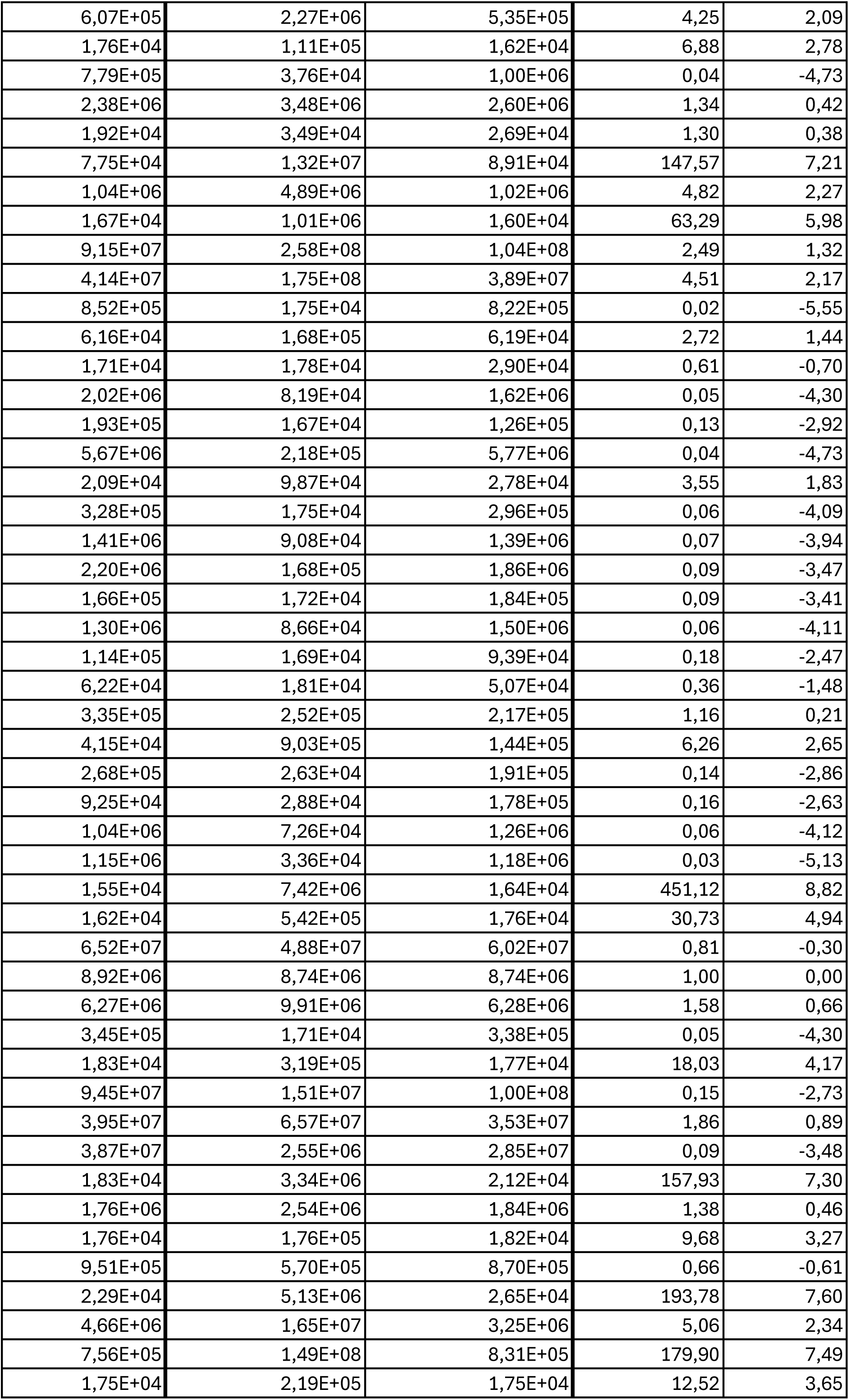
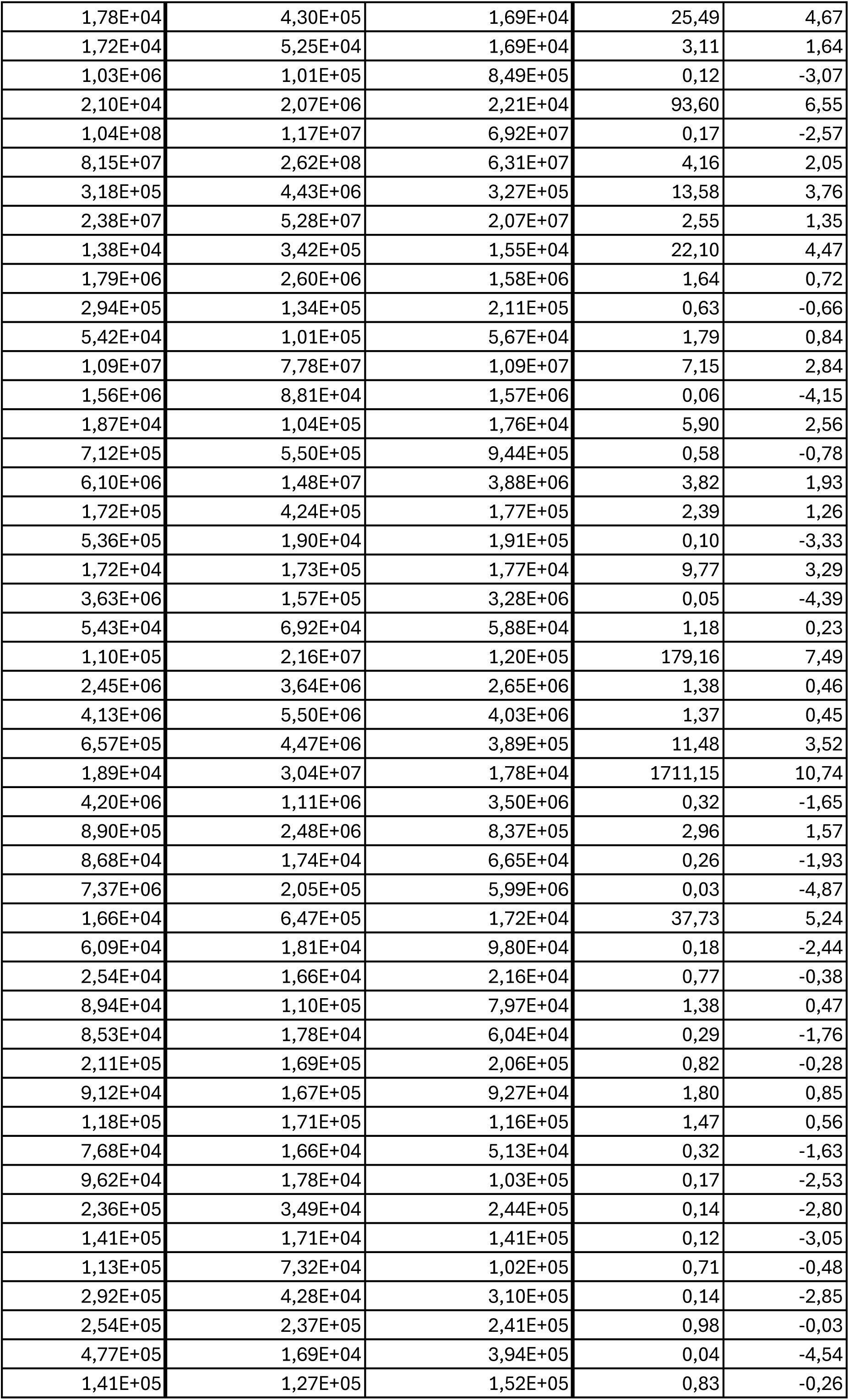
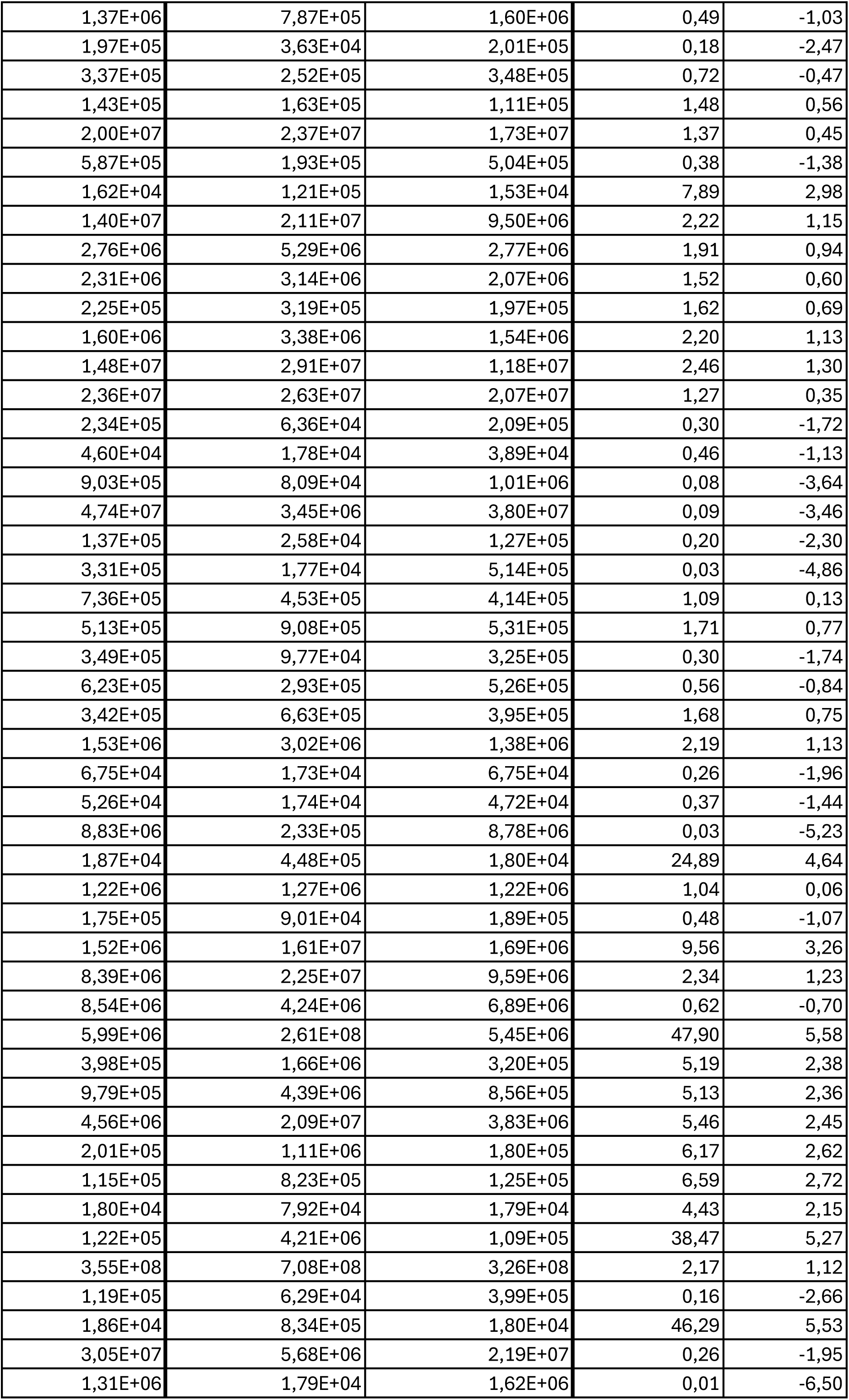
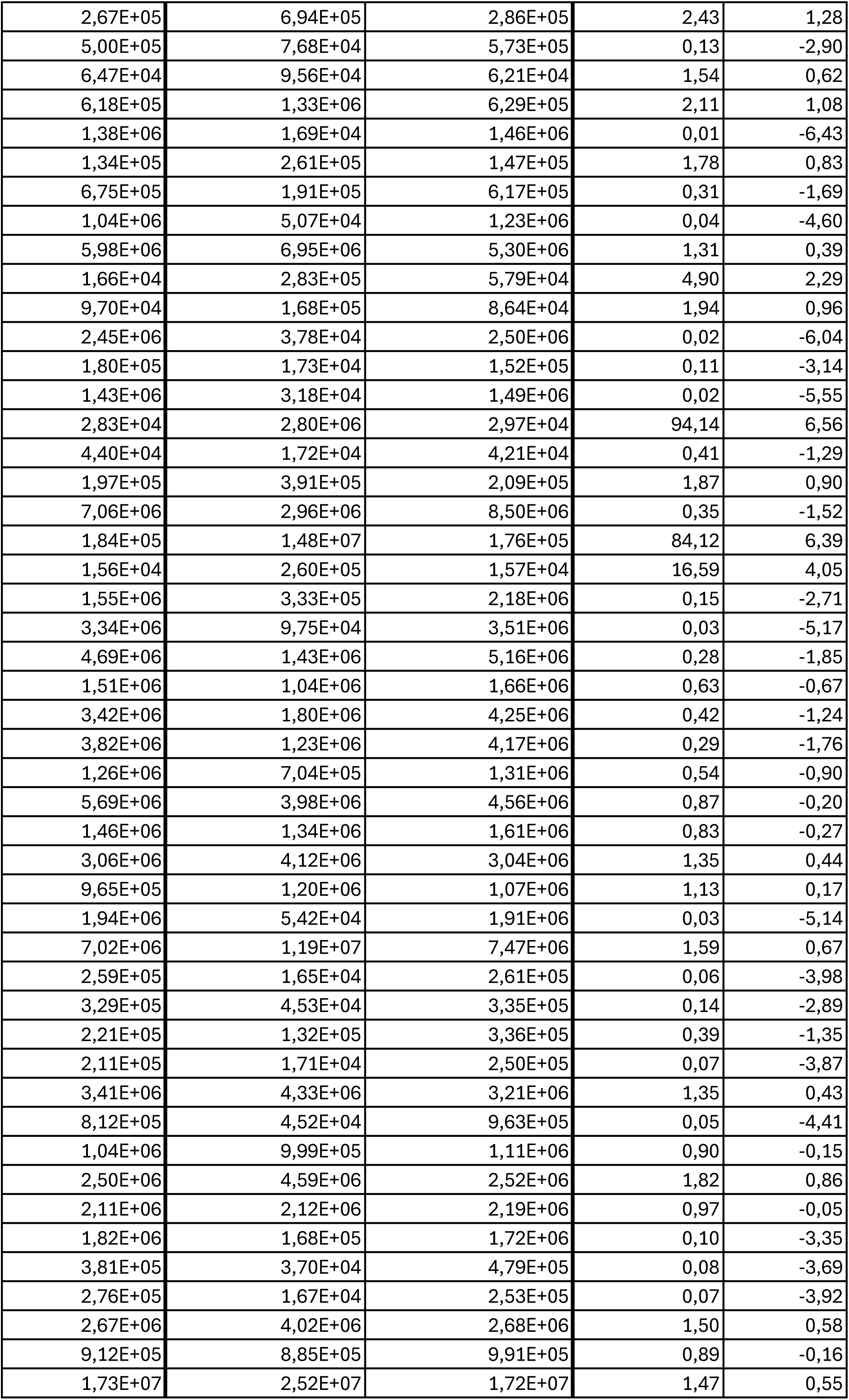
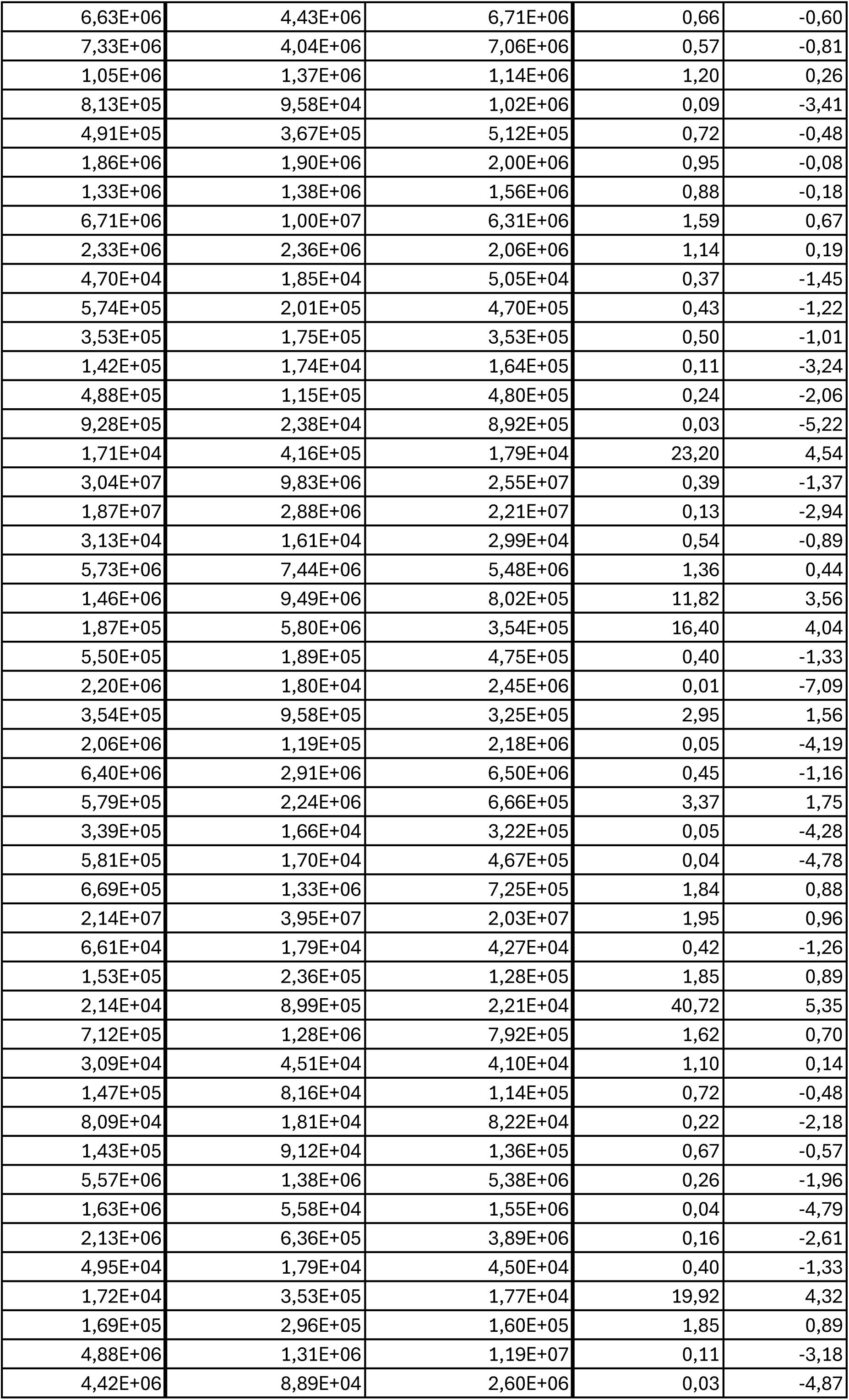
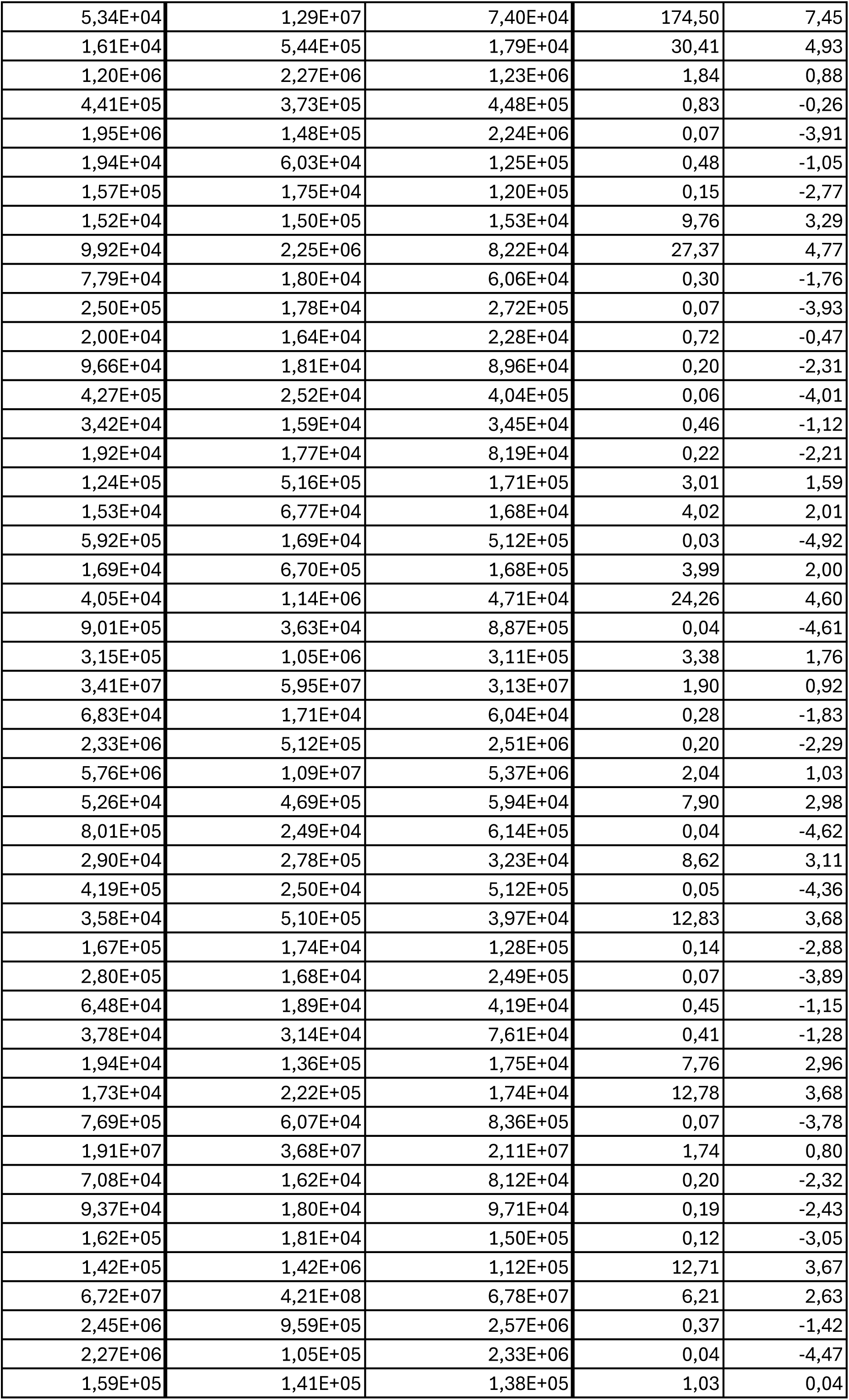
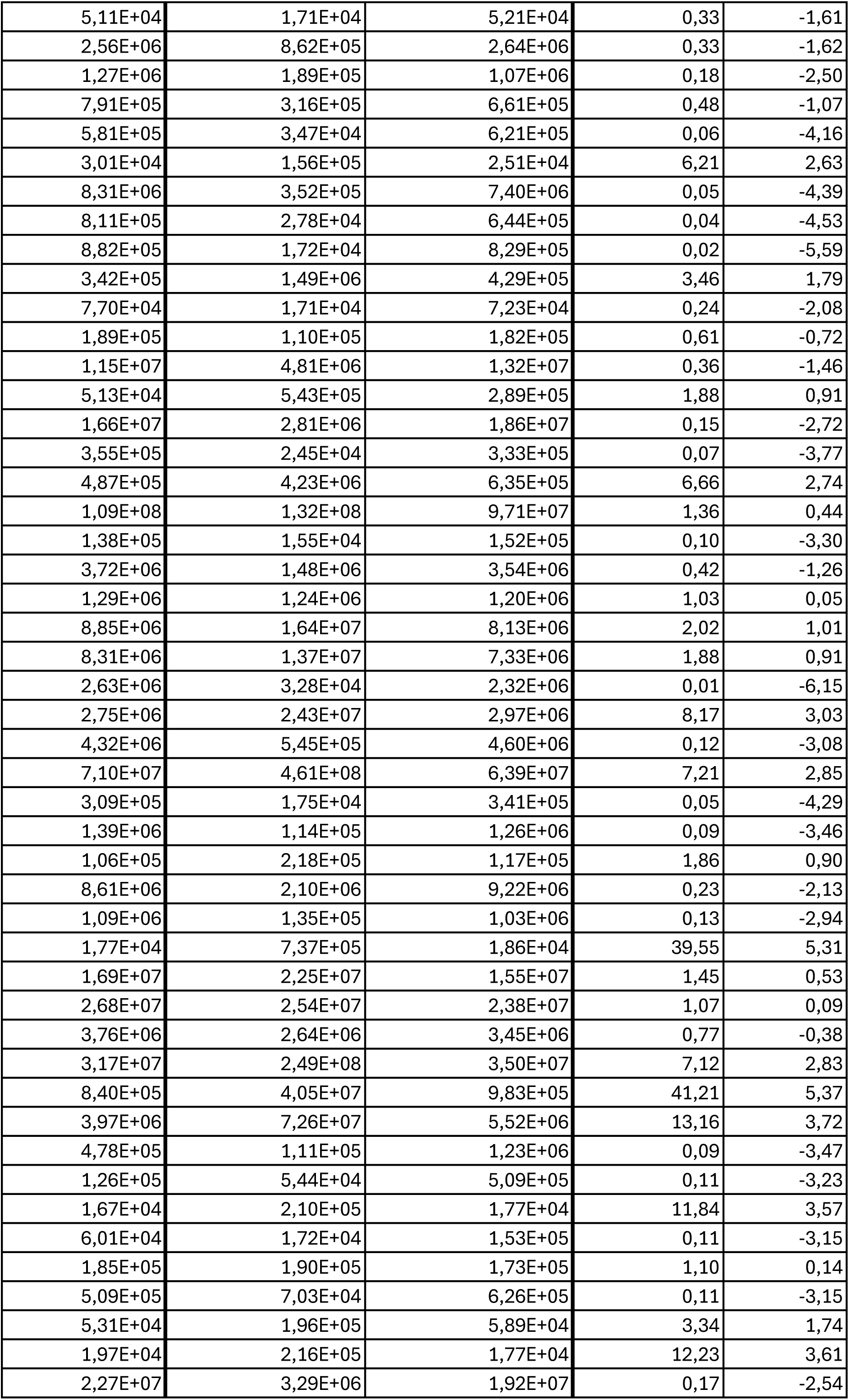
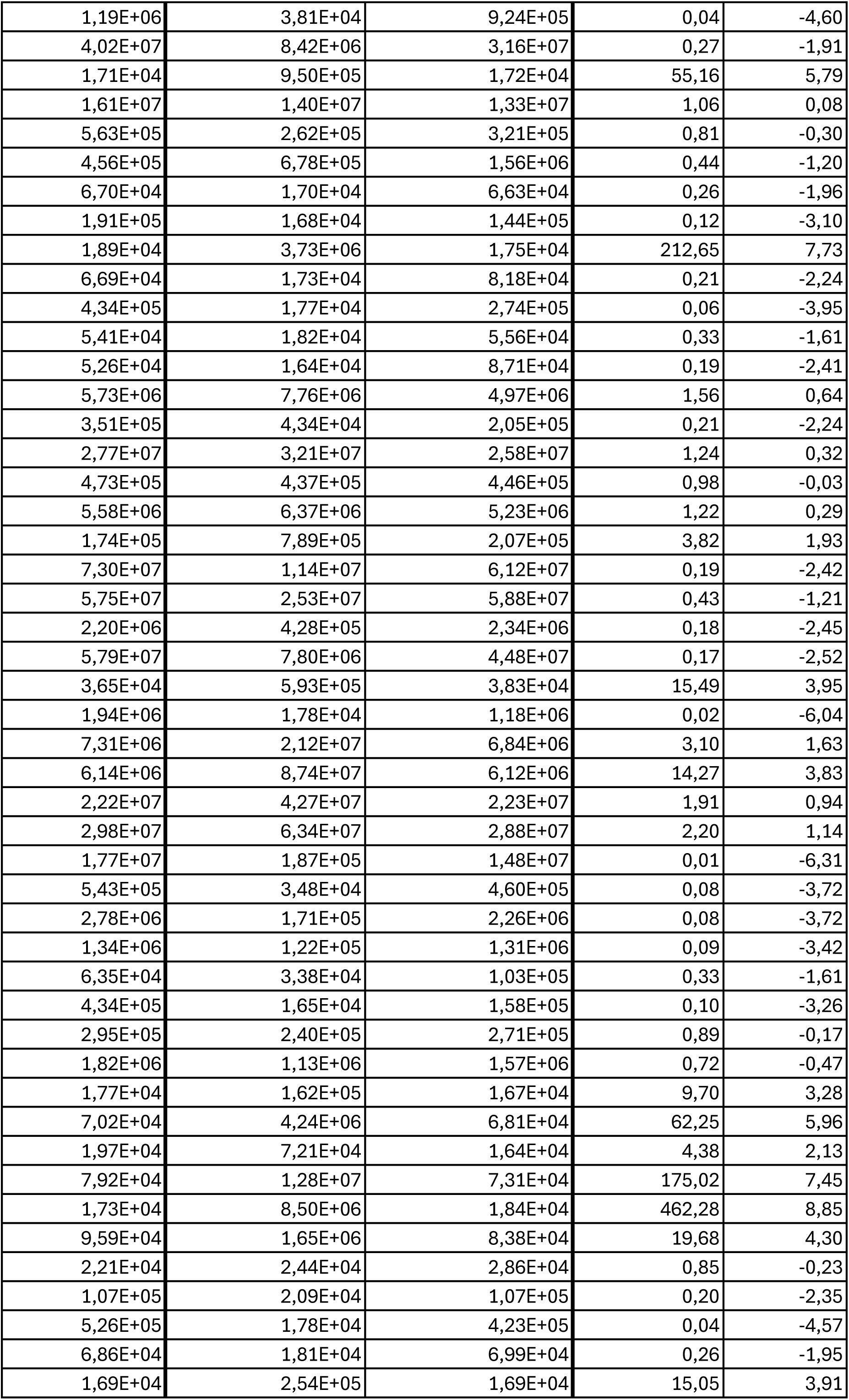
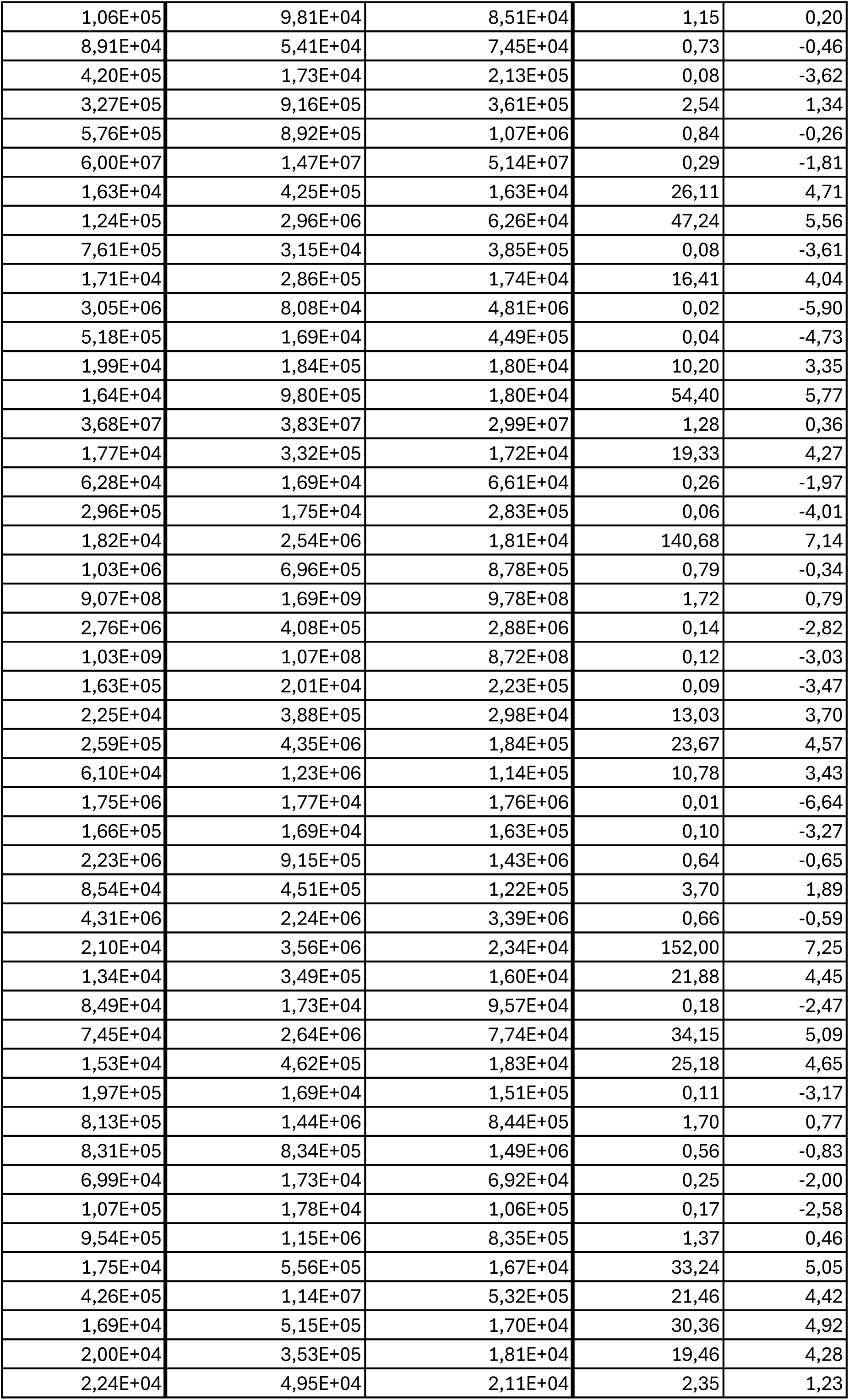
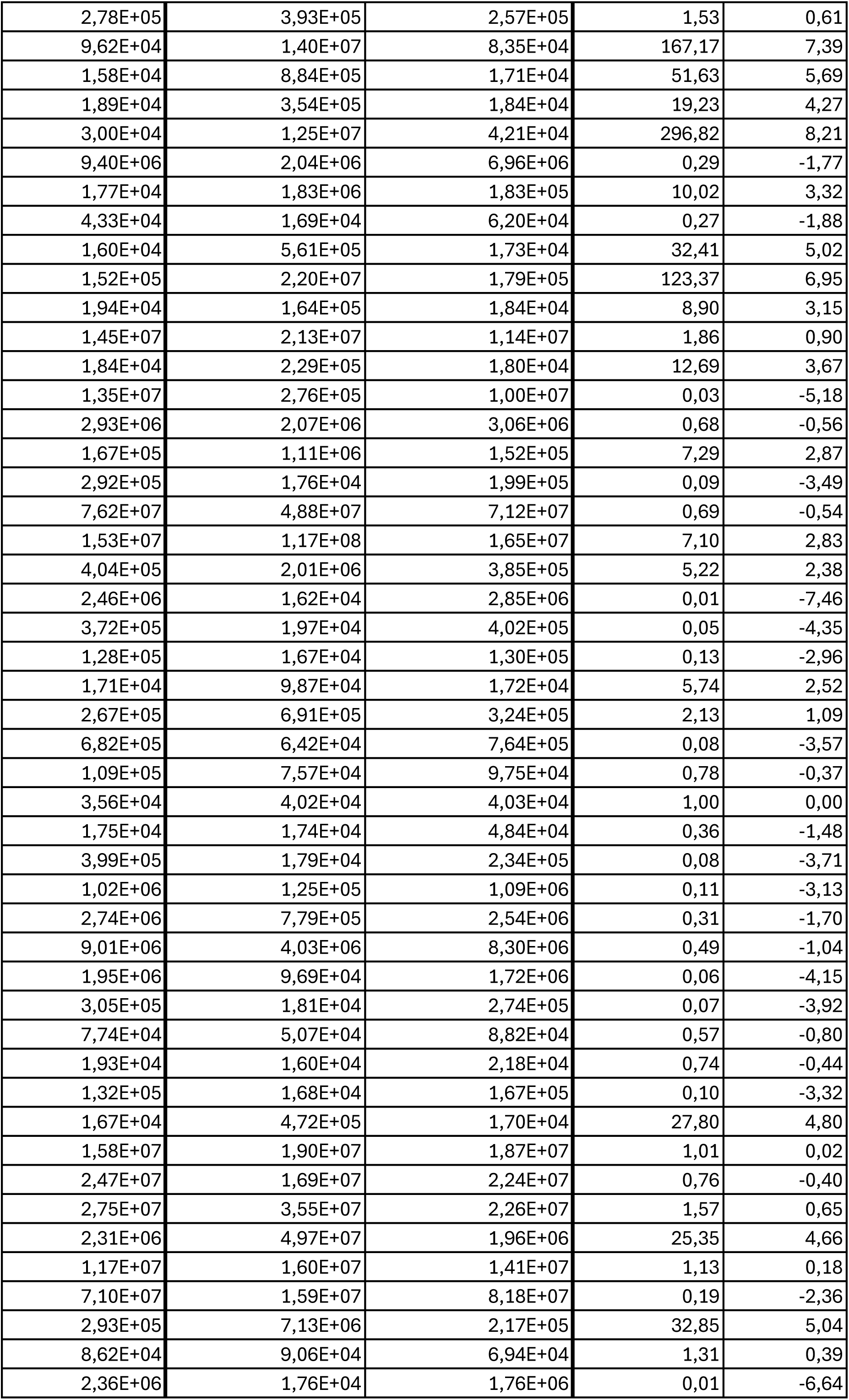
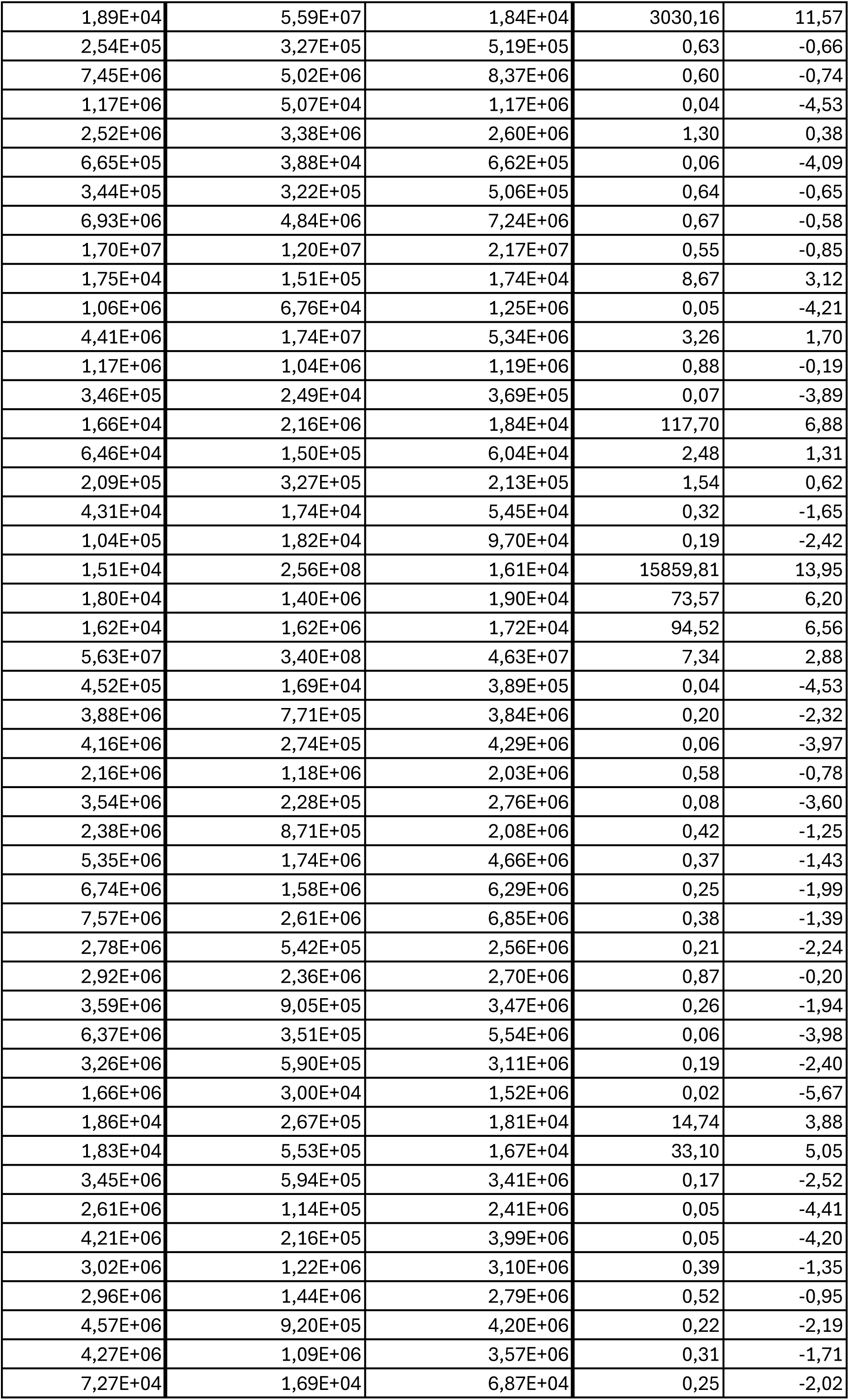
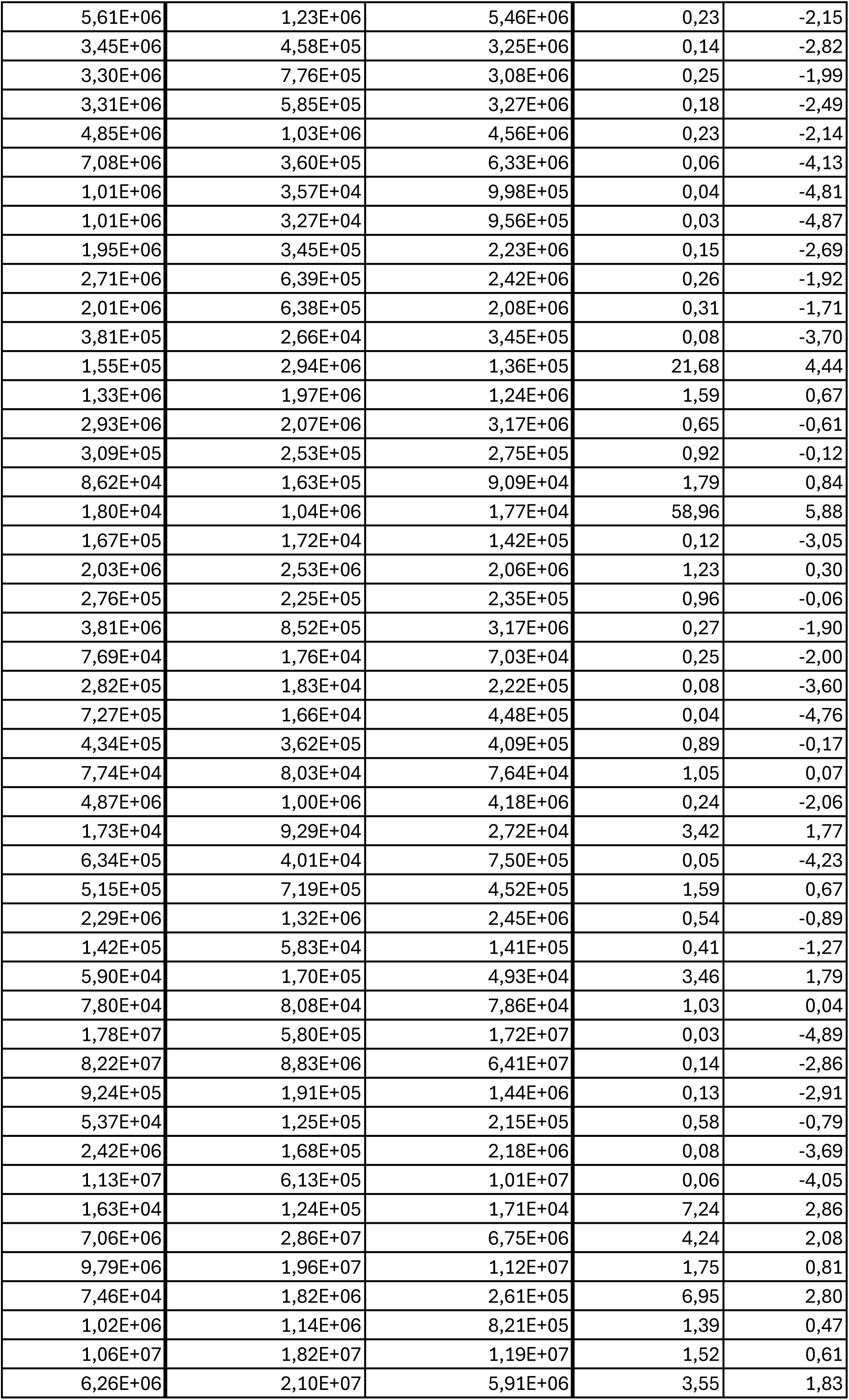
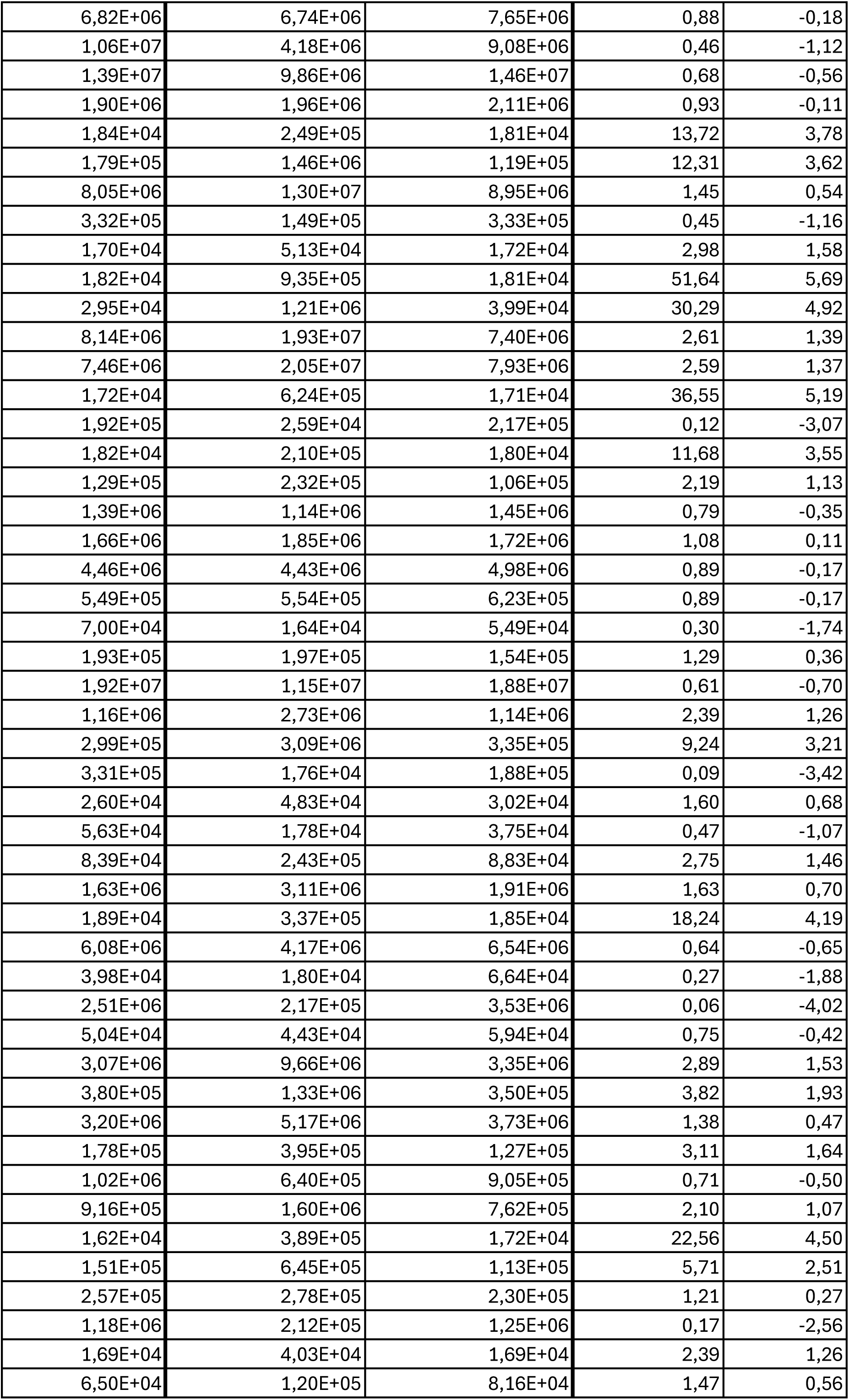
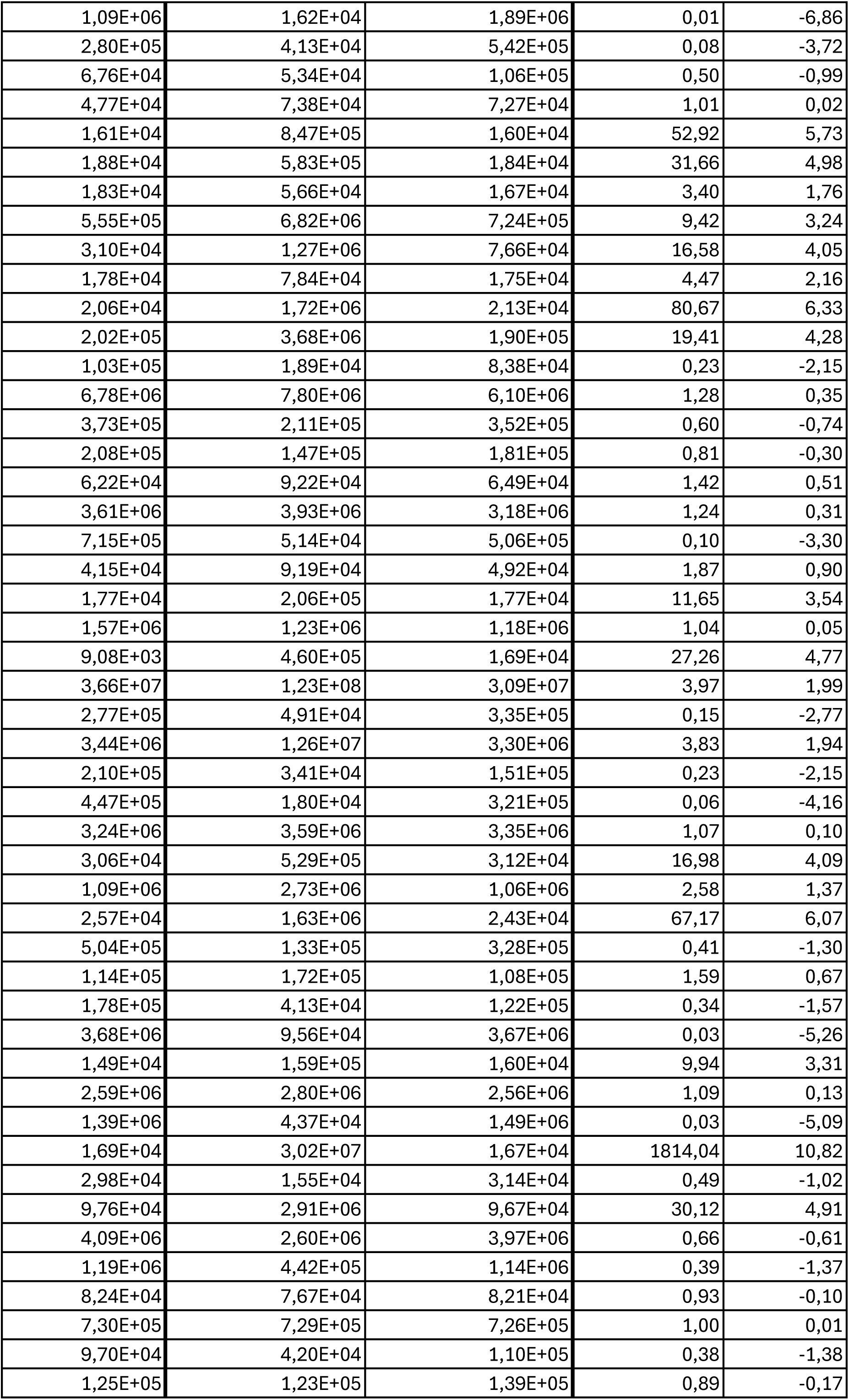
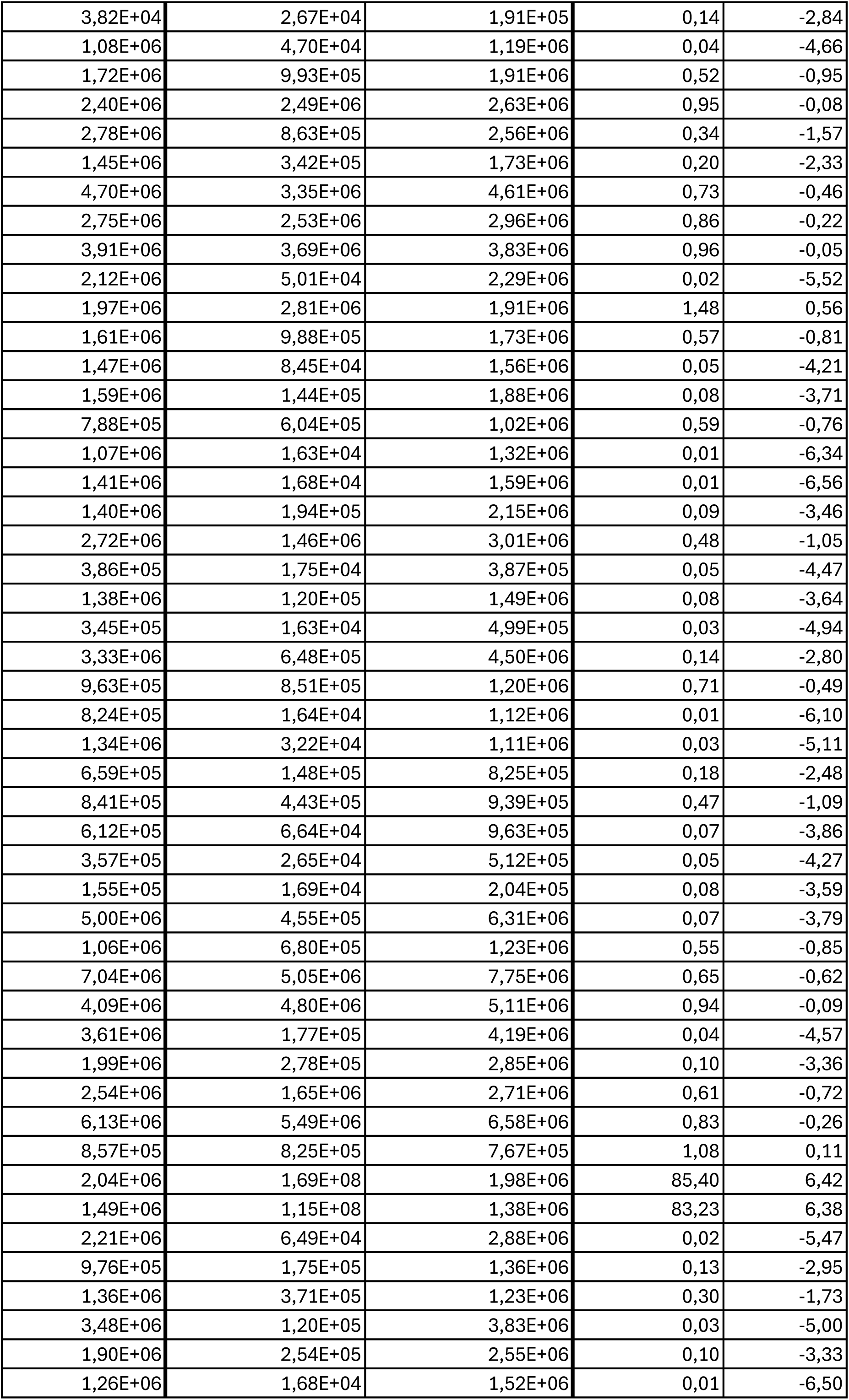
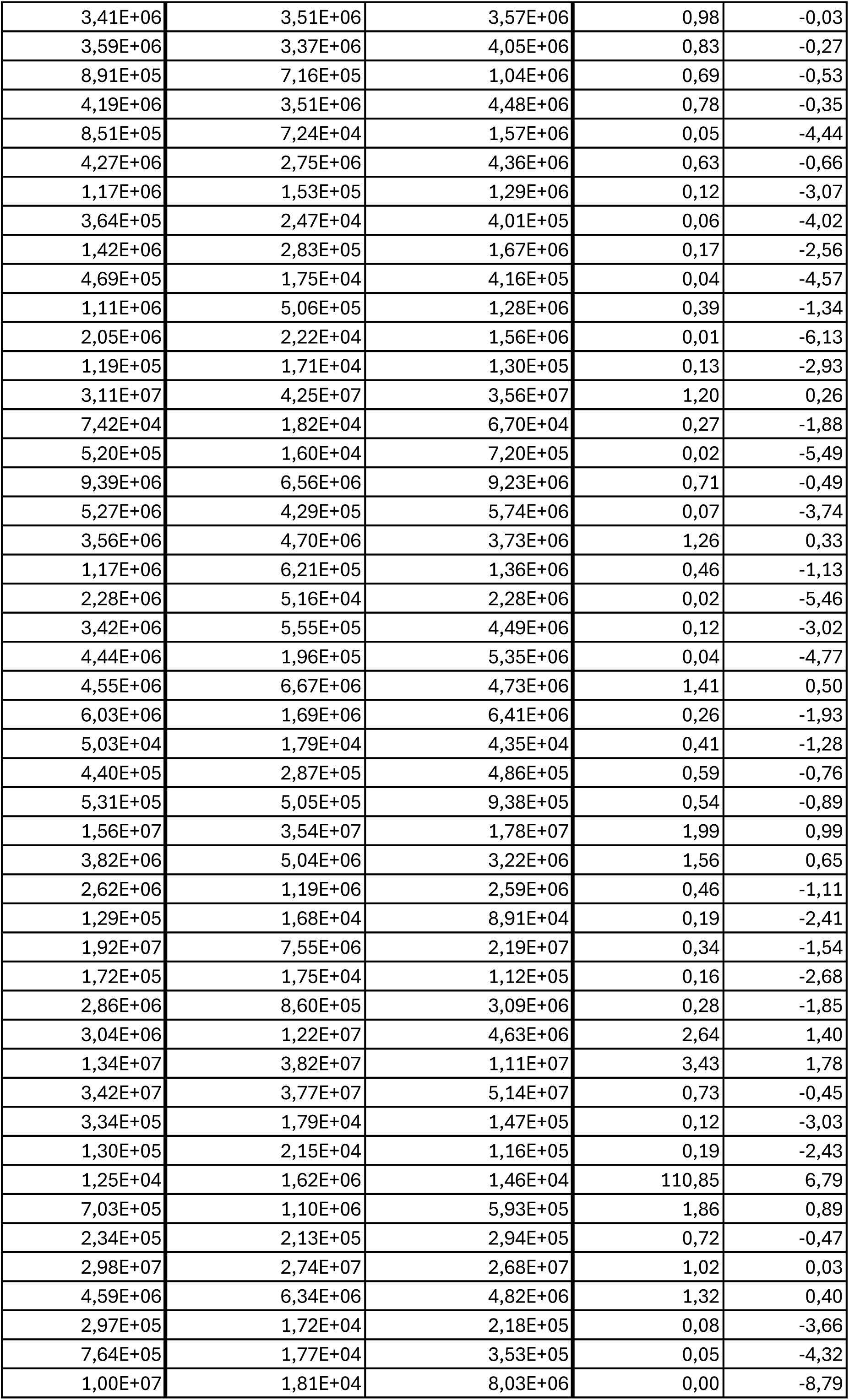
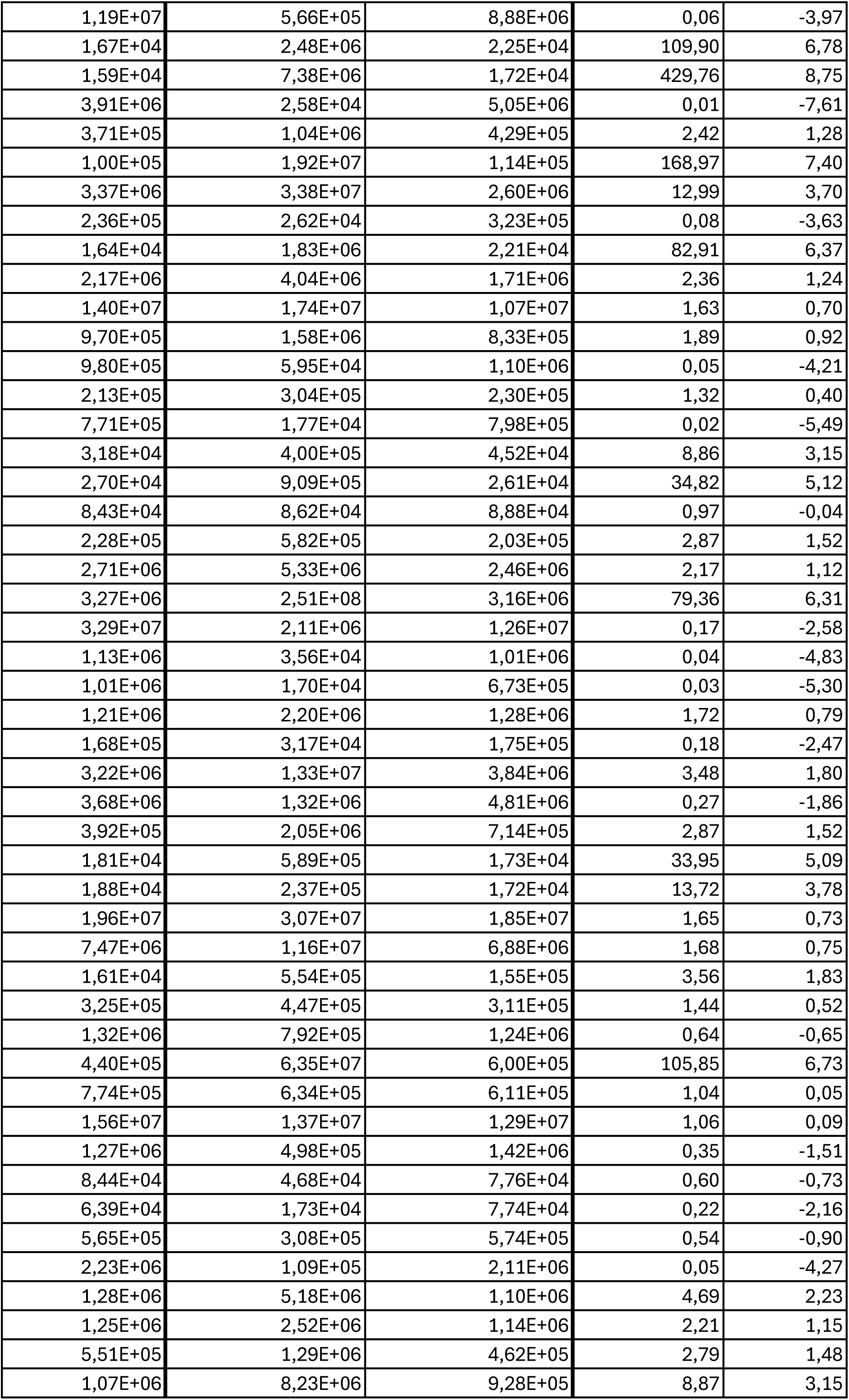
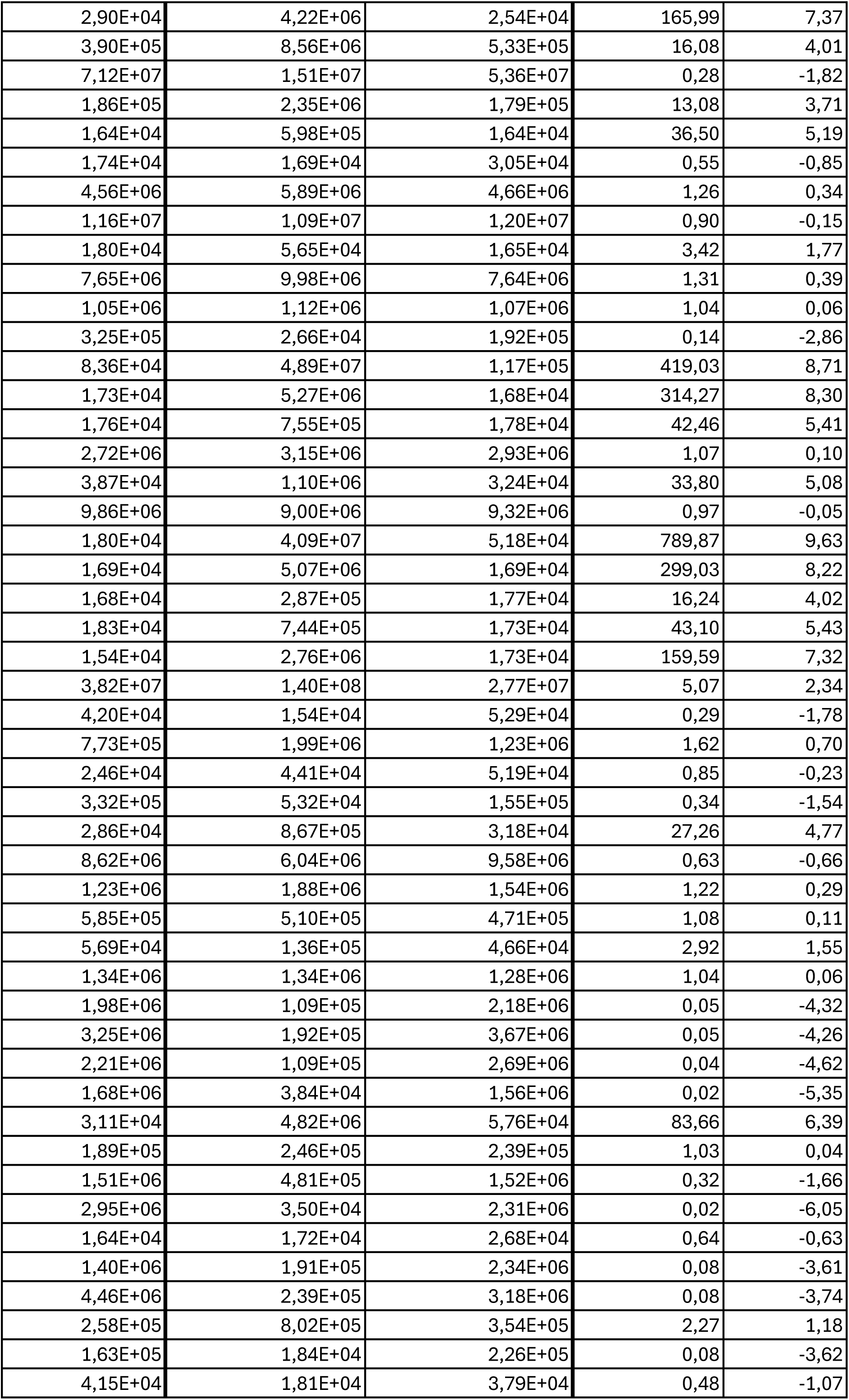
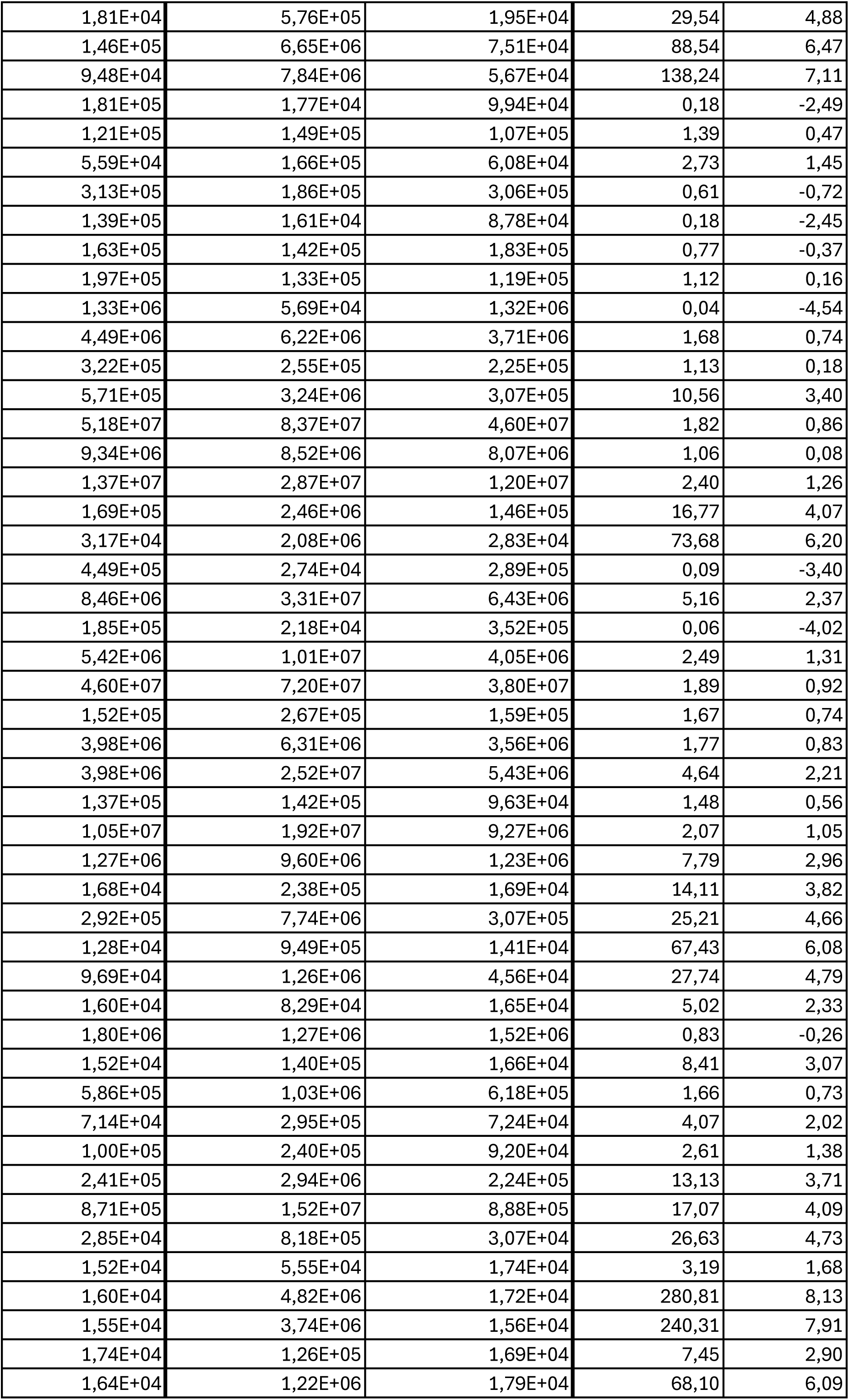
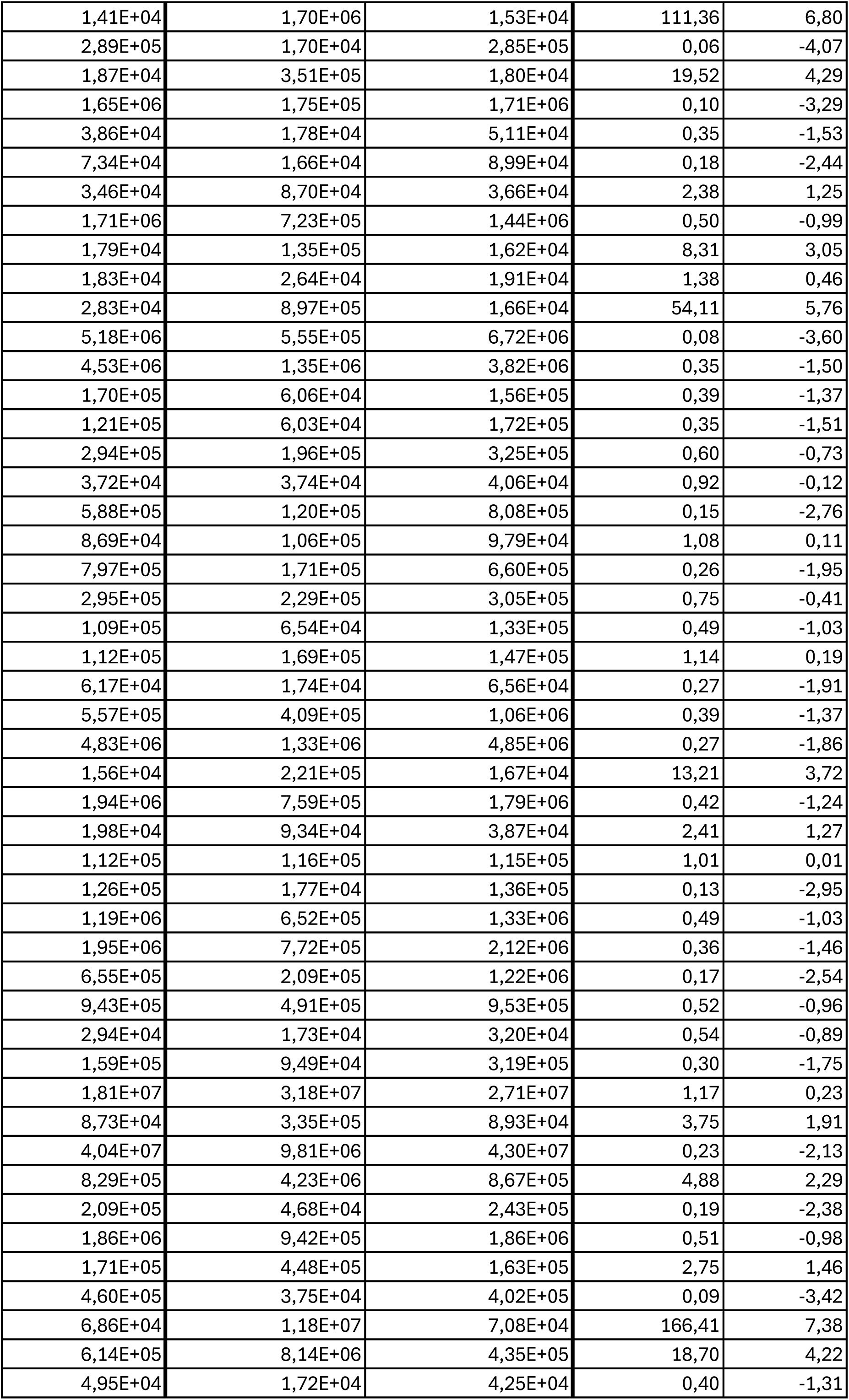
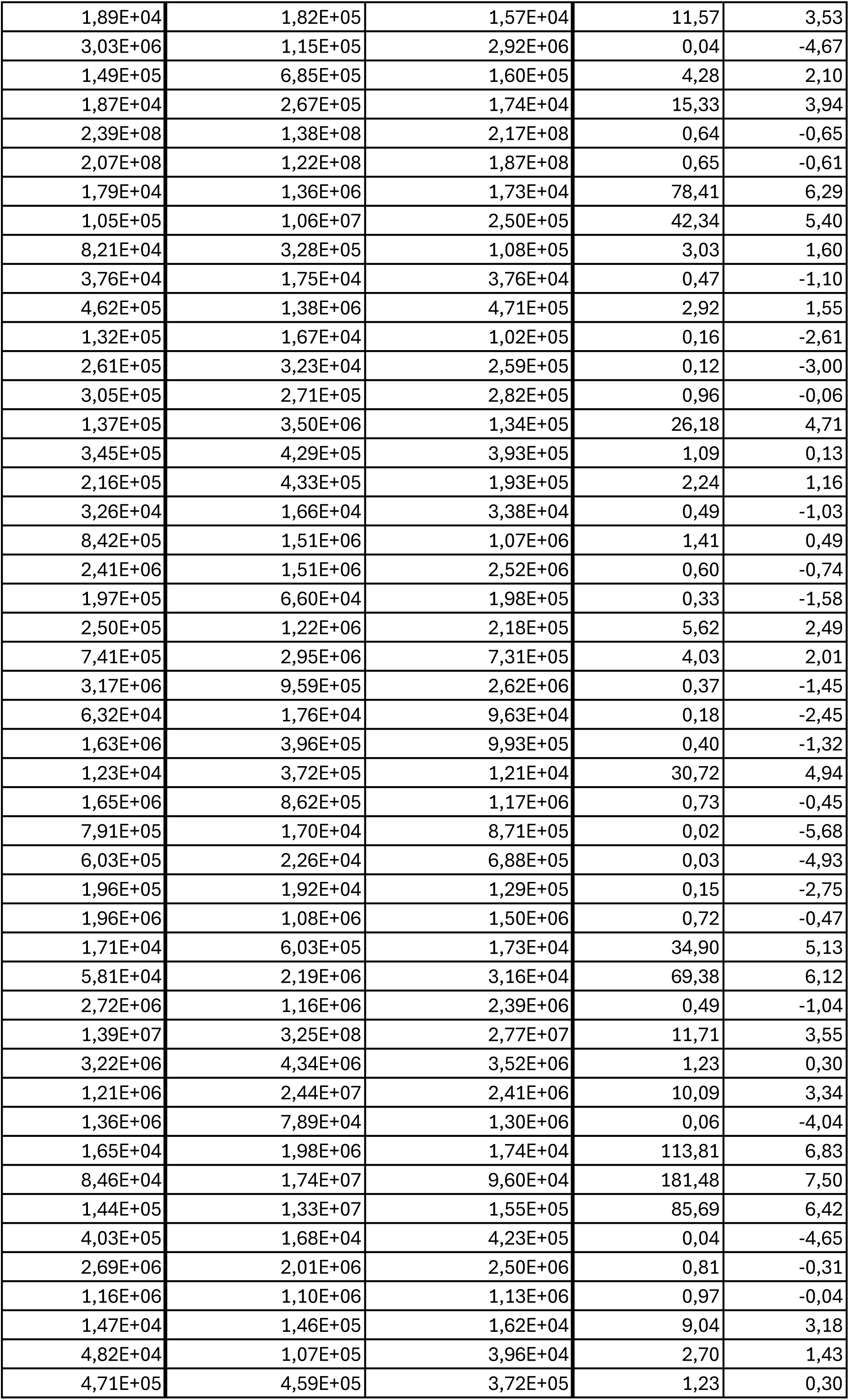
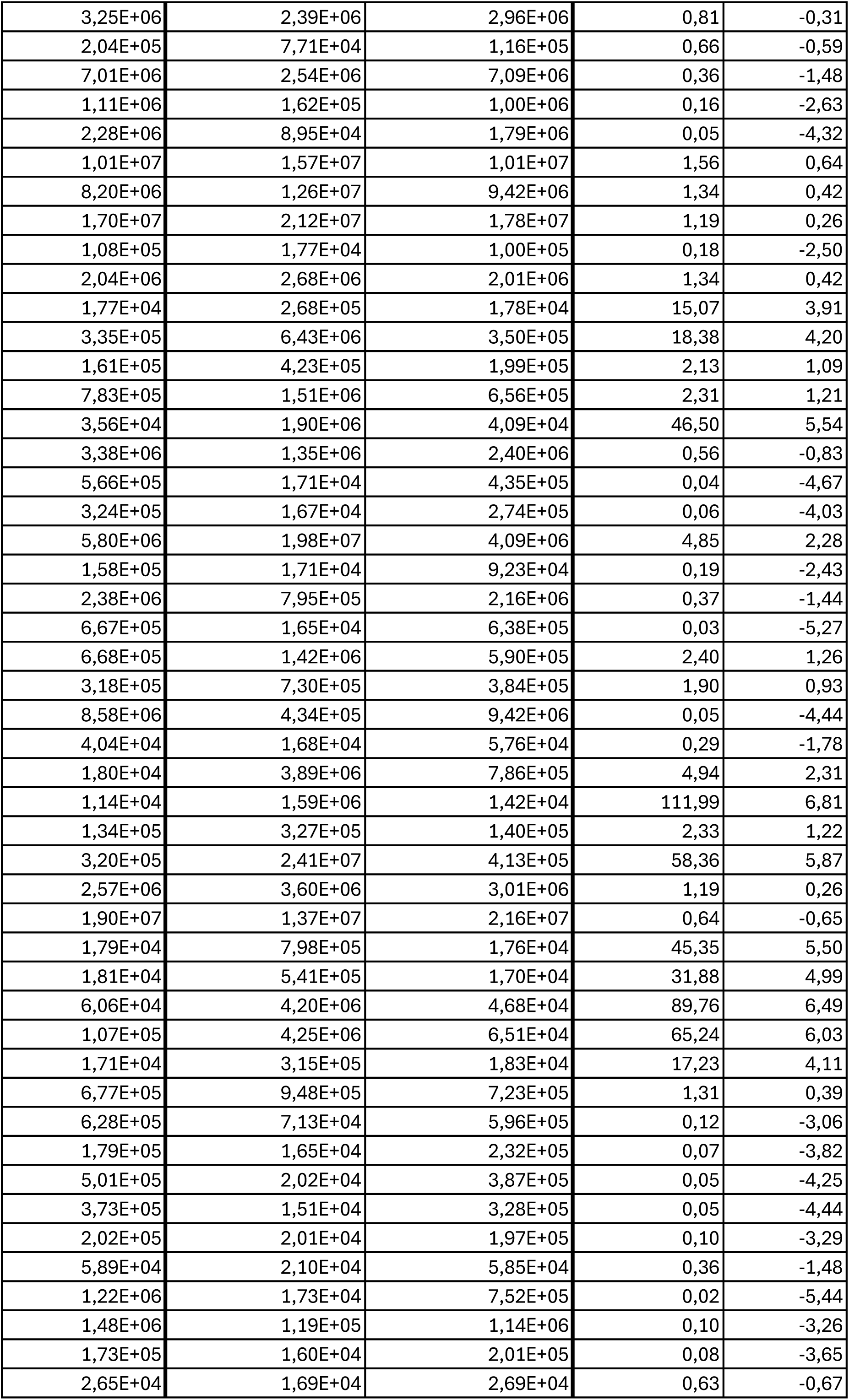
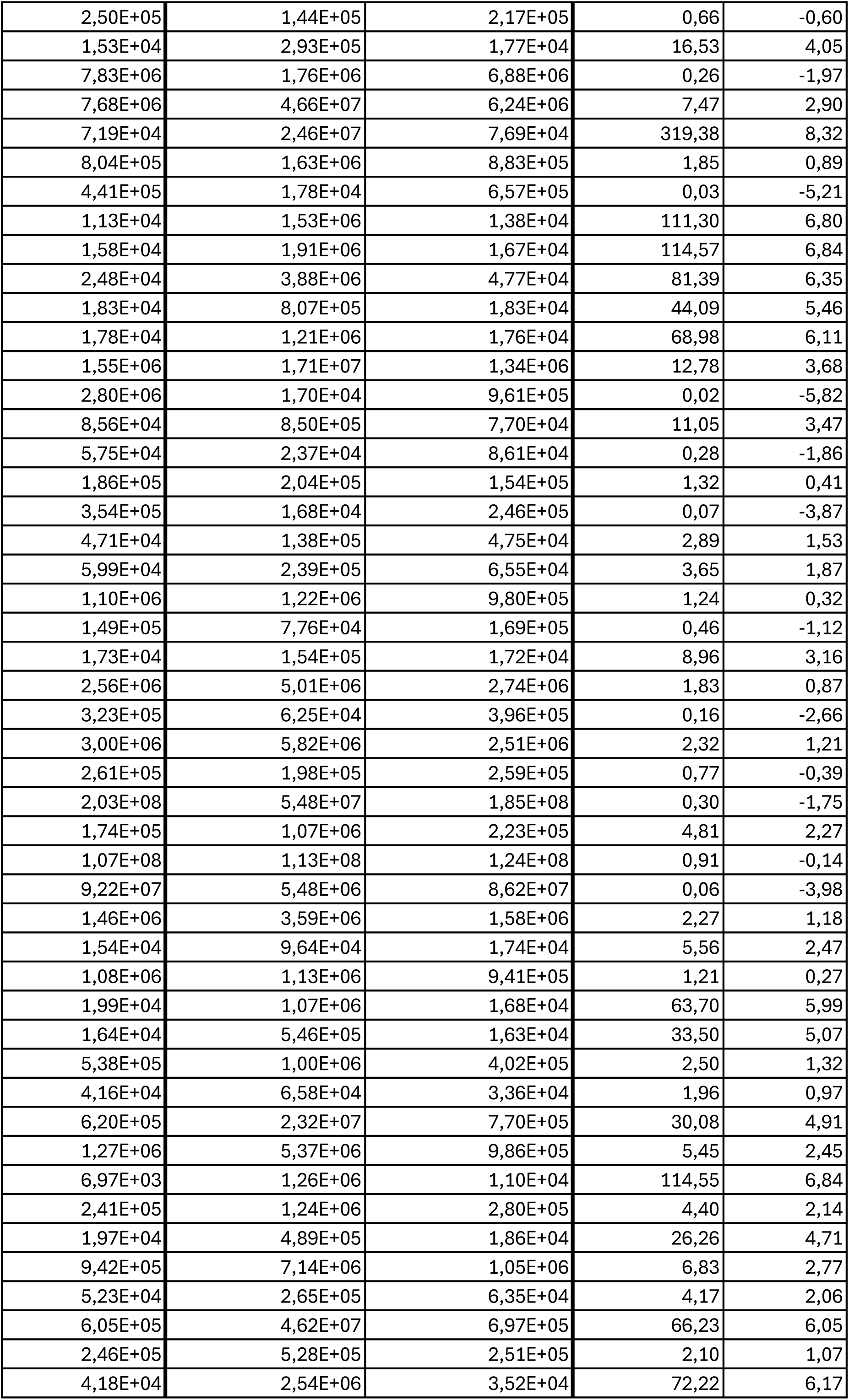
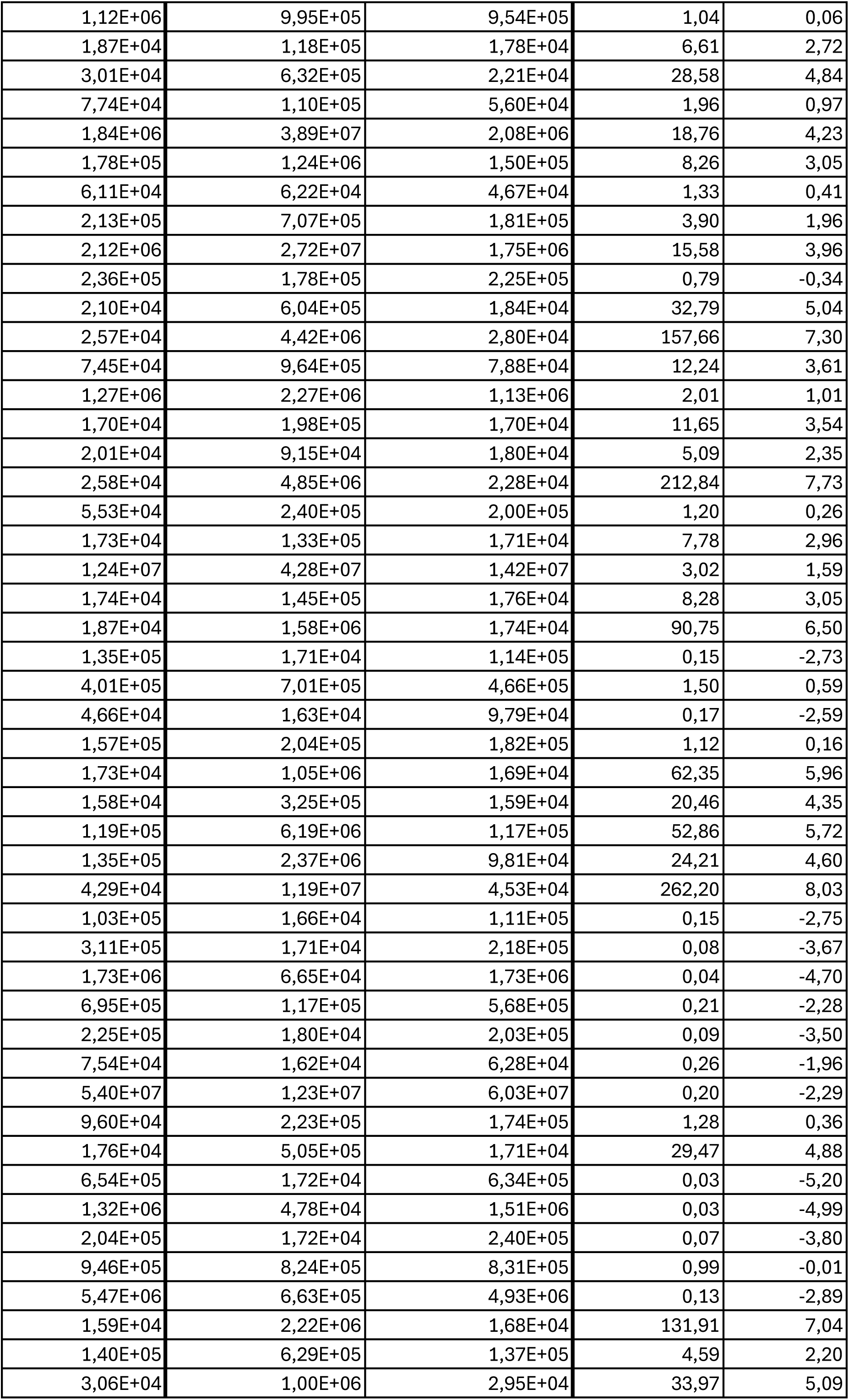
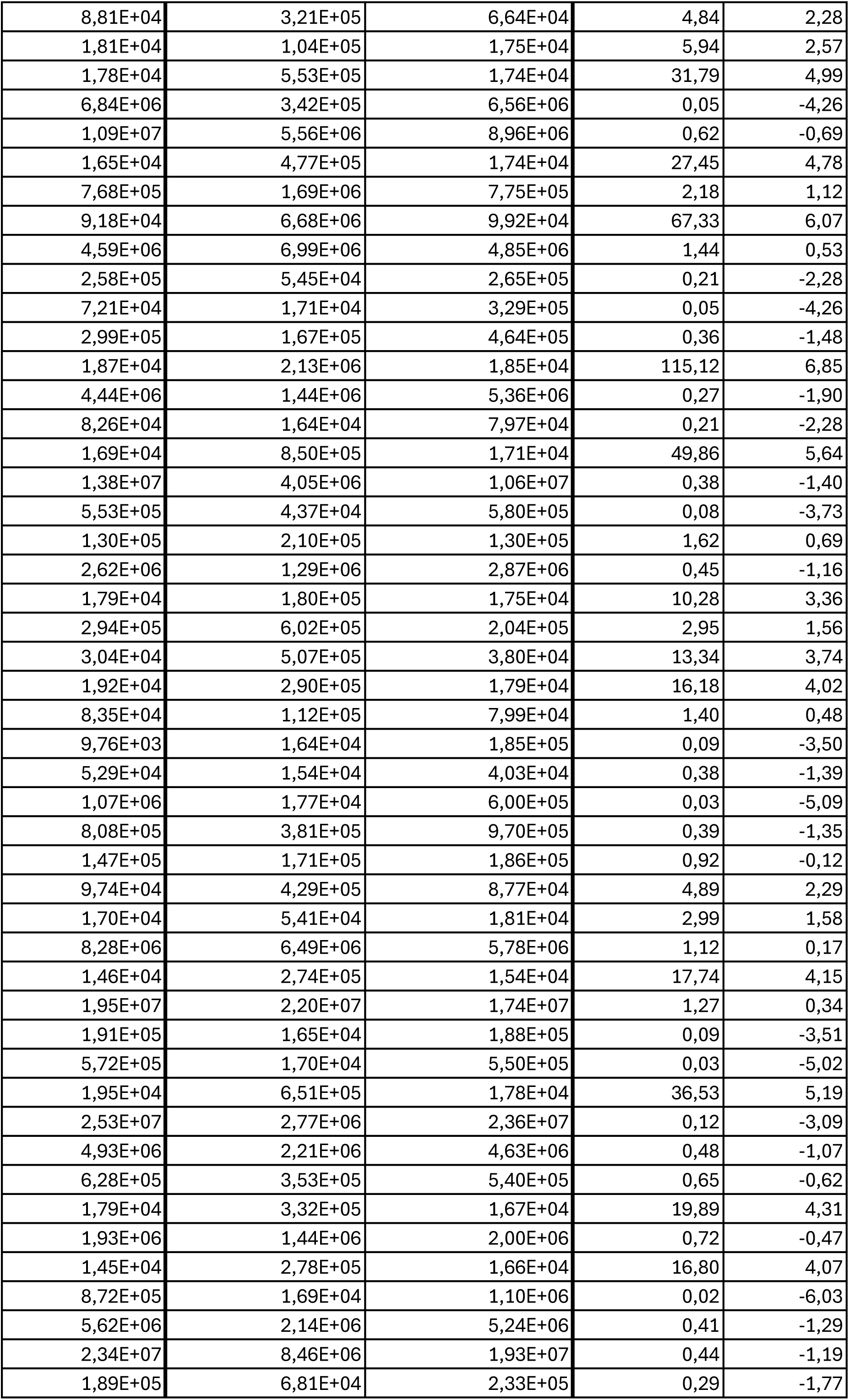
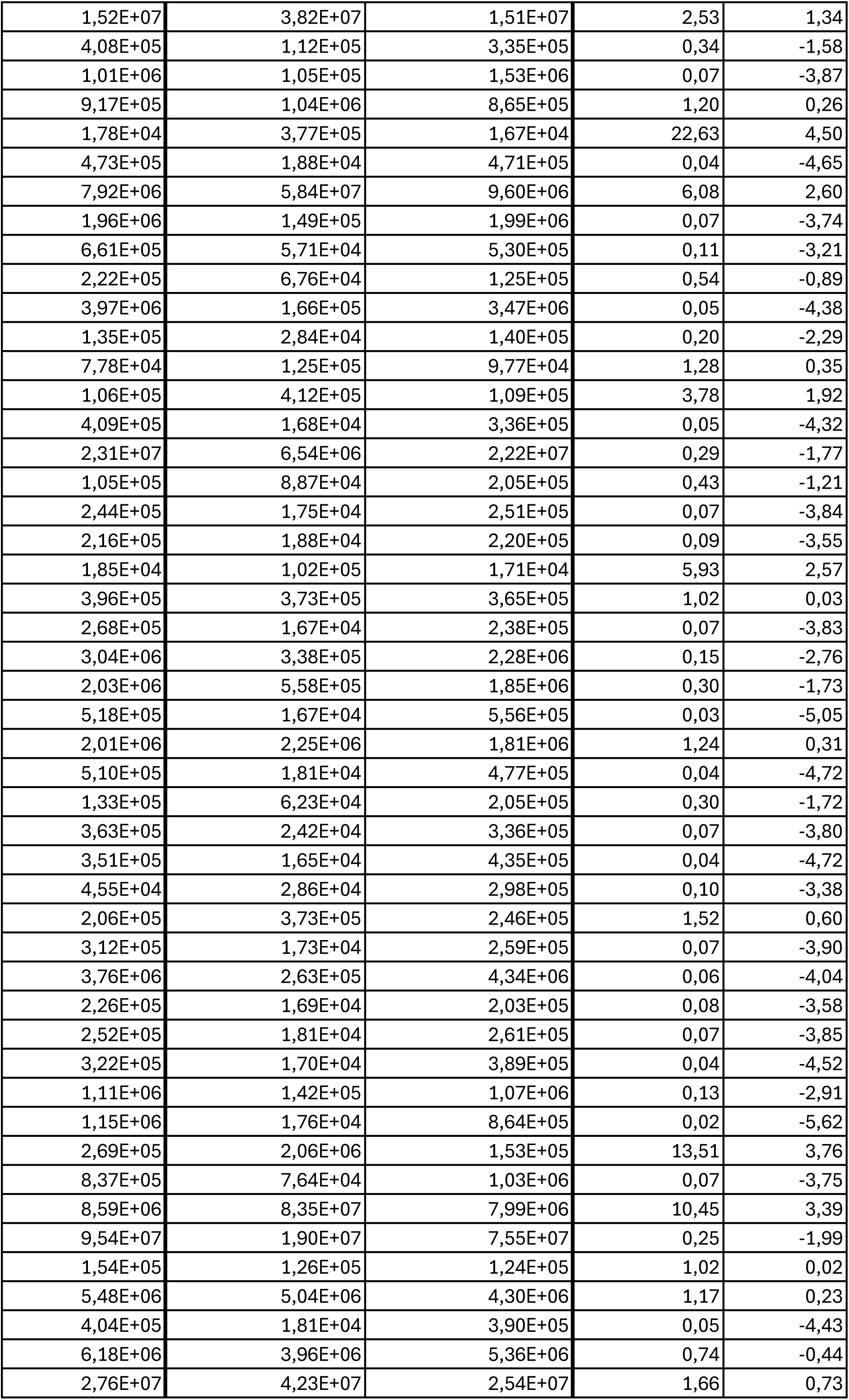
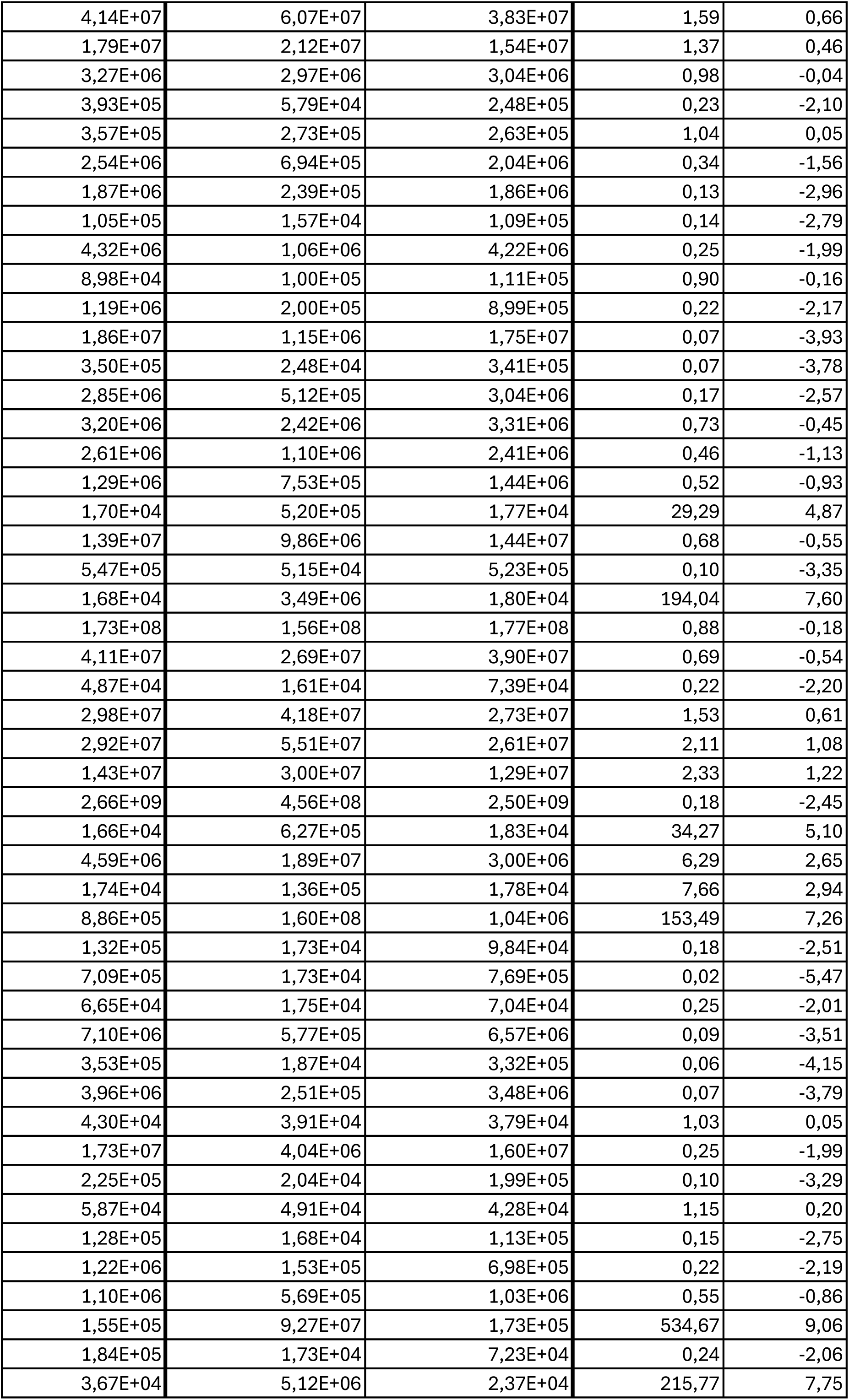
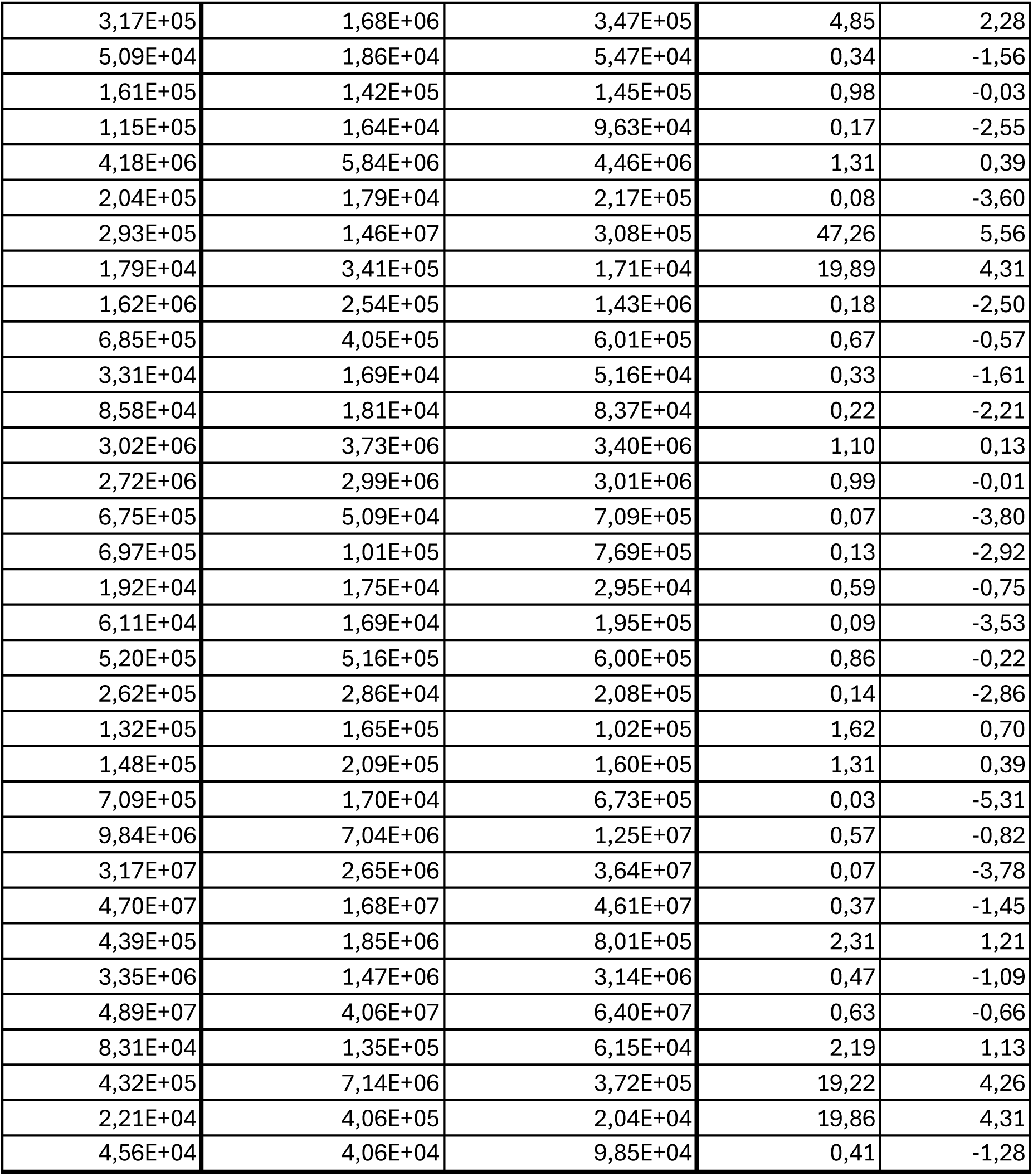
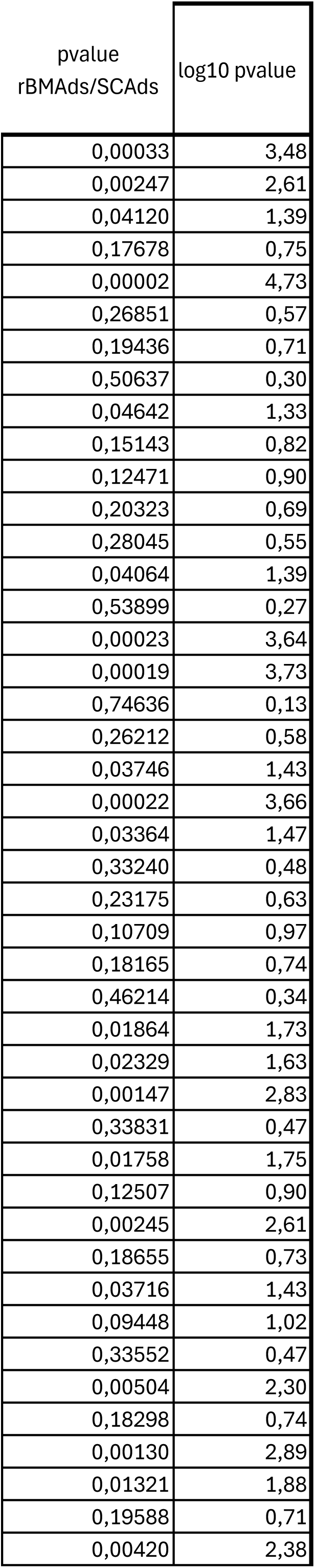
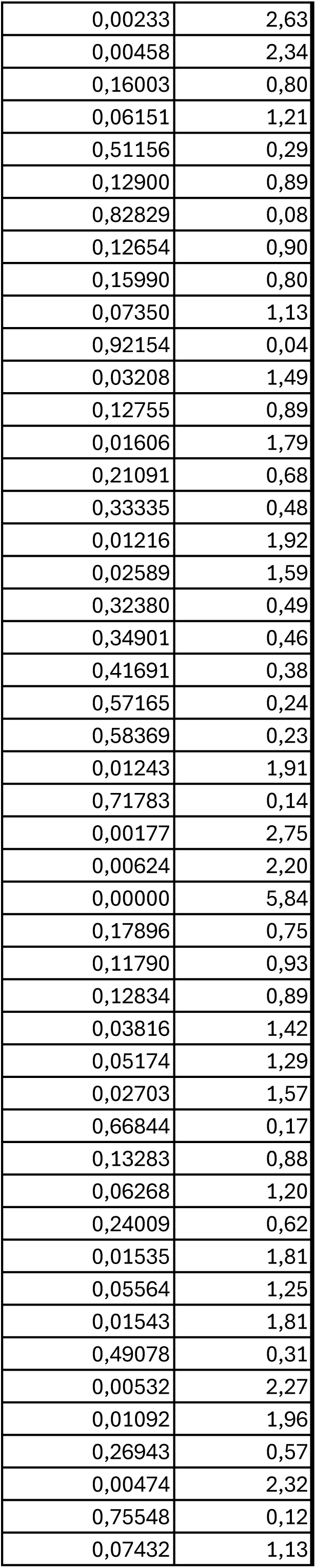
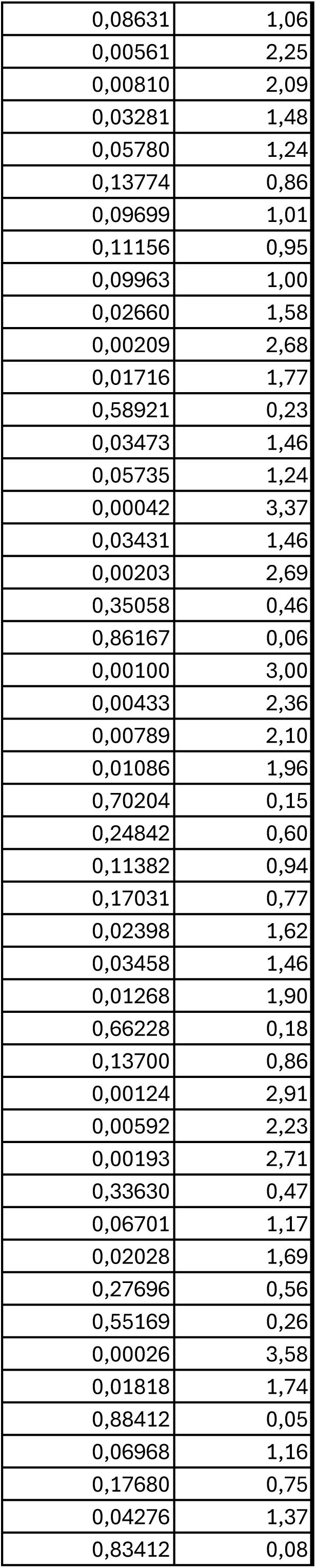
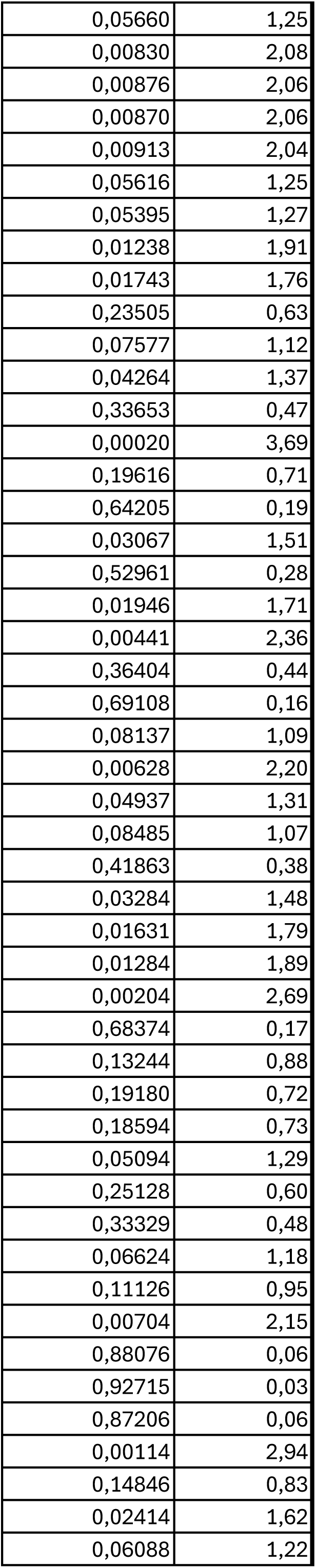
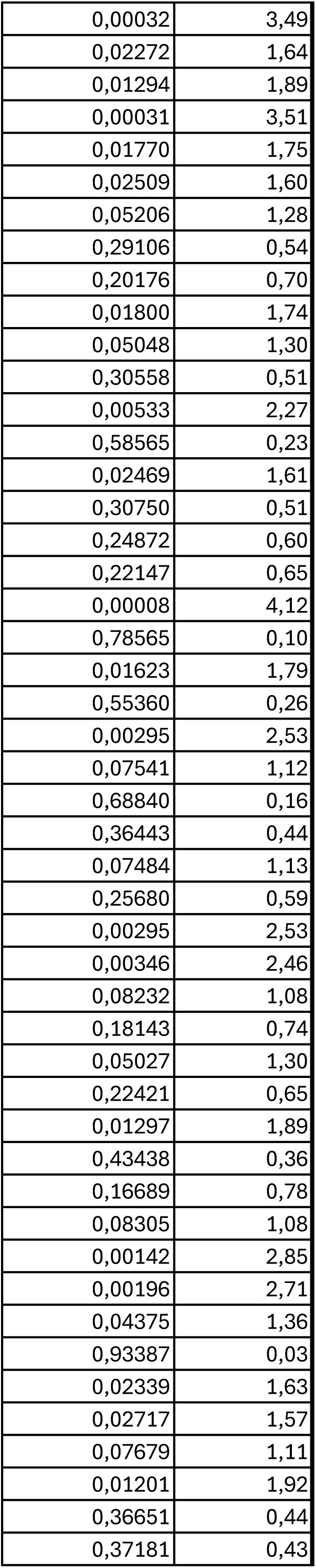
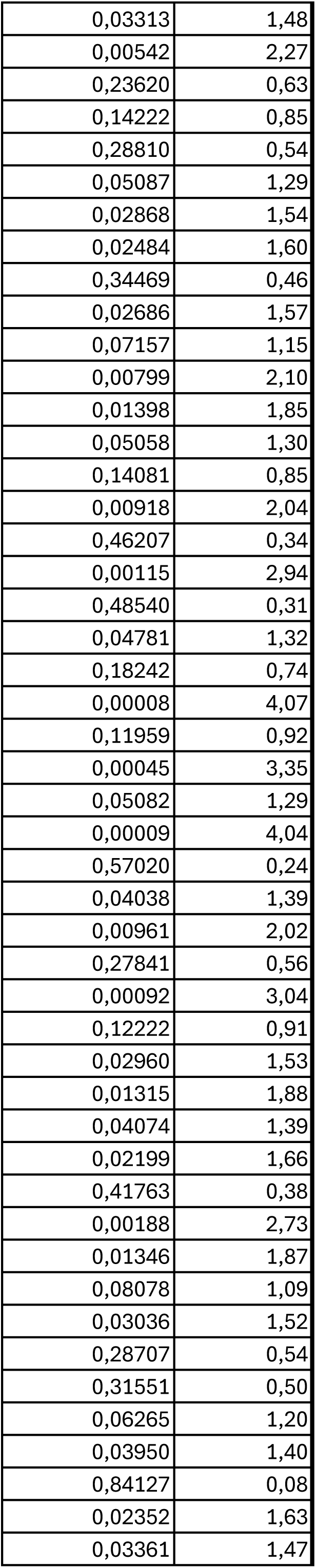
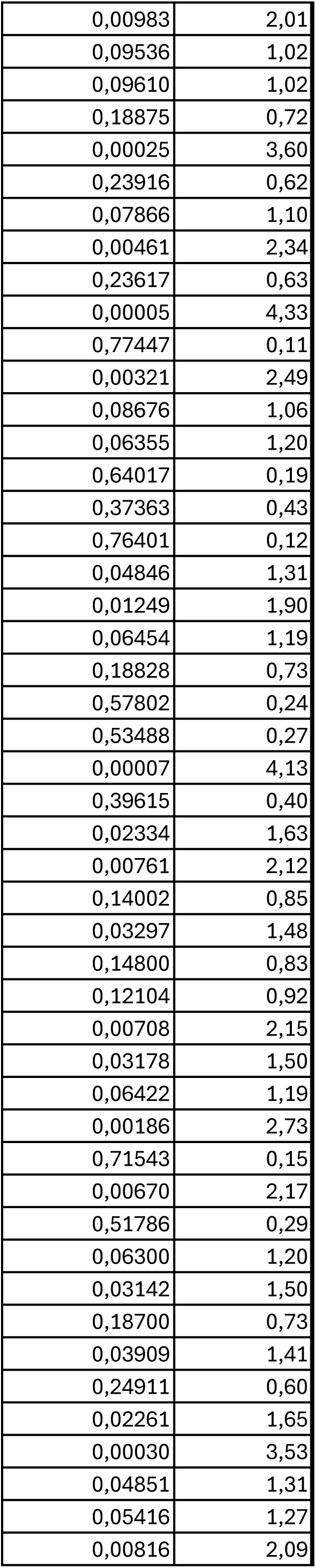
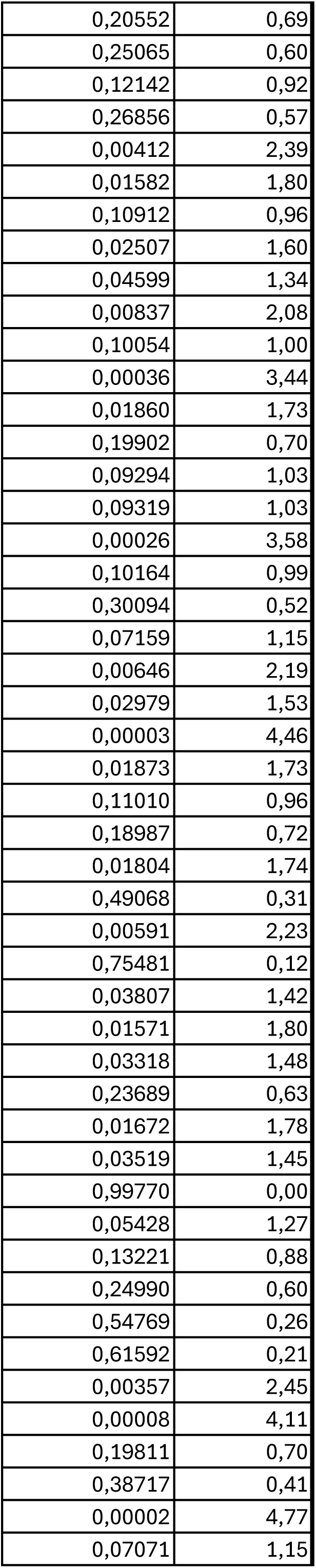
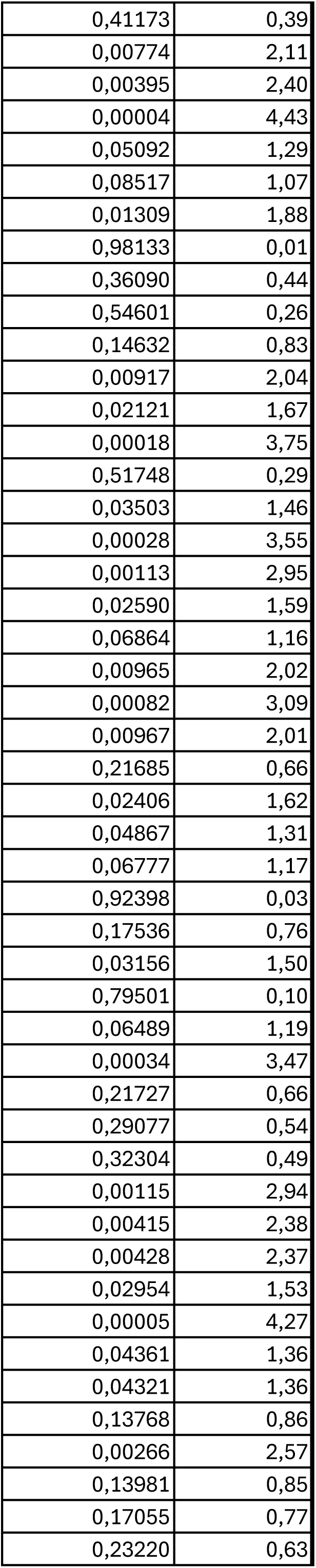
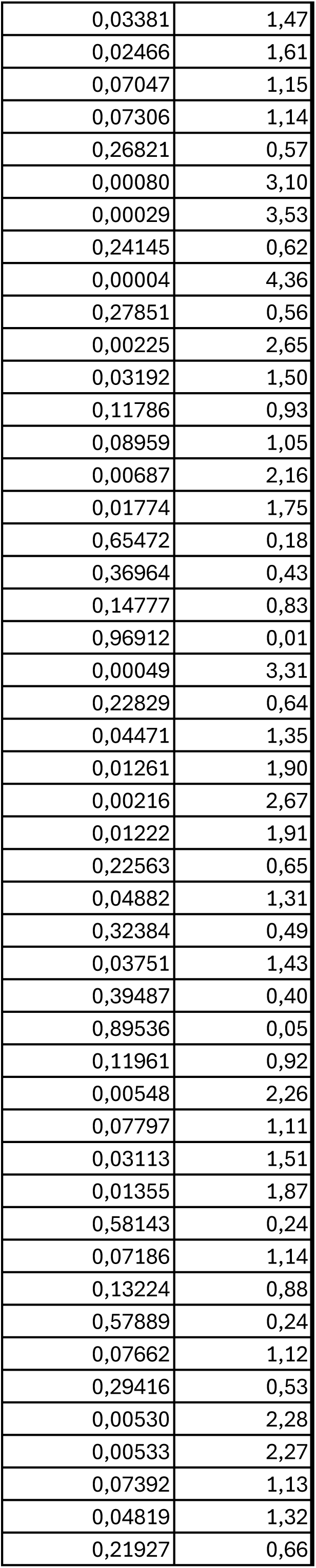
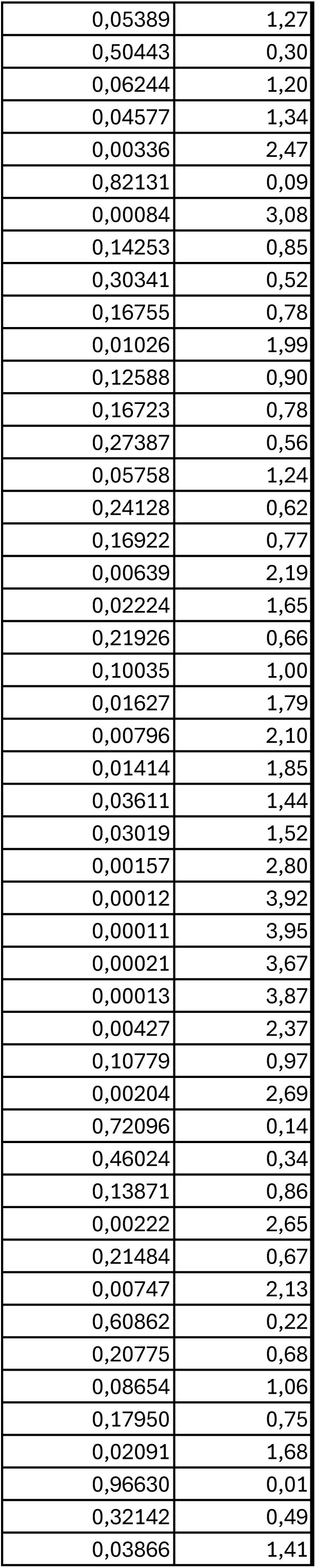
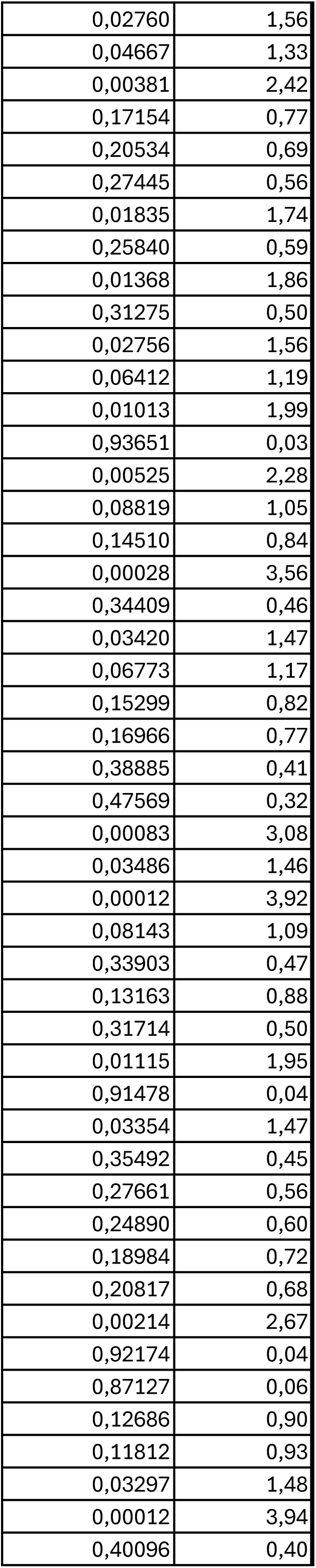
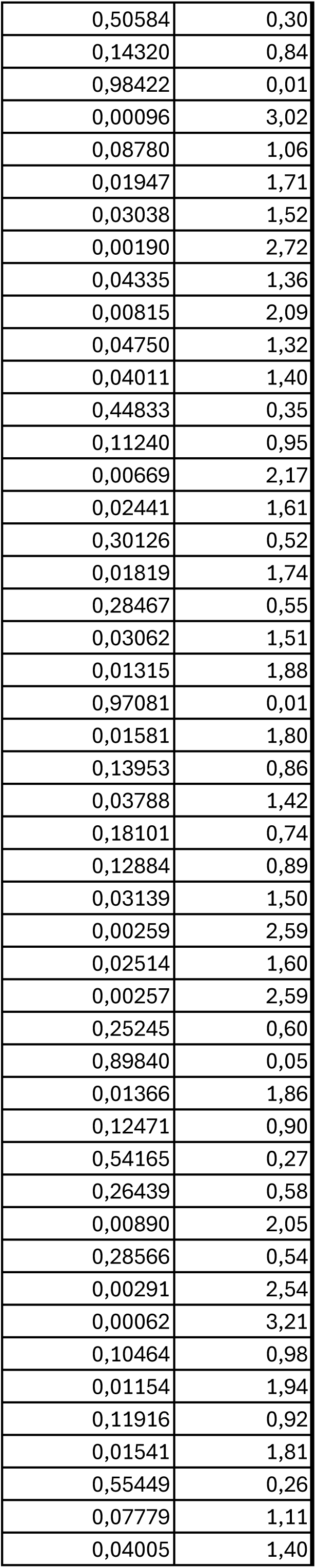
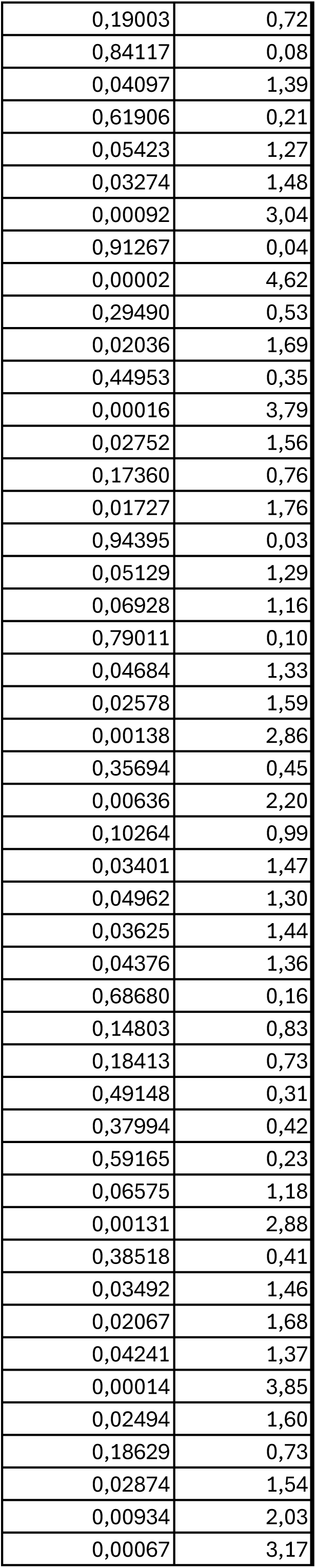
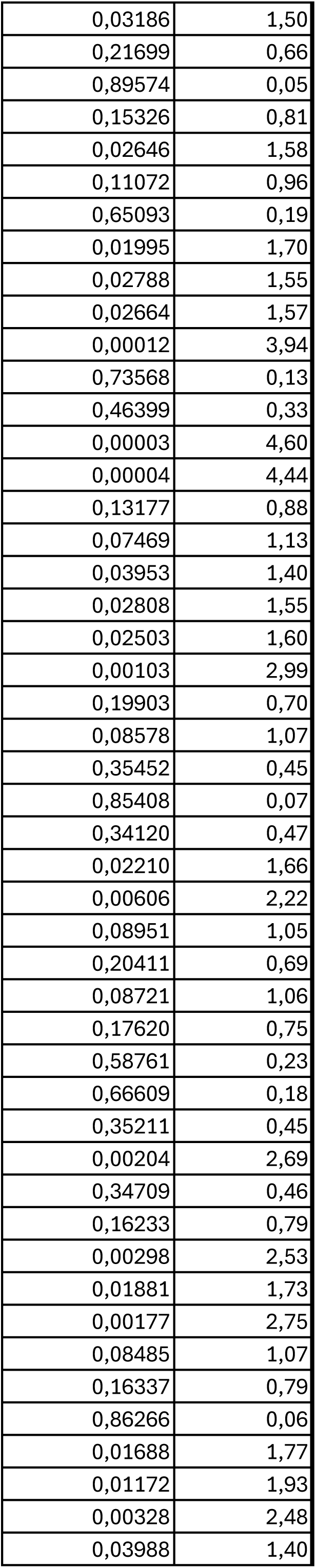
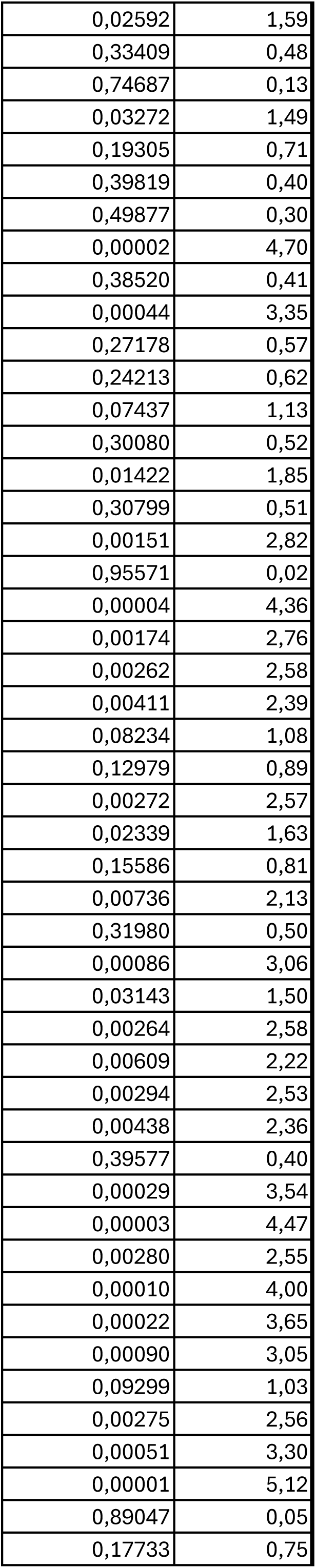
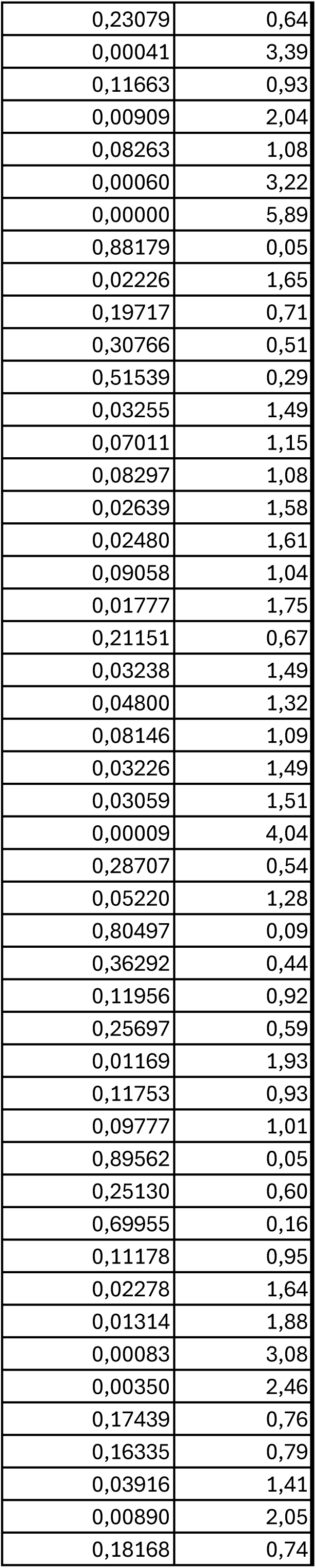
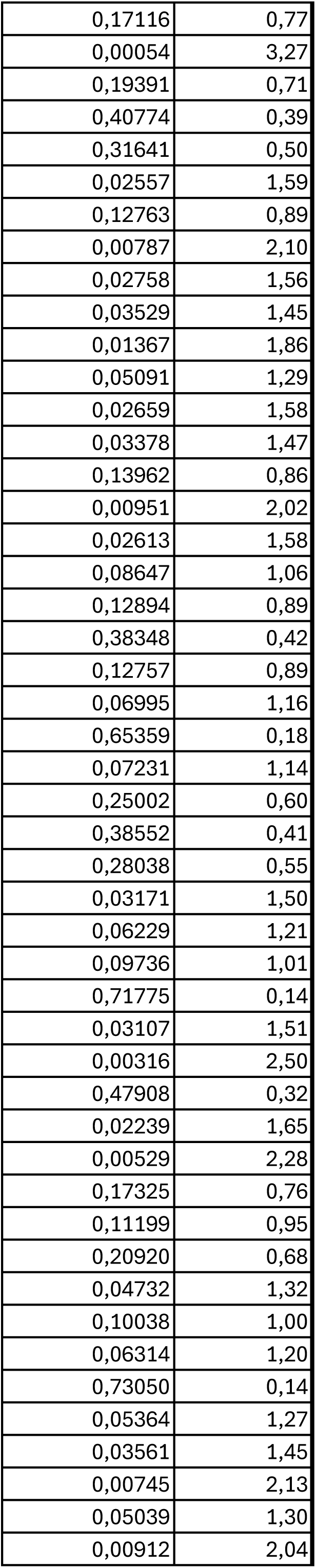
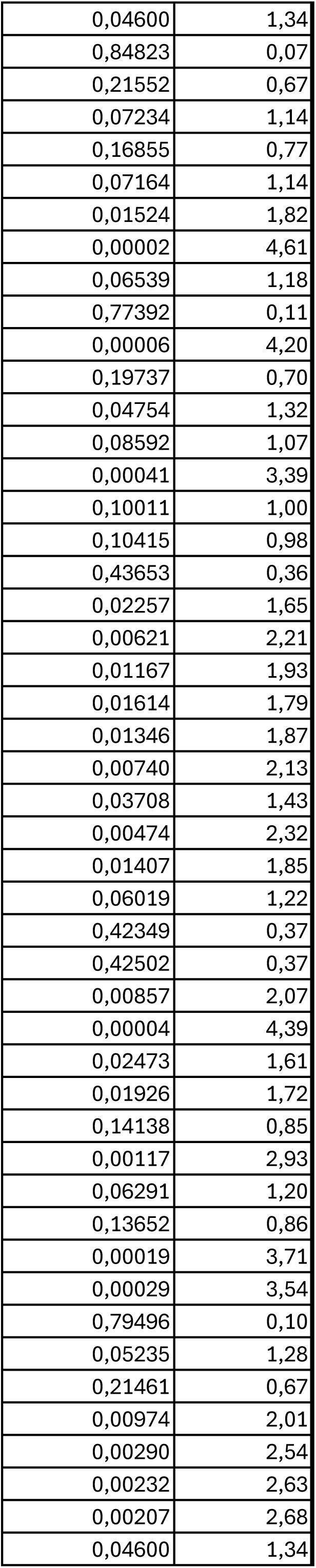
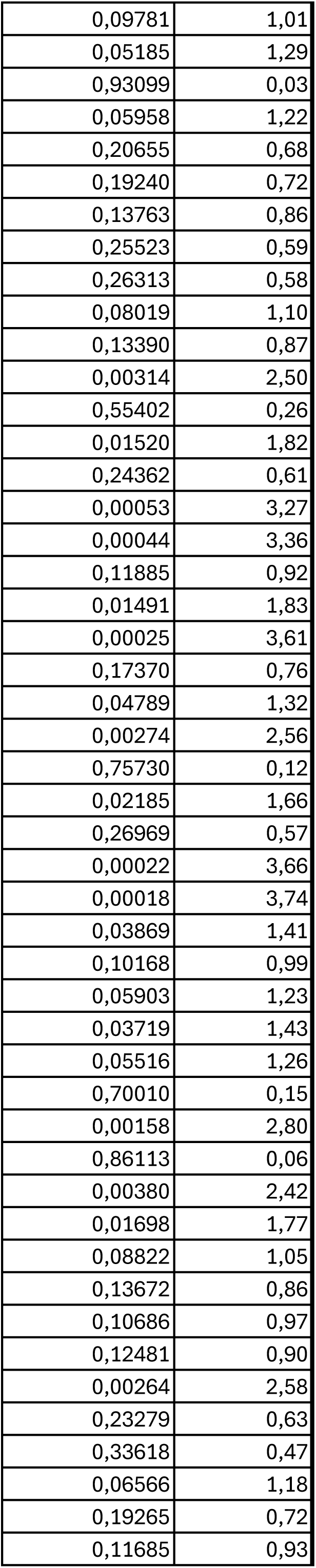
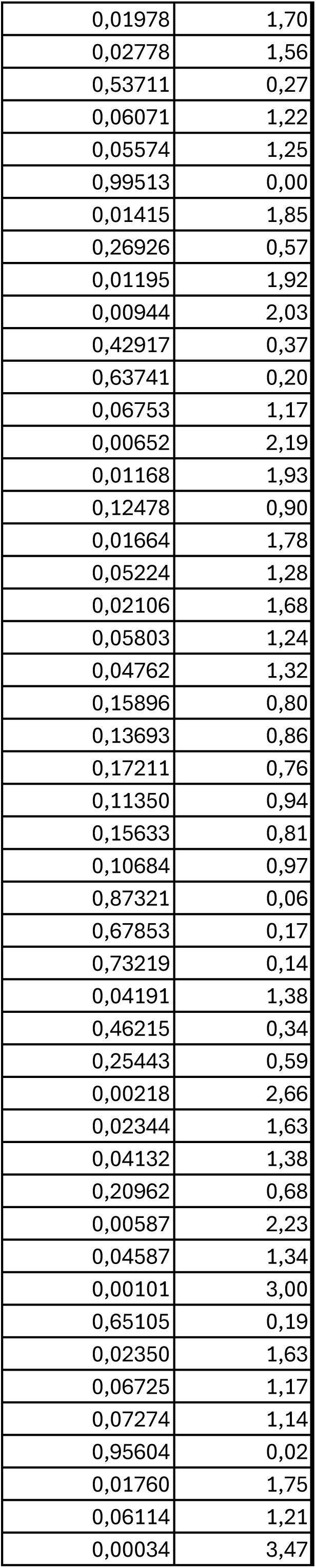
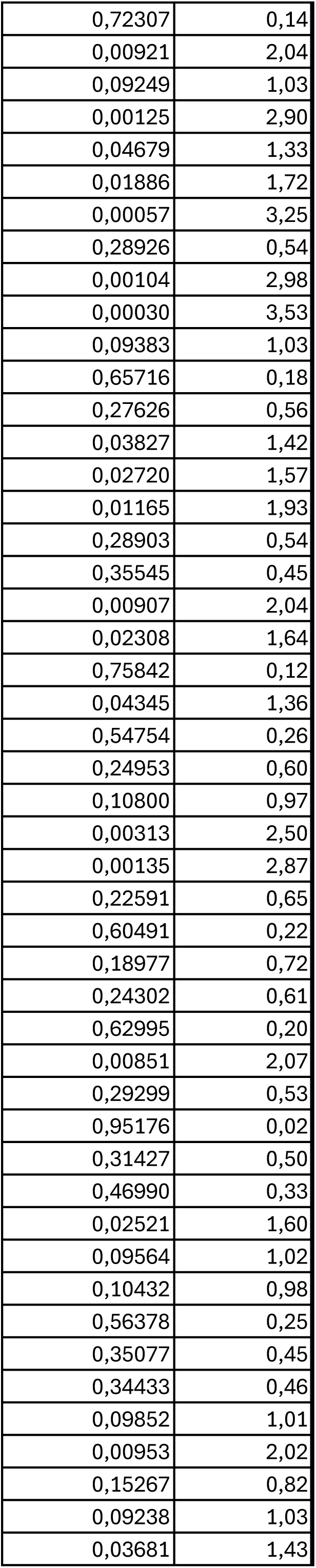
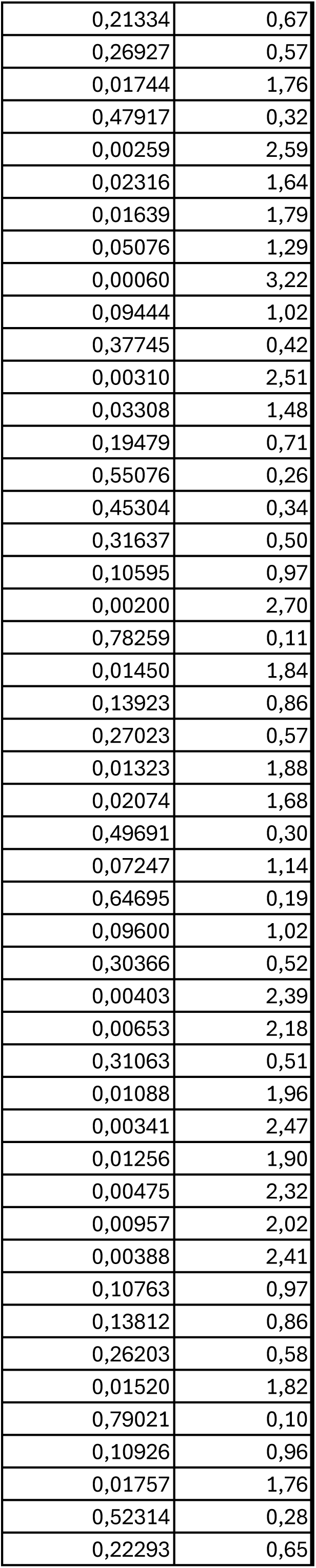
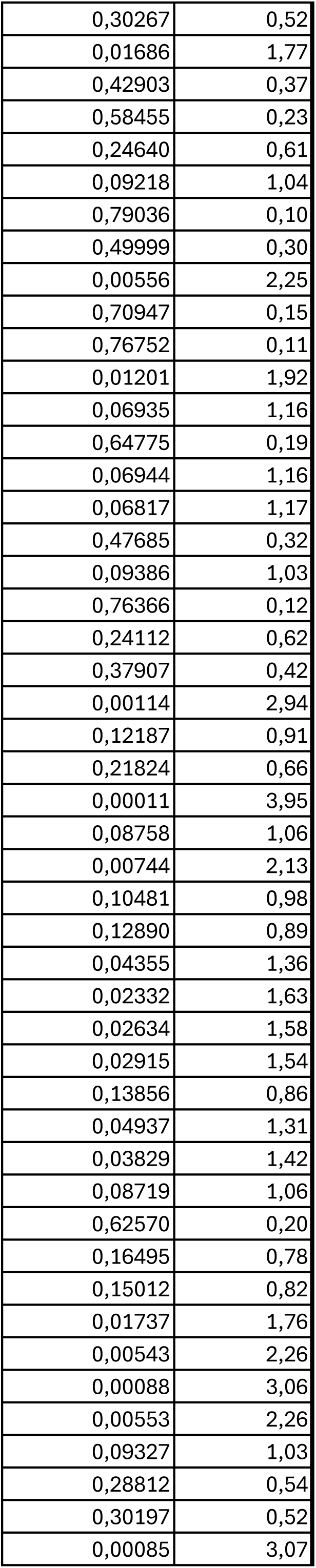
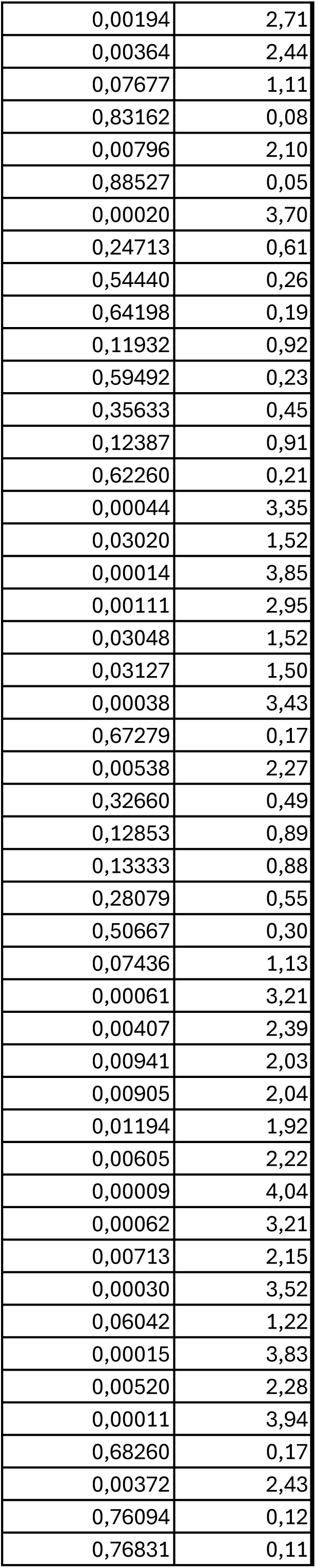
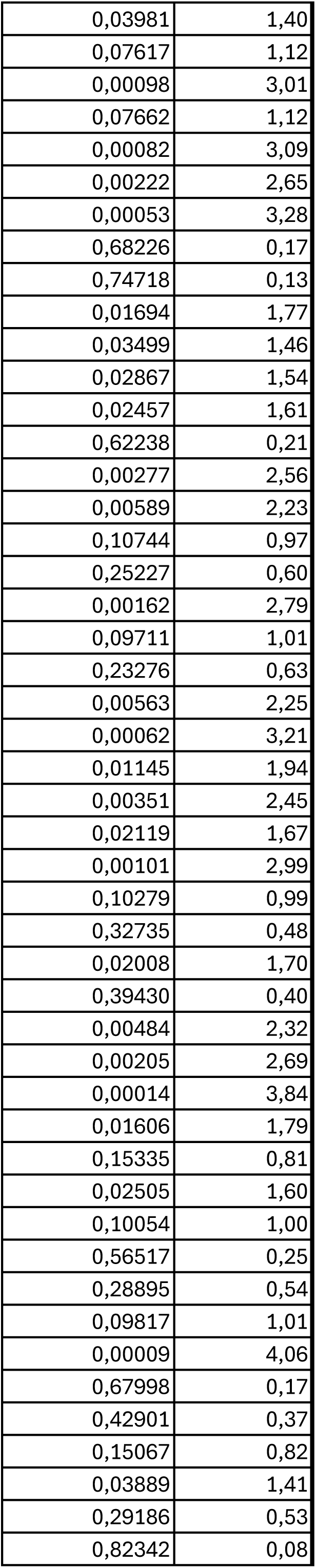
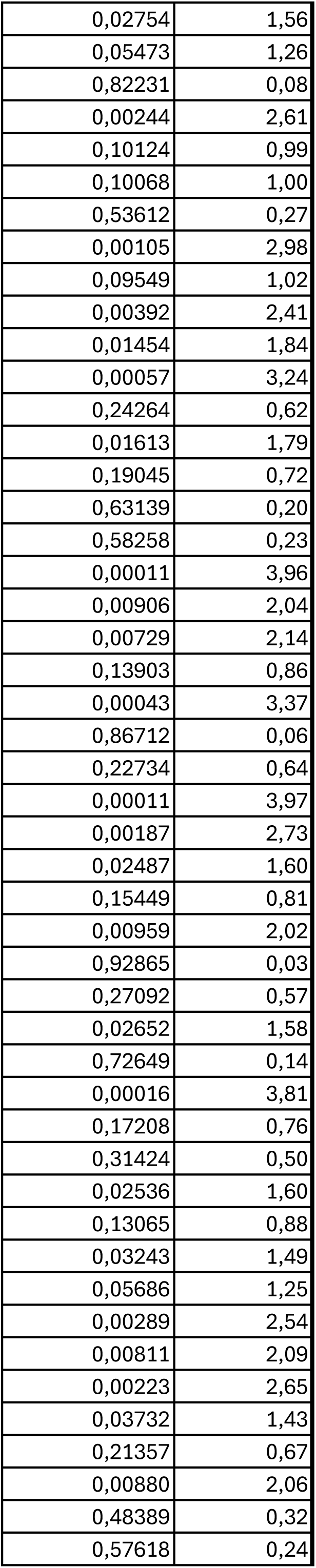
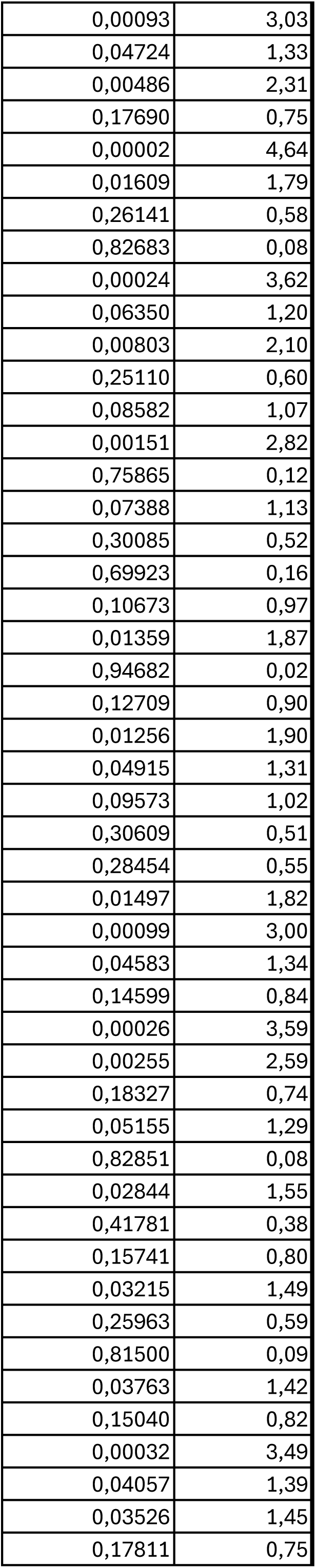
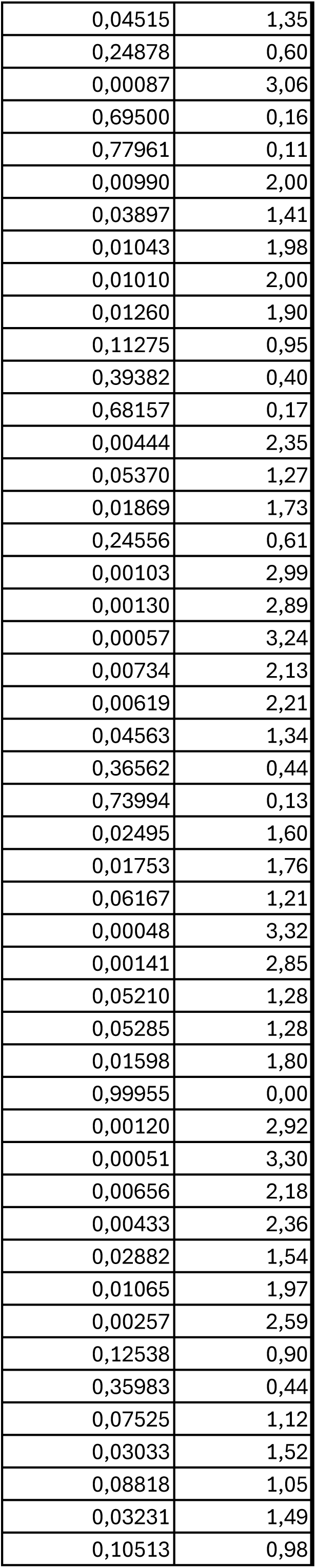
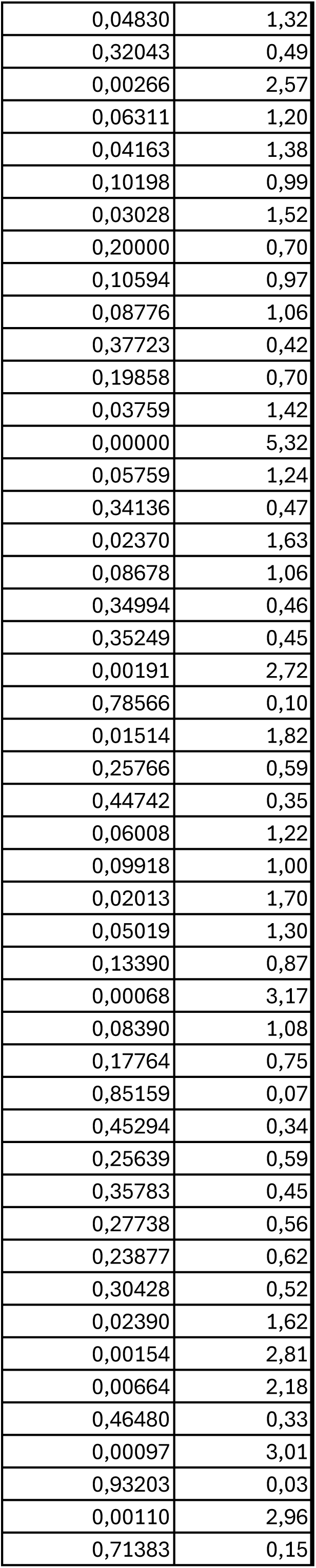
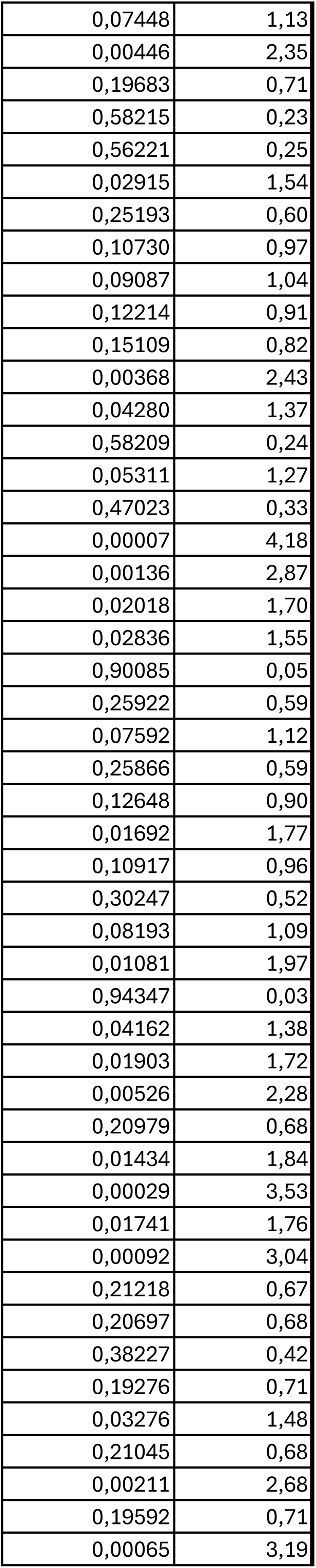
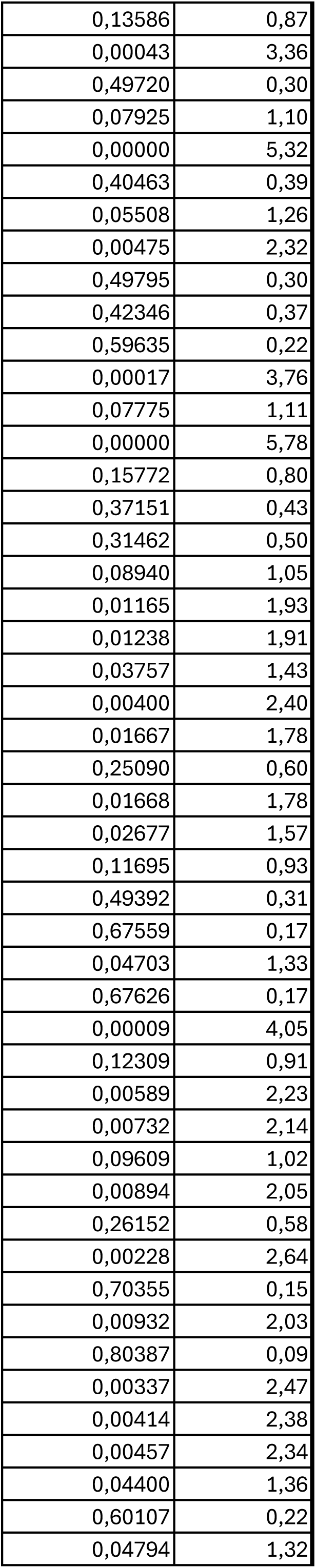
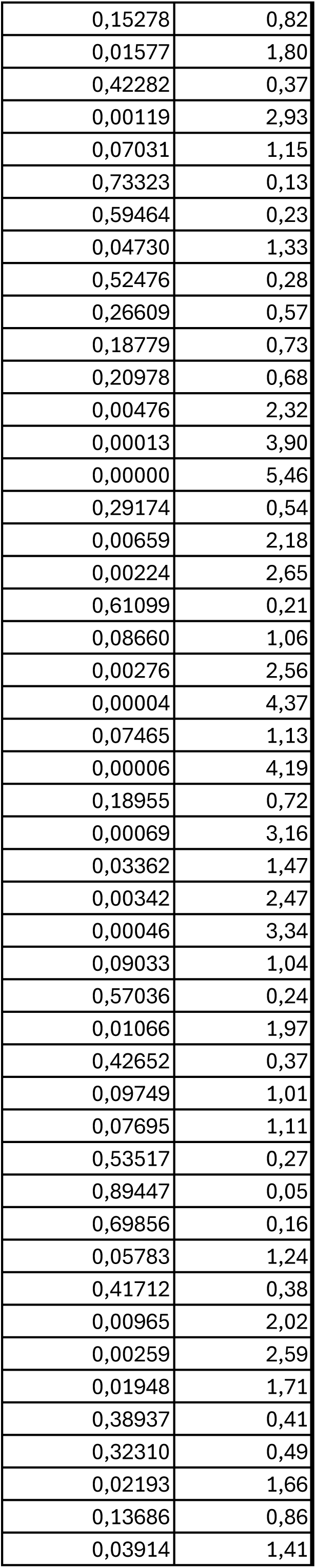
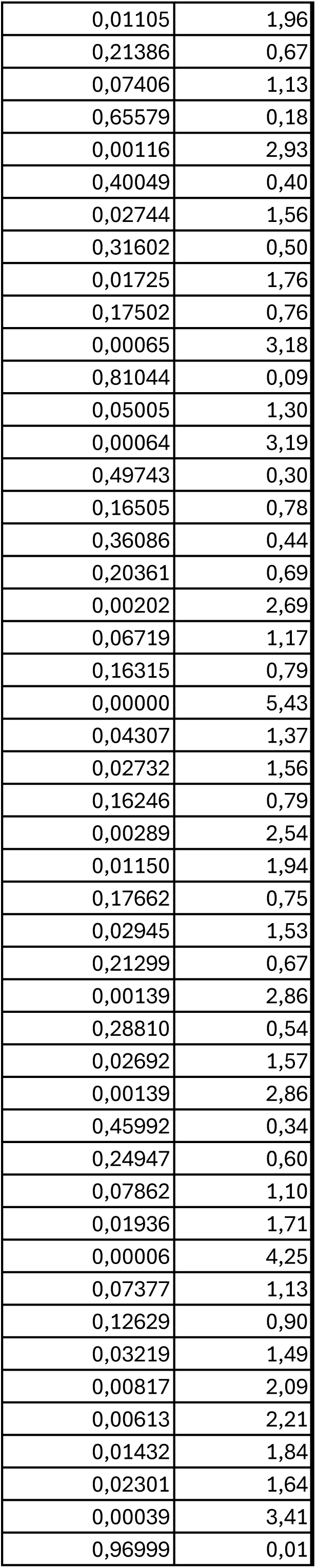
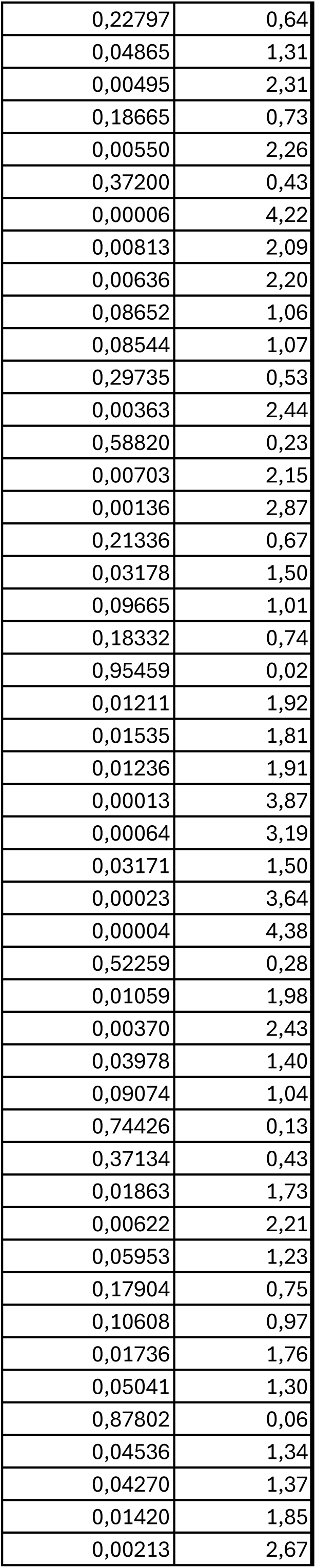
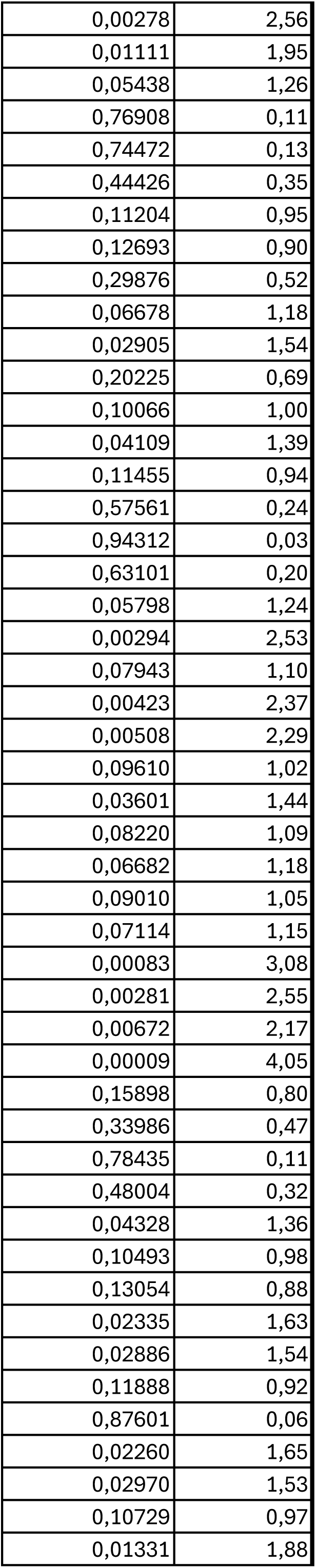
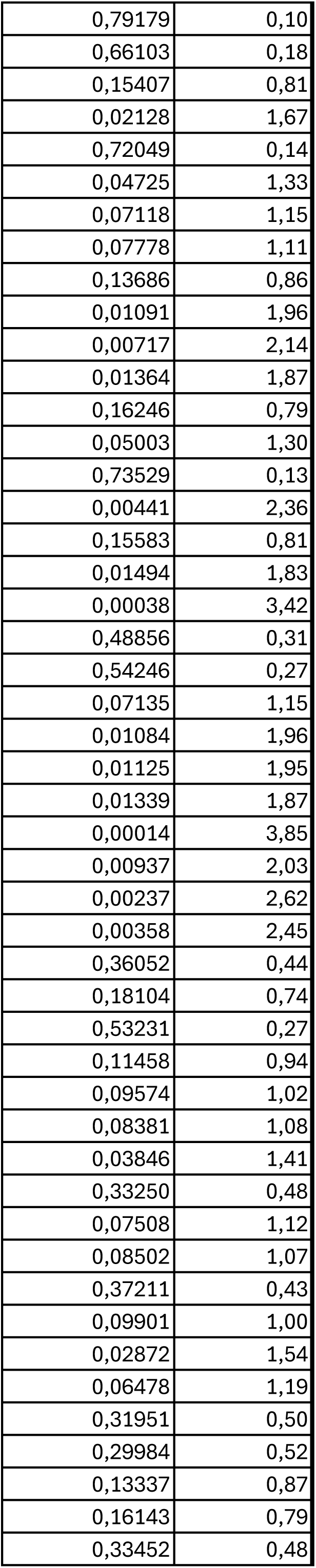
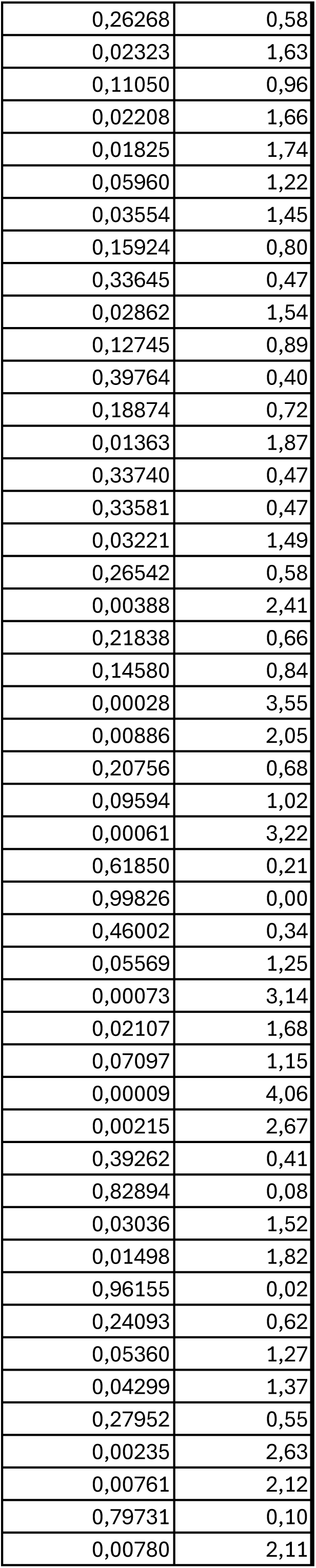
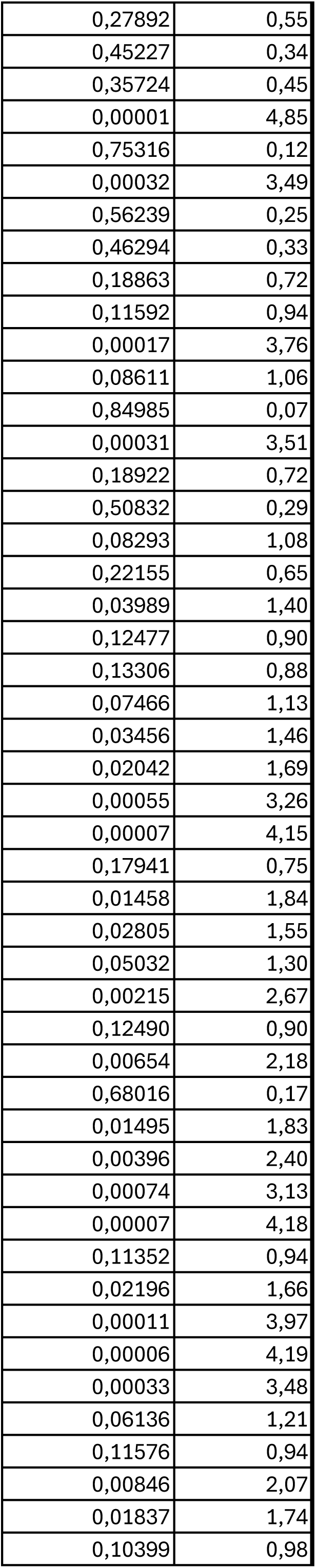
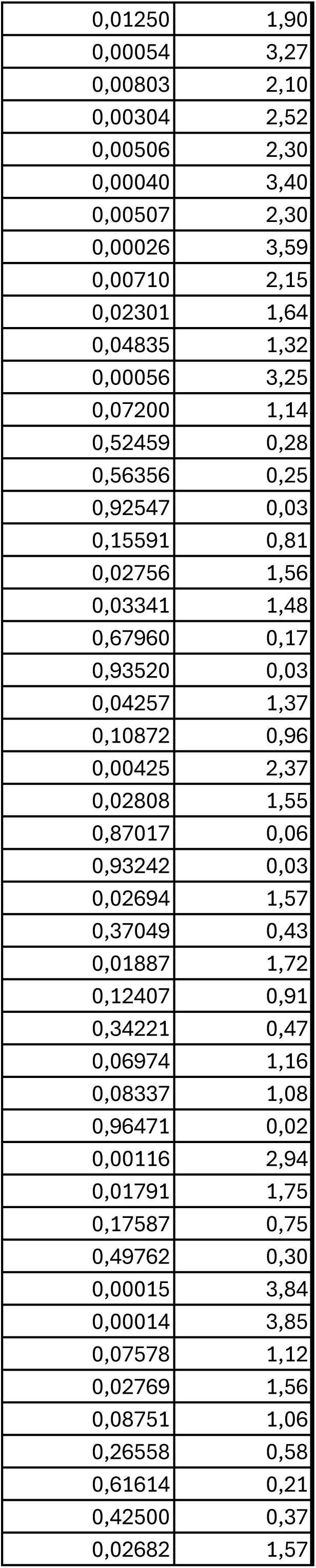
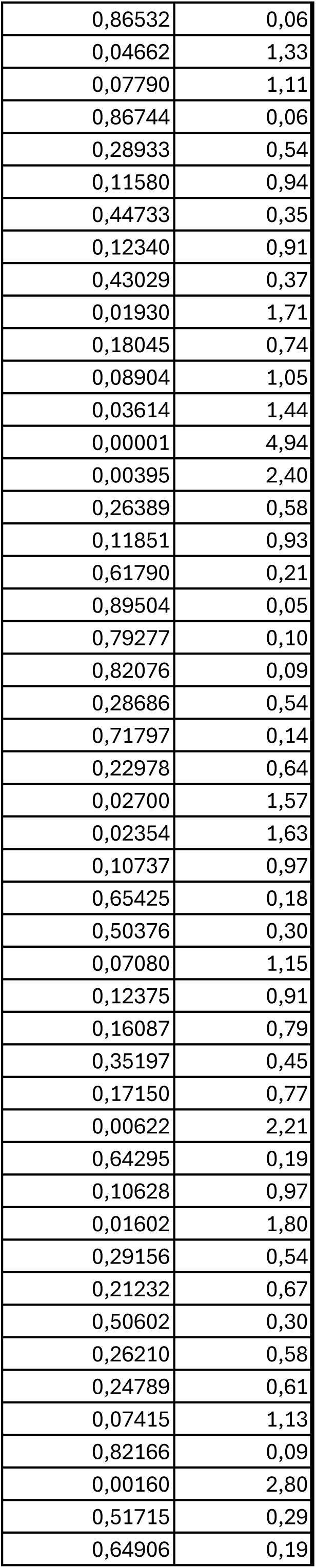
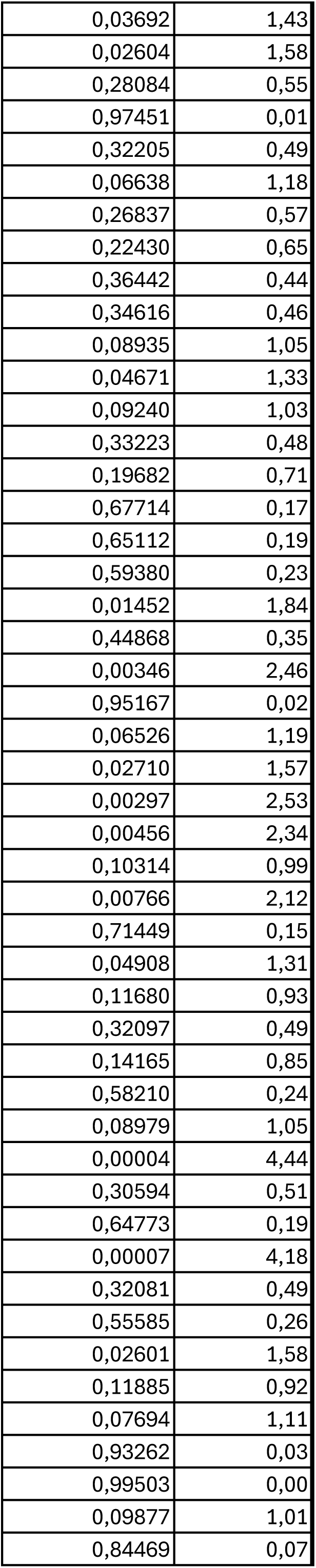
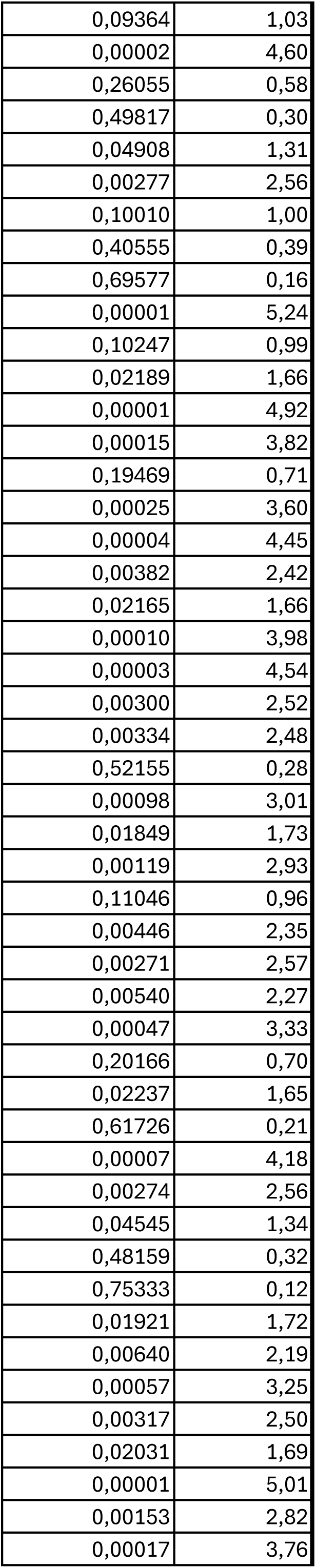
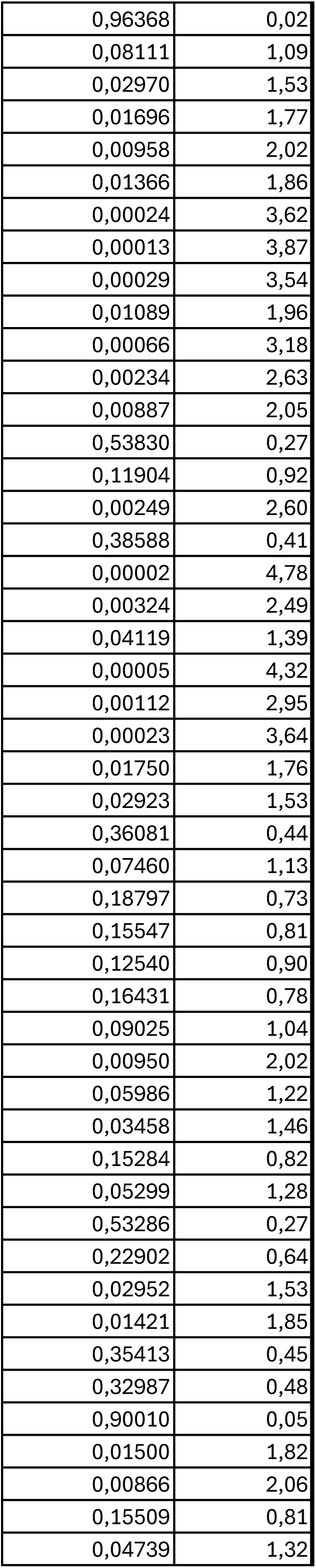
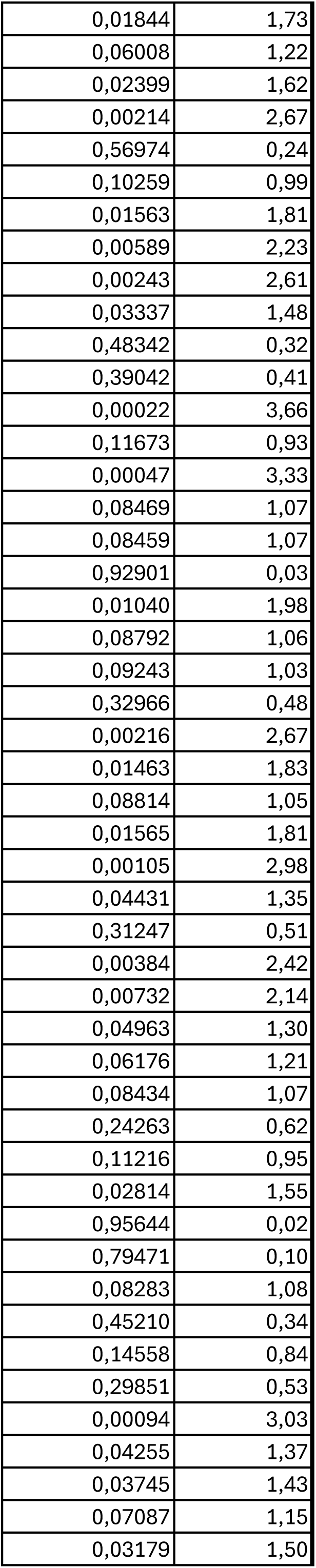
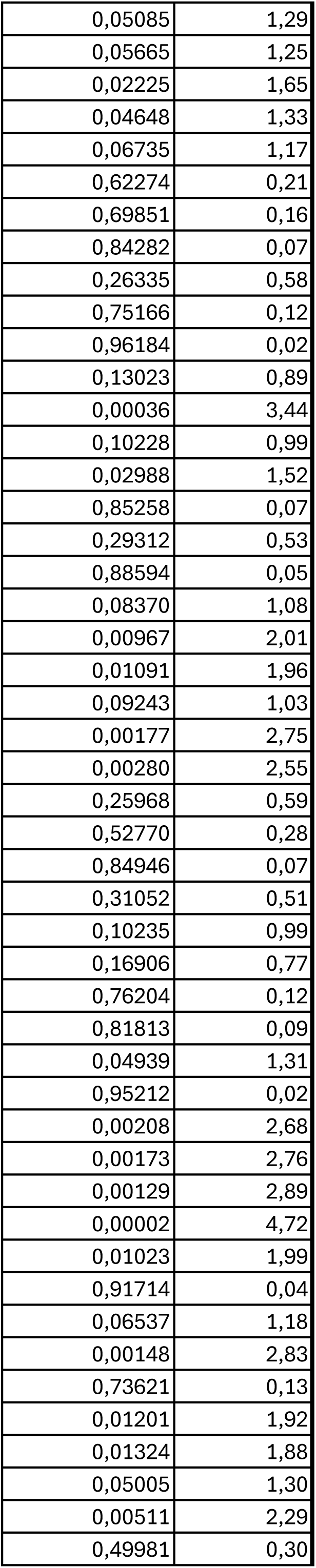
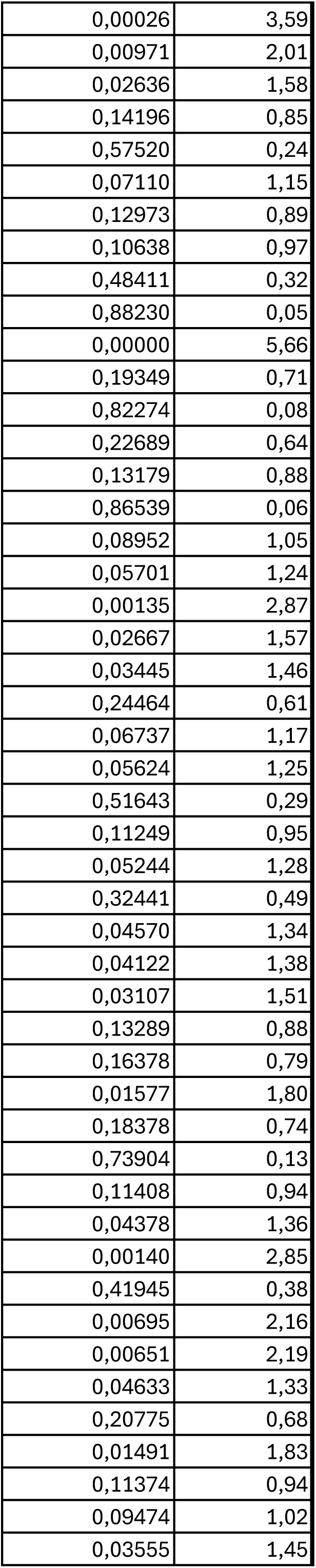
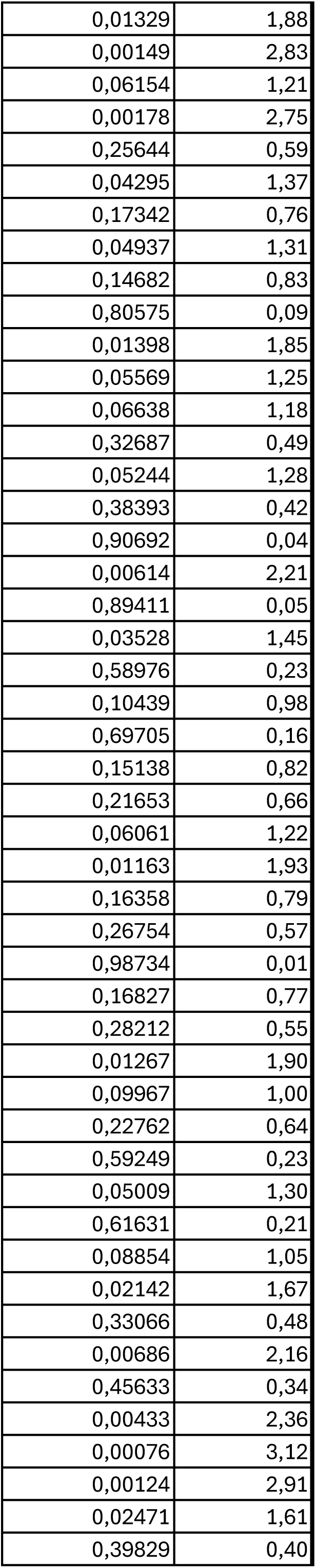
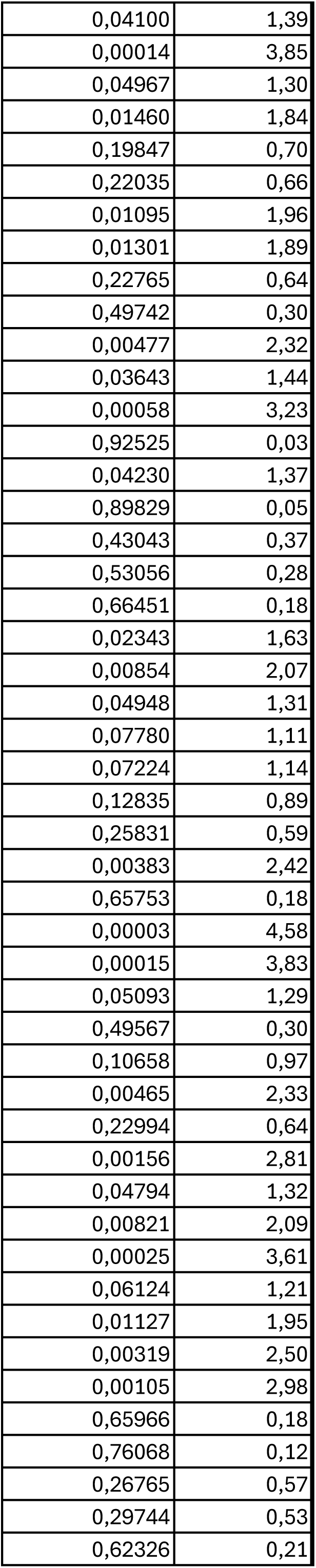
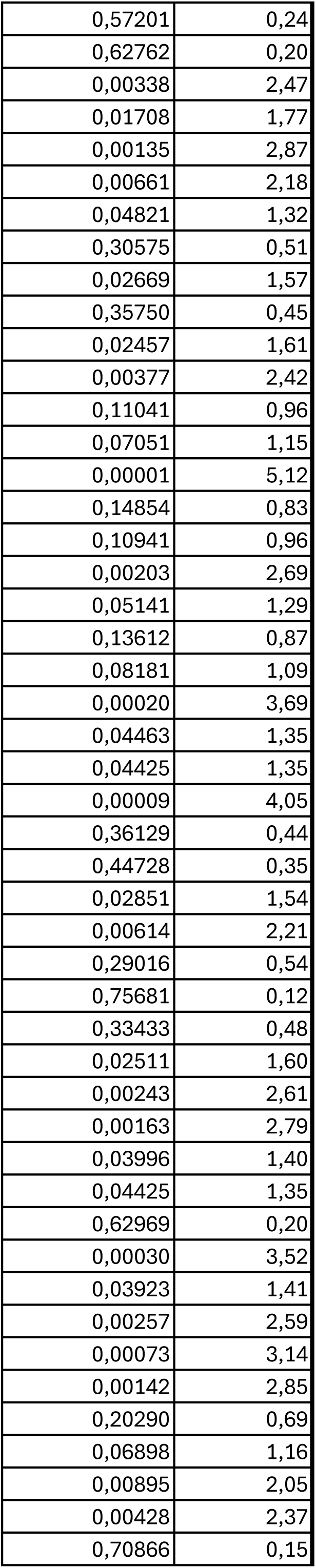
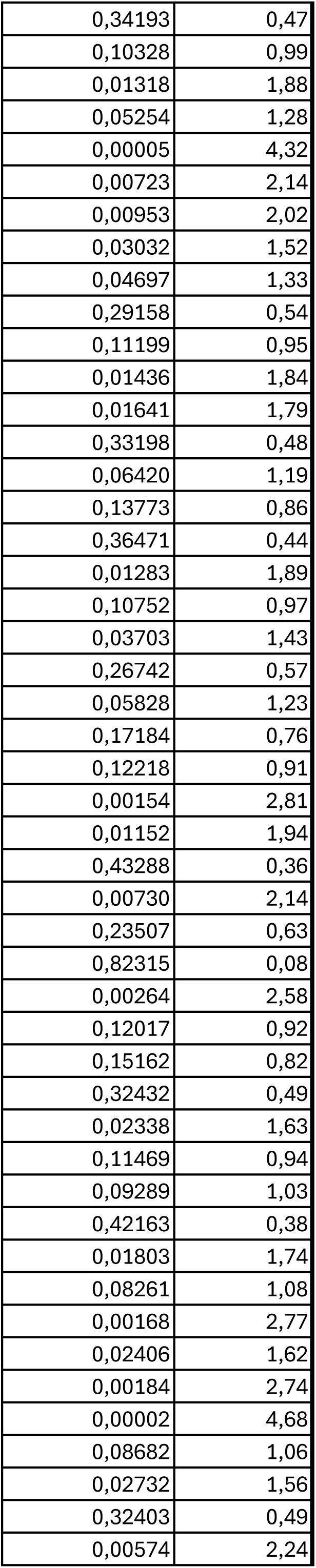
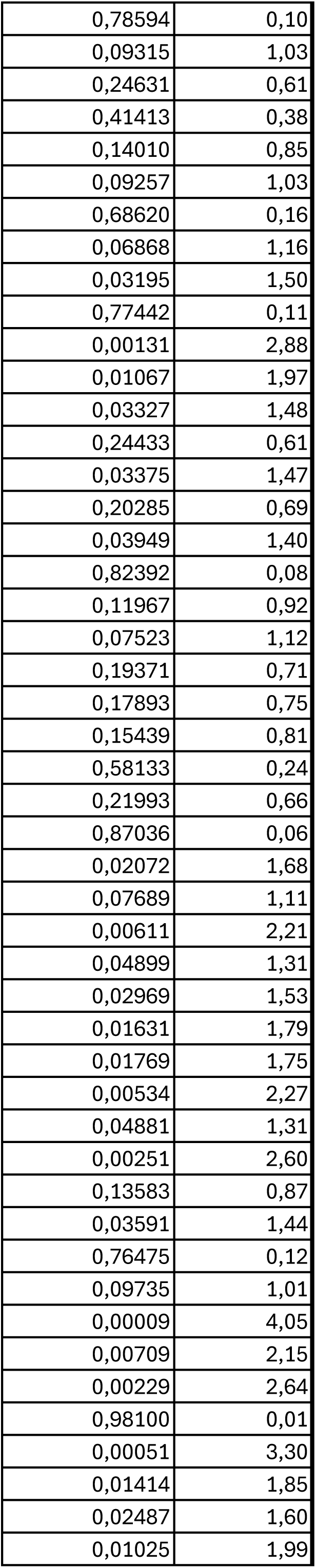
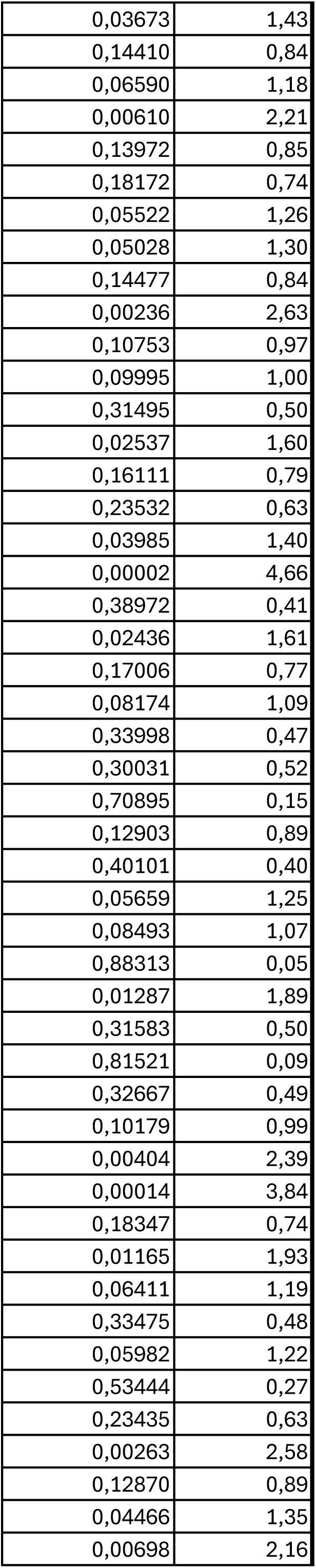
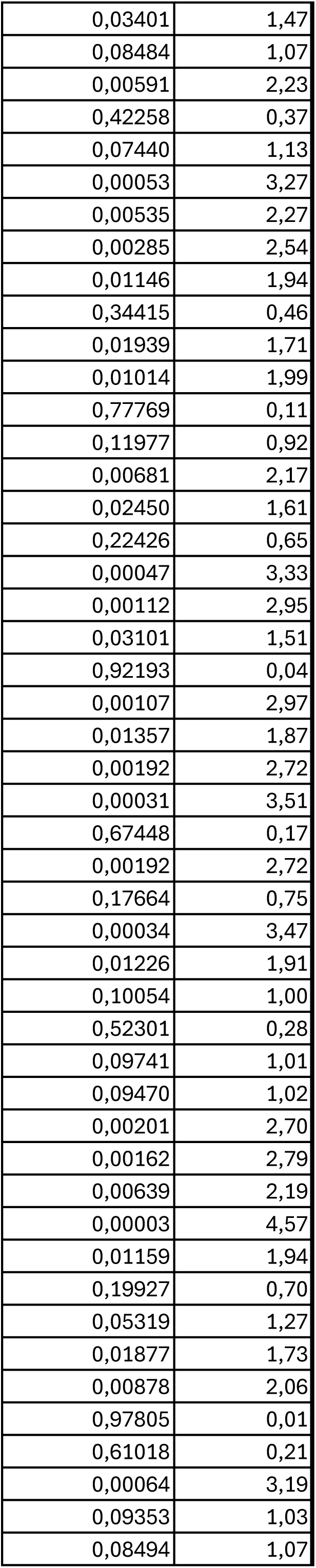
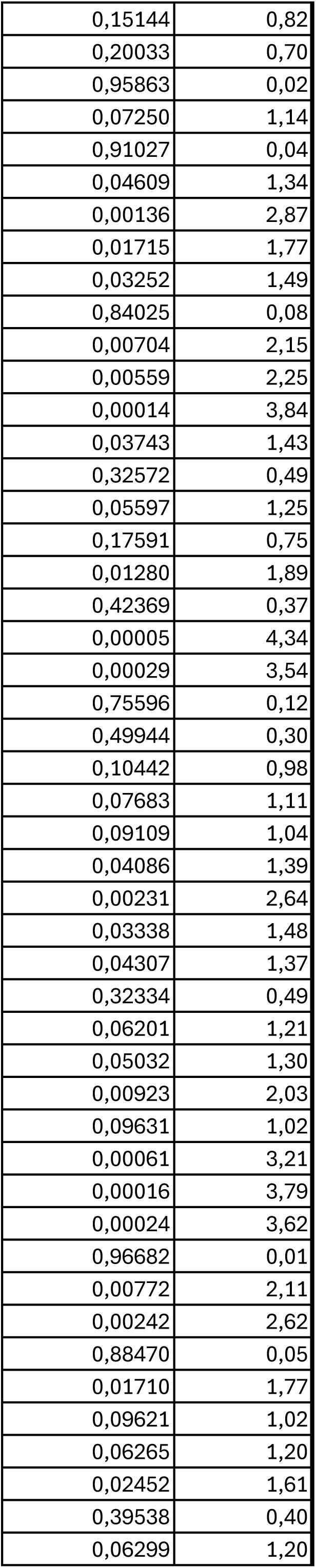
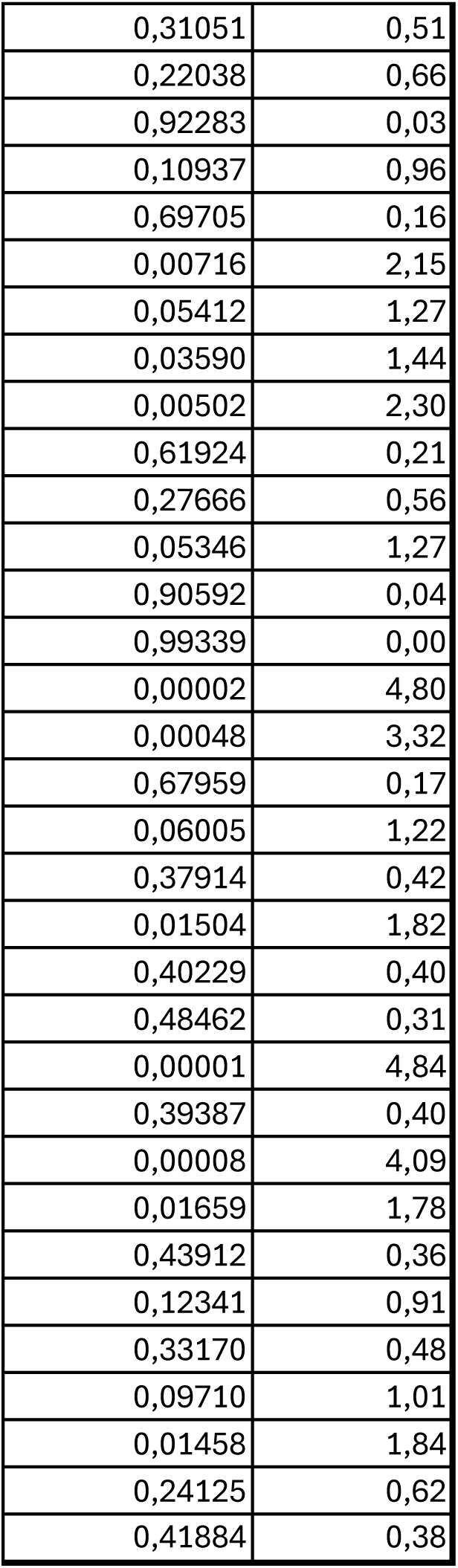
High-throughput MS-based proteomic analyses on paired rBMAds and SCAds.

**Table 2:**
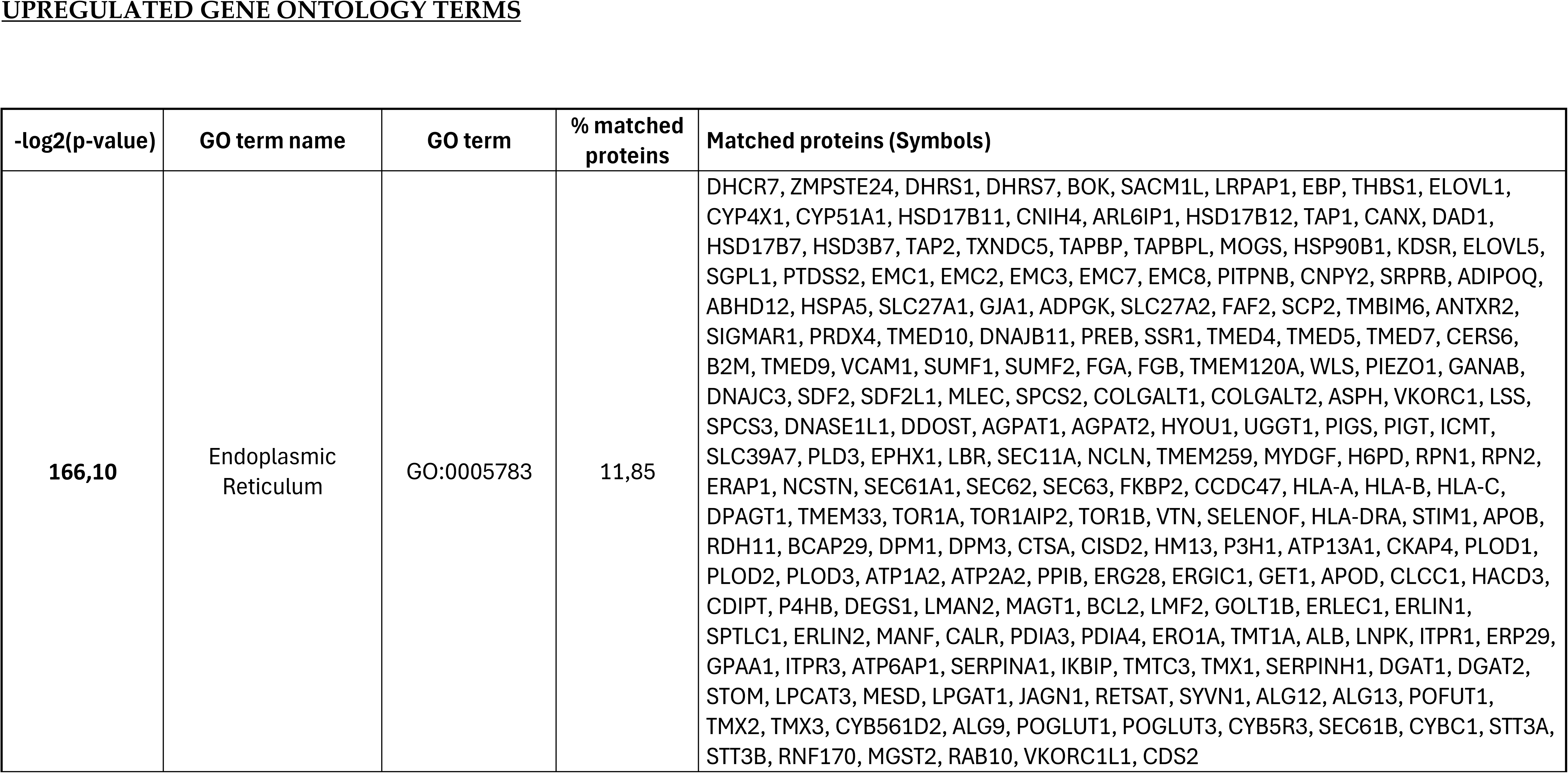
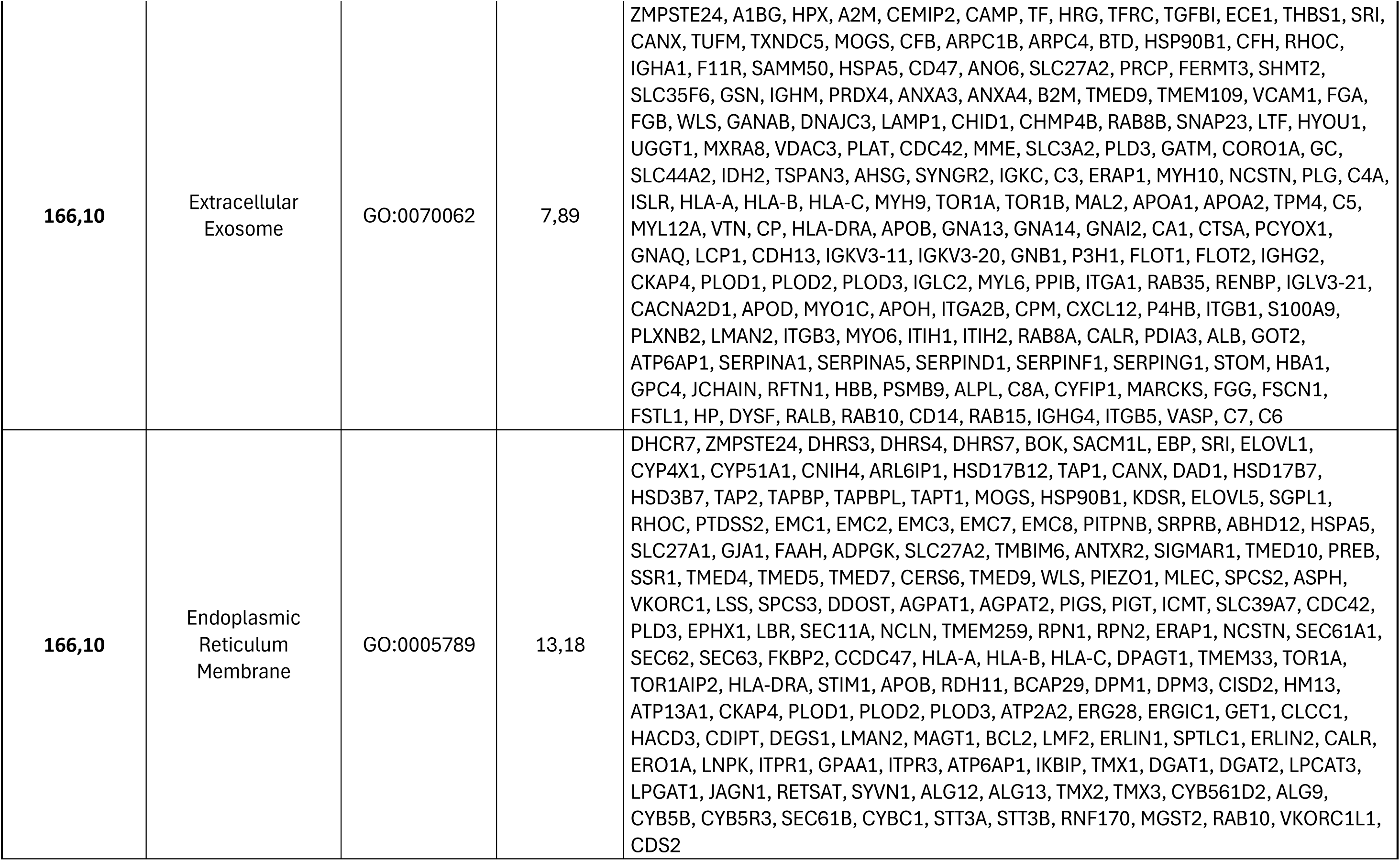
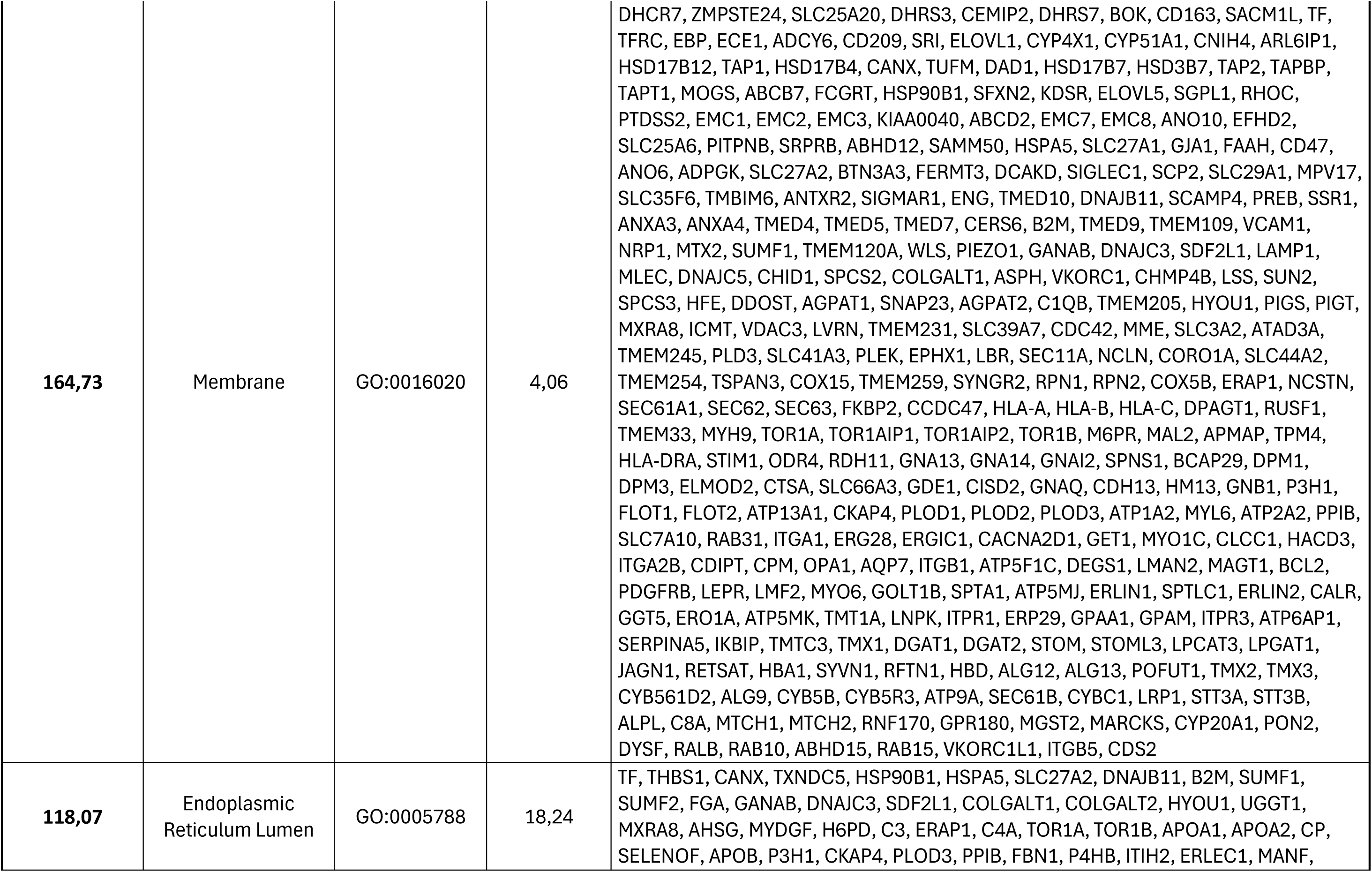
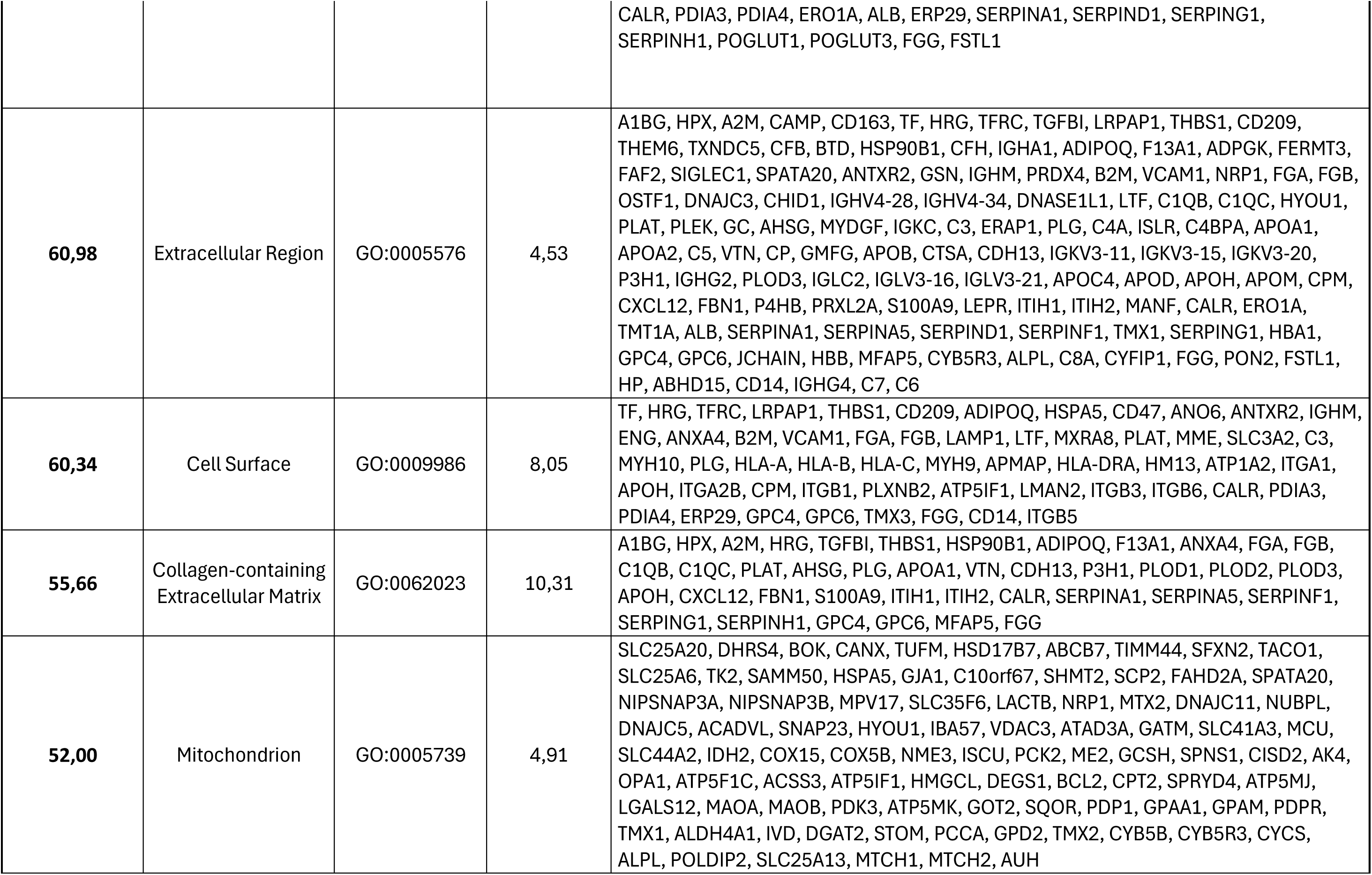
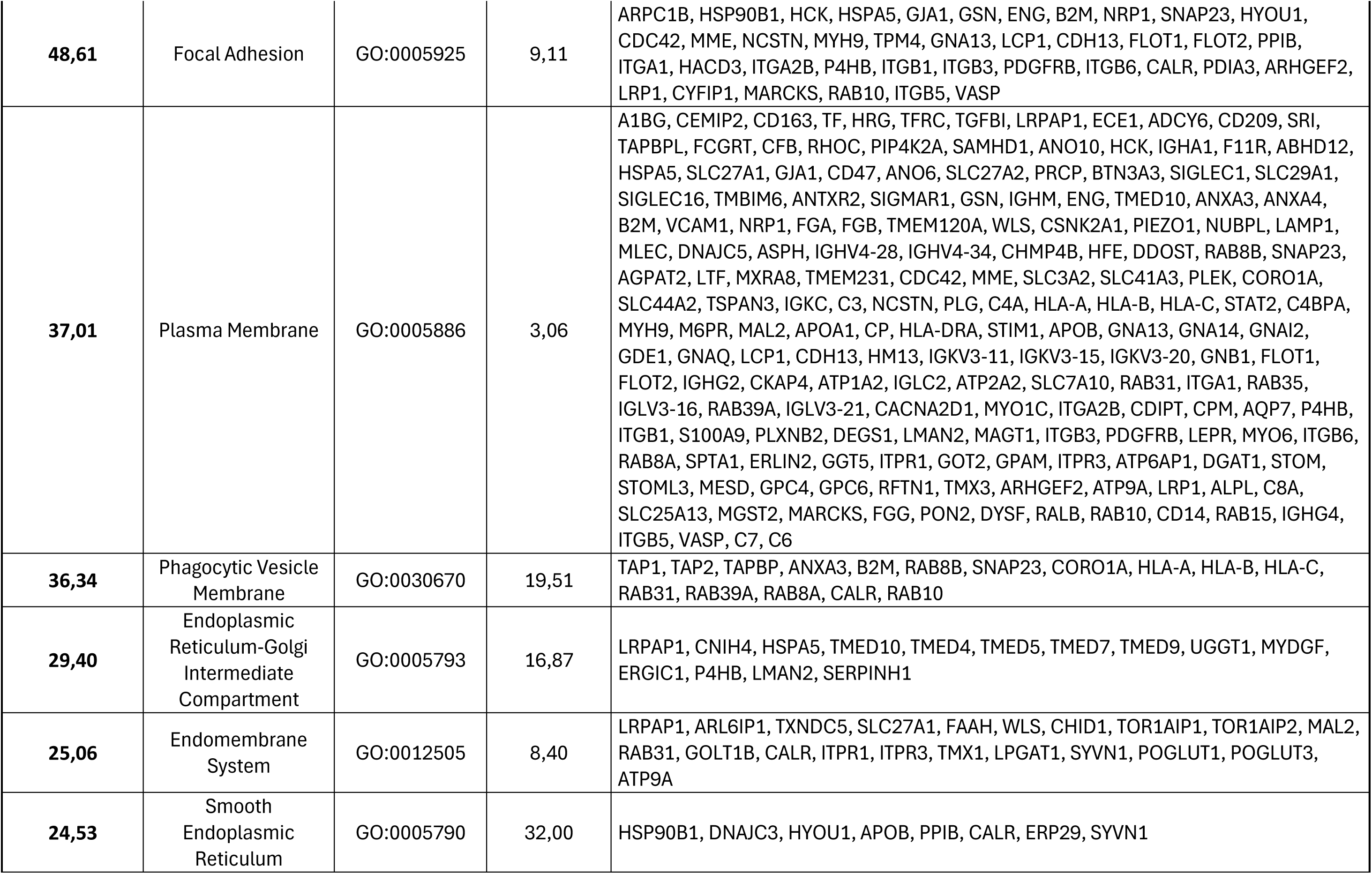
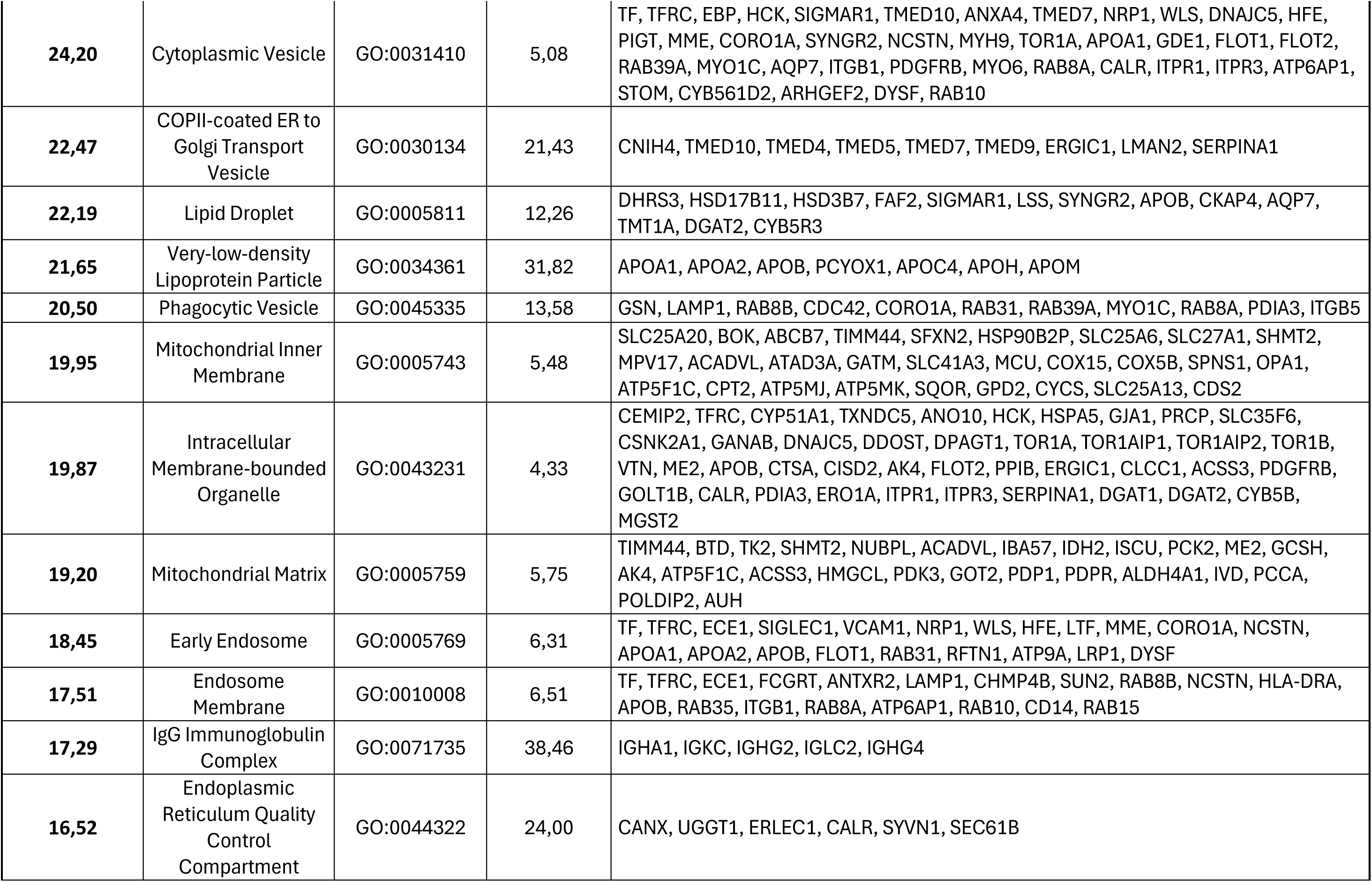
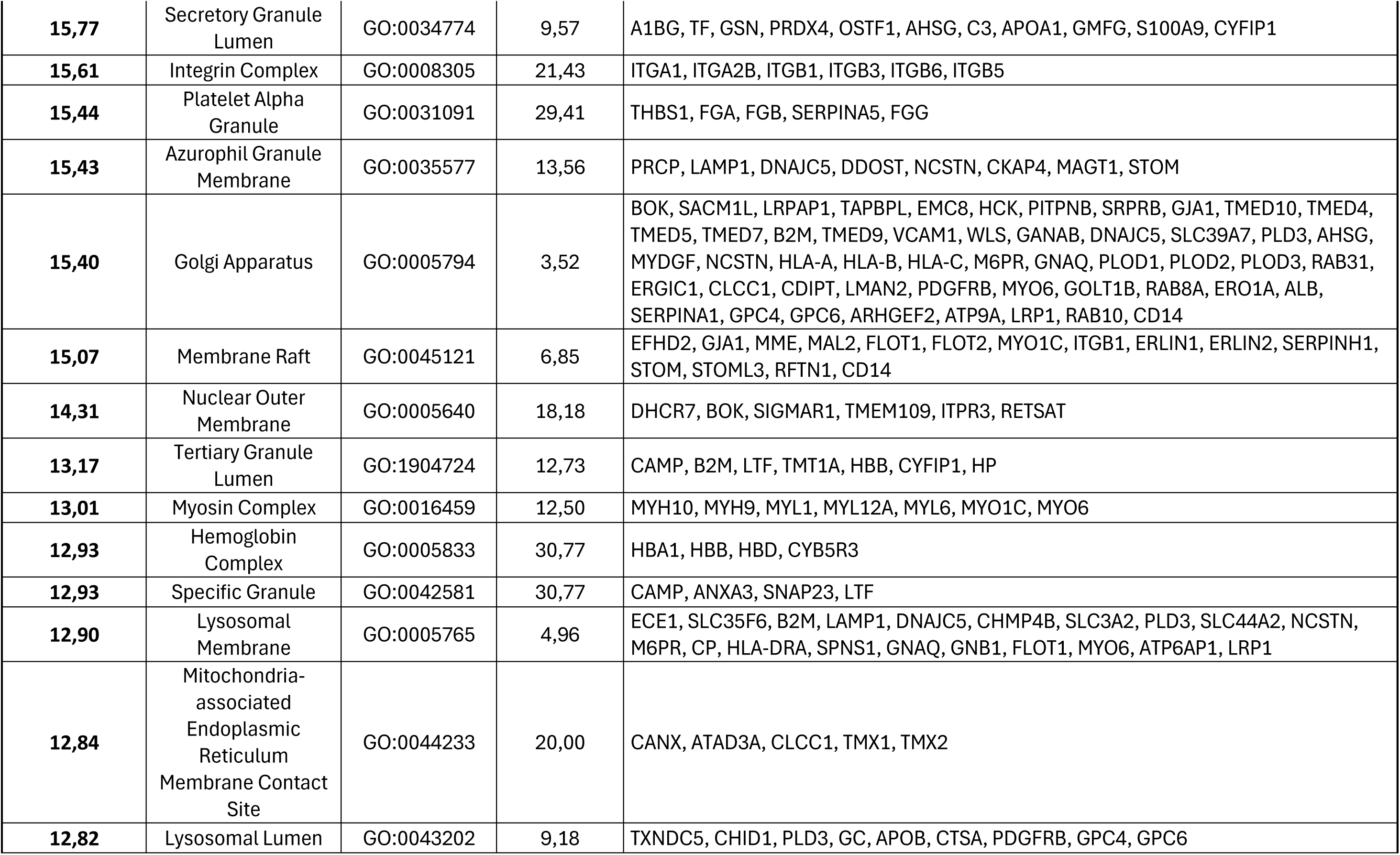
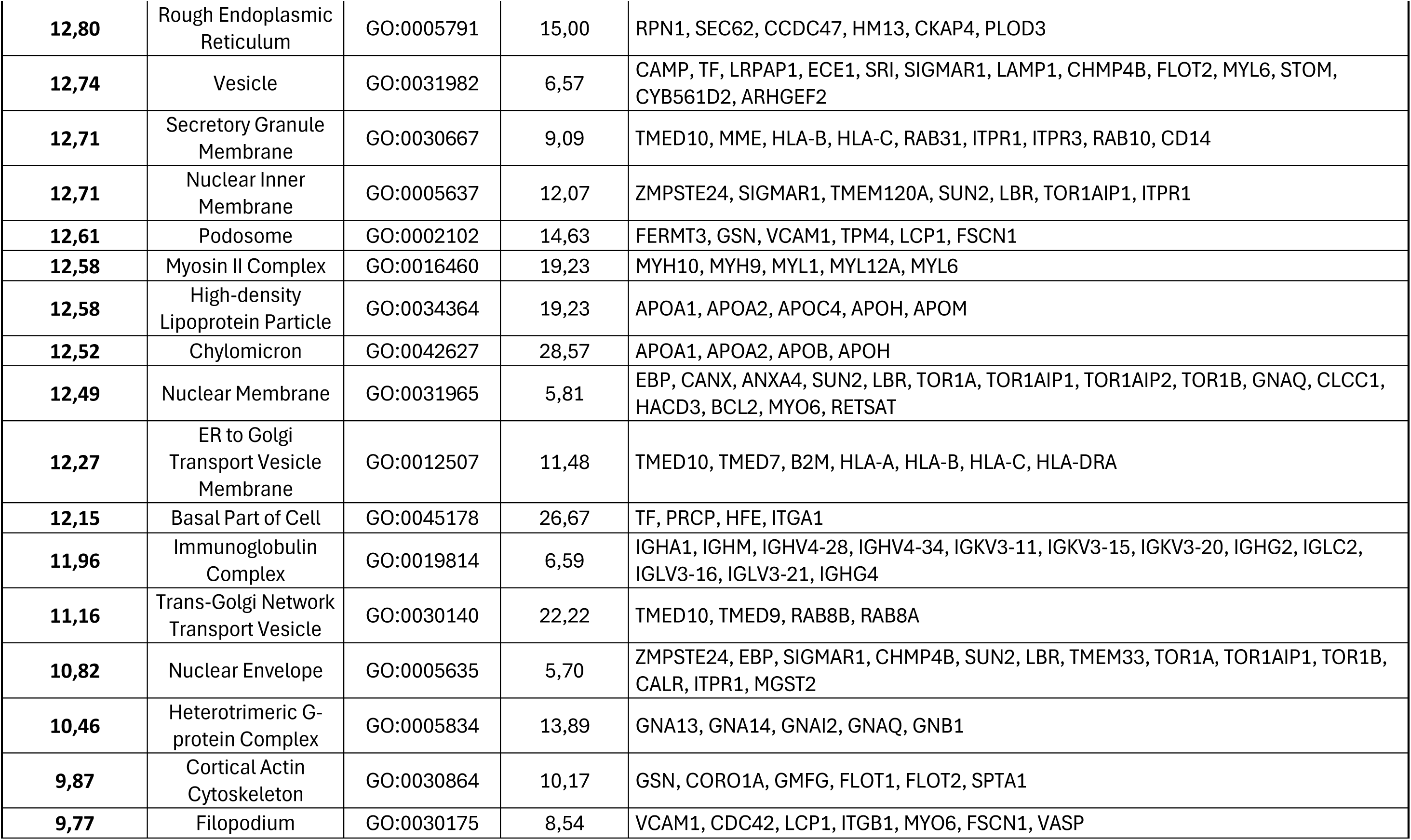
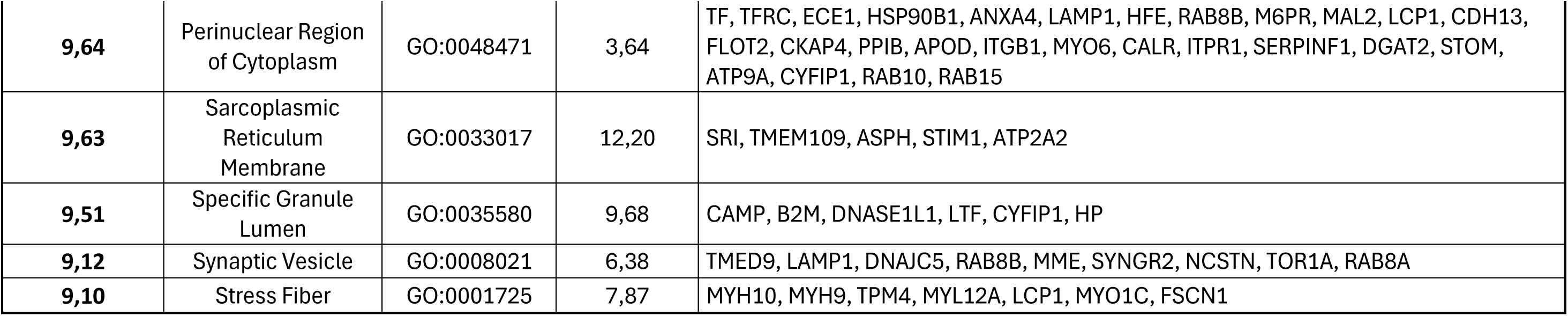
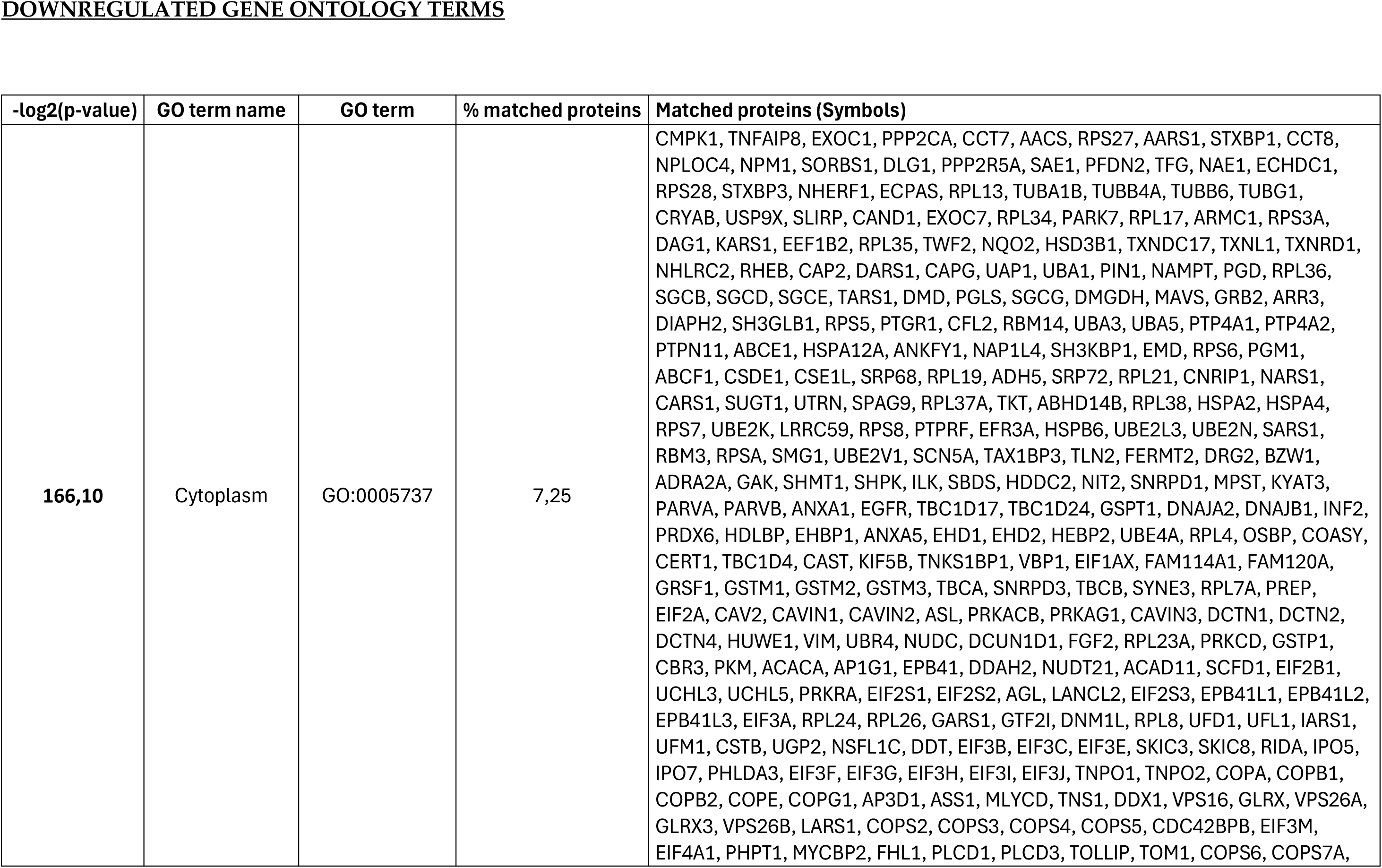
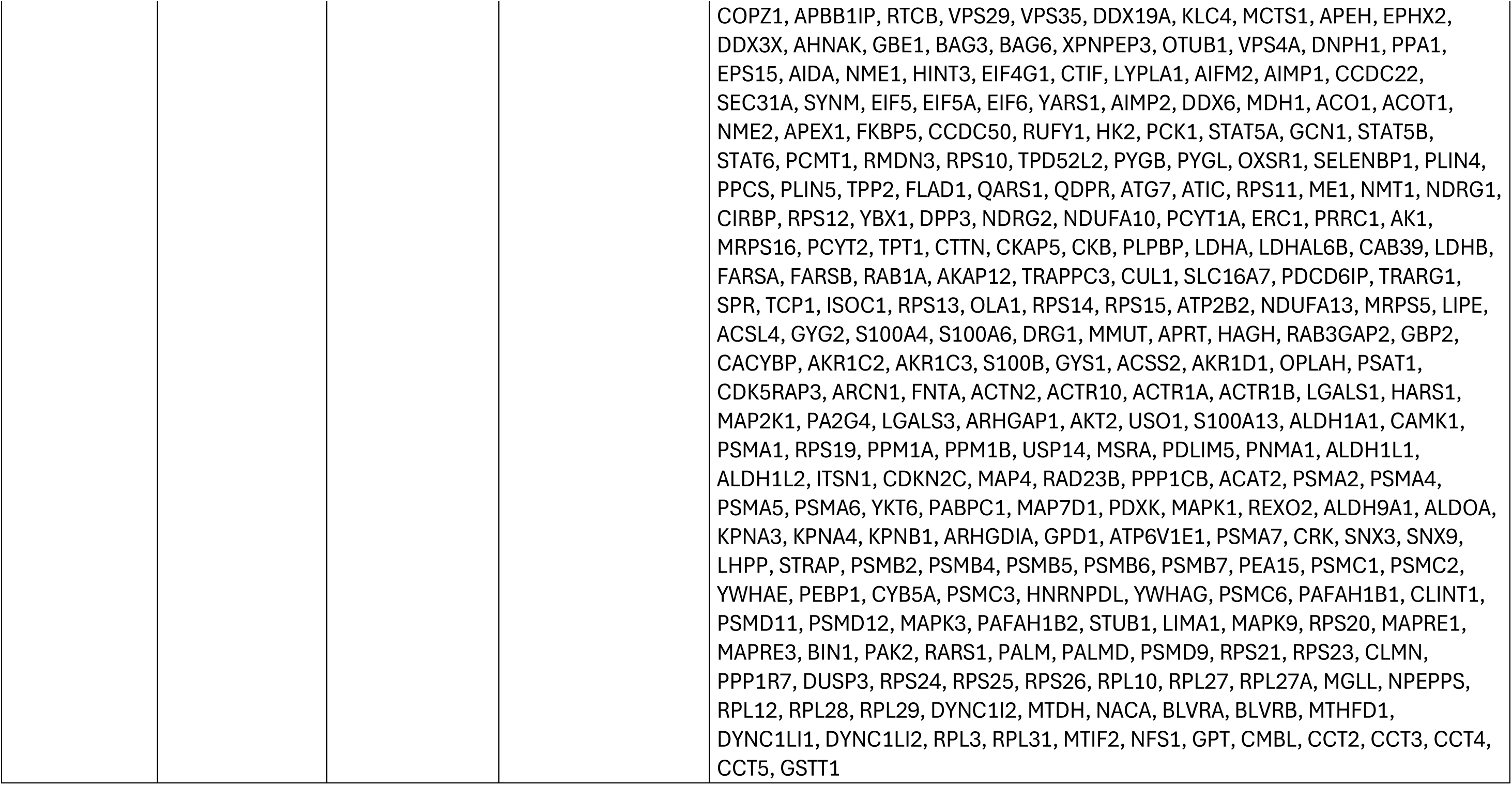
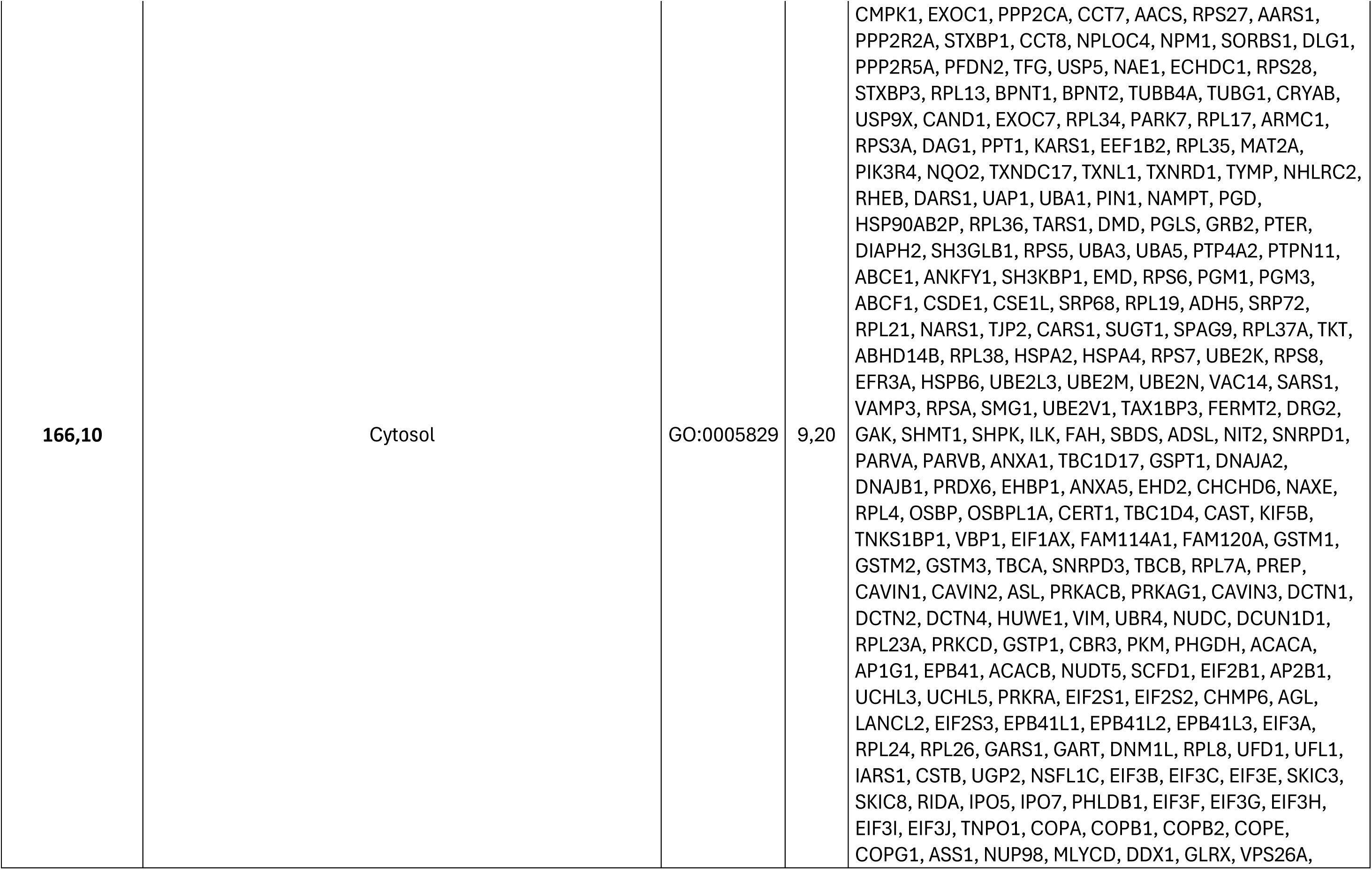
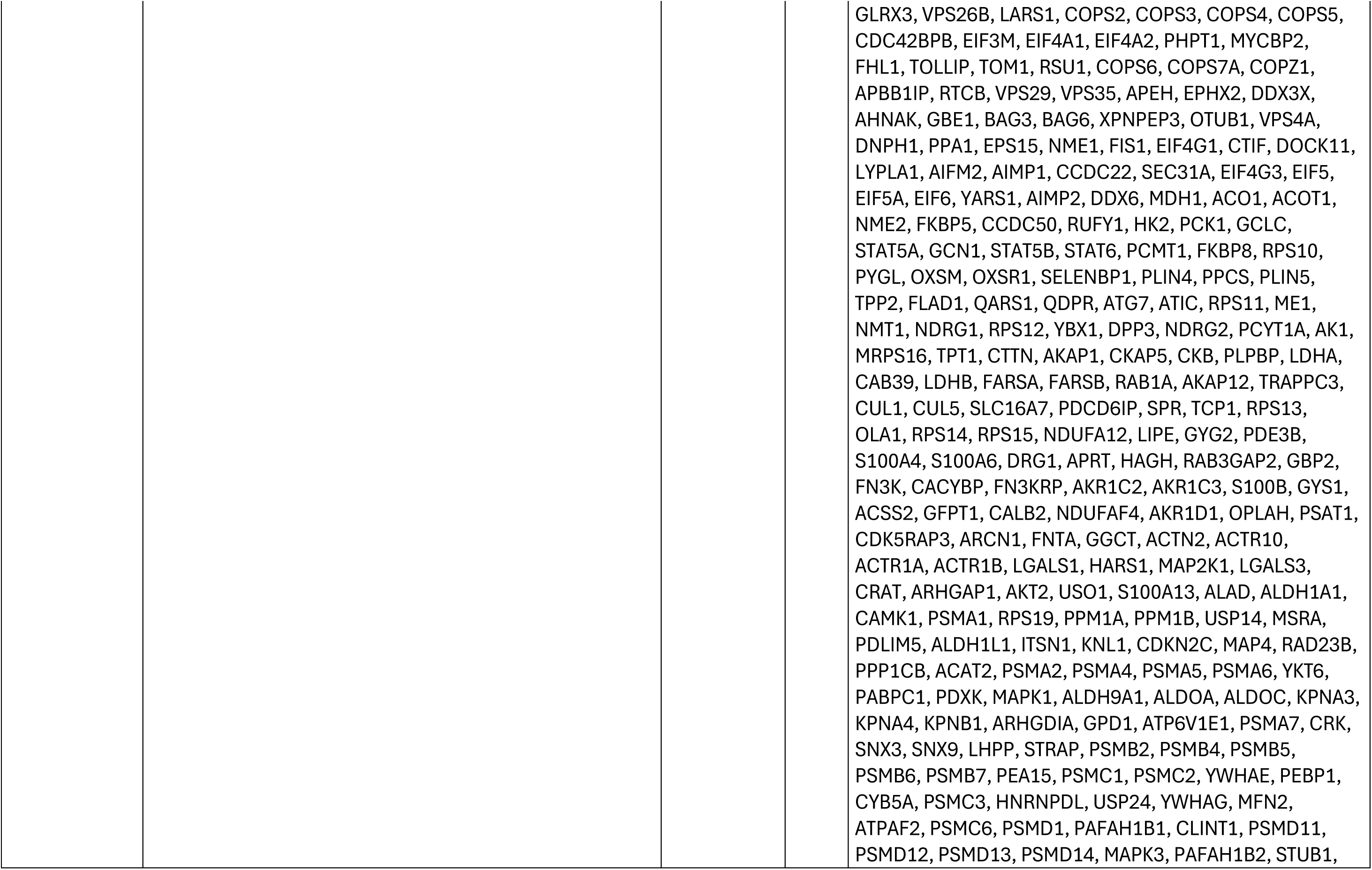
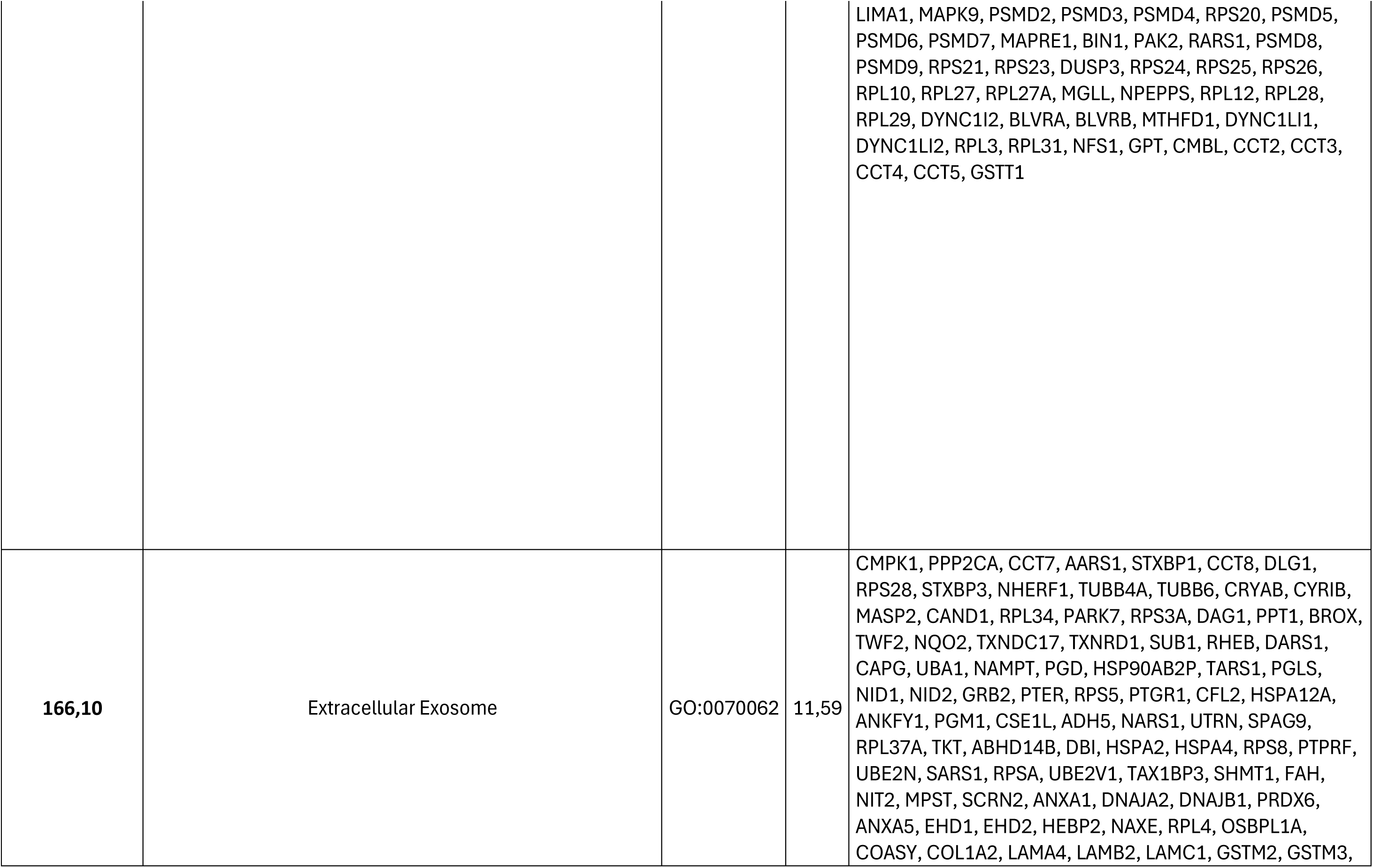
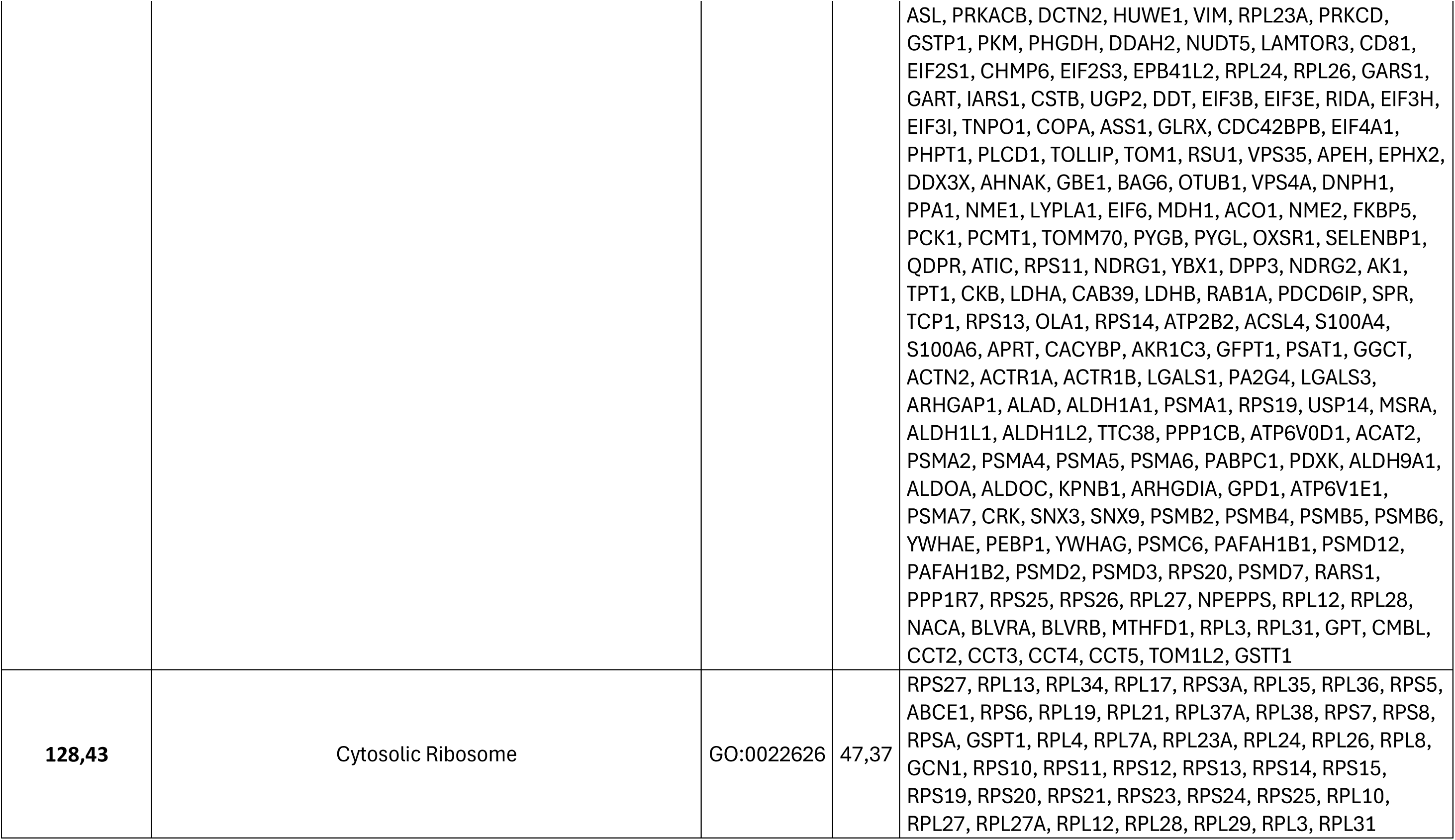
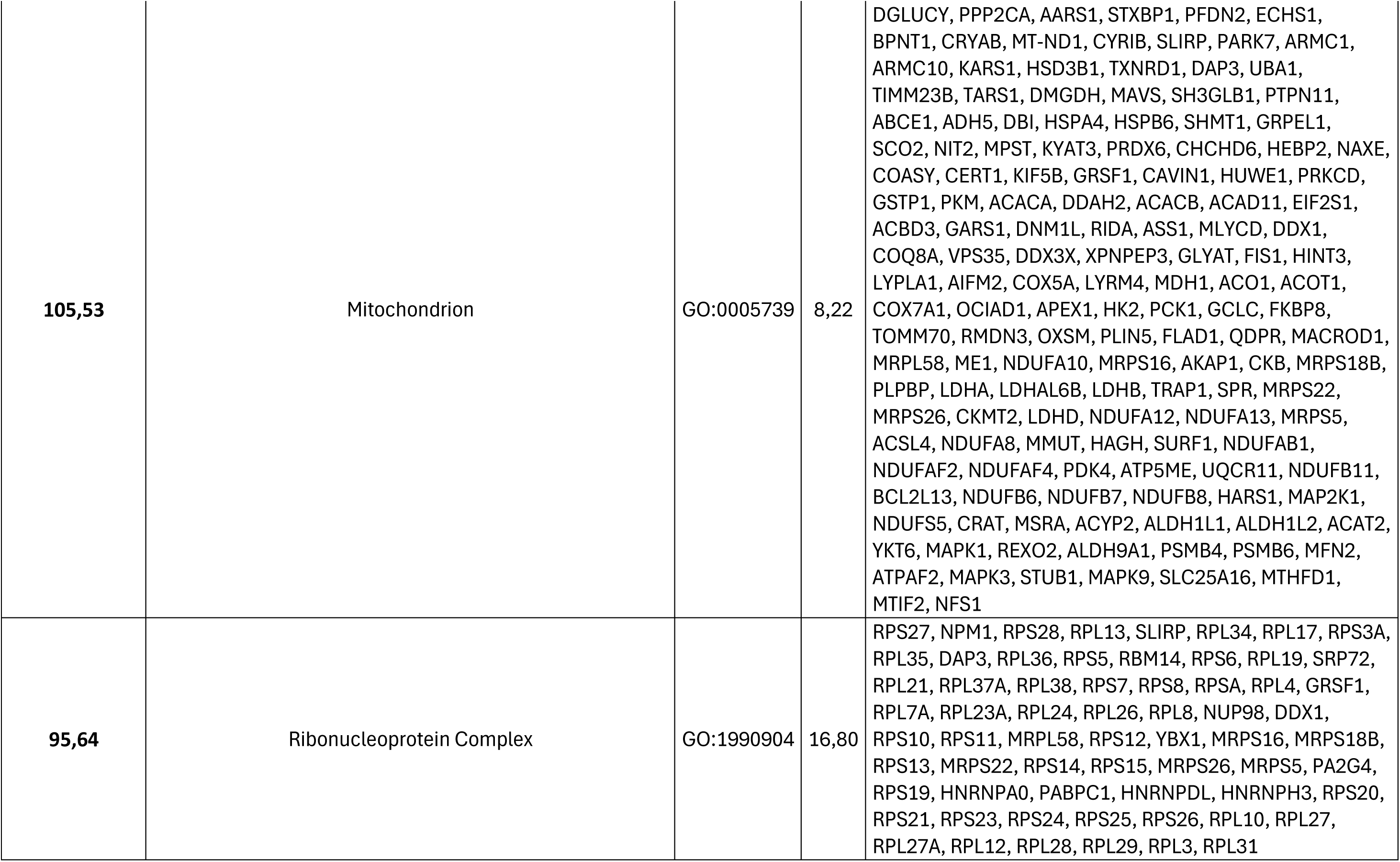
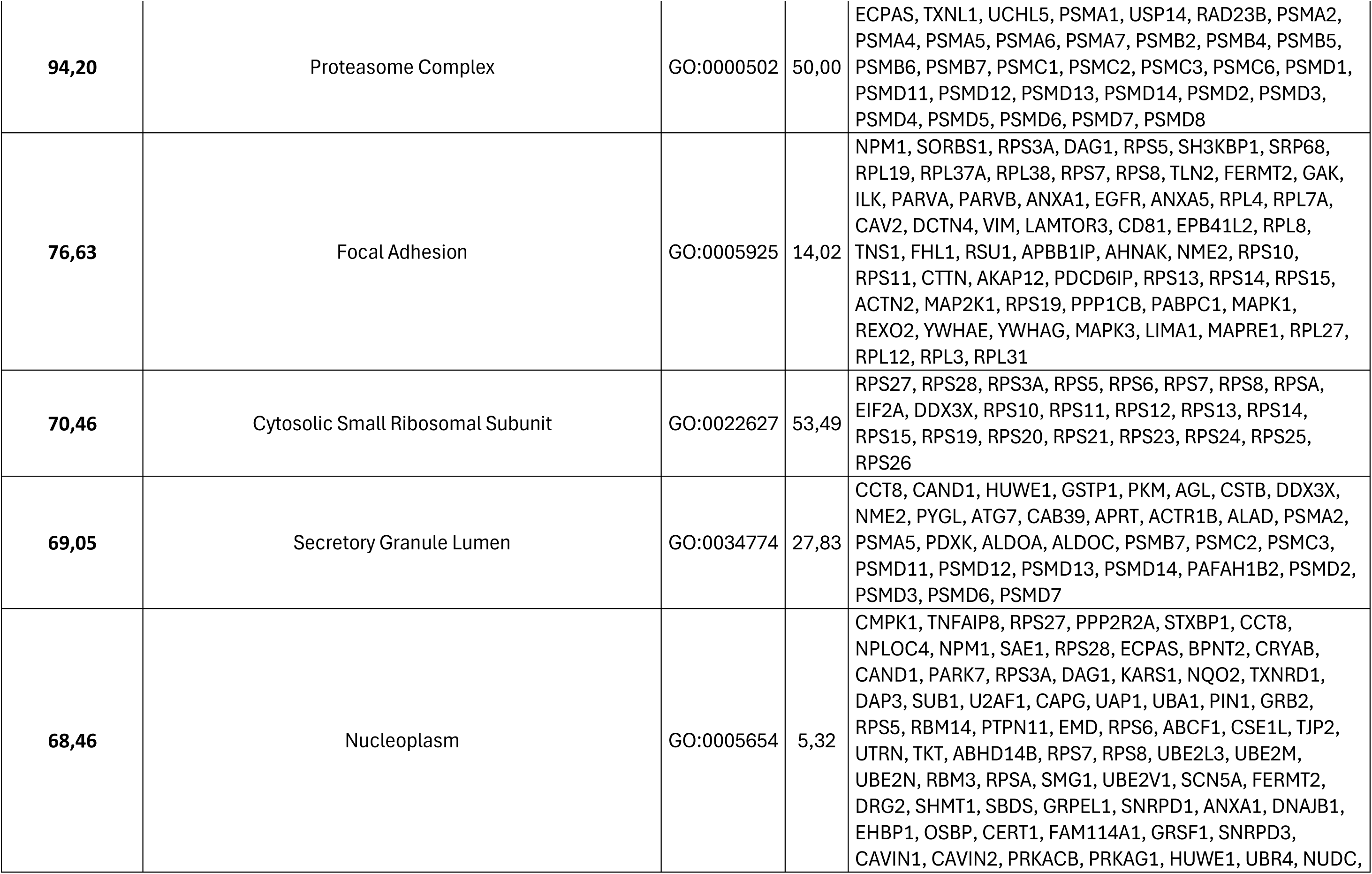
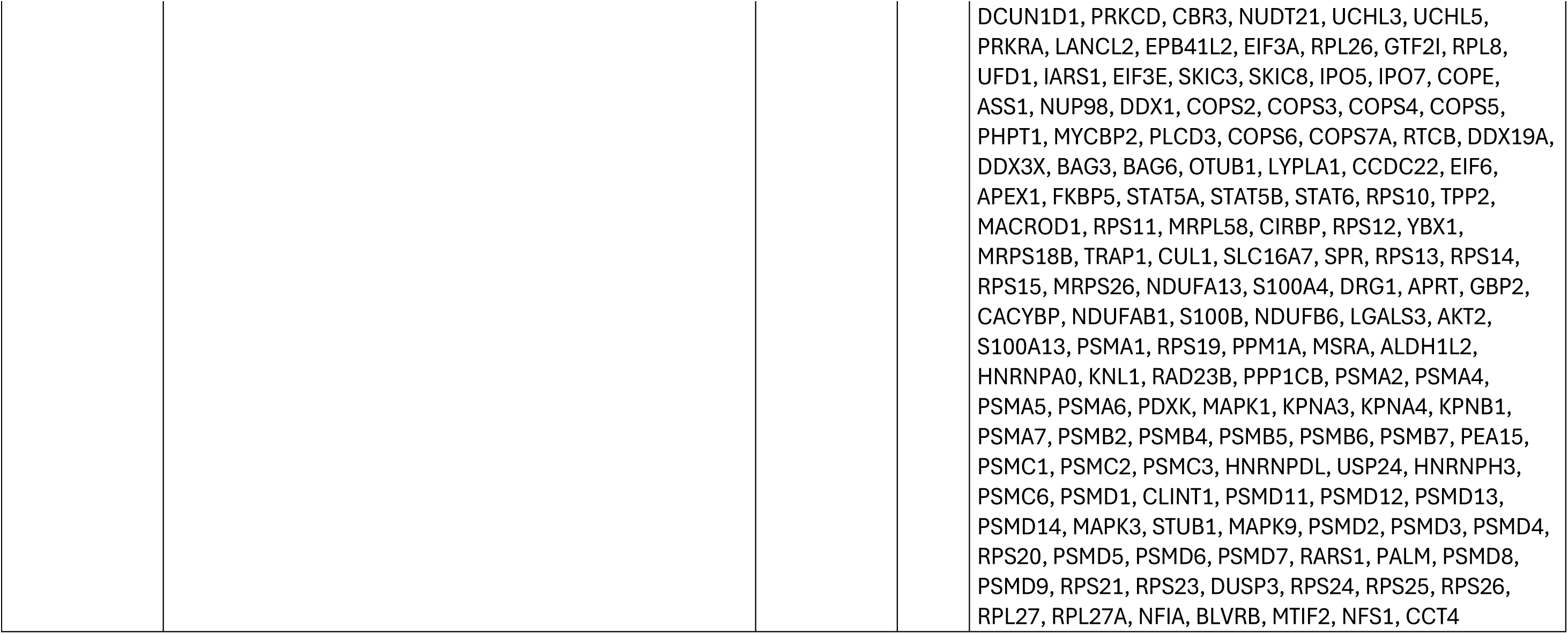
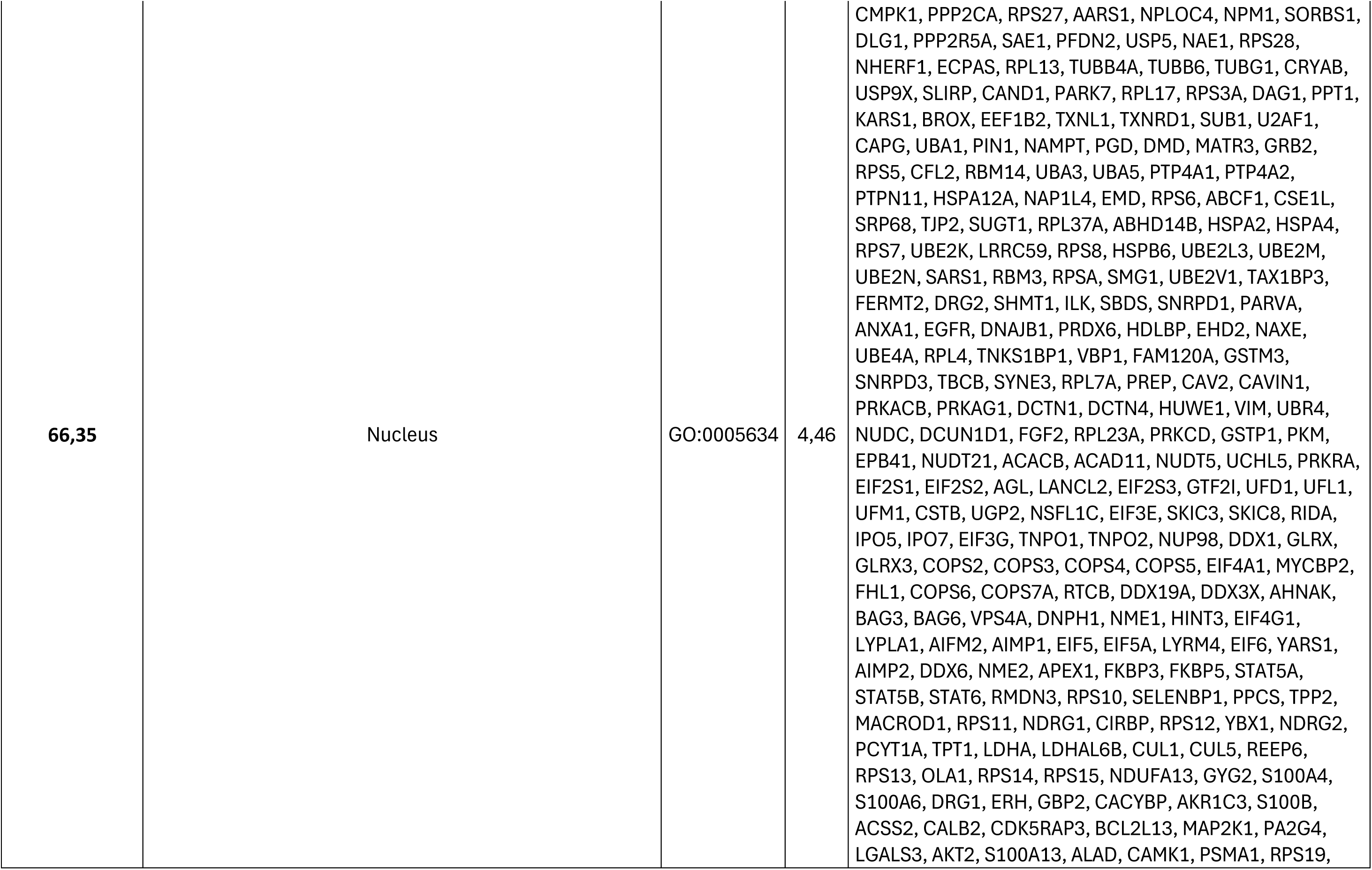
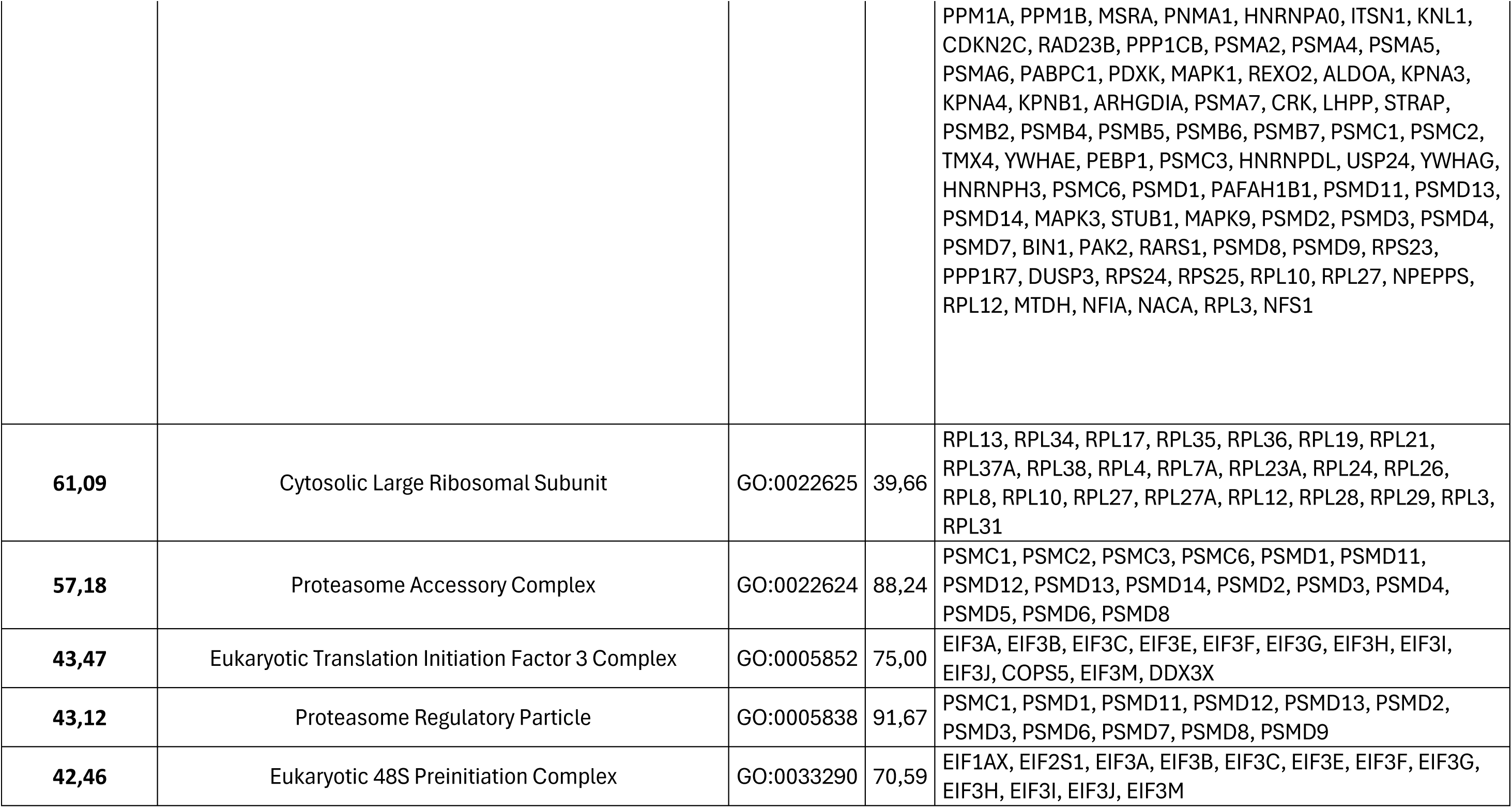
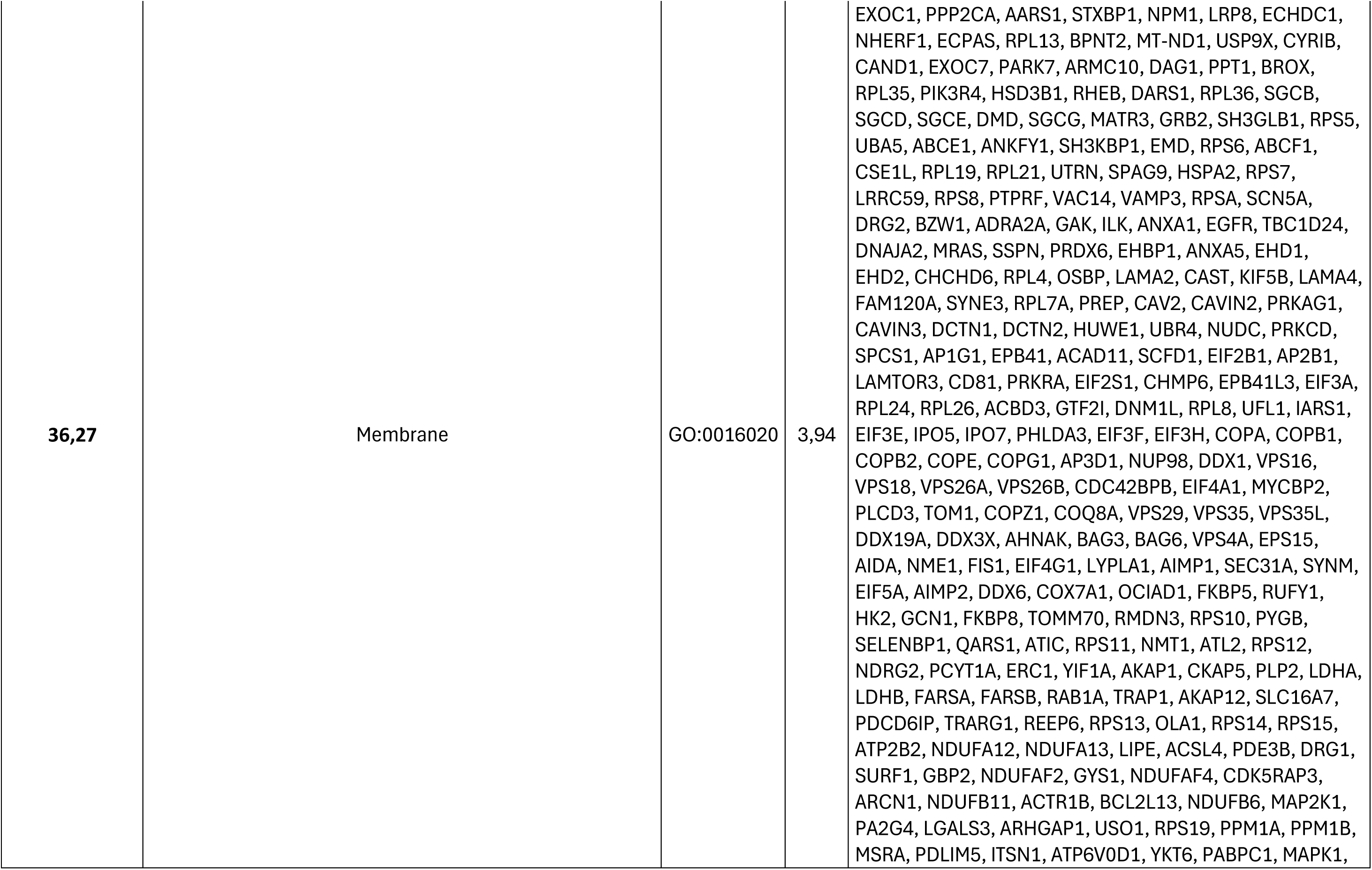
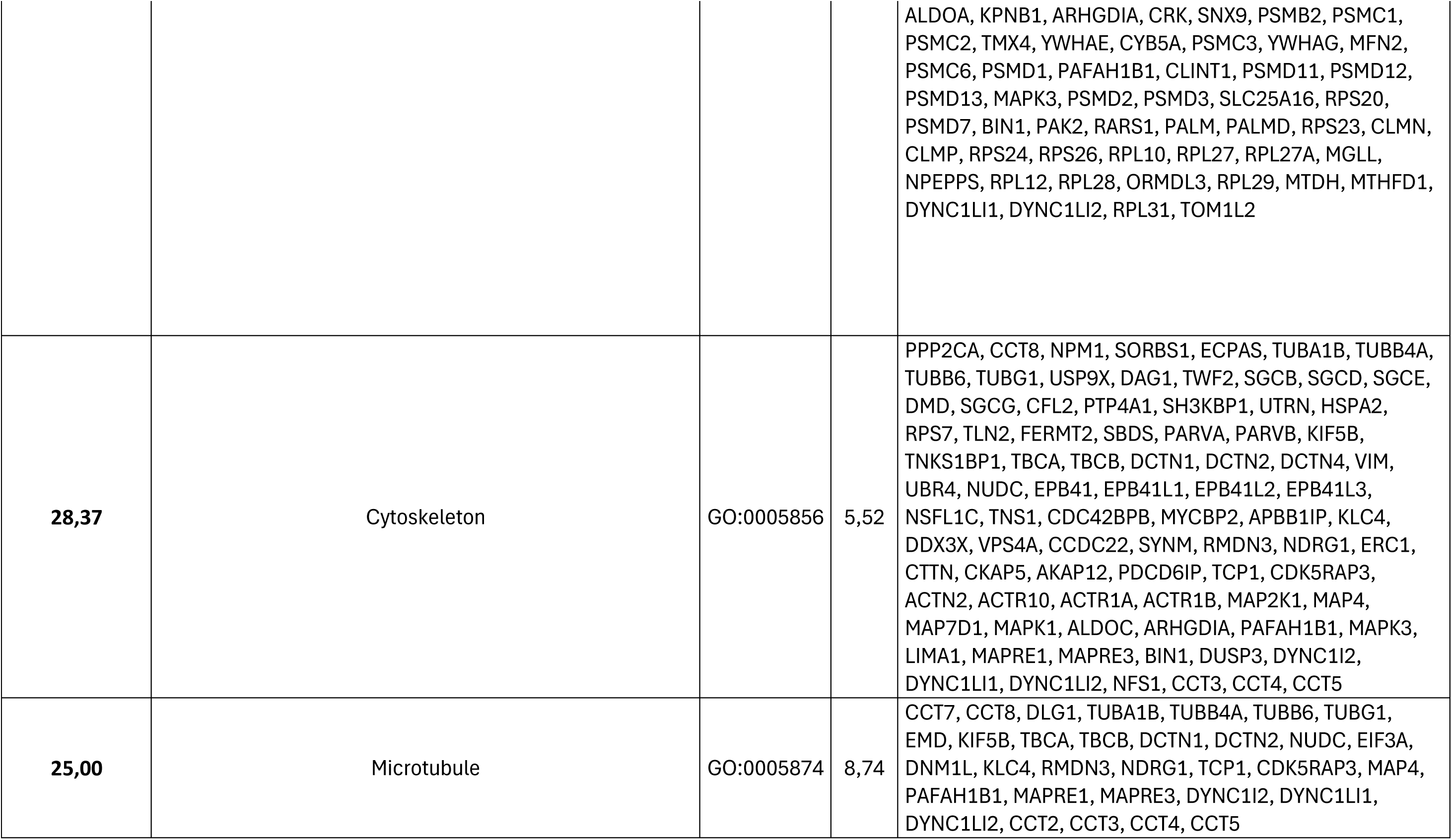
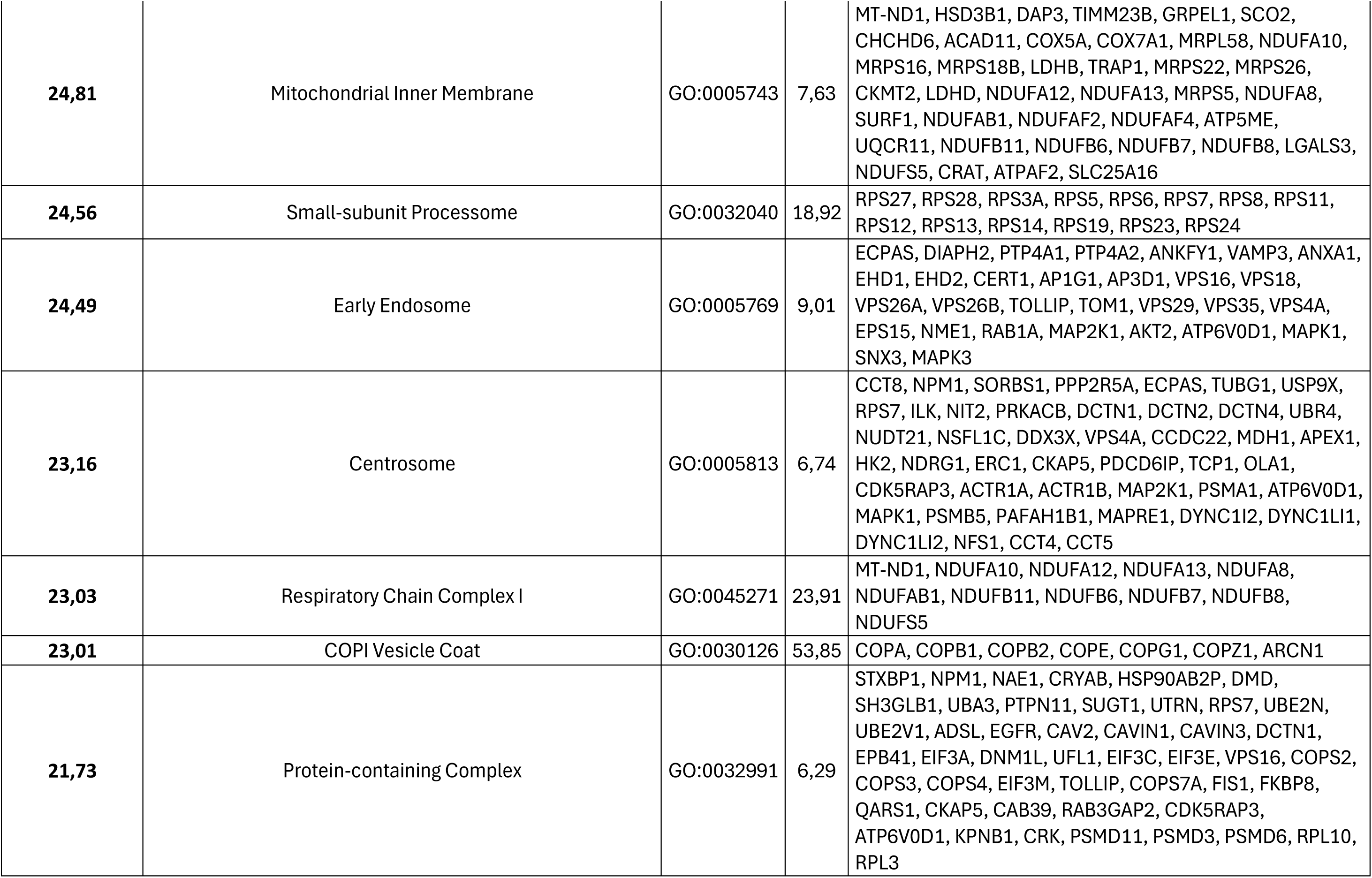
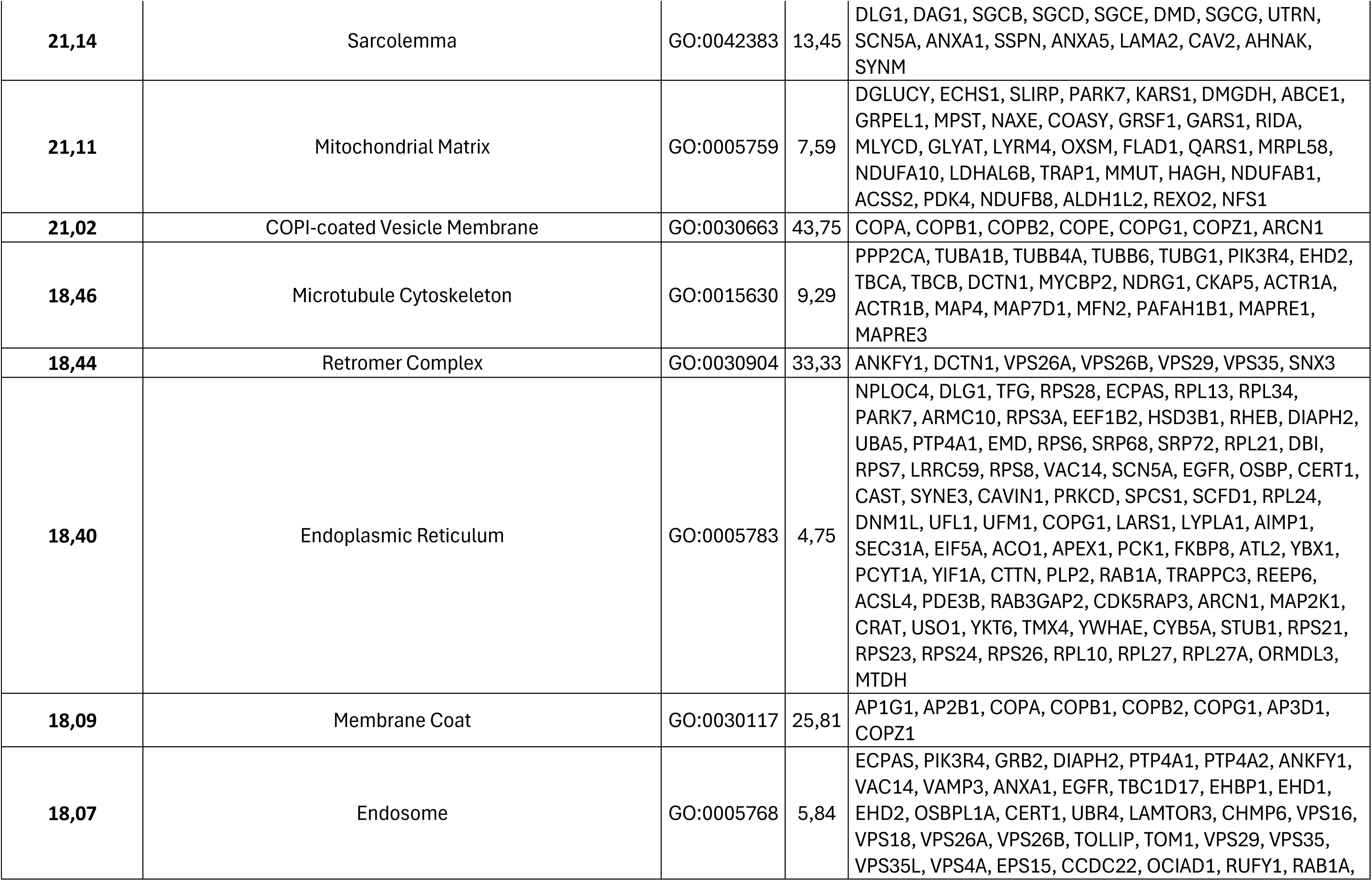
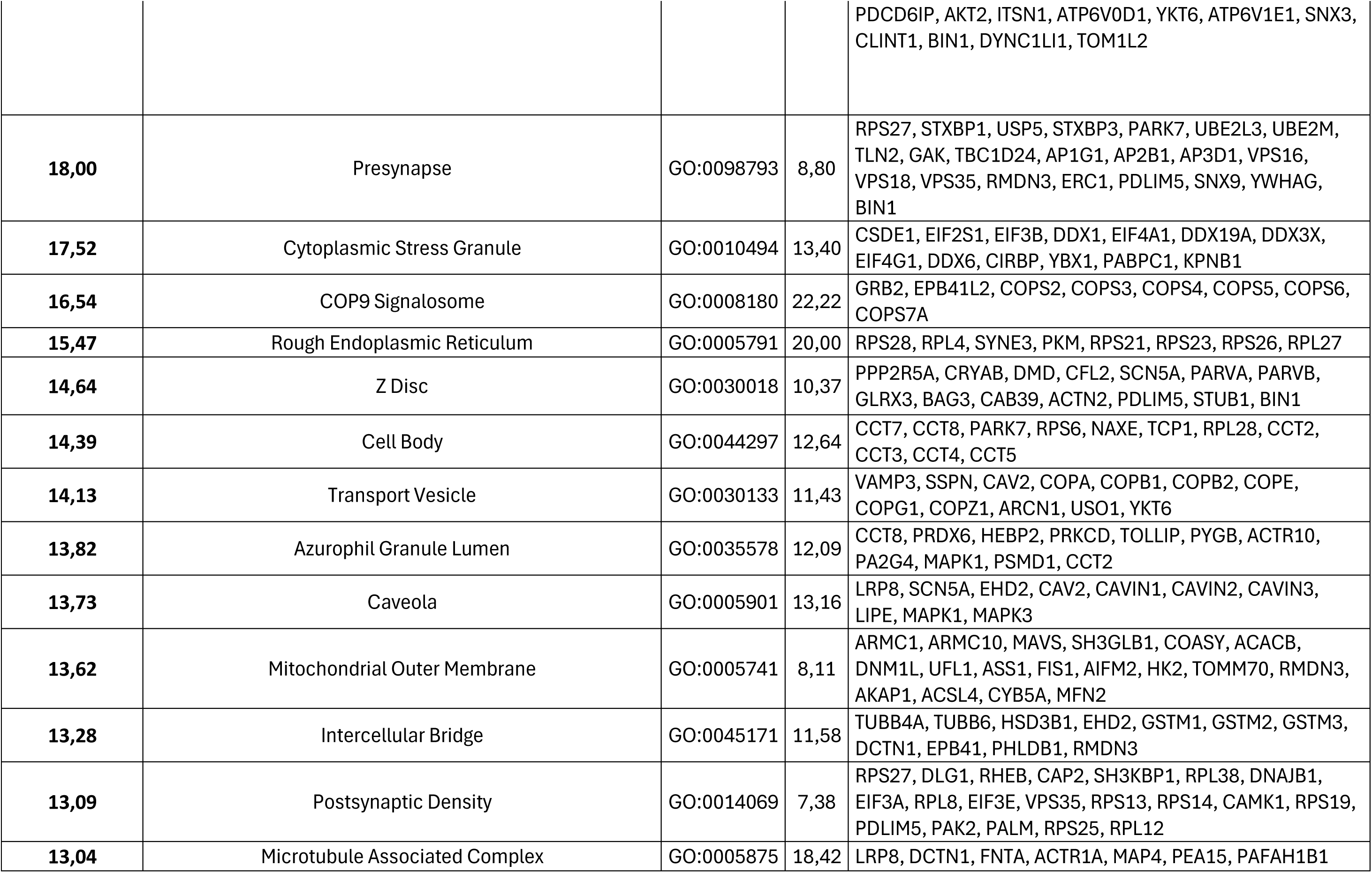
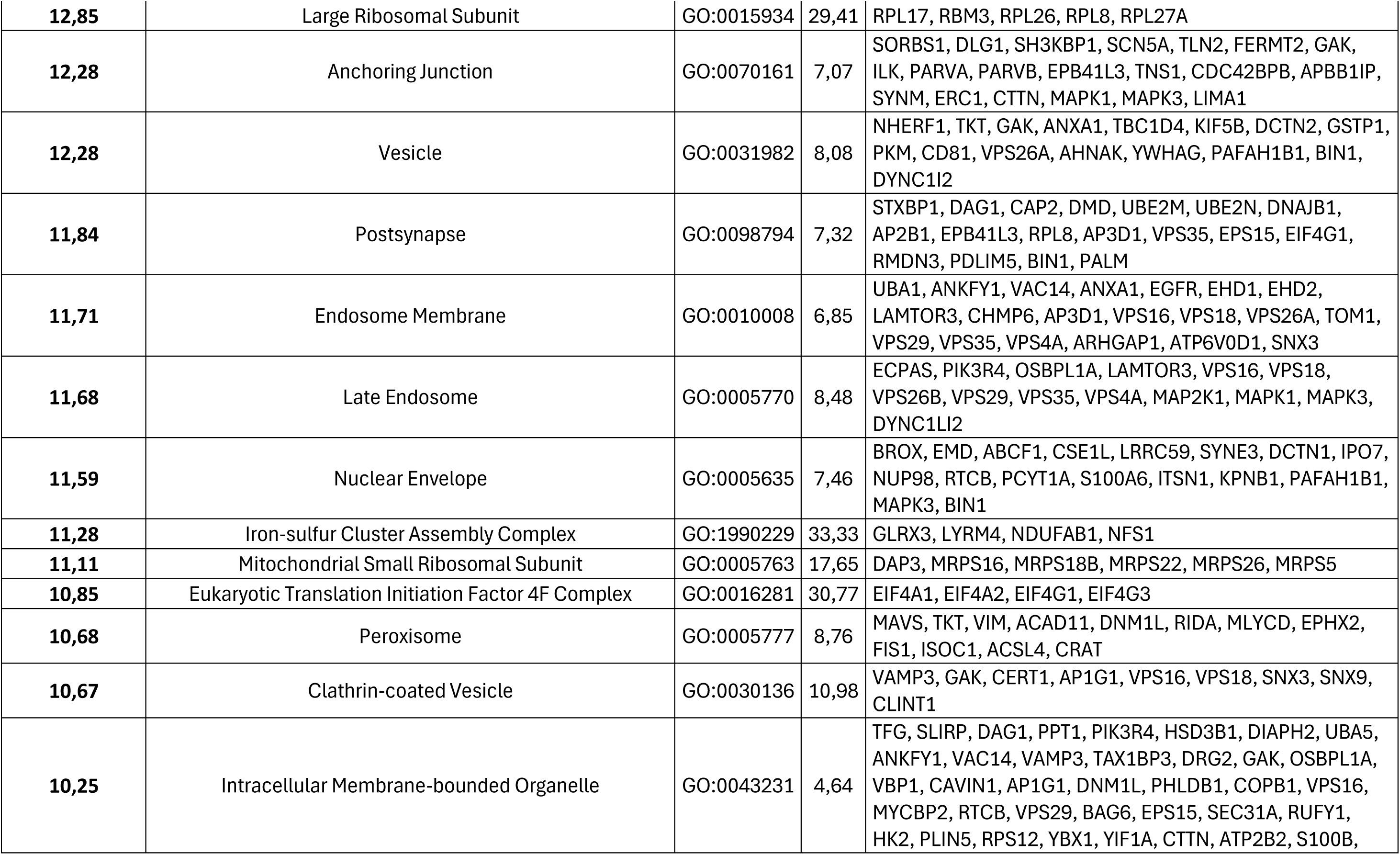
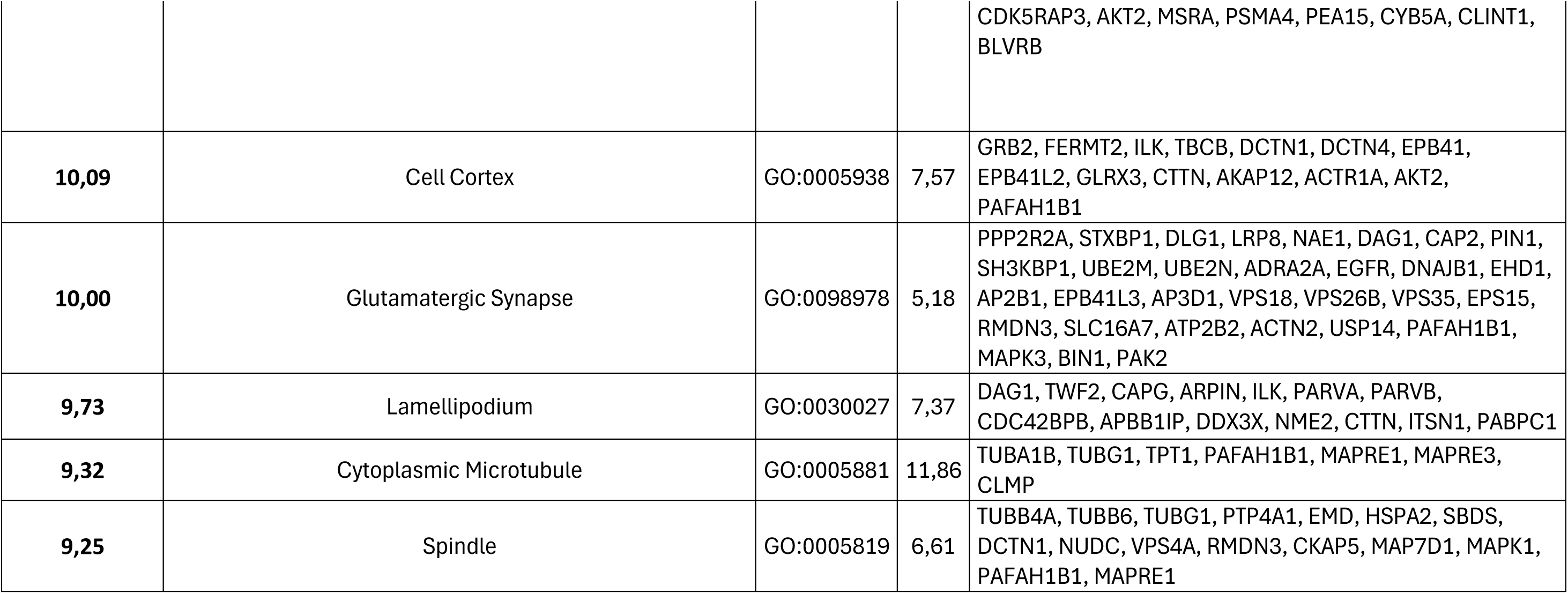
Pathway enrichment analysis of cellular component processes gene ontology terms.

### rBMAds exhibit distinct metabolic and endocrine characteristics and support normal granulo-monocytic hematopoiesis *in vitro*

We next investigated whether these anucleate cells retain metabolic and secretory functions. A primary role of adipocytes is the storage of FAs, either taken up from the circulation or synthesized *de novo* from glucose and their subsequent mobilization through lipolysis^38^. Proteomic analysis of proteins specifically involved in FA metabolism revealed that both adipocyte types express similar levels of FA transporters (most notably CD36 and FABP4) and key enzymes involved in FA oxidation (Figure 4A). In contrast, enzymes responsible for FA esterification into TGs, including AGPAT2 (1-acylglycerol-3-phosphate O-acyltransferase 2) and DGAT1 (Diacylglycerol O-acyltransferase 1), were more abundant in rBMAds than in SCAds (Figure 4A). Functional assays of intracellular FA fate confirmed that FA oxidation rates were comparable between rBMAds and SCAds (Figure 4B), as well as the levels of esterification into mono-, di-, and triglycerides (MG, DG, and TG, respectively) (Figure 4C), despite the elevated expression of esterification enzymes in rBMAds (Figure 4A). Interestingly, rBMAds exhibited a slightly reduced capacity to incorporate glucose into TGs under insulin stimulation (Figure 4D), potentially due to modest reductions in the expression of two key *de novo* lipogenesis enzymes: acetyl-CoA carboxylase-A (ACACA) and fatty acid synthase (FASN) (Figure 4A). Thus, despite lacking a nucleus, rBMAds remain metabolically active, displaying capacities for FA and glucose utilization for TG storage, as well as insulin responsiveness, comparable to those of SCAds. These findings contrast with imaging studies in humans using positron emission tomography, which have shown increased uptake of FAs and glucose in BMAT^21–23^. These discrepancies likely stem from the imaging technique capturing substrate uptake at tissue level, so the higher rBMAT signal may reflect metabolic activity in surrounding hematopoietic cells rather than rBMAds themselves. We next assessed lipolytic activity under basal conditions and upon stimulation with isoprenaline and forskolin, two canonical activators of lipolysis. In both rBMAds and rBMAT, glycerol release was low at baseline and remained unaltered upon stimulation, in contrast to the robust response observed in SCAds (Figure 4E) and SCAT (Figure S4A). This marked impairment of lipolysis in rBMAds is consistent with our proteomic results showing their low expression of HSL and MGLL, the terminal enzymes in the lipolytic cascade, whereas ATGL, the initiating enzyme, and its coactivator CGI-58 (α-β hydrolase domain-containing 5) were more abundant in rBMAds than SCAds (Figure 4F). These findings indicate that, like cBMAds^17^, rBMAds are devoid of classical lipolytic activity, contributing to explain the well-documented preservation of BMAT during caloric restriction in contrast to the active lipolysis observed in white AT^1,2^. Our findings in isolated rBMAds underscore key human–rodent differences. In rodents, rBMAds respond to β-adrenergic stimuli, although their glycerol release is very modest compared to SCAds^15,19^ and rBMAds decrease in size and number following cold exposure in control mice whereas this response is lost in mice with BMAd-specific deletion of ATGL^19^. Notably, our proteomic data show higher ATGL levels in human rBMAds than in SCAds, suggesting a possible role for ATGL in non-classical or partial lipolysis in human rBMAds. Consistently, human biopsies from acute myeloid and lymphoblastic leukemia, both arising in hematopoietic regions, show reduced rBMAd size and number, indicating that rBMAds also undergo dynamic remodeling in humans through an unknown mechanism^39^.

**Figure 4.**
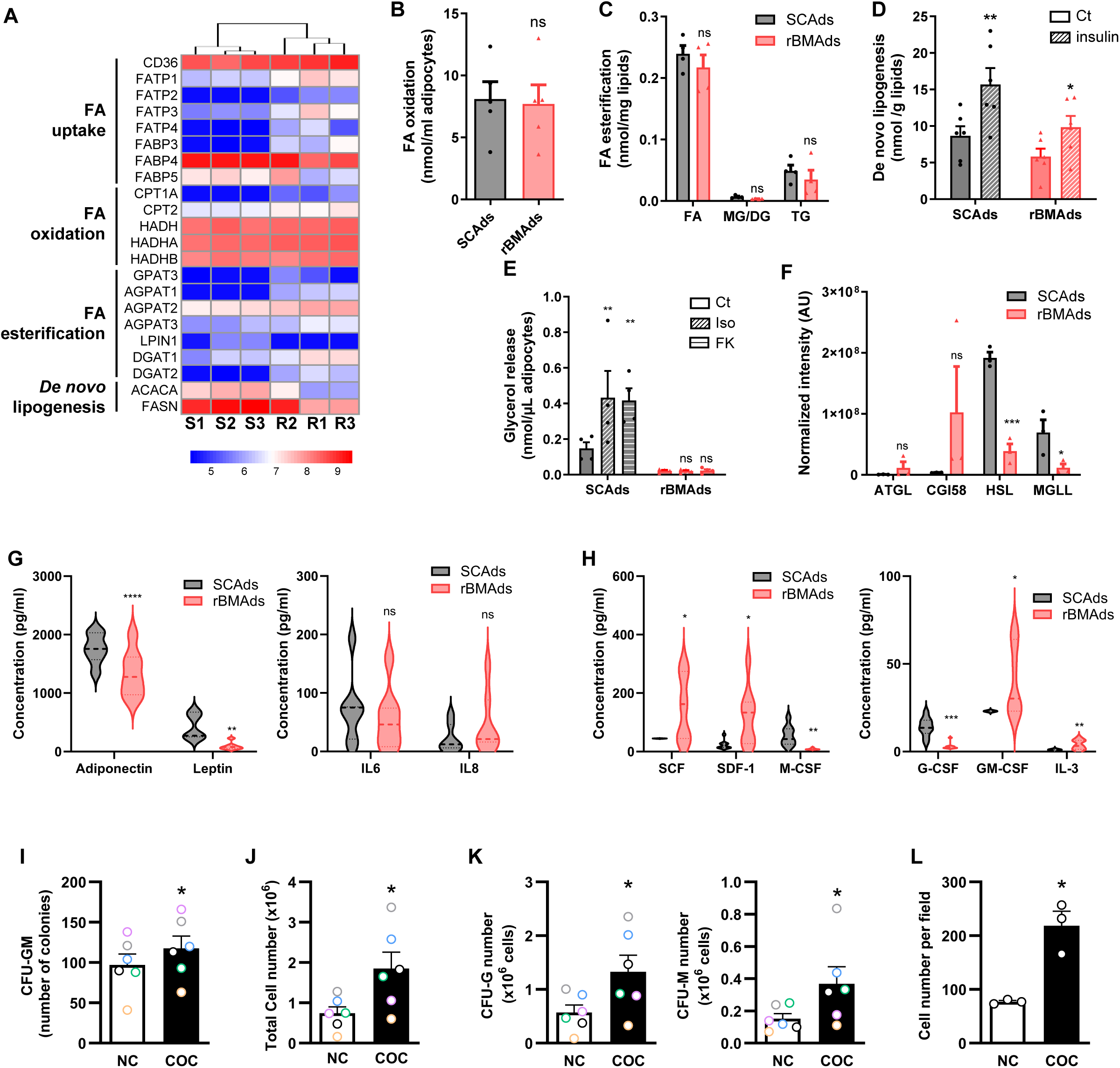
rBMAds exhibit distinct metabolic and endocrine characteristics and support normal granulo-monocytic hematopoiesis *in vitro*. **(A)** Heatmap showing expression of protein involved in FA metabolism in rBMAds (R) and SCAds (S). **(B-E)** Lipid metabolism assessed in paired SCAds and rBMAds: (B) FA oxidation capacity (n=5); (C) FA esterification into MG, DG and TG (n=4); (D) *De novo* lipogenesis under basal condition or after insulin stimulation (n=6). (E) Lipolysis activity assessed by measuring glycerol release under basal conditions or after stimulation with isoprenaline (Iso) or forskolin (FK) (n=4). **(F)** Expression of lipolytic enzymes from proteomic dataset. **(G)** Adipokines and cytokines released by paired SCAds and rBMAds (n=7). **(H)** Hematopoietic factors released by paired SCAds and rBMAds (n=9). **(I-K)** CD34+ cells cocultivated (COC) or not (NC) with rBMAds in methylcellulose (n=6). After 14 days, CFU-GM colonies (I) and total cell number (J) were evaluated as well as the number of CFU-G and CFU-M assessed by FACS (K, left and right panel respectively). **(L)** CD34+ cells cocultivated or not with rBMAds for 24h, cell number was evaluated (n=3). The data are the mean ± SEM. ns: non-significant; *p<0.05; **p<0.01; ***p<0.001; ****p<0.0001 according to Student t test (B and I-L) or two-way ANOVA followed by Bonferroni post-test (C - H).

We next assessed, for the first time, the secretory function of a pure population of human rBMAds by measuring adipokines release by ELISA. Previous characterization, at both transcriptomic and protein levels, has only been performed in human BMAds isolated from femoral head wich contained a mixed population with high predominance of cBMAds^4,25^, or in tissue^24,25^. We found human rBMAds release less adiponectin than SCAds (Figure 4G), in line with previous observations in human rBMAT^25^, while earlier reports showed higher secretion from cBMAT-enriched regions in rodents and human^24,25,40^. Moreover, rBMAds released less leptin than SCAds, whereas IL-6 and IL-8 levels were similar (Figure 4G) and IL-1β and TNFα were undetectable. This contrasts with tissue enriched in human cBMAds, which secrete comparable leptin levels and higher amounts of pro-inflammatory cytokines than SCAds^40,41^. Overall, these results show that human rBMAds are actively secretory cells and highlight differences between BMAd subtypes.

Since rBMAds reside in regions of active hematopoiesis, they could support this process. Although human BMAds from femoral head in ceiling culture (i.e. with a dedifferentiated phenotype) have been shown to support hematopoiesis^41^, this has never been investigated using primary human rBMAds directly used after isolation. Moreover, rodent studies have been controversial with studies showing that BMAds can promote stem cell regeneration and hematopoiesis by secreting SCF^11^ whereas opposite results have been reported in areas mainly containing cBMAds^9^. Using multiplex ELISA assay for hematopoietic factors, we showed rBMAds secrete SCF and IL3 known to support hematopoietic stem cells (HSC) ^42,43^, SDF-1 (also known as CXCL12), involved in HSC retention and quiescence^44^ and GM-CSF, involved in the differentiation of granulocyte and myeloid progenitors^43^ (Figure 4H). By contrast, factors involved in the late differentiation of monocytes/macrophages (M-CSF) and granulocytes (G-CSF) were decreased in rBMAds compared to SCAds (Figure 4H). Thus, the secretion profile of rBMAds suggests that they likely function as niche-supporting stromal cells, similar to mesenchymal stromal cells, playing a key role in HSC maintenance, retention, and balanced myeloid support^45^.

According to these results, we investigated whether rBMAds could support normal hematopoiesis using a colony-forming unit (CFU) cell assay with human CD34+ HSC grown in methylcellulose alone or in the presence of isolated rBMAds. After 14 days of culture, we counted the number of CFU-GM and the total cell number after methylcellulose dissociation and we determined the number of CFU-G (CD235-/CD15+) and CFU-M (CD235-/CD15-) by flow cytometry (Figure S4B). The number of CFU-GM colonies (Figure 4I), the total cell number (Figure 4J) as well as the number of CFU-G and CFU-M (Figure 4K) were significantly increased in HSC cells co-cultivated with rBMAds compared to controls. In presence of erythropoietin, a slight increase in the number of BFU-E colonies was observed in the presence of rBMAds (Figure S4C), which was not associated with an increase in total cell number (Figure S4D) nor in the presence of the highest number of CD235+/CD15- cells (Figure S4E). According to the pattern of cytokine secretion of rBMAds, we investigated if rBMAds can stimulate HSC proliferation. These experiments were performed over a short period (24 h) to ensure rBMAds optimal survival in 2D. As shown in Figure 4L, a significant increase in the number of CD34+ cells was observed in the presence of rBMAds compared to cells grown alone. In summary, our data reveal that human rBMAds possess a distinct secretory profile that includes hematopoietic-supporting cytokines such as SCF, GM-CSF, SDF-1, and IL-3 and promote the proliferation and differentiation of human HSCs, particularly toward myeloid lineages, supporting their role as niche-like stromal cells. Although limited by the technical difficulty of cultivating rBMAds in long-term culture assay, these findings provide the first evidence in humans that rBMAds may contribute to active hematopoiesis in the BM microenvironment.

### Concluding remarks

Our study uncovers rBMAds as a previously unrecognize anucleate adipocyte within human BM. These human rBMAds exhibit distinct structural and functional features, separating them from classical adipocytes. Despite lacking nuclei, rBMAds retain mitochondria and ER. Functionally, rBMAds are devoid of lipolytic activity but maintain lipid and glucose metabolism, alongside a secretory profile rich in hematopoietic-supportive cytokines. We provide the first direct evidence in humans that rBMAds promote HSC proliferation and early differentiation *in vitro*. These findings suggest that rBMAds may act as stromal niche cells, directly supporting hematopoiesis. Their anucleate state and phenotype hint at a terminal differentiation process, potentially from cBMAds. The in-depth characterization of rBMAds from human BM provides a foundation for exploring their role in aging, disease and hematologic disorders. Collectively, our findings redefine adipocyte biology within the BM.

## STAR Methods

### Human subcutaneous and bone marrow tissue samples

Paired subcutaneous adipose tissue (SCAT) and bone marrow adipose tissue (BMAT) were harvested from patients undergoing hip surgery at the Orthopedic Surgery and Traumatology Department of the Hospital Pierre Paul Riquet (Toulouse, France) as previously reported^17^. All patients gave informed consent and the samples were obtained according to national ethics committee rules (authorization DC-2019-3784). The samples were immediately placed in 37°C pre-warmed KRBHA (Krebs Ringer Buffer HEPES Albumin buffer) corresponding to Krebs Ringer buffer (Sigma-Aldrich) supplemented with 100mM HEPES (Sigma-Aldrich) and 0.5% FA-free bovine serum albumin (BSA) (Sigma Aldrich) and rapidly transferred to the laboratory (within 1h) where they were processed. BMAT were sorted to obtain “yellow” and “red” BMAT. For all experiments, samples were obtained from 55 men and 54 women (mean age: 64.4 years and mean body mass index (BMI): 24.6 kg/m^2^) (Table S1). Images of femur sections were collected from body donors for scientific research.

### Adipocyte isolation and purification

“Red” (regulatory) BMAds (rBMAds), “yellow” (constitutive) BMAds (cBMAds) and SCAds were isolated as we previous described ^17,18^. Briefly, SCAT and BMAT were rinsed several times with KRBHA. After sorting the “yellow” and “red” BMAT, all ATs were digested with collagenase (250 UI/ mL) diluted in calcium and magnesium free PBS supplemented with 2% FA-free BSA during 15 min for cBMAT and 30 min for rBMAT and SCAT at 37°C under constant shaking. Samples were then filtered with 100 and 200µm cell strainers (for BMAT and SCAT respectively) to remove cellular debris, undigested fragments and bone trabeculae. Floating adipocytes were collected and rinsed with KRBHA several times to obtain pure adipocyte cell suspensions. rBMAds from rBMAT were gently centrifuged for 5 min at 300 g at room temperature (RT) to exclude other contaminants.

### Hematoxylin and eosin staining

Samples of SCAT, cBMAT and rBMAT were fixed in 3.7% paraformaldehyde (PFA) solution from Electron Microscopy Sciences (EMS; Hatfield, PA, USA) for 24 hours prior to paraffin embedding and sectioning (10 µm tissue sections using HistoCore MULTICUT R). Paraffin-embedded tissue sections were incubated at least 30 min at 60°C before being deparaffinized and rehydrated. Slides were incubated in hematoxylin (BioGnost #HEMML-OT-100) for 1 min, and washed in water during 2 min. After 30 seconds in bluing solution (0.028% NH3 in H2O), slides were soaked in 95% ethanol before being incubated in eosin (Sigma #HT110116) for 20 seconds. Slides were successively soaked in 70%, 80%, 90% and 100% ethanol and mounted using cover safe (Microm microtech F/MMCSA100).

### Picrosirius red staining

Pieces of SCAT, rBMAT and cBMAT were fixed overnight in 3.7% paraformaldehyde (PFA) solution from EMS (Hatfield, PA, USA). Samples were rinsed and incubated in Picrosirius Red solution (1 g/L Sirius red in 1.3% picric acid) during 6 hours at RT under gentle shaking. Samples were washed and dehydrated successively in 60%, 80%, 100% ethanol during respectively 10, 10 and 30 minutes. Samples were kept in 100% ethanol at 4°C until image acquisition using a LSM 710 confocal microscopy (Zeiss).

### Confocal microscopy

Pieces of SCAT, cBMAT and rBMAT were fixed overnight in 3.7% PFA solution from EMS (Hatfield, PA, USA). Fixed tissues were blocked and permeabilized for 1 h at RT in calcium and magnesium free PBS supplemented with 3% BSA and 0.1% Triton X-100 (Sigma Aldrich). Tissues were incubated overnight with primary antibody against β-tubulin (Abcam #ab6046) 1:200, vimentin (Abcam #ab195878) 1:500 or perilipin-1 (Origene #AM09128SU-N) 1:5 in PBS. The following day, tissues were rinsed 5 times in PBS and then incubated for 1 hour with secondary antibody αRabbit IgG AlexaFluor 594 (Invitrogen #A11072) 1:500 or αMouse IgG CF647 (Ozyme #BTM20042) 1:1000 depending on the species, together with 5 µg/mL of Bodipy 493/503 (Invitrogen #D3922) and 50 µg/mL of DAPI (Invitrogen #D1306). For F-actin staining, fixed tissues were permeabilized for 1 hour at RT in calcium and magnesium free PBS with 0.1% triton X-100 (Sigma Aldrich) and incubated with rhodamine-coupled phalloidin (Thermofisher #R415) 1:100, 5 µg/mL Bodipy 493/503 (Invitrogen #D3922) and 50 µg/mL of DAPI (Invitrogen #D1306) for 2 hours at RT. Z-stack images were acquired using a LSM 710 confocal microscope with a 40X or 60X objective (Zeiss). Maximum intensity projections were performed using Fiji software (Image J, Bethesda, MD, USA).

SCAds, cBMAds and rBMAds were isolated as described above. Immediately after isolation, primary adipocytes were embedded in a fibrin gel to maintain cellular integrity. Fifty microliters of isolated adipocytes were gently mixed with 50 µL of 18 mg/mL fibrinogen (Sigma Aldrich) solution in 0.9% NaCl buffer and 50 µL of thrombin, 5 units in 50 µL of CaCl2 solution (Sigma-Aldrich) as previously described ^17,18^. After polymerization, gels containing isolated cells were fixed in 3.7% PFA for 1 hour. Cells were permeabilized for 15 minutes at RT in PBS with 0.1% triton X-100 (Sigma Aldrich) and incubated with 5 µg/mL Bodipy 493/503 (Invitrogen #D3922) and 50 µg/mL of Hoescht 33258 (Sigma #94403) or 50 µg/mL of DAPI (Invitrogen #D1306), or CellMask (Thermofisher #C10046) 1:500 or Mitotracker 1µM (Thermofisher #M7512) for 1 hour at RT. For caveolin-1 and PDI, cells were permeabilized for 15 minutes in 0.1% triton X-100 and blocked in calcium and magnesium free PBS supplemented with 3% BSA for 1 hour. Gels were then incubated overnight in primary antibody against caveolin-1 (BD biosciences #610059) 1:200 or PDI (abcam #ab2792) /1:100 in PBS. The following day, gels were rinsed 5 times in PBS and then incubated with secondary antibody αRabbit IgG AlexaFluor 594 (Invitrogen #A11072) 1:500, following by 1 hour of staining, together with 5 µg/mL of Bodipy 493/503 (Invitrogen #D3922) and 50µg/ml of DAPI (Invitrogen #D1306). Z-stack images were acquired using a LSM 710 confocal microscope with a 10X, 40X, 60X objective (Zeiss). Maximum intensity projections were performed Fiji software (Image J, Bethesda, MD, USA).

### Transmission electron microscopy on tissue

Pieces of SCAT, cBMAT and rBMAT were fixed overnight at 4°C with 2.5% glutaraldehyde and 2% PFA from EMS (Hatfield, PA, USA) in Cacodylate buffer (0.1M, pH 7.2) and post-fixed at 4°C with 1% OsO4 and 1.5% K3Fe(CN)6 in the same buffer. Samples were treated for 1 h with 1% aqueous uranyl acetate and were then dehydrated in a graded ethanol series and embedded in EMBed-812 resin (EMS). After 48 h of polymerization at 60°C, ultrathin sections (80 nm thick) were mounted on 75 mesh formvar-carbon coated copper grids. Sections were stained with 2% uranyl acetate (EMS) and 3% Reynolds lead citrate (Chromalys). Grids were examined with a Jeol JEM-1400 transmission electron microscope (JEOL Inc) at 80 kV and images were acquired with a Gatan Orius digital camera (Gatan Inc, Pleasanton, CA, USA).

### Cell counting and size measurement

SCAds and cBMAds were diluted in KRBHA buffer at a ratio 1:10, while rBMAds were diluted at a ratio 1:20 and mounted on a Malassez hemocytometer. For each adipocyte sample, at least 5 images were acquired with Olympus CKX53 inverted microscope with a 10X objective (Zeiss). Cell number and diameter measurement were performed manually using Fiji software (Image J, Bethesda, MD, USA).

### DNA dosage on isolated cells

Aliquots of 50 µL of isolated SCAds, cBMAds or rBMAds were used for DNA dosage. Cells were lysed in TE 1X buffer (10 mM Tris-HCl, 1 mM EDTA, pH 7.5) supplemented with 0.1% Triton X-100, and were sonicated (20 kHz) for 10 seconds. Samples were centrifuged at 10 000 rpm for 5 minutes at 4°C and supernatants were collected. Double strand DNA (dsDNA) was quantified in each sample using the Quant-iT™ PicoGreen® dsDNA Reagent and Kits (Thermofisher #P7589) following the manufacturer’s instruction. For each sample, DNA content was normalized to cell number.

### RNA extraction and dosage on isolated cells

1 mL of Qiazol (Qiagen) was added to 300 µL of isolated SCAds, cBMAds or rBMAds suspension. Samples were mixed thoroughly then centrifuged for 10 minutes at 12000 rpm. 200 µL of chloroform was added to samples and centrifuged for 10 min at 12000 rpm. Aqueous phases (chloroform) were collected and were added to 1:1 v/v 70% ethanol. RNA was purified using the RNeasy® Mini kit columns (Qiagen) following the manufacturer’s instructions. RNA was dosed using a nanodrop (Nanodrop^TM^ 2000, Thermosfisher). For each sample, RNA content was normalized to cell number.

### Lipid content

Tissue (□100mg) or isolated adipocytes (□100µl) were first lysed in 100µl NaOH at 55°C for 15 minutes and 100µl HCl is added. Dole’s Reagent (40:10:1 isopropanol: heptane:H_2_SO_4_ 1N) is then added and the upper phase, containing lipids, was extracted again with heptane, evaporated under a nitrogen stream and weighed as dried lipids.

### Lipidomic analysis

#### Untargeted lipidomic

Lipid extraction was performed with methyl tert-butyl ether (MTBE; Sigma-Aldrich) on 400µl of adipocyte suspension as previously described^17^. One milliliter of the lipid phase was evaporated under a nitrogen stream and dried samples were used for untargeted lipidomic profiling at the Harvard mass spectrometry core as previously described^17^. Lipidomics statistical analysis and visualization were carried out in R working environment and RStudio interface using the default base and stats packages, and tidyverse meta-package^46^. Probabilistic Quotient Normalization was performed using the lipidr package^47^. Only lipid species detected in at least three out of the five patients for every adipocyte type were considered. Lipid species belonging to the same class were summed to obtain relative abundances. Percentage of each species were defined considering the total relative abundance within each sample.

#### Total FA content

Analysis of FA composition was performed at the metabolomics, lipidomics & fluxomics facility of Toulouse (MetaToul), after lipid extraction with MTBE. Ten microliter of the lipid phase was hydrolyzed in KOH (0.5 M in methanol) at 55°C for 30 min, and transmethylated in boron trifluoride methanol solution 14% (Sigma) and at 80°C for 1h. After addition of H20 to the crude, Fatty Acid Methyl Esters (FAMEs) were extracted with heptane, centrifugated 1min at 2500rpm and evaporated to dryness and dissolved in 200µL ethyl acetate. FAME (1 µL) were analyzed by gas chromatography equipped with a FID (Lillington, J.M., Trafford, D.J., and Makin, H.L. 1981, Clin Chim Acta 111:91-98) on a Trace 1300 Thermo scientific system using a Famewax RESTEK fused silica capillary columns (30 m x 0.32 mm i.d, 0.25 µm film thickness). Finally, peak detection, integration and quantitative analysis were done using MassHunter Quantitative analysis software (Agilent Technologies). Unsaturation index was calculated by dividing MUFA (C16:1 + C18:1) by SFA (C16:0+C18:0) as previously reported^48^

#### Plasma membrane lipid quantification

The phospholipid an sphingolipids analysis was performed at the metabolomics, lipidomics & fluxomics facility of Toulouse (MetaToul). Lipid extraction was done on 400µl of adipocytes and adapted from Bligh and Dyer^49^ in dichloromethane/methanol (2% acetic acid) / water (2,5 :2,5 :2 v/v/v), in the presence of the internals standards (Cer d18:1/15:0; PE 12:0/12:0; PC 13:0/13:0; SM d18:1/12:0; PI 16:0/17:0; PS 12:0/12:0). The solution was centrifuged at 1500 rpm for 3 min. The organic phase was collected and dried under nitrogen, then dissolved in 100µL of dichloromethane. For positive analysis, samples were diluted: 10 µL of final extract with 50 µL of isopropanol and for negative analysis: 20 µL of final extract with 40 µL of isopropanol. The lipid extract was then stored at −20°C prior analysis. Lipid extracts were analyzed using an Agilent 1290 UPLC system coupled to a G6460 triple quadripole mass spectrometer (Agilent Technologies) and a Kinetex HILIC column (Phenomenex, 50 x 4.6 mm, 2.6 µm) was used for LC (liquid chromatography) separations. Electrospray ionization was performed in positive mode for Cer, PE, PC and SM analysis and in negative mode for PI and PS analysis. Finally, peak detection, integration and quantitative analysis were done using MassHunter Quantitative analysis software (Agilent Technologies). For each sample, lipid content was normalized to cell surface evaluated after quantification of adipocyte number and size for each sample (see detail above).

### Proteomic analysis

#### Sample preparation for proteomic analysis

Three paired samples, injected twice each, were used (two men and one woman, mean age: 59 years; mean BMI: 24.5 kg/m^2^). For each sample, 5µg of protein in 50 mM triethylammonium bicarbonate (TEAB), 5% SDS are reduced and alkylated in 10 mM Tris(2-carboxyethyl)phosphine hydrochloride, 40 mM Chloroacetamide, 40 mM KOH for 5 min at 95°C. After acidification with 2.5% of phosphoric acid, proteins were precipitated with 7 vol of 90% MeOH-10 % TEAB and then deposited on S-Trap Micro column. After 6 washes with 1 volume of 90% MeOH-10 % TEAB, trypsin digestion was performed on the column with 1µg of trypsin (modified sequencing grade trypsin (Promega)) for 5 µg of proteins overnight at 37°C. The resulting peptides were eluted by several steps: 50 mM ammonium bicarbonate followed by 2% formic acid and then with 50% acetonitrile (ACN), 0.2% formic acid. The pooled peptidic fractions were dried under speed-vacuum and resuspended with 2% ACN and 0.05% trifluoroacetic acid (TFA) for MS analysis.

#### LC-MS/MS analysis

Peptides were analyzed by nanoLC-MS/MS using an UltiMate 3000 RSLCnano system coupled to a Orbitrap Fusion mass spectrometer (Thermo Fisher Scientific). Five µL of each sample was loaded onto a C-18 precolumn (300 µm ID x 5 mm, Thermo Fisher) in a solvent consisting of 2% ACN and 0.05% TFA. After 5 min of desalting, the precolumn was switched online with the analytical C-18 column (75 µm ID x 15 cm, Reprosil C18) equilibrated in 95% solvent A (5% ACN, 0.2% formic acid) and 5% solvent B (80% ACN, 0.2% formic acid). Peptides were eluted using two gradients: a 5 to 25% gradient over 75 min and a 25 to 50% over 30 min of solvent B at a flow rate of 300 nL/min. The Orbitrap Fusion was operated in a data-dependent acquisition mode, on the 400-1500 m/z range with the resolution of 120000 in MS and 30000 in MSMS after HCD fragmentation. Dynamic exclusion was employed within 60 seconds and duplicate technical LC-MS measurements were performed for each sample.

#### Data processing

Raw MS files were processed with Mascot 2.8.3 software for database search and Proline 2.1 for label-free quantitative analysis^50^. Data were searched against human sequences of the database (release UniProt SwissProt 2022; 20,376 reviewed sequences). Carbamidomethylation of cysteines was set as a fixed modification, whereas oxidation of methionine and protein N-terminal acetylation were set as variable modifications. Up to two missed trypsin cleavage sites were allowed. The mass tolerance was set to 5 ppm for the precursor and to 0.8 Da in tandem MS mode. Minimum peptide length was set to 7 amino-acids, and identification results were further validated in Proline by the target decoy approach using a reverse database at both a peptide spectrum-match and protein false-discovery rate (FDR) of 1%. Label free relative quantification was performed with Proline 2.1 using default parameter set.

Only proteins identified in at least 2 out of 3 samples in the same location (i.e. rBMAds or SCAds) were considered to be robustly detected, and quantities were log2-transformed and median-centered. The statistical analysis of differentially expressed proteins was performed with the Limma statistical test with a paired design. Protein expression was considered significantly different between conditions when i) protein expression ratio was more than 2-fold and ii) the p-value was less than 0.05. Perseus 2.0.10.0 was used to perform principal component analysis on all proteins after replacement of missing values by random numbers drawn from a normal distribution of 1.8 standard deviation down shift and with a width of 0.3 of each sample. Pathway enrichment analysis was performed with Gene Analytics (GeneCards). Heatmaps was made with the gplots package (v3.0.1.1).

### Colony forming cell assay

Three hundred human CD34+ progenitor cells (ABCell-Bio) resuspended in IMDM media (Thermofisher #12440053) were cultured alone or in co-culture with 50 µL of rBMAds in MethoCult□□ (Stem Cell technologies #84534) supplemented or not with 3U of human erythropoietin (Tebu bio #11965-1) in a total final volume of 1 mL per 6-plate-well. After 14 days, whole wells were scanned using the EVOS™ M700 Imaging System for colony count. Each of the well from the colony forming cell assay were resuspended in magnesium and calcium free PBS, washed and centrifuged twice at 1400 rpm for 7 minutes at RT. After counting the total number of cells using a Malassez counting chamber, cells were incubated in anti-CD235a-APC-Vio770 (Miltenyi Biotech #130-100-268) 1:5, anti-CD14-PerCP-Vio700 (Miltenyi Biotech #130-110-523) 1:20 and anti-CD15-eFluor™ 450 (Fisher scientific #15559636) 1:20 antibodies for 30 minutes on ice. Cells were washed and analyzed using Fortessa X20 flow cytometer using DiVa software (BD Biosciences) and analyzed using FlowJo 10.1 software (Tree Star).

### Quantification of secreted adipokines

Two hundred microliters of isolated SCAds and rBMAds were incubated in 500 µL of RPMI 5mM glucose (without serum) during 3 hours at 37°C in 5% CO_2_. Conditioned media (CM) were cautiously collected and centrifuged at 12000 rpm for 5 minutes to eliminate cell debris. CM were then divided into aliquots and stored at −20°C for further ELISA or Luminex assay. Fifty microliters of each CM were used to quantify adiponectin, leptin, IL-6, IL-8, IL-1β and TNF-α using Human DuoSet ELISA kits from R&D Systems according to the manufacturer’s instruction. Twenty-five microliters of each CM were used to quantify SDF-1, SCF and TPO using HCYP2MAG-62K Luminex plate (Merck Millipore) and G-CSF, GM-CSF, Flt-3 ligand, IL-3, IL-4, IL-5, M-CSF using HCYTA-60K Luminex plate (Merck Millipore) according to the manufacturer’s instruction. Missing values were imputed using half of the lowest value from the standard curve for each specific cytokine.

### Lipolysis assay

Isolated adipocytes (100 µL) or AT explants (∼100 mg) were incubated in 200 µL KRBHA with or without isoprenaline 1µM (Sigma Aldrich) or forskolin 10µM (Sigma Aldrich) to evaluate stimulated and basal lipolysis. After 3 hours incubation at 37°C under gentle shaking, media were collected and stored at −20°C to measure glycerol release using commercial kits (Sigma F6428). Lipolysis was normalized by the volume of adipocyte suspension or the weight of AT.

### Measure of *de novo* lipogenesis

One hundred microliters of SCAds and rBMAds were incubated 2 hours at 37°C in 5% CO_2_ in 200µl of RPMI glucose-free RPMI medium without FBS. Then, adipocyte suspensions were pre-incubated during 15 min in a 5mM glucose medium supplemented with or without 100nM of insulin (Sigma Aldrich, I2643), followed by incubation with 0.5µCi/mL of [U-^14^C]-D-glucose, (Perkin Elmer, NEC042X250UC). After 2 hours, adipocytes were rinsed 2 times with PBS. Lipids were extracted after adding Dole solution (40:10:1 isopropanol/heptane/1N H_2_SO_4_) and heptane solution at equal volume. Lipid organic phase was transferred to scintillation tube, evaporated and the lipid residue was weighed for data normalization. Cell-associated radioactivity was evaluated by scintillation counting (CytoScint, MP Biomedicals).

### Measure of FA oxidation (FAO) and esterification

One hundred microliters of SCAds and rBMAds were incubated 2 hours at 37°C in 5% CO_2_ in 200µl of RPMI glucose-free RPMI medium without FBS. Then, adipocyte suspensions were incubated with 400 µL warmed (37°C) modified Krebs Ringer buffer pH 7.4 containing 5mM glucose, 1mM palmitate complexed with 1.3% FA-free BSA and 0.5µCi/mL ^14^C-palmitate (Perkin Elmer, NEC075H250UC) for 3 hours as we previously described^51^. To evaluate FA esterification, adipocytes are washed 2 times with PBS and lipids were extracted after adding Dole solution (40:10:1 isopropanol/heptane/1N H_2_SO_4_) and heptane solution at equal volume. Lipid organic phase was transferred to a glass vial, evaporated and the lipid residue was weighed for data normalization. FA esterification was quantified after separation of TG, DG, MG and FA by thin-layer chromatography using hexane-diethyl ether-formic acid (55:45:1 v/v/v) as the eluent^52^. For quantification, silica gel zones at the Rf values corresponding to authentic chemical standards were scraped and radioactivity was measured using a β-counter, as previously described^53^. To evaluate FAO, 1mL of incubation media was transferred to a glass tube and a microtube containing 300µL of NaOH 1N was placed in the vial to capture the ^14^CO_2_. The buffer was acidified with 1mL of 1M H_2_SO_4_ and the vial was sealed. After 24h, the radioactivity was counted (CytoScint, MP Biomedicals). FAO was normalized by the volume of adipocyte suspension.

### Statistical analysis

Statistical analyses were performed using Prism v10 (GraphPad Software). Two group comparisons were performed using paired Student’s t test and multiple comparisons were performed by one-WAY or two-way ANOVA followed by Bonferroni post-test. All the statistical details (statistical tests used, n numbers and data dispersion) can be found in the figure legend.

## Supplementary Figure legends

**Figure S1. Distinct structural and cellular features of rBMAT compared to cBMAT and SCAT**

**(A)** Representative images showing sample heterogeneity from 6 patients with different amount of BMAT. Gender (women or men), age and body mass index (BMI) is mentioned.

**(B)** Longitudinal cross section of human femur from 2 patients.

**(C)** Total lipid content in paired adipose tissues (n=7). Bar plots represent mean ± SEM. Ns: non-significant; *p<0.05; **p<0.01 according to One-way ANOVA followed by Bonferroni post-test.

**(D)** HE staining of two punch biopsies of BMAT harvested from the longitudinal cross section of one human femur.

**Figure S2. rBMAds are devoid of nucleus but retain metabolic organelles**

**(A)** Primary SCAds, cBMAds and rBMAds stained with Bodipy 493/503 and DAPI. Representative maximum intensity projection of z-stack images is shown. Scale bar, 50µm.

**Figure S3: Lipidomic and proteomic characterization define rBMAds identity in humans**

**(A)** Principal component analysis performed on the lipidomic dataset from paired adipocytes (n=5).

**(B)** FA species in paired isolated adipocytes (n=5). The data are the mean ± SEM. Ns: non-significant; *p<0.05; according to two-way ANOVA followed by Bonferroni post-test.

**(C)** Pathway enrichment analysis of biological process gene ontology terms performed with Gene Analytics. The top 20 pathways upregulated or downregulated in rBMAds compared to SCAds are presented.

**Figure S4. rBMAds exhibit distinct metabolic and endocrine characteristics and support normal granulo-monocytic hematopoiesis in vitro**

**(A)** Lipolysis activity of paired SCAT and rBMAT assessed by measuring glycerol release under basal conditions or after stimulation with isoprenaline (Iso) or forskolin (FK) (n=4).

**(B)** Experimental workflow designed to assess the effects of rBMAds on hematopoiesis.

**(C-E)** CD34+ cells cocultivated (COC) or not (NC) with rBMAds in methylcellulose containing erythropoietin (n=6). After 14 days, BFU-E colonies (C) and total cell number (D) were evaluated as well as the number of BFU-E assessed by FACS (E).

The data are the mean ± SEM. Ns: non-significant; *p<0.05; ***p<0.001 according to Student t test (C-E) or two-way ANOVA followed by Bonferroni post-test (A).

Table S1: Characteristics of the patients

## Acknowledgements

This work was supported by INCa PLBio (2020-28), *Ligue Nationale contre le Cancer* (équipe labélisée) and AFERO (*Association Française d’étude et de recherche sur les obésités*) for running costs. David Estève received a post-doctoral fellowship from the *Fondation pour La Recherche Médicale* (SPF201809007124). Sauyeun Shin and Philippe de Medina received a post-doctoral fellowship from INCa PLBio (2020-28). Marine Hernandez received a PhD grant from the French Ministry of Research and by the *Ligue Nationale contre le Cancer*.

We acknowledge the TRI (*Toulouse Réseau Imagerie*) imaging facility, member of the national infrastructure France-BioImaging supported by the French National Research Agency (ANR-10-INBS-04). We acknowledge the METi imaging facility (Genotoul-TRI), member of the national infrastructure France-BioImaging supported by the French National Research Agency (ANR-24-INBS-0005 FBI BIOGEN). This study benefited from the national proteomic infrastructure funded by the French Ministry of Research with *the Investissement d’Avenir Infrastructures Nationales en Biologie et Santé* program (ProFI, Proteomics French Infrastructure project, ANR-10-INBS-08 and ANR-24-INBS-0015). Lipidomic analysis were performed at MetaToul (Toulouse metabolomics & fluxomics facilities, www.mth-metatoul.com on the site dedicated to Lipids Analysis Inserm I2MC) and was supported by the MetaboHUB infrastructure funded by the Agence Nationale de la Recherche under the France 2030 program (MetaboHUB ANR-11-INBS-0010; MetEx+ ANR-21-ESRE-0035; MetaboHUB (JVCE) ANR-24-INBS-0012). We thank the Genotoul Anexplo facility for luminex analysis and the Mass Spectrometry Facility of the Beth Israel Deaconess Medical Center (Boston, USA) for global lipidomic analysis. We thank Life Science Editors for editorial assistance.

## Author contributions

NR set up the conditions for harvesting BMAT and SCAT in close collaboration with CA and CM and supervised the samples collection. MR supervised collection and analysis of femur from body donors for scientific research. SS, MH, DE, CA and MM handled the AT samples and isolated adipocytes. DE, SS, MH and SD performed the immunofluorescence experiments and the transmission electron microscopy (with the help of the METi platform) and as well as image data analysis. SS, MH and CA performed sample preparation for proteomic and lipidomic studies and the lipolysis experiments. MDP and EMB performed the proteomics studies and statistical analysis under the supervision of OS and CA performed pathway analysis. SS, CA, OD and JBM conducted lipidomic analysis and DR performed statistical analysis. MH and CA performed metabolic experiments and PDM, MP, SSP performed FA esterification quantification. SS performed hematopoiesis analysis and CD and PV performed multiplex analysis of hematopoietic factors. CA and CM conceived the idea for this project and wrote the manuscript with significant inputs from all authors. CA and CM supervised the study.

## Data availability

The MS proteomics data are available via ProteomeXchange with identifier PXD071439.

## Declaration of interests

The authors declare they have no conflict of interest

## Notes

### Competing Interest Statement

The authors have declared no competing interest.

