## Supplementary figures and images for "Red bone marrow hosts metabolically active anucleate adipocytes that support hematopoiesis"

### Supplemental figures

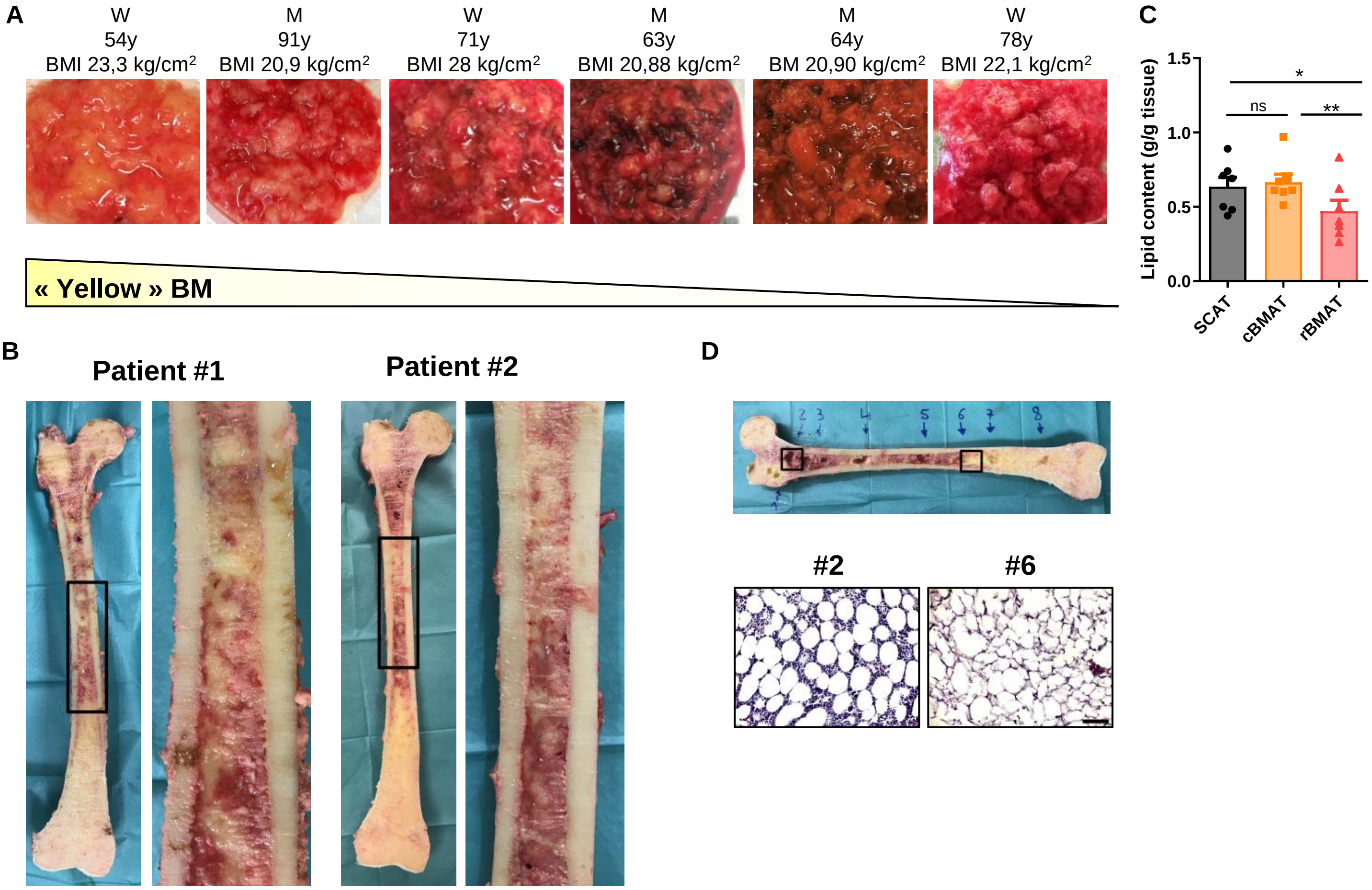

#### Figure S2

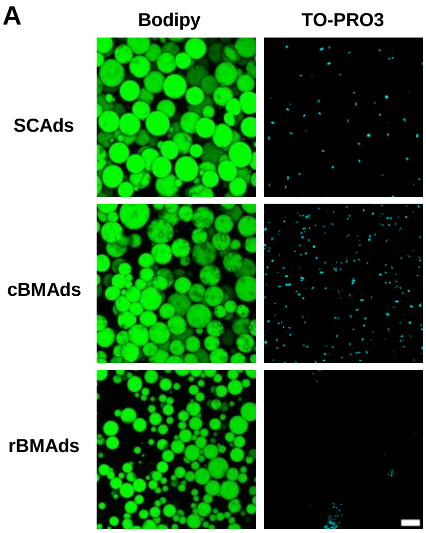

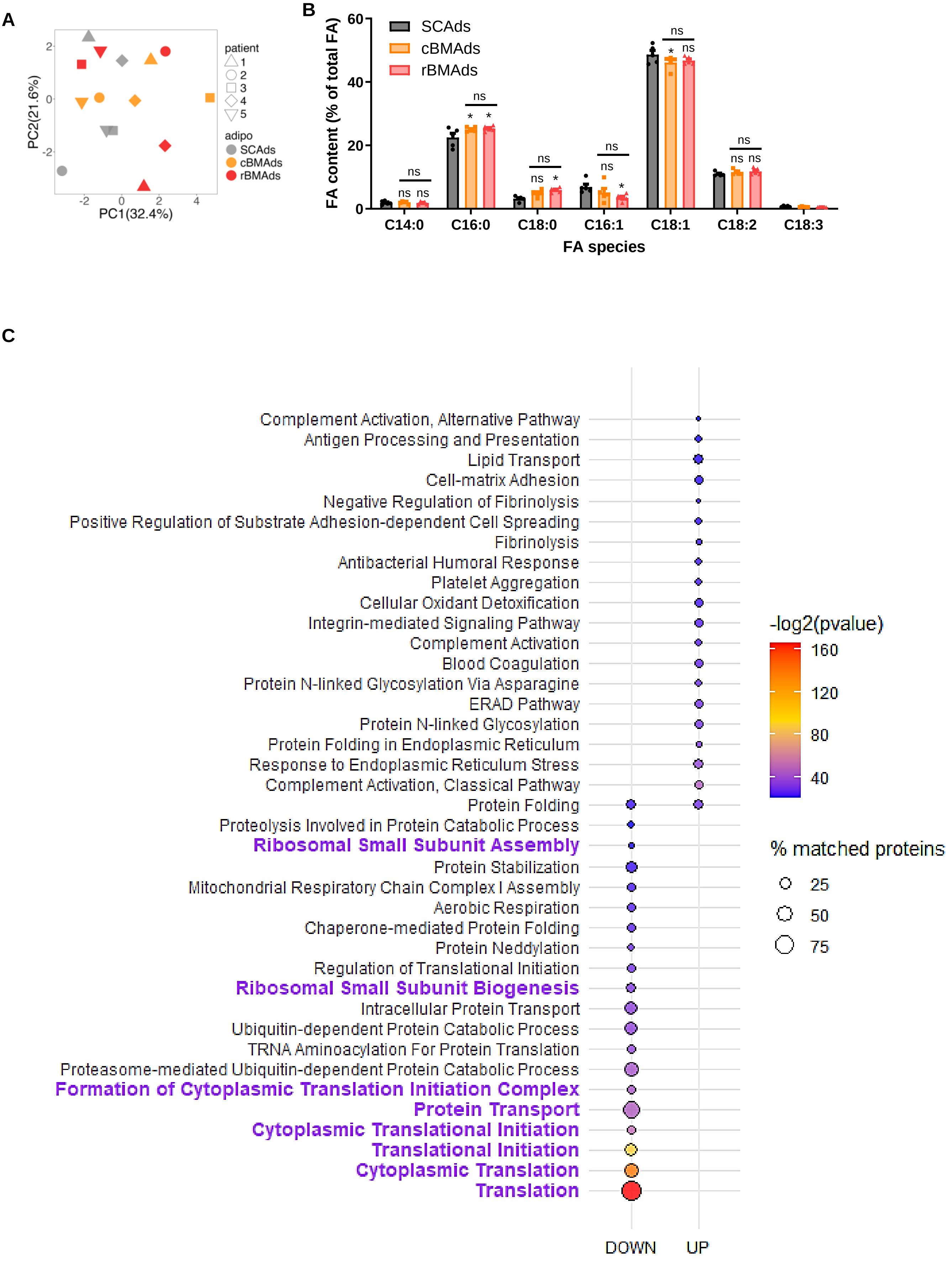

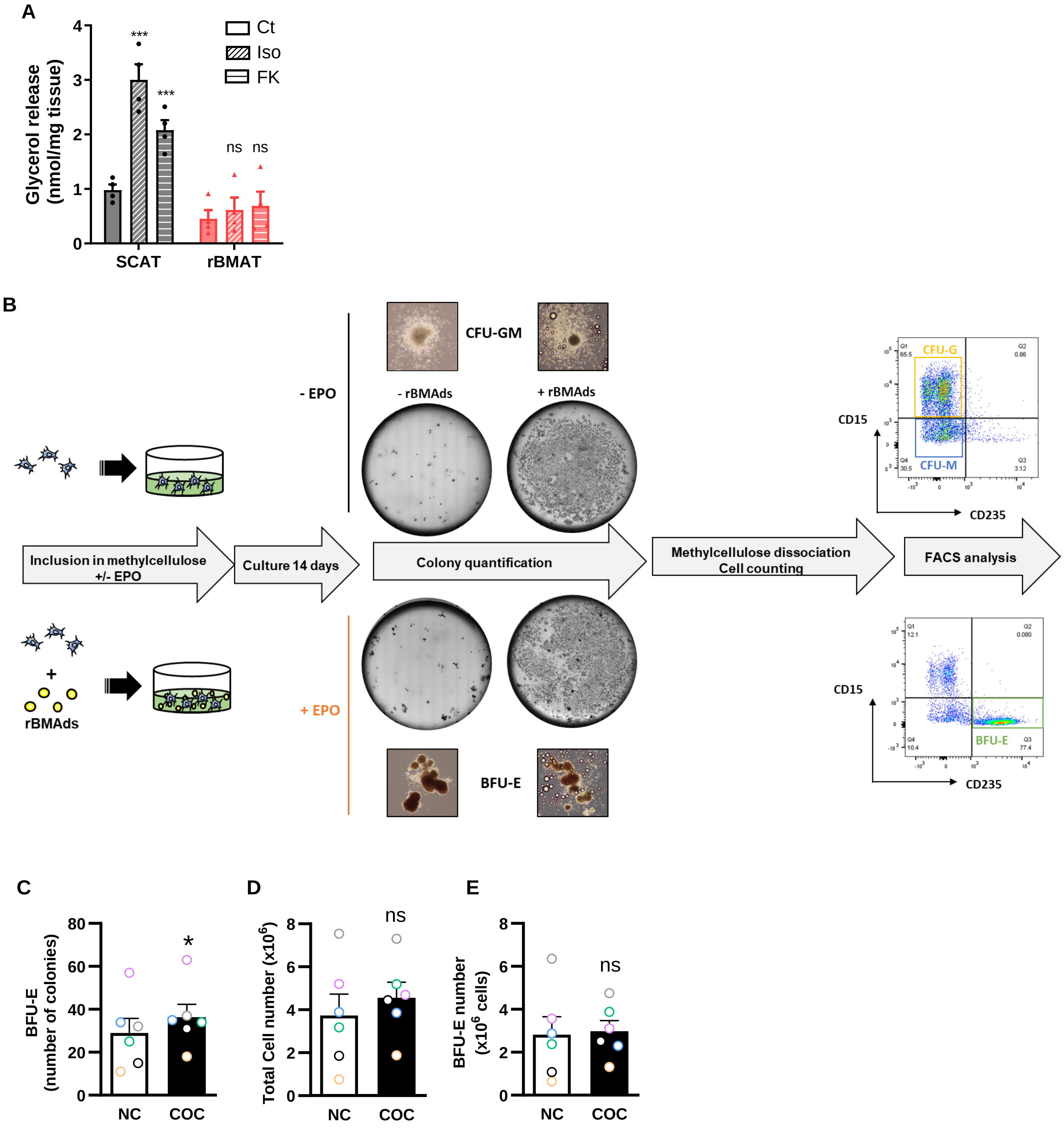
