## Supplemental table for "Red bone marrow hosts metabolically active anucleate adipocytes that support hematopoiesis"

| Gender | Age | BMI |
|---|---|---|
| W | 74 | 25,4 |
| W | 82 | 28,7 |
| W | 88 | 22,8 |
| W | 54 | 28,5 |
| W | 77 | 27,9 |
| W | 74 | 24,0 |
| W | 71 | 29,8 |
| W | 77 | 27,7 |
| W | 69 | 26,0 |
| W | 83 | 30,2 |
| W | 82 | 29,0 |
| W | 57 | 1,7 |
| W | 50 | 23,1 |
| W | 77 | 1,6 |
| W | 74 | 1,6 |
| W | 58 | 23,5 |
| W | 40 | 23,0 |
| W | 77 | 19,5 |
| W | 49 | 26,6 |
| W | 70 | 21,1 |
| W | 59 | 34,6 |
| W | 67 | 22,4 |
| W | 61 | 27,4 |
| W | 68 | 21,6 |
| W | 60 | 28,7 |
| W | 75 | 27,9 |
| W | 50 | 28,3 |
| W | 72 | 23,5 |
| W | 66 | 21,6 |
| W | 42 | 20,2 |
| W | 58 | 22,7 |
| W | 75 | 22,9 |
| W | 51 | 24,6 |
| W | 76 | 23,0 |
| W | 63 | 30,5 |
| W | 59 | 23,2 |
| W | 74 | 25,0 |
| W | 58 | 25,3 |
| W | 80 | 20,5 |
| W | 64 | 23,7 |
| W | 87 | 22,9 |
| W | 34 | 23,8 |
| W | 77 | 21,6 |
| W | 71 | 25,3 |
| W | 68 | 19,8 |
| W | 58 | 32,9 |
| W | 63 | 18,4 |
| W | 57 | 22,5 |
| W | 78 | 22,1 |
| W | 71 | 19,5 |
| W | 45 | 20,1 |
| W | 62 | 19,4 |
| W | 71 | 28,1 |

| | | |
|---|---|---|
| W | 54 | 23,3 |
| M | 78 | 31,0 |
| M | 71 | 25,3 |
| M | 47 | 30,1 |
| M | 66 | 27,7 |
| M | 56 | 28,0 |
| M | 57 | 23,0 |
| M | 40 | 30,0 |
| M | 65 | 29,7 |
| M | 82 | 26,2 |
| M | 72 | 25,2 |
| M | 55 | 24,2 |
| M | 77 | 22,6 |
| M | 30 | 23,8 |
| M | 61 | 28,0 |
| M | 76 | 33,2 |
| M | 63 | 27,8 |
| M | 21 | 16,0 |
| M | 74 | 26,3 |
| M | 61 | 28,0 |
| M | 66 | 21,9 |
| M | 72 | 23,7 |
| M | 79 | 28,7 |
| M | 73 | 28,0 |
| M | 69 | 23,7 |
| M | 67 | 26,9 |
| M | 36 | 25,4 |
| M | 51 | 28,4 |
| M | 67 | 26,8 |
| M | 52 | 28,4 |
| M | 79 | 24,8 |
| M | 36 | 22,9 |
| M | 67 | 25,6 |
| M | 75 | 25,9 |
| M | 72 | 30,8 |
| M | 71 | 23,3 |
| M | 72 | 23,3 |
| M | 39 | 22,8 |
| M | 59 | 27,0 |
| M | 70 | 27,7 |
| M | 51 | 28,7 |
| M | 57 | 24,8 |
| M | 70 | 26,4 |
| M | 88 | 25,5 |
| M | 83 | 23,4 |
| M | 35 | 22,7 |
| M | 48 | 26,8 |
| M | 63 | 20,9 |
| M | 70 | 24,2 |
| M | 69 | 25,1 |
| M | 61 | 22,8 |
| M | 72 | 32,5 |
| M | 77 | 26,6 |
| M | 83 | 25,4 |
| M | 54 | 22,7 |
| M | 64 | 20,9 |
